# Long-read detection of transposable element mobilization in the soma of hypomethylated *Arabidopsis thaliana* individuals

**DOI:** 10.1101/2025.02.07.637047

**Authors:** Andrea Movilli, Svitlana Sushko, Fernando A. Rabanal, Detlef Weigel

## Abstract

**Background:** Because transposable elements (TEs) can cause heritable genetic changes, past work on TE mobility in *Arabidopsis thaliana* has mostly focused on new TE insertions in the germline of hypomethylated plants. It is, however, well-known that TEs can also be active in the soma, although the high-confidence detection of somatic events has been challenging. Here, we leveraged the high accuracy of PacBio HiFi long reads to evaluate the somatic mobility of TEs in individuals of an *A. thaliana* non-reference strain lacking activity of METHYLTRANSFERASE1 (MET1), a major component of the DNA methylation maintenance machinery. Most somatically mobile families coincided with those found in germline studies of hypomethylated genotypes, although the exact TE copies differed. We also discovered mobile elements that had been missed by standard TE annotation methods. Somatic TE activity was variable among individual plants, but also within TE families. Finally, our approach pointed to the possible involvement of alternative transposition as a cause for somatic hypermutability in a region that contains two closely spaced VANDAL21 elements. We conclude that long-read sequencing can reveal widespread TE transposition in the soma of *A. thaliana* hypomethylated mutants. Assessing somatic instead of germline mobilization is a fast and reliable method to investigate different aspects of TE mobility at the single plant level.

## Introduction

Within the dynamics of ongoing evolutionary genomic conflict, TEs, being ‘genomic parasites’, should be silent – or at least minimally disruptive – in both somatic and germline cells of the host (Haig 2016; Chang *et al*. 2022). In animals, the germline is frequently specified early during development, thereby minimizing the number of cell divisions in the germline and also the opportunity for TEs to proliferate. In contrast, the separation between soma and germline is not as clear in plants, and the later and more plastic determination of the plant germline (Berger and Twell 2011) allows for rare mutagenic events in the soma, including TE insertions, having a non-zero chance to be passed on to the next generation (Bourque *et al*. 2018).

Germline mobilization of TEs in *Arabidopsis thaliana* has been studied both in epigenetic mutants (Singer *et al*. 2001; Miura *et al*. 2001; Kato *et al*. 2004; Tsukahara *et al*. 2009; Baduel *et al*. 2021; Zhang *et al*. 2023; Vendrell-Mir *et al*. 2024) and in epigenetic recombinant inbred lines (epiRILs) *(Reinders et al. 2009; Mirouze et al. 2009; Catoni et al. 2019; Quadrana et al. 2019)*, but there has been much less focus on assessing somatic TE mobilization. In addition to direct detection of mobile TEs by whole-genome sequencing, TE mobility can be assessed by isolation of virus-like particles (VLP) (Lee *et al*. 2020) or extrachromosomal circular DNA molecules (eccDNAs) (Merkulov *et al*. 2023; Zhang *et al*. 2023).

A plethora of enzymes belonging to different pathways synergistically and with different degrees of redundancy establish TE repression in plants, through DNA methylation in all cytosine contexts, deposition of histone variants, and post-translational modification of histones (Lippman *et al*. 2003; Mirouze *et al*. 2009; Tsukahara *et al*. 2009; Deleris *et al*. 2012; Stroud *et al*. 2013; Creasey *et al*. 2014; Zhang *et al*. 2018; Osakabe *et al*. 2021; Déléris *et al*. 2021; To *et al*. 2022; Oberlin *et al*. 2022; Sasaki *et al*. 2022b; Hure *et al*. 2025). The chromatin remodeller DECREASE IN DNA METHYLATION 1 (DDM1), which defines heterochromatin by depositing the H2A.W histone variant, mediates methyltransferases accessibility, ultimately establishing methylation in all cytosine contexts, CG, CHG and CHH, in *A. thaliana* (Osakabe *et al*. 2021, 2024; Lee *et al*. 2023). On the other hand, METHYLTRANSFERASE1 (MET1) – a DNA methyltransferase 1 (DNMT1) homolog – is directly responsible for only CG methylation maintenance, and its disruption leads to massive gene expression changes and the stochastic generation of epialleles (Mathieu *et al*. 2007; Rigal *et al*. 2016; Srikant *et al*. 2022). *MET1* activity can be further modulated by the genetic background and, as a consequence, mutations of *MET1* in different accessions cause phenotypes that range from near-wild-type to dwarves with reduced fertility and compromised survival (Srikant *et al*. 2022). Although much of the phenotypic variation due to loss of *MET1* activity is explained by the generation of epialleles (Reinders *et al*. 2009), germline insertions in a range of genes have also been shown to be a cause for morphological defects in later-generation *met1* mutants and *met1*-derived epiRILs (Mirouze *et al*. 2009; Quadrana *et al*. 2019).

Despite there being hundreds of TE families, in *met1* and *met1*-derived epiRILs only a few TE families of both class I (“copy-and-paste” retrotransposons) and class II (“cut-and-paste” DNA transposons) become active. These are the ATCOPIA93 ÉVADÉ (EVD), CACTA/EnSpm (ATENSPM3) and Pack-TYPE CACTA elements (Kato *et al*. 2004; Reinders *et al*. 2009; Mirouze *et al*. 2009; Marí-Ordóñez *et al*. 2013; Catoni *et al*. 2019). Compared to inactivation of *MET1*, disruption of *DDM1* triggers much more extensive TE mobilization (Tsukahara *et al*. 2009). In addition the three families that become active in *met1* mutants, several Ty1/Copia families (ATCOPIA93, -13, -51, -63, -21, -31, ATRE1), several Ty3 families (ATGP3-1, -2, ATGP2N), CACTA/EnSpm elements (ATENSPM3), and MuDR elements (VANDAL21, ATMU1/5) transpose in *ddm1* mutants and *ddm1*-derived epiRILs (Singer *et al*. 2001; Miura *et al*. 2001; Tsukahara *et al*. 2009; Fu *et al*. 2013; Quadrana *et al*. 2019). Some of these TEs have also been found to mobilize in mutants defective in the RNA-dependent DNA methylation pathway (RdDM), which is responsible for *de novo* DNA methylation of TEs (Baduel *et al*. 2021; Zhang *et al*. 2023), and in plants lacking all DNA methyltransferases (He *et al*. 2022).

A limitation of studying only *de novo* TE insertions that have survived selection during the haploid phase of the germline is that not all insertions can be passed on to the next generation. Biases in the detection of TE insertions may in turn result in important aspects of TE biology being overlooked. Moreover, the study of somatic TE mobilization is worthwhile in its own right given that it can have phenotypic consequences, which has been known since McClintock linked revertant maize kernel colors to “controlling element” excision (McClintock 1950; Döring and Starlinger 1984). More recently, somatic TE activity has been suggested to underlie neuronal heterogeneity in diverse animals including humans, contributing to aging and disease pathophysiology (Muotri *et al*. 2005; Kazazian 2011; Baillie *et al*. 2011; Li *et al*. 2013; Gerdes *et al*. 2022). Unfortunately, because somatic events are only found in a subset of cells, their detection by short-read sequencing has been challenging (Cameron *et al*. 2019). Here, we assess whether one can study even very rare somatic TE transposition events with highly-accurate single-molecule long-read sequencing, similar to approaches developed elsewhere (Siudeja *et al*. 2021; D’Costa and Simpson 2023). Because with this method each molecule is read multiple times, the per-base error rates are low, potentially allowing for very rare events that are present only on a single chromosome in a single cell of the sample to be detected. While this approach would have limitations for estimating single-base mutations, the longer sequences of TEs make a low per-base error rate acceptable. We demonstrate the potential of our approach in methylation-defective *A. thaliana* mutants by measuring somatic TE insertions and excisions in 10 first-generation *met1* siblings in a non-reference accession. We describe extensive interindividual, within- and between-family variation in TE mobilization and uncover family-specific biases. Finally, we identify a case of sequence hypermutability, probably driven by alternative transposition, in a region where two TEs are spaced closely apart.

## Results

### Detection of somatic TE mobilization events in *met1* single individuals

To investigate the extent to which TEs mobilize somatically in *A. thaliana*, we made use of *met1* mutants, which are known to have increased TE activity (Reinders *et al*. 2009). First, we *de novo* assembled the genome of the non-reference accession Tsu-0 (1001 Genomes Project ID: 7373) with PacBio HiFi reads, followed by annotation of protein-coding genes and TEs (Supplementary Table 1, Supplementary Fig. 1,2,3). Next, we sequenced, also with PacBio HiFi reads, 10 first-generation Tsu-0 *met1* individual plants (Line 2, (Srikant *et al*. 2022)) (Supplementary Fig. 4), obtaining on average a genome-wide coverage of 38x (Supplementary Fig. 3).

To detect rare somatic transposition events, we mapped reads with Winnowmap2 (Jain *et al*. 2022) from *met1* individuals, and, as a control, from wild type, to the Tsu-0 genome. We identified candidate reads spanning new insertions of annotated TEs because they either (1) were flagged as “chimeric”, *i.e.*, had one or more discontiguous supplementary alignments (SA tag) in addition to a primary alignment, or (2) showed ≥500 bp insertions or deletions relative to the target genome, as reported in their CIGAR strings (Fig. 1a). CIGAR strings have been used before to assess structural variations (SVs) in *A. thaliana* mutants (Zhang *et al*. 2023). If either (1) one alignment of a “chimeric” read, or (2) the insertion/deletion section of a read mapped precisely to an annotated TE, we considered this *bona fide* evidence for a mobilization event (Fig. 1a, reads 2,3,4). To reduce false positives, we visually inspected the genome alignments of reads describing candidate insertion/deletion events (Additional file 4). First, we discarded all events at one locus, because an insertion of organellar DNA into the nuclear genome had been erroneously annotated as a TE (Additional file 4). After visual inspection of candidate insertion events, we discarded 26 events, of which 15 were apparent rearrangements overlapping with a region that appeared to be highly variable in our *met1* mutants (Additional file 4), which we decided to investigate further (see below). Of 23 candidate excisions events, we confirmed eight, while of the other thirteen, seven overlapped with satellite DNA rearrangements and five with the above-mentioned hypermutable region, and were therefore excluded (Additional file 4).

**Figure 1.**
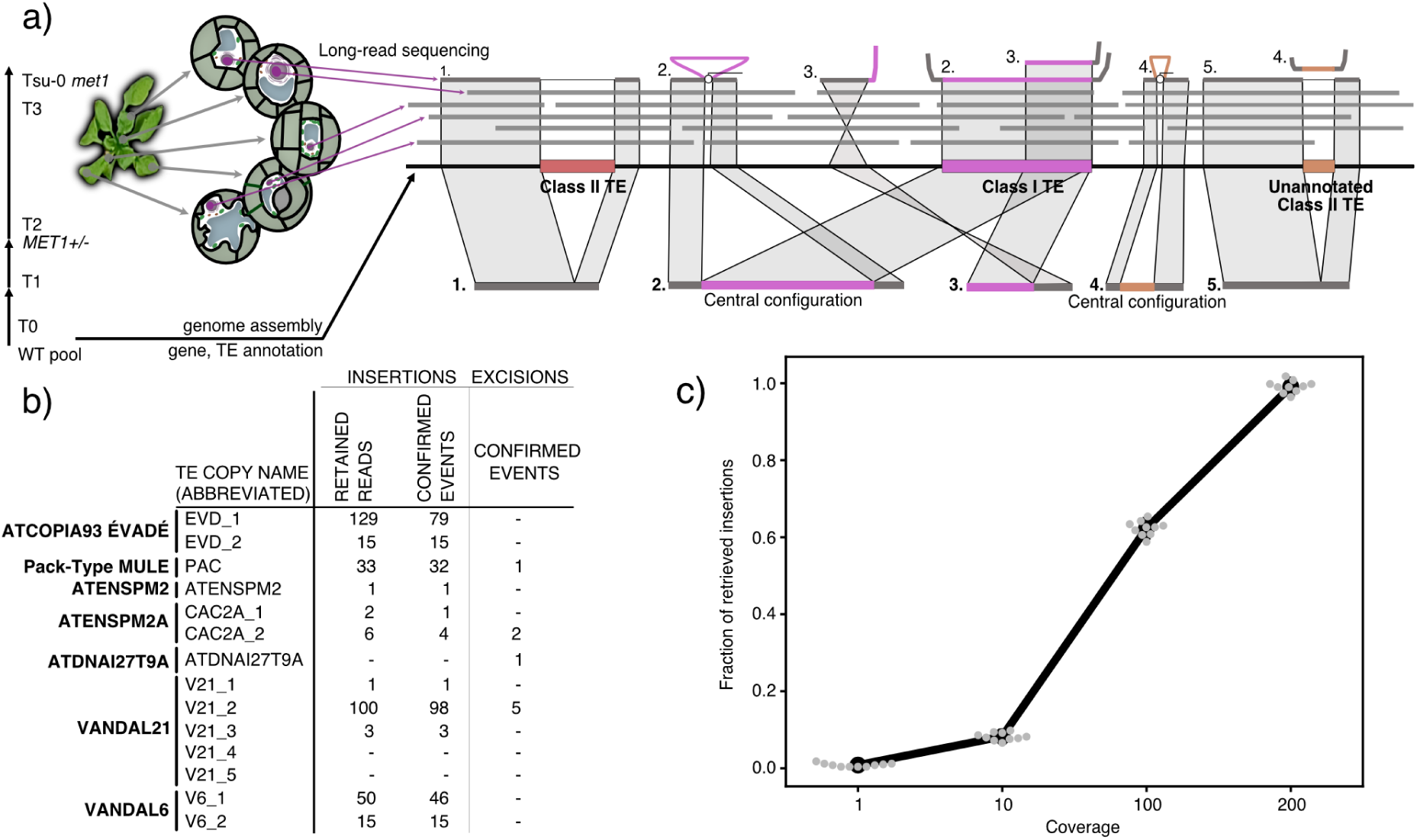
Single-molecule method for detecting somatic TE insertions, excisions in *met1* mutants. **a** Experimental design and diagram that show how different configurations of mapped long reads were used to call somatic mobilization events of TEs. The sequenced *met1* homozygous mutants were three generations away from the plants that were used for *de novo* genome assembly. **b** Numbers of retained candidate reads for insertions and number of non-redundant insertion and excision events. The region around V21_4 and V21_5 had signs of non canonical TE-mediated SV formation (see Fig. 5). **c** Fraction of mock ÉVADÉ somatic insertions retrieved with our method using HiFi read simulation at different coverage. TE: transposable element, TSD: target site duplication, SV: structural variation.

When both ends of the *de novo* TE insertions could be resolved by “chimeric” reads (Fig. 1a, “Central configuration”, “read 2” and “read 4”, see also Additional file 4), which was the case for 27% of our candidate insertion events (63/235), we could almost always identify apparent target site duplications (TSDs). This was the case for 60 out of 63 such events.

Our pipeline strongly relies on the annotation of TEs, and a limitation is that not all TEs might have been properly annotated (see Methods). To reduce false negatives introduced by lack of TE annotation of a mobilizing sequence, we took advantage of the fact that class II TEs are expected not only to transpose, but also to excise (Fig. 1a, reads 1,5), as they move by a “cut-and-paste” mechanism (Quesneville 2020). Therefore, we also retained every read with global evidence for overlapping insertions and deletions of a specific sequence (Fig. 1a, reads 4,5). In this manner, we recovered somatic mobilization events associated with elements that appear to be class II TEs that had been incorrectly annotated or missed altogether during TE annotation. Because this is a relatively specific approach, we might have missed somatic insertions of un-annotated TEs that are not mobilized by a “cut-and-paste” mechanism.

Overall we identified 364 reads that fulfilled our criteria for somatic transposition, which in total explained 295 independent insertions and nine deletions in the ten *met1* individuals (Fig. 1b, 2b), which, since they overlap with class II TE annotations, might correspond to repaired excision events. By contrast, we could not confirm any somatic mobilization in the wild type (Fig. 2b).

**Figure 2.**
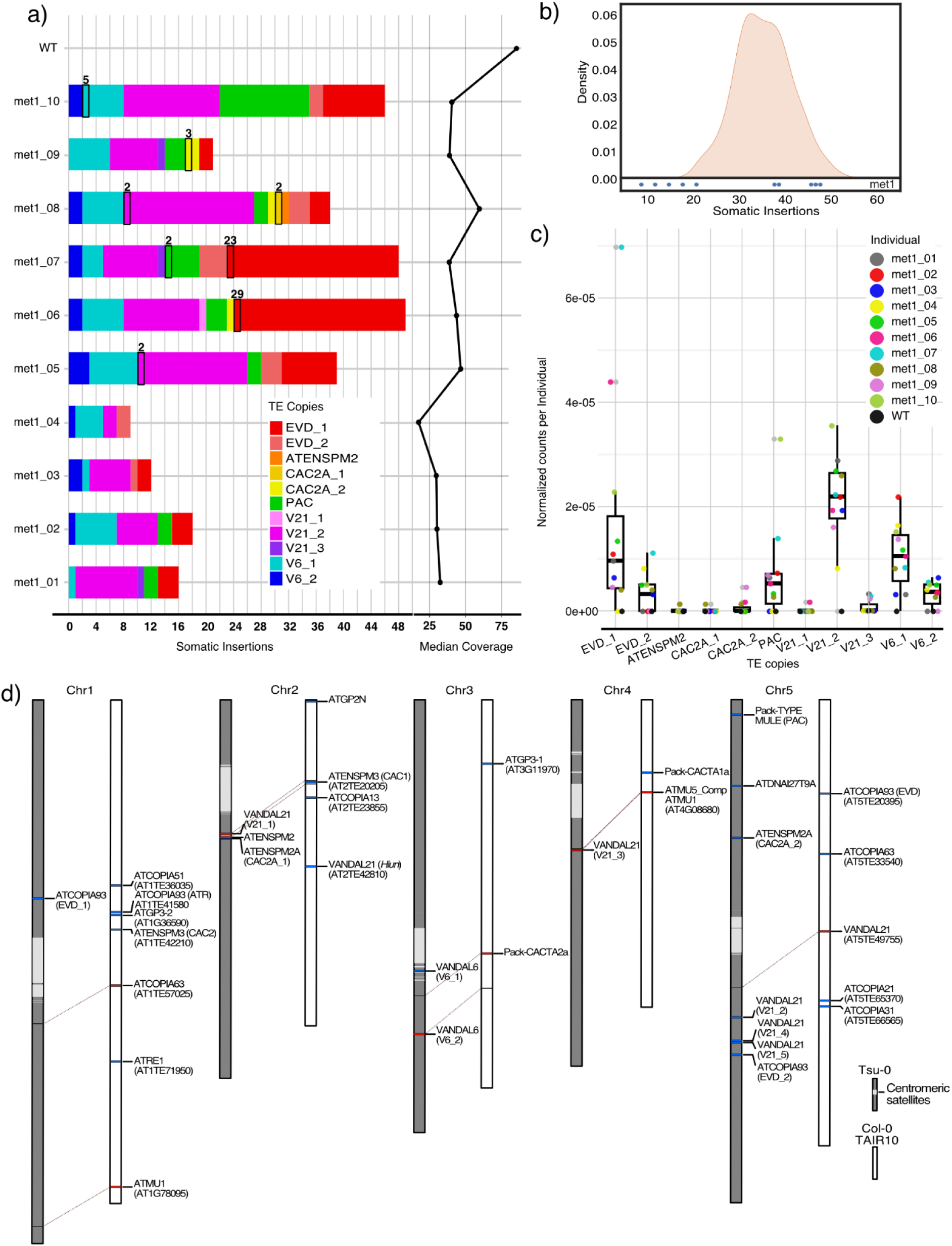
TE mobilization at the individual, family and species level. **a** Stacked barplot indicating the raw count of somatic insertions in each *met1* individual and wild types. Events that were supported by multiple reads are framed by black rectangles and the number of reads supporting them are reported above. Median coverage for the *met1* individual sequencing is shown on the right. **b** Density plot of 1,000 counts of “reads explaining insertions” that were sampled with replacement *n* times, where *n* is the average number of reads per individual, from the entirety of reads from all *met1* individuals. The insertion count for the *met1* individuals is reported at the bottom of the plot with blue dots. **c** Boxplot and jitterplot of individual insertion counts normalized by library size per different TE copy. **d** Double karyoplot showing the difference between somatically mobilizing copies in non-reference Tsu-0 accession (this study) (grey) and published mobile copies from TE mobilization germline studies in *met1*, *ddm1* mutants and epiRILs in the Col-0 reference background (TAIR10) (Singer *et al*. 2001; Miura *et al*. 2001; Kato *et al*. 2004; Tsukahara *et al*. 2009; Catoni *et al*. 2019; Quadrana *et al*. 2019) (white). Syntenic TEs are highlighted in red and a line links them to the position in the Tsu-0 genome. Non-syntenic TEs are highlighted in blue.

Tsu-0 *met1* mutants at 6 weeks after germination have ∼15 leaves (Supplementary Fig. 4), with a size comparable to a 3-week-old wild-type plant. Assuming that cell size is largely unaffected in *met1* mutants and that each leaf has ∼20,000 cells (Gonzalez *et al*. 2010), we estimated that we extracted DNA from a total of ∼300,000 cells per *met1* individual. With an average of 421,821 mapped reads, each cell would have been roughly sampled once. Assuming in addition that most sampled TE mobilization events would occur relatively late during development, the vast majority should only be present in one or a few cells and therefore observed only once. This is indeed what we see, with 96% of apparent TE mobilization events being associated with a single read (Fig 1b, 2b).

To test the sensitivity of our semi-automated method, we randomly inserted 5 ATCOPIA93 ÉVADÉ TE sequences per genome in 100 concatenated *A. thaliana* Tsu-0 genomes and used these modified genomes to simulate PacBio HiFi reads at 2x genome-wide coverage using PBSIM3 software (Ono *et al*. 2022). The simulated reads were either subsampled beforehand or directly mapped to obtain genome-wide coverages of 1x, 10x, 100x or 200x, and subsequently used to call somatic insertions. At the highest coverage we considered, we could recover at least 96% of simulated insertions (Fig. 1c).

### High interindividual variability in active TEs of first-generation *met1* plants

We analyzed the annotations of the detected mobilization events in Tsu-0 *met1*s, finding a total of 14 distinct active TE copies (Fig. 1b) that had evidence of insertions, excisions or both.

Specifically, these included two ATCOPIA93 ÉVADÉ (EVD_1 and EVD2) copies, three VANDAL21 (V21_1 to V21_3) copies, two VANDAL6 copies (V6_1 and V6_2), an uncharacterized non-autonomous element (PAC), one ATENSPM2, two previously uncharacterized ATENSPM family members (CAC2A_1 and CAC2A_2) and a copy of ATDNAI27T9A, a MuDR element with one somatic excision. Two adjacent VANDAL21 copies (V21_4 and V21_5) showed evidence of having caused many SVs, but no canonical somatic transpositions (see below).

Because we had generated independent PacBio HiFi libraries from 10 individual siblings, we could investigate interindividual variation in TE mobilization. The number of detected mobilization events in our individuals ranged from nine to 48, and there were up to 79 reads in a single individual supporting these events (Fig. 2a). To assess whether TE mobility differed significantly between individuals, we used a bootstrap approach to compare the observed insertion counts with a simulated distribution of expected positive reads, *i.e.*, reads that cover a mobilization event (Fig. 2b). Assuming that the true TE insertion frequency in a *met1* individual is close to the arithmetic mean in our sample of 10 individuals, we combined the reads from all 10 individuals and sampled with replacement 1,000 times reads corresponding to the average number of reads obtained from each individual. The results supported the conclusion that the differences in observed transposition events between *met1* individuals were very unlikely to be due to chance, especially for the individuals with few detected events.

In addition, different TE copies showed different degrees of somatic mobilization in different sister plants (Fig. 2c). The mobilization of some families, such as VANDAL6 and VANDAL21, was more uniform than that of other TEs such as EVDs and PAC (Fig. 2c).

In some cases, we detected somatic excisions of full-length class II TEs, most likely underlying repaired excisions (Fig. 1b, Additional file 4). For any given class II TE, somatic excisions were rarer compared to somatic insertions (Fig. 1b).

### Somatic transposition as evidence for annotation of associated TEs

The somatic mobilization events we detected did not always overlap with TE annotations, which we had generated with the EDTA pipeline (Ou *et al*. 2022) based on a curated library of TE models of the Col-0 reference accession (https://github.com/oushujun/TAIR12-TE). We exploited this finding to refine our TE annotation, both in terms of correcting mis-annotated copies and annotating non-reference TEs missing from the curated library.

As an example, all VANDAL elements mobilizing in our experiments had been annotated as non-contiguous segments and were therefore merged. In addition, overlapping insertions and excisions helped us to identify two non-reference TEs that we curated using approaches proposed by Goubert and coauthors (Goubert *et al*. 2022). In one such case, we detected overlapping excisions and insertions of two 8.1 kb fragments that were nearly identical to other two fragments in the Tsu-0 genome, all four of which were predicted to encode an ATENSPM2-like transposase (Supplementary Fig. 5). As these sequences did not comply with the 80-80-80 rule for any of the *consensus* sequences in our Col-0-based library (Wicker *et al*. 2007; Wells and Feschotte 2020), we concluded that they belong to a new TE family that we call ATENSPM2A (CAC2A), with two actively mobilizing elements (Supplementary Fig. 5). Since the CAC2A *consensus* is absent from the Col-0-based library, the EDTA pipeline apparently mis-annotated portions of CAC2As as fragments of different

ATENSPM families (Supplementary Fig. 5). Looking for matches to the Tsu-0 CAC2A *consensus* in the Col-0 reference genome, we found one copy that was apparently mis-annotated as a series of separate TEs, mirroring the fragmented EDTA annotation of Tsu-0, but lacking one of the terminal inverted repeats (TIRs) (Supplementary Fig. 5), most likely compromising its mobilizability. With the PANTERA software (Sierra and Durbin 2024), which can make use of polymorphic copies in different *A. thaliana* genomes (see Methods), we could *de novo* annotate four full copies of CAC2A in the Tsu-0 genome (Supplementary Fig. 5).

A second instance of non-reference TE identification was a ∼1 kb mobilizing element, for which we found several somatic insertions and one somatic excision. This appears to be a Pack-TYPE element (here abbreviated as “PAC”). Pack-TYPE TEs are often missed during *de novo* TE annotation (Gisby and Catoni 2022). Tsu-0 PAC had two direct repeats that could potentially be targeted by VANC6 demethylases (Sasaki *et al*. 2022a), and the mode of the TSD lengths extracted from its somatic insertions coincided with those of VANDAL elements in our dataset (Supplementary Fig. 6). These observations point to the fact that PAC is a non-autonomous VANDAL transposon and possibly a Pack-TYPE MULE (Jiang *et al*. 2004).

Taken together, these results highlight the detection of somatic mobilization as a source for discovering new elements and assisting TE annotation. In this manner, we could identify new TEs based on the ability of sequences to transpose, independently of sequence similarity to known elements.

### Intraspecific variation of the mobilizable mobilome

We compared our inventory of mobile TEs in Tsu-0 *met1* mutants with a collection of TEs that have been reported to be active in the germline of *ddm1, met1* or epiRILs in the Col-0 reference accession (Singer *et al*. 2001; Miura *et al*. 2001; Kato *et al*. 2004; Tsukahara *et al*. 2009; Fu *et al*. 2013; Catoni *et al*. 2019; Quadrana *et al*. 2019). Among the mobilizing TE copies in Tsu-0, only four were present at syntenic locations in Col-0, but none of these had been previously reported as mobile in *met1* and *ddm1* mutants and epiRILs (Fig. 2d). Likewise, none of the TEs that are mobile in Col-0 mutants were syntenic to TEs that are mobilizable in Tsu-0.

At the family level, we did not find evidence for mobilization of many Ty1/Copia, CACTA/EnSpm, Ty3 and MuDR elements, which often actively transpose in Col-0 *ddm1* epiRILs (Singer *et al*. 2001; Quadrana *et al*. 2019; Zhang *et al*. 2023) (Fig. 2d). At the same time, we detected the mobilization of families either absent or previously unknown to mobilize in Col-0.

Both copies of intact full-length EVDs, ÉVADÉ and ATTRAPÉ (ATR), are activated in Col-0 epigenetic mutants, with ÉVADÉ arguably being the most active TE in the Col-0 background (Mirouze *et al*. 2009; Marí-Ordóñez *et al*. 2013; Quadrana *et al*. 2019). Tsu-0 has five EVDs that are highly similar to each other. Four of these were intact, but only two were mobile in our material (here abbreviated as EVD_1, EVD_2) (Fig. 1b,2a).

We found three copies of VANDAL21 (V21_1, V21_2, V21_3) and two copies of VANDAL6 (V6_1, V6_2) to be somatically mobilizing in Tsu-0. As mentioned before, we did not further consider a region in chromosome 5 with two head-to-head VANDAL21s (V21_4 and V21_5), which showed high somatic variability, but where we could not confidently call somatic transposition events. Of the mobilizing VANDAL21s in Tsu-0, only V21_1 and V6_2 were syntenic in Col-0 (Fig. 1b,2a).

We detected mobilization of a complete class II ATENSPM2 element, which is lacking from Col-0, and of three copies of ATENSPM2A, a previously uncharacterized ATENSPM/CACTA family member (Fig. 1b, Supplementary Fig. 5). A different ATENSPM/CACTA family member, ATENSPM3, becomes mobile in Col-0 *ddm1* and *met1* epiRILs (Quadrana *et al*. 2019) (Fig. 2a).

We found one mobilizing non-autonomous Pack-TYPE MULE (Jiang *et al*. 2011), present in a single non-reference copy. While, to our knowledge, Pack-TYPE MULEs are not active in Col-0, non-autonomous Pack-TYPE CACTA elements have been reported to become mobile in *met1* epiRILs (Catoni *et al*. 2019).

We also found one excision of ATDNAI27T9A, a MuDR TE for which we did not find any associated insertion. As host-mediated mechanisms could give rise to the same outcome through TE-independent activity, we were not confident in excluding it from being a false positive. Methylation at ATDNAI27T9A is highly sensitive to temperature (Dubin *et al*. 2015) and it is mobile in Tsu-0 wild-type plants (Baduel *et al*. 2021).

In conclusion, our approach can be used to investigate intraspecific variation of actively mobilizing TEs.

As expected, identity between the two long terminal repeats (LTRs) seemed to be a necessary condition for EVD mobilization, as the LTRs of both mobile copies were 100% identical. The immobile EVD_3 was also intact, without ORF truncating mutations, and had 100% LTR identity (Fig. 3a), indicating that LTR identity and ORF intactness are necessary but not sufficient for mobilization of EVDs.

**Figure 3.**
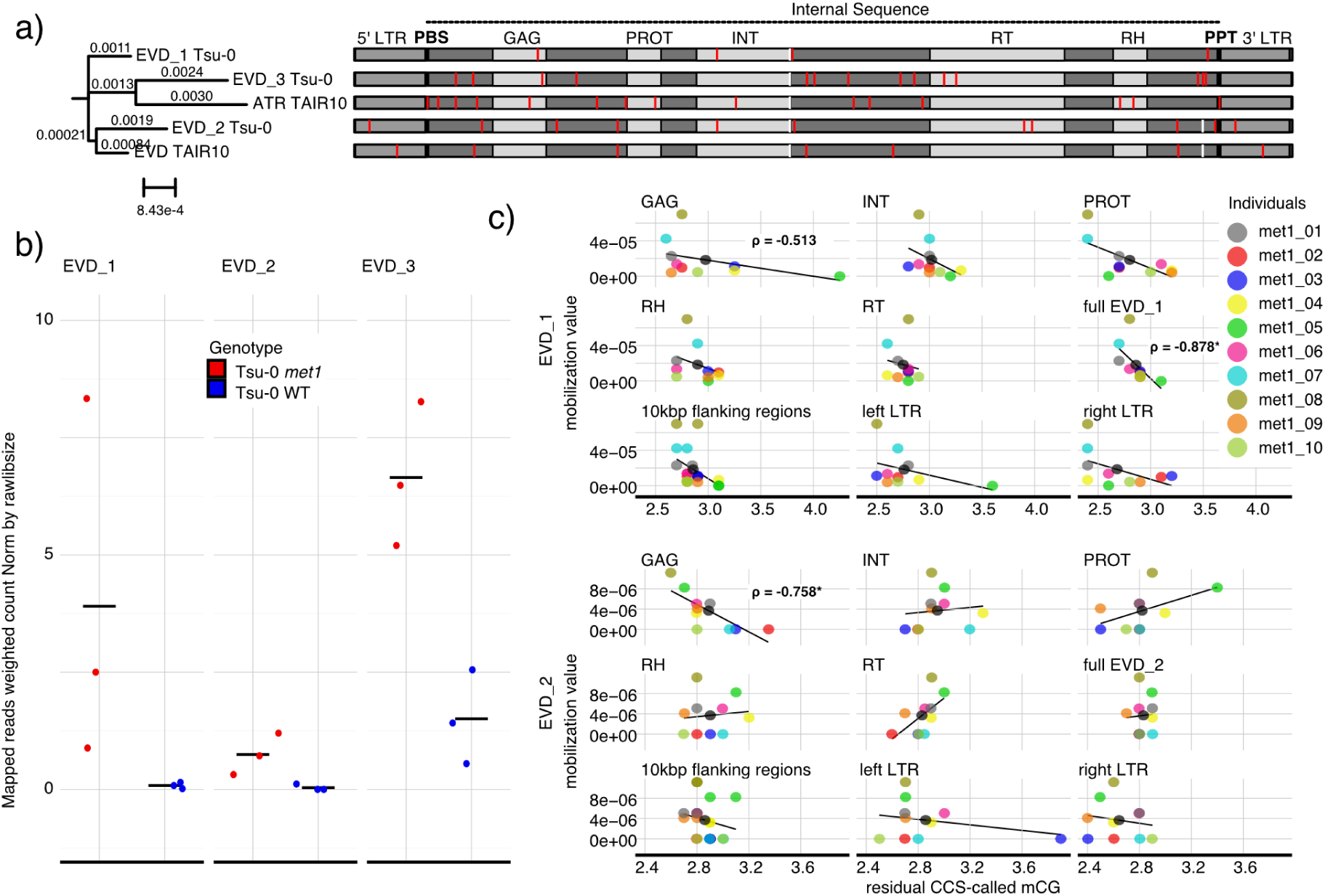
Differential mobility of ATCOPIA93 ÉVADÉ copies. **a** Phylogenetic t ree (left) and multiple sequence alignment (right) of intact non-degenerate EVD copies in Tsu-0 and Col-0. LTRs (grey), primer binding site (PBS, black), polypurine tract (PPT) (black), and protein domains (GAG, PROT, INT, RT, RH) (light grey) are indicated. Substitutions are shown in red, small deletions in white. b Dotplot of expression levels of intact and non-degenerate Tsu-0 EVDs. Average is indicated as a black solid line. c Scatterplot with regression line of mobilization values, i.e., individual counts normalized for library size as in Fig. 2d, as a function of residual PacBio HiFi-called CpG methylation for EVD_1 and EVD_2. Values are plotted for the full copies, LTRs, flanking sequences and sequences encoding for protein domains. *Spearman correlation p-value<0.05; ***p-value<0.001; n.s., p-value≥0.05.

Re-mapping of published RNA-seq data (Srikant *et al*. 2022) indicated a slightly higher expression level of EVD_2 than EVD_1 (Fig. 3b) in Tsu-0 *met1*, but comparable accessibility and residual DNA methylation levels (Supplementary Fig. 7). EVD_3 had the lowest expression level (Fig. 3b), with the proviso that estimating expression levels of elements with closely related copies is inherently difficult.

To look into residual methylation as a driver for interindividual differences in EVD mobility, we extracted CG methylation profiles of mobilizing EVDs from HiFi circular consensus sequencing (CCS) reads. Residual CG methylation on EVDs was generally very low in *met1* compared to the wild type (Supplementary Fig. 7). Across individuals, EVD_2, and less so EVD_1, showed an inverse correlation between residual CG-methylation in the GAG portion and mobilization level (Spearman correlation coefficient = -0.76/-0.51, p-value = 0.01/0.13) (Fig. 3c). This result is in line with EVD’s GAG being methylated via translation-dependent RdDM (Oberlin *et al*. 2022) as a first line of defense against its pervasive mobility. EVD_1 as a whole showed a strong inverse correlation between total residual CG-methylation and mobilization level (Spearman correlation coefficient = -0.88, p-value < 0.001) (Fig. 3c).

Taken together, expression, accessibility and residual methylation are not good predictors for differential mobilization of EVDs in *met1* Tsu-0 mutants.

### Somatic insertion preferences generally mirror germline bias

Actively mobilizing transposons did not show any evident bias towards or against specific regions of each chromosome, except for centromeres (Fig. 4a). We detected only five somatic transpositions within centromeric satellite repeats and three within centrophilic Athila elements in all chromosomes but chromosome 3 (Fig. 4a).

**Figure 4.**
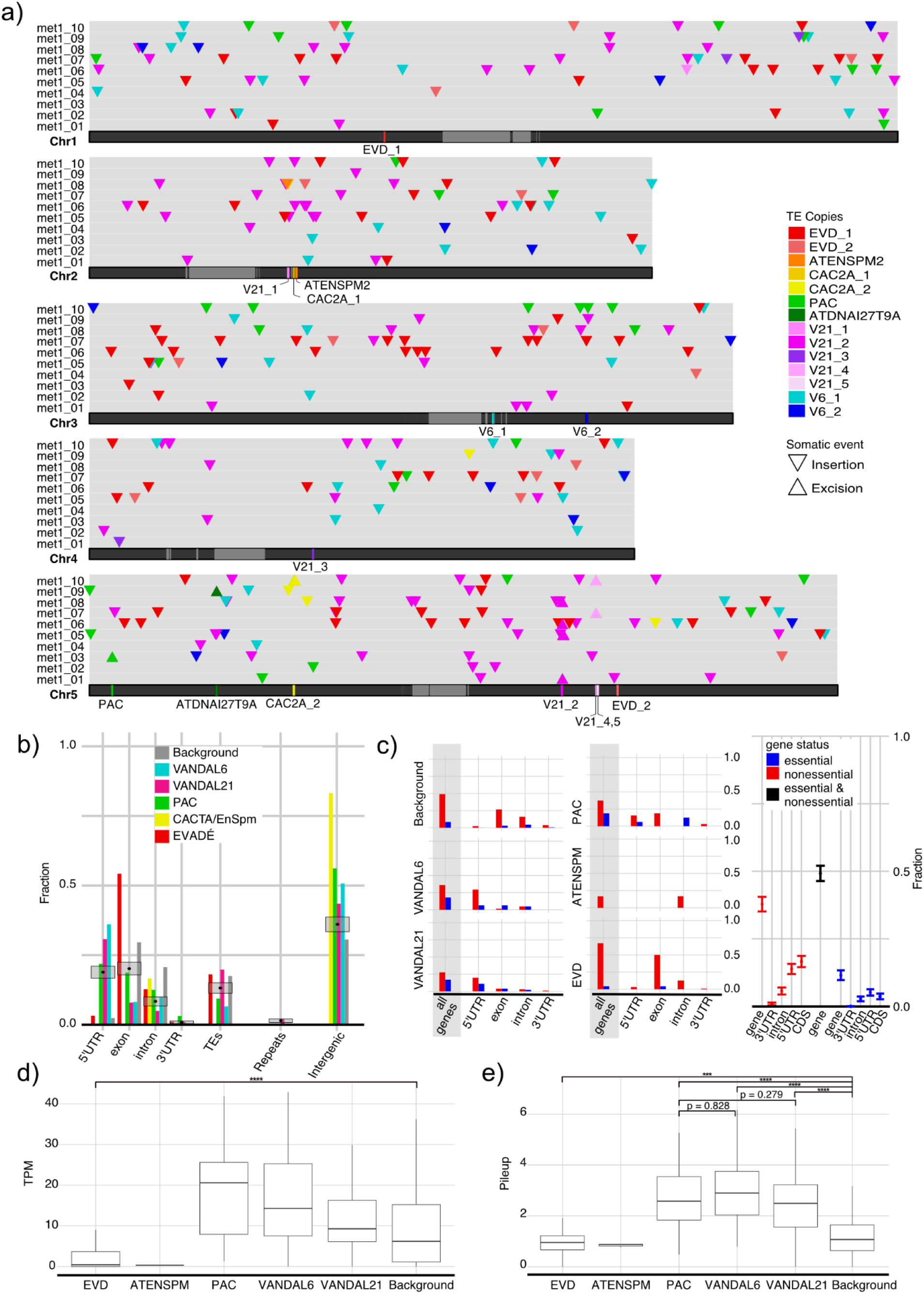
Insertion preference in Tsu-0 *met1* mutants. **a** Karyoplot of genome-wide somatic insertion landscape. Centromeric satellites in lighter grey. Mobile TE copies are highlighted with different colors, with insertions as downward pointing and excisions as upward pointing triangles. **b** Insertion preferences towards genomic features. “Background” describes the fraction of randomly generated sites. Grey boxes represent the random distributions of genomic features at somatic insertions generated by 1,000 iterations of sampling with replacement from the entirety of the *met1* reads. Black dots indicate the means. **c** Barplot of the somatic insertions in genes colored by “essentiality”. “Background” describes the fraction of randomly generated insertions. On the right, distributions of “essentiality” of genes disrupted by mobilizing TEs generated by 1,000 iterations of sampling with replacement from the entirety of the *met1* reads. **d** Expression levels of genes disrupted by insertions in *met1*. “Background” represents randomly generated sites. Outliers are not shown. **e** Accessibility of insertion sites in *met1*. “Background” represents randomly generated sites. Outliers are not shown. ***Welch Two-sample t-test p-value<0.001; ****p-value<0.0001. Kolmogorov-Smirnov test was performed beforehand to test for normality. TPM, transcripts per million.

Concerning somatic insertion bias towards specific genomic features, we characterized the insertion sites (Fig. 4b) and measured their distance to the nearest gene, TE, and repeat type (rDNA, centromere, telomere, nuclear insertions of organellar DNA) (Supplementary Fig. 8). We compared the insertion preferences with randomly generated insertion sites and with a random distribution of insertion site features assuming the arithmetic mean of all the insertions to be the true mean. Since ATENSPM insertions were too few, we could not assess biases towards any particular genomic feature. For the other TE categories, we found the insertion preference towards genomic features to be different between EVDs, and VANDALs together with PAC (Fig. 4b; Supplementary Fig. 8).

In our material, VANDALs preferentially inserted near genes, especially 5’ UTRs and upstream regions, with PAC showing similar insertion preferences. EVDs showed a bias towards exons (Fig. 4b. Supplementary Fig. 8), consistent with previous studies (Fu *et al*. 2013; Quadrana *et al*. 2019) (Fig. 4b).

Somatic insertion preference of EVDs also mirrored the germline bias towards genes, especially non-essential genes (Quadrana *et al*. 2019) (Fig. 4c), suggesting that this pattern might not be due to purifying selection at the cellular level.

### Sites disrupted by different TEs show different expression and accessibility

We assessed the expression of genes disrupted by somatic insertions in *met1* with published RNA-Seq data (Srikant *et al*. 2022) by re-mapping reads to our *de novo* assembled Tsu-0 genome. EVDs showed a bias towards genes with a significantly lower expression in Tsu-0 *met1* (Fig. 4d). In contrast, VANDALs and PACs more often transposed within genes with a higher expression level compared to EVDs, but not significantly different from random (Fig. 4d).

To test whether the pattern of somatic TE insertions reflects chromatin accessibility landscape, we made use of ATAC-Seq data generated for Tsu-0 *met1* (Srikant *et al*. 2022). We found that EVDs had opposing insertion preferences compared to the majority of TE families, as they were more likely to insert in less accessible regions, whereas the insertion sites of the other families were in general significantly more accessible (Fig. 4e). The similarity in insertion preferences between PACs and VANDALs were reflected in similar accessibility preferences (Fig. 4e). Accessibility of somatic insertion sites was not explained by residual DNA methylation, as deduced from CCS-called and BS-Seq methylation data (Supplementary Fig. 9).

### VANDAL21-associated SVs display patterns of alternative transposition

We found evidence of extensive somatic rearrangements in a region in chromosome 5 centered around two head-to-head VANDAL21s, which we abbreviated as V21_4 (left) and V21_5 (right) and which were separated by a ∼200 bp spacer (Fig. 5). These two VANDAL21 copies disrupted a region with homology to a pseudogene of the TIR-NLR family in Col-0, AT5G40920 (Supplementary Fig. 11). Many reads with supplementary alignments (“chimeric” reads) mapped to this region (Fig. 5a), but visual inspection did not support the presence of *bona fide* transposition events, as there were no reads where both ends simultaneously mapped to the same region and included a complete or partial VANDAL21 element in their center, while respecting the TE boundaries (Additional file 4). The structural rearrangements in this region were diverse, but none could be straightforwardly associated with simple insertion/excision events. These SVs included inversions, deletions, apparent translocations where reads described a VANDAL21 being contiguous with a different region of the genome (“out-region”) and “complex” SVs where a duplication and a deletion in addition to the inversion of the spacer co-occurred (Fig. 5b). The most often found SV was the deletion of a median of 185 bp spacer DNA between the two VANDAL21s (Fig. 5b, Supplementary Fig. 11), also evident from a drop in read coverage (Fig. 5a), which did not always coincide with the borders of V21_4 and V21_5 (Supplementary Fig. 11). In addition, there were excisions of V21_4 and V21_5 (Fig. 5b), four longer deletions comprising the left flanking region of V21_4, inversions of either V21_4, V21_5 or the internal spacer, along with other “complex” SVs (Fig. 5b). “Chimeric” reads mapping to this region very often spanned either V21_4 or V21_5 and the end of the mapping was right before or after the spacer sequence respectively (Fig. 5a). The break-end (BND) coordinates of those “chimeric” reads that described “out-region” non-local rearrangements (Fig. 5b) were located almost entirely on chromosome 5, with more than 75% within a 10 Mb region around the head-to-head VANDAL21s (Fig. 5c), pointing to a spatial constraint. BND coordinates of “out-region” SVs also showed a bias towards 5’ UTRs (Fig. 5d), reminiscent of known VANDAL insertion preferences.

**Figure 5.**
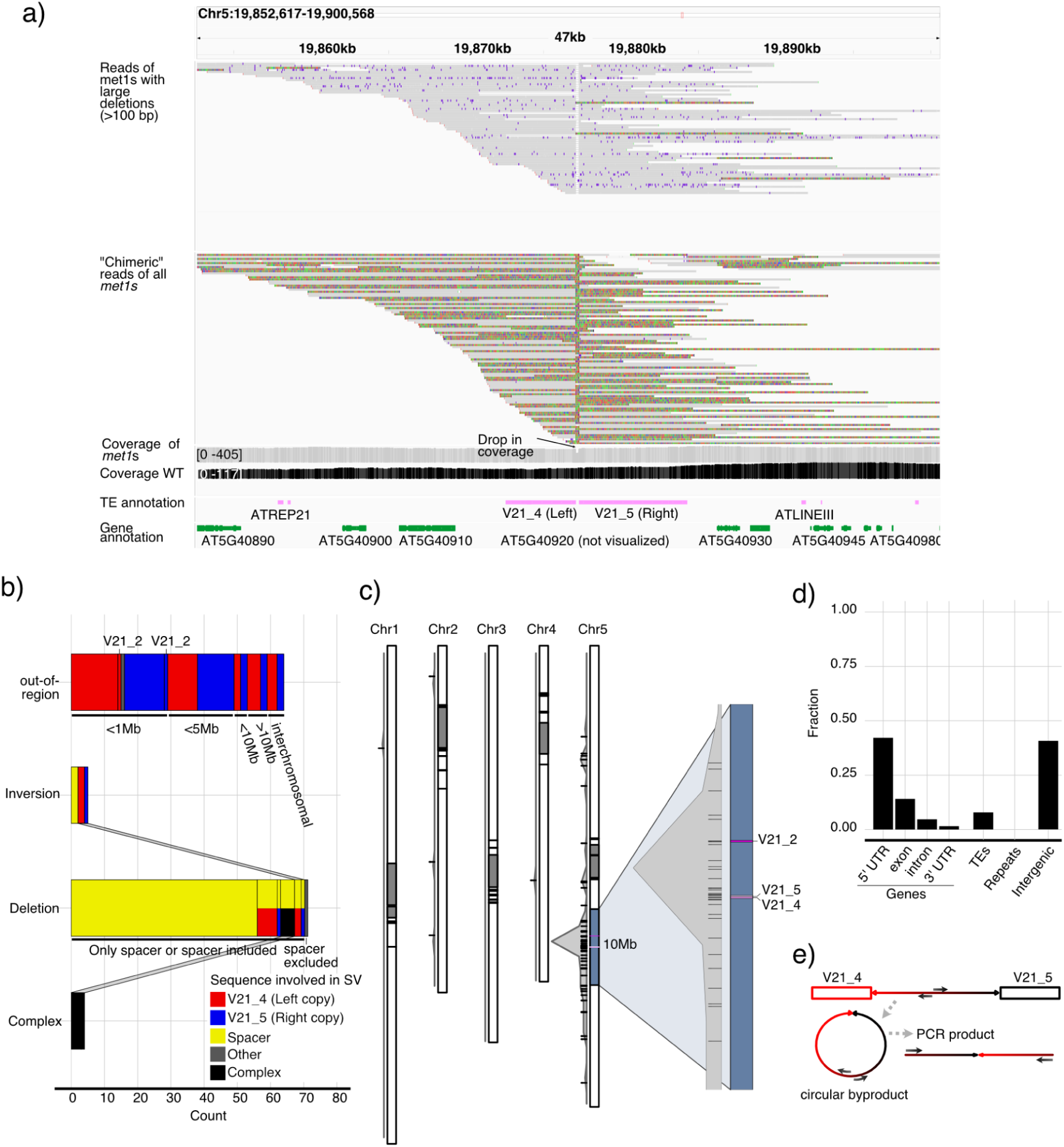
Local VANDAL21-mediated somatic hypermutability. **a** IGV screenshot of reads mapping to the hypermutable region. Tracks, from top: reads with spacer deletion, reads with supplementary alignment (colored portion) indicating putative somatic hypermutability, *met1* reads (unfiltered) coverage in grey, wild-type reads (unfiltered) in black, TE annotation in pink, gene annotation in green. **b** Counts of different classes of SVs overlapping with the region colored by involved sequence. The distance from the region and the mapping to the close V21_2 copy is indicated. It is specified if deletions contain the spacer or not. If a deletion involves two different sequences it is colored with two different colors. SVs that could be classified in two different ways are linked. **c** Karyoplot with density of BND mapping coordinates of “out-region” SVs not overlapping with the hypermutable region. On the right, close-up of the 10 Mb centered around V21_4 and V21_5. **d** Barplot of the fraction of genomic features overlapping with BND coordinates of “out-region” SVs. **e** Schematic representation of spacer circular DNA detection via PCR. SV: structural variation, BND: break-end.

It is plausible that the specific configuration and proximity of the two adjacent VANDAL21s could lead to alternative transposition, a phenomenon that has been extensively studied for Ac/Ds transposons in maize and which can generate deletions, duplications, composite insertions (Zhang and Peterson 2005; Zhang *et al*. 2014; Su *et al*. 2020), inversions (Sharma and Peterson 2023), as well as interchromosomal translocations (Wang *et al*. 2015). It takes place during DNA replication, after multivalent transpososomes have bound to distinct elements, transposition intermediates are aberrantly resolved, generating different types of SVs in combination with DNA repair mechanisms (Zhang *et al*. 2014). As a result, the spacer between the transposons is circularized and lost (Zhang and Peterson 2004). Virtually all the

SVs that were identified at this head-to-head VANDAL21 region could be explained by alternative transposition, although the reads were not long enough to fully resolve all the SVs including TSDs (Fig. 5b).

That alternative transposition could underlie the observed hypervariability of this region was supported by direct detection of the circular excised spacer using PCR with outward-facing primers (Fig. 5e), including when PCR was preceded by rolling circle amplification (RCA) (Supplementary Fig. 11). This pointed to the presence of alternative transposition byproducts.

To search for additional alternative transposition events in our dataset, we looked for similar patterns of “chimeric” reads mapping to regions in which either two annotated class II TEs with the same classification or simply two TIR sequences – extracted from our TE *consensus* library – were arranged in a head-to-head fashion. We did not find any additional instances, which perhaps is not surprising, given the greater activity of VANDAL21s compared to any other TE in our material.

## Discussion

By long-read sequencing of genomic DNA from hypomethylated *met1* mutants in *Arabidopsis thaliana*, we detected somatic insertions and excisions of transposable element (TE) copies. The analysis of closely related siblings (Fig. 1a) allowed us to assess interindividual variation in TE mobility (Fig. 2a, 4a), while use of the non-reference Tsu-0 background revealed TEs absent or not mobile in Col-0 reference accession (Fig. 2d). Our approach also helped to generate support for an underappreciated aspect of TE biology, namely the hypermutability linked to alternative transposition, a phenomenon previously studied for Ac/Ds transposon in maize and snapdragon, and P-elements in Drosophila, but not in *A. thaliana* (Gray 2000; Su *et al*. 2018).

Plants have flexible germline specification, allowing the inheritance of somatic TE variations (Bourque *et al*. 2018) and in some cases for clonal propagation. Even if somatically variable cells do not contribute to the germline, *i.e.*, becoming ‘germ-track’ – as per Haig’s definition (Haig 2016) –, they still can significantly impact phenotypes (McClintock 1950), making it important to assess their prevalence. Measuring somatic events in plants bypasses some biases found in germline studies. For instance, the plant life cycle, with its extended gametophytic phase (Bowman *et al*. 2016), may act as a bottleneck for the transmission of deleterious mutations, limiting the passage of somatic transpositions to future generations.

We classified all low-coverage TE mobilizations as “somatic” because it was unlikely that they had become fixed in the L2 layer, which gives rise to the gametes (“germ-track”).

As Siudeja, van de Beek and coauthors (Siudeja *et al*. 2021) have pointed out, long reads can span complete insertions, thus resolving both the central TE and their flanking sequences, intrinsically increasing confidence in detection of somatic events – even if supported only by single reads. Several aspects of our analyses further boost the confidence we have in our somatic insertion calls: (1) absence of mobilization in wild-type plants; (2) reads having been precisely mapped to full or partial TEs; (3) target site duplication (TSD) being detected in “chimeric” reads; and (4) mobilizing TEs belonging to families known to become active in the germline of epigenetic mutants.

In Tsu-0 *met1* mutants, somatic TE mobilization rates appear to be higher than in Col-0 *met1* mutant (Mathieu *et al*. 2007; Reinders *et al*. 2009; Mirouze *et al*. 2009; Rigal *et al*. 2016), which may be due to Tsu-0 belonging to an miRNA haplotype that reduces TE silencing efficiency (Borges *et al*. 2018), although we cannot exclude the involvement of germline biases. In our dataset, the extent of TE mobilization seemed to vary across individuals (Fig. 2a), with only some elements showing high variation in mobilization rates (Fig. 2c). We confirmed previous findings on TE insertion preferences (Fig. 4b,c), such as VANDAL elements targeting 5’ UTRs (Fu *et al*. 2013) and EVDs preferring non-essential genes as targets (Quadrana *et al*. 2019). We detected centromeric insertions of ÉVADÉ and VANDAL elements, probably as a consequence of changes in chromatin architecture change and increased accessibility of centromeres in *met1* mutants (Feng *et al*. 2014; Zhong *et al*. 2021; Naish *et al*. 2021). The insertion of EVDs into genes with lower expression levels may be associated with transcriptional repression associated with H2A.Z enrichment (Coleman-Derr and Zilberman 2012; Gómez-Zambrano *et al*. 2019).

Beyond TE derepression, hypomethylation impairs genome stability more broadly, generating SVs (Zhang *et al*. 2023) and TE-driven mechanisms may contribute to region-specific hypermutability (López-Cortegano *et al*. 2023). We propose that alternative transposition between closely positioned VANDAL21 transposons (V21_4 and V21_5) may explain local hypermutability, similar to alternative transposition of Ac/Ds TEs in maize (Zhang *et al*. 2014; Su *et al*. 2018). While VANDAL transposons are class II TEs, like Ac/Ds, they have not been linked to alternative transposition before. A VANDAL5-mediated paracentric inversion event has been suggested as having generated the polymorphic chromosome 4 heterochromatin knob in *A. thaliana (Fransz et al. 2016)*. The proposed causal mechanism involves the 5’ end of one of two proximal VANDAL5s having inserted into a distal gene while its 3’ end remained attached to its origin (Fransz *et al*. 2016). An alternative transposition event could have been the underlying cause of this paracentric inversion in agreement with directly demonstrated mechanisms in maize (Zhang *et al*. 2009). The narrow 10 Mb break-end coordinates interval of V21_4 and V21_5 rearrangements (Fig. 5c) likely reflect positional constraints or reduced fitness of cells with extreme rearrangements, such as translocations or large deletions. These alternative transposition-driven SVs may contribute to the reduced viability of *met1* by destabilizing chromosome structure and accumulating DNA damage, which is prominent in severely dwarfed *met1* plants (Liang *et al*. 2022).

With our approach we detected non-autonomous and mis-annotated class II TEs not found in the Col-0 reference-based TE library. While the contribution to the improvement of TE annotations was small here, improvements might be more substantial in other plant species with more TE-rich genomes, especially those with prominent clonal propagation abilities, where somatic variation is intrinsically more impactful. For species where epigenetic mutants and epiRILs are difficult to obtain, this method could provide insights into TE mobility in cases where germline studies are impractical – which is the case even for some *A. thaliana* accessions (Srikant *et al*. 2022).

Lastly, the lack of synteny in actively mobilizing TEs between Tsu-0 and Col-0 (Fig. 2d) suggests considerable diversity in the mobilizable mobilome of *A. thaliana*. Future efforts to explore this diversity at the species level are warranted.

## Conclusions

With the use of long-read DNA sequencing of *A. thaliana met1* mutants we revealed widespread somatic TE transposition, with variable mobilization extents across individuals. Unlike traditional germline-focused approaches, methods such as ours enable the interrogation of the mobilome in mutant individuals already in the first generation, without the need for inherited events, and thus is applicable to infertile, semi-lethal genotypes. Our approach taught us new aspects of TE biology as, for instance, we found evidence for alternative transposition events as the cause for local hypermutability.

## Methods

### Plant material

Seeds of *Arabidopsis thaliana* Tsu-0 accession (1001 Genomes Project ID 7373) were sterilized with gaseous chlorine and stratified in 0.1% w/v agarose for 7 days at 4°C in the dark. Tsu-0 wild-type seeds from the pool used for transformation with the *MET1* targeted CRISPR-Cas9 guide construct in (Srikant *et al*. 2022) were directly sown on soil. Because of their delayed development, Tsu-0 *met1* mutants (line 2, (Srikant *et al*. 2022)) were first grown for 2 weeks on ½ strength MS agar plates supplemented with 1% w/v sucrose and then transferred on soil. Before harvest, plants were placed for 24 h in the dark to reduce chloroplast DNA. The aerial parts were harvested and snap-frozen with liquid nitrogen. Wild-type material was collected from pools of plants four weeks after germination and ten *met1* plants were collected individually three weeks after transfer on soil, at a stage of development comparable with that of the wild types. Mutant plants at harvest are shown in Supplementary Fig. 4.

### DNA extraction, PacBio HiFi library preparation and sequencing

The plant material was finely ground with mortar and pestle in liquid nitrogen, and high-molecular-weight (HMW) DNA extraction was carried out as published (Rabanal *et al*. 2022).

HMW DNA was sheared with Covaris gTUBEs and was then directly used for PacBio HiFi SMRTbell library preparation, as specifically described for the Col-0 HiFi library in (Rabanal *et al*. 2022). 5 µg of Tsu-0 wild-type DNA was used as input for HiFi SMRTbell library using SMRTbell Express Template Prep Kit 2.0 with the barcode “bc1010”, multiplexed with two other libraries and sequenced on a single SMRTcell.

Sheared DNA from individual plants *met1_01* to *met1_10* (Supplementary Fig. 4) was used as input for PCR amplification of *MET1* exon 7 (Srikant *et al*. 2022). Sanger sequencing of the PCR product was used to assay the known mutant sequence. Different amounts of input DNA (1480 - 5000 ng) were used for preparation of individual *met1* mutant HiFi SMRTbell libraries with SMRTbell Express Template Prep Kit 3.0 with 10 adapters “bc1001” to “bc1003” and “bc1008” to “bc1016”, sequenced on a single SMRTcell and resequenced on another SMRTcell multiplexed with other libraries. Prior to sequencing the final pooled libraries were size-selected on a BluePippin (SageScience) with 10 kb cutoff in a 0.75% DF Marker S1 High Pass 6–10 kb v3 gel cassette (Biozym). Sequencing was performed on the Sequel II system using Binding Kit 2.2 for wild-type material and Binding Kit 3.2 for mutants.

### Genome assembly

High Fidelity (HiFi) reads for all samples were generated with the DeepConsensus pipeline (https://github.com/google/deepconsensus): generation of draft consensus sequences with pbccs v6.4.0 (‘--min-rq=0.88’) (https://ccs.how/), alignment of subreads to the draft consensus sequence with v0.2.0, with DeepConsensus v1.2.0 run in CPU mode (Baid *et al*. 2023). Assembly of the Tsu-0 genome was carried out with hifiasm v.0.19.2 (‘-l0 -f0’) (Cheng *et al*. 2021). Contigs larger than 50 kb were scaffolded into chromosomes with RagTag v.2.1.0 (‘scaffold -q 60 -f 30000 -i 0.5 --remove-small’) (Alonge *et al*. 2022), using as reference genome a version of TAIR10 in which organellar DNA nuclear insertions, and repetitive sequences had been masked (Rabanal *et al*. 2022).

The contiguity of the genome assembly was assessed using Quast v5.2.0 (Mikheenko *et al*. 2023) (https://github.com/ablab/quast). The completeness was checked with BUSCO v5.4.2 (Manni *et al*. 2021) (https://busco.ezlab.org) in ‘genome’ mode using ‘viridiplantae_odb10’ lineage gene models.

### Annotation

#### Repeats

Repeats were annotated with a previously described pipeline that uses RepeatMasker (Smit *et al*. 2013-2015) (http://www.repeatmasker.org) with a custom library to identify centromere satellite repeats, ribosomal RNA genes, and telomeres, and minimap2 (Li 2018) (https://lh3.github.io/minimap2/minimap2.html) to find nuclear insertions of organellar DNA (Rabanal *et al*. 2022).

#### Genes

Genes were lifted over to the Tsu-0 genome from the TAIR10 annotation (12 July 2019) with liftoff v1.6.3 (Shumate and Salzberg 2021) (https://github.com/agshumate/Liftoff), with ‘-copies’ option and a list of features was provided for lift-over: ‘CDS, exon, five_prime_UTR, gene, miRNA, mRNA, ncRNA, snoRNA, snRNA, three_prime_UTR’. Exclusively primary isoforms were included. Gene annotation was translated with gffread (Pertea and Pertea 2020) (https://github.com/gpertea/gffread) and quality-checked via BUSCO v5.4.2 (Manni *et al*. 2021) (https://busco.ezlab.org) in ‘protein’ mode using ‘viridiplantae_odb10’ lineage.

#### Transposable elements

For primary TE annotation, we ran the EDTA v2.2 (Ou *et al*. 2019) (https://github.com/oushujun/EDTA) pipeline providing the Col-CC curated library (https://github.com/oushujun/TAIR12-TE). To overcome fragmentation of seemingly intact annotated TEs we increased the ‘-maxdiv’ parameter from 3.5 to 5, as EDTA tries to combine RepeatMasker lines that are likely the same element (https://github.com/oushujun/EDTA/blob/v2.2.0/EDTA.pl#L694).

We applied a custom script to merge all elements in the TE annotation file that had the same name and were at most 1 bp apart (https://github.com/aerilli/Somatic-transposition_met1). TE annotations overlapping repeats (rDNA, centromere, telomere, nuclear insertions of organellar DNA) annotations were excluded.

We made use of LTRpred v1.1.0 (Drost and Sanchez 2019) (https://github.com/HajkD/LTRpred) to calculate LTR identity and annotate primer binding site (PBS) and polypurine tract (PPT) of EVDs, and DANTE v0.1.9 (Neumann *et al*. 2019) (https://github.com/kavonrtep/dante) to annotate domains. We used EMBOSS getorf (https://emboss.sourceforge.net/apps/cvs/emboss/apps/getorf.html) to predict orfs of the different EVDs as a measure of their degeneration state.

To determine whether improved TE annotation methods would substantially affect our results, we additionally ran PANTERA v0.2.1 (Sierra and Durbin 2024) (https://github.com/piosierra/pantera) to detect TEs in a pangenome graph that had been generated from 67 *A. thaliana* genome assemblies from different accessions. TE models detected by PANTERA were classified by TEsorter v1.4.6 (Zhang *et al*. 2022) (https://github.com/zhangrengang/TEsorter) and only the models with a confident classification were kept. Afterwards, the models were inspected using TEtrimmer v1.4.0 (Qian *et al*. 2024) (https://github.com/qjiangzhao/TEtrimmer) and new consensus sequences were built for some TE families. We added 93 non-redundant TE models detected by PANTERA to the Col-CC curated library and used this library to annotate the Tsu-0 genome with EDTA v2.2.1. Also, TE models generated by *de novo* EDTA annotation step that overlapped with a CDS or with more than 50% of a protein-coding gene were removed. By using this improved TE annotation we could detect single insertion events of few new TE copies.

### Read simulation

To simulate HiFi reads of somatic TE insertions, we concatenated 100 Tsu-0 genomes and inserted the sequence of the Tsu-0 ÉVADÉ transposon (EVD_1) with seqkit v2.8.2 ‘mutate’ function (Shen *et al*. 2024) (https://bioinf.shenwei.me/seqkit/usage/#mutate) at 500 randomly generated positions, *i.e.*, on average 5 per genome. We then used the concatenated, modified genomes to run pbsim3 v3.0.4 software (Ono *et al*. 2022) (https://github.com/yukiteruono/pbsim3) as follows: ‘pbsim --strategy wgs --method errhmm --length-mean 15775.0 --length-sd 7500 --length-min 5000 --difference-ratio 22:45:33 --errhmm ./pbsim3/data/ERRHMM-SEQUEL.model --depth 2 --genome ${concatenated_genome_with_random_insertions} --accuracy-mean 0.9 --pass-num 7’.

Length parameters were chosen to coincide with the specific details of our dataset. With the depth parameter set to ‘2’, the final apparent coverage would be around 200x since 100 concatenated genomes were used as input.

We then generated HiFi reads using pbccs v6.4.0 (https://ccs.how/) with ‘--all’ option and extracthifi v1.0.0 (https://github.com/PacificBiosciences/pbtk). We repeated this process ten times.

### Somatic TE insertion and excision calling

HiFi reads, either simulated or from PacBio sequencing, were mapped to the Tsu-0 genome using winnowmap v2.03 (Jain *et al*. 2022) (https://github.com/marbl/Winnowmap), using ‘meryl count k=15’, ‘meryl print greater-than distinct=0.9998’ and ‘winnowmap -Y -L -ax map-pb’. We used the sorted bam files as input for our somatic TE mobilization pipeline (https://github.com/aerilli/Somatic-transposition_met1). Candidate somatic insertions were initially recognized as reads with “Supplementary alignment”, *i.e.*, “chimeric” reads. Because the aligner sometimes does not generate supplementary alignments but describes insertions within CIGAR strings, we also made use of CIGAR strings, using an approach inspired by (Zhang *et al*. 2023). We discarded somatic insertions not overlapping with repeats (centromeres excluded) or close (≤50 bp) to assembly gaps.

In more detail, we first collected IDs of all reads that mapped to an annotated TE and then retained the IDs of reads that had a “Supplementary alignment” tag. Different sections of “chimeric” reads can be independently aligned to different regions of the genome and the “Supplementary alignment” tag is assigned to one of at least two independent alignments. We retained as reads from candidate insertions only those with at least 99% overlap of the annotated TE. Regions of a read with a mapping quality <50 were removed. In other words, the part of a “chimeric read” that maps to a TE should respect the TE boundaries and be fully contained in the TE. Additional formatting steps are required to generate a file with TE insertion site, name, and position of the original TE copy and the IDs of the reads supporting the same insertion site. All candidate insertions, including possible TSDs, were visually inspected.

In a second step, we identified reads including newly inserted sequences ≥500 bp, as deduced from their CIGAR strings, and with mapping quality ≥50. We extracted the insertion sequences and re-mapped them to the genome with minimap2 v2.24 (Li 2018) (https://lh3.github.io/minimap2/minimap2.html) (‘-ax map-pb’). We intersected the mappings with the TE annotation of the genome, followed by additional formatting steps. All candidate insertions except for the simulated ones were visually inspected.

From the two steps, we compiled a final curated list of confirmed somatic insertions for downstream analyses. We merged reads pointing to insertions within a 5bp range into the same event.

To retrieve somatic excisions, we retrieved the IDs of all the reads that had a deletion greater or equal than 500 bp, as described in the associated CIGAR string, filtered them by mapping quality (≥50) and intersected the filtered bam file with TE annotation. We extracted the coordinates of the deletions and checked for a 90% overlap with an annotated TE. All instances were visually inspected.

To minimize the annotation bias for un-annotated Class II TEs, we collected all reads from all *met1* plants that had “supplementary alignment” tag and indicated deletions ≥500 bp, and visually inspected them to assess whether we had missed any mobilizing TE.

### Curation of non-reference TE families

The borders of VANDAL21_3 were redefined based on which sequence was subject to *de novo* somatic insertion, and multiple VANDALs with different names, as annotated by EDTA, were merged.

The PAC annotation was manually added based on the sequence that was subject to *de novo* somatic insertion and excision.

TSDs were retrieved using a custom bash script (https://github.com/aerilli/Somatic-transposition_met1) agnostic of the TSD length expected for different TE families. We compared with blastn v2.9.0 with default parameters the sequence of each read with the flanking sequences. As mapping of the sequences immediately flanking a TE can be misleading depending on the sequence context of the insertion site, we collected all putative TSDs shorter than 15 bp and determined the most often found lengths (modes) for each family.

For CAC2A curation, we used steps inspired by (Goubert *et al*. 2022). In brief we used blastn to find additional copies in the genome, we extended the borders of the putative copies. To determine the exact borders, we inspected their multiple sequence alignment from mafft (Katoh and Standley 2013) https://github.com/GSLBiotech/mafft) ‘--adjustdirection --autò. We produced a *consensus* with cons (EMBOSS), obtained its ORFs via getorf (EMBOSS) and ran pfam_scan.pl to identify its domains. We also ran TE-Aid (https://github.com/clemgoub/TE-Aid) to obtain additional information. Conserved domains were extracted using DANTE v0.1.9 (https://github.com/kavonrtep/dante) and sequences were aligned with mafft ‘--autò. Multiple sequence alignments and iqtree v2.3.0 (Minh *et al*. 2020) (https://github.com/iqtree/iqtree2) (‘-bb 1000’) were used to build trees of the full sequences of CAC2A *consensus* and other CACTA/EnSpm (https://github.com/oushujun/TAIR12-TE) and of their extracted domains.

### Synteny of mobilizable TEs in different A. thaliana accessions

To determine whether TEs that are mobilizable in Col-0 are found in syntenic positions in Tsu-0,we collected TAIR10 positions of TEs that have been reported as mobilizable in Col-0 *met1* and *ddm1* mutants and in epiRILs from (Singer *et al*. 2001; Miura *et al*. 2001; Kato *et al*. 2004; Tsukahara *et al*. 2009; Fu *et al*. 2013; Catoni *et al*. 2019; Quadrana *et al*. 2019). We manually checked whether the presence of such TEs at the same positions in the Tsu-0 genome was supported in a whole-genome alignment (WGA) of Tsu-0 and Col-0 TAIR10 that we had generated with wfmash v0.12.6-0-g7205bf7 (https://github.com/waveygang/wfmash) (‘wfmash -a -s 10000 -n 1 -p 90 -k 19 -H 0.001 --hg-filter-ani-diff 30’) (Additional file 4). To determine whether Tsu-0 *met1* mobilizable TEs were found in syntenic positions in TAIR10, we extracted the TEs with 500 bp upstream and downstream flanking sequences and mapped these to TAIR10 using blastn with default parameters.

### Estimated distribution of detected somatic insertions

We used a bootstrap approach to estimate how many detected somatic insertions we should expect from our sample. We drew with replacement a sample of *n* reads, with *n* equaling the average reads per *met1* individual, from the pool of total reads of all the *met1* individuals, and tallied how often we retrieved a read that corresponded to an observed somatic insertion. We repeated this process 1,000 times.

### DNA methylation

#### BS-Seq

BS-Seq reads of the Tsu-0 *met1* line 2 (Srikant *et al*. 2022) were analyzed as before (Srikant *et al*. 2022) with minor differences: reads were trimmed using skewer v0.2.2 (Jiang *et al*. 2014) (https://github.com/relipmoc/skewer) and processed using Bismark suite v0.23.1 (Krueger and Andrews 2011) (https://github.com/FelixKrueger/Bismark). Reads were aligned to the newly assembled Tsu-0 genome, bam files were deduplicated, methylation calls in all contexts were extracted and the output was converted to BedGraph format for downstream analyses.

#### PacBio CG methylation calling

PacBio HiFi reads were extracted from the sequencing movies (PacBio raw format) of *met1* individuals via pbccs (‘--all --hifi-kinetics’) and extracthifi, demultiplexed with lima v2.6.0 (https://github.com/PacificBiosciences/pbbioconda) (‘--same --ccs --min-score 70 --min-scoring-regions 2 --min-ref-span 0.8 --peek-guess --split-bam-named’). We called cytosine methylation for every *met1* individual using jasmine v2.0.0 (https://github.com/PacificBiosciences/pbbioconda), mapped them to the Tsu-0 reference genome using the ‘align’ option of pbmm2 v1.10.0 (https://github.com/PacificBiosciences/pbbioconda), and generated site methylation probabilities with the ‘aligned_bam_to_cpg_scores’ tool v2.3.2 from pb-CpG-tools suite (https://github.com/PacificBiosciences/pb-CpG-tools) using the provided model.

### Chromatin accessibility

ATAC-Seq reads of Tsu-0 *met1* line 2 (Srikant *et al*. 2022) were analyzed as before (Srikant *et al*. 2022) with minor differences: reads were trimmed using skewer v0.2.2 and mapped to the newly assembled Tsu-0 genome using bowtie2 v2.3.5.1 (Langmead and Salzberg 2012) (https://github.com/BenLangmead/bowtie2) and duplicates were marked with picard ‘MarkDuplicates’ v2.27.5 (https://broadinstitute.github.io/picard), we then called ATAC-seq peaks independently for the three replicates with total gDNA as control with macs2 ‘callpeak’ v2.2.9.1 (Zhang *et al*. 2008) (https://github.com/macs3-project/MACS) (‘--nomodel --extsize 147 --keep-dup=all -g 1.35è). “Treatment” pile-up BedGraph output files were used for downstream analyses.

### RNA-expression

RNA-Seq data of Tsu-0 *met1* line 2 (Srikant *et al*. 2022) were mapped to the newly assembled Tsu-0 genome and transcripts per million (TPM) were extracted as before (Srikant *et al*. 2022).

To estimate expression of highly identical ÉVADÉ (EVD) copies, we mapped RNA-Seq reads of Tsu-0 wild-type and *met1* line 1,2 (Srikant *et al*. 2022) with bowtie2 ‘--mp 13 --rdg 8,5 --rfg 8,5 --very-sensitivè to the newly assembled Tsu-0 genome. We estimated the expression of EVD_1,2,3 as the summation of the mapping qualities of each mapping read divided by the maximum mapping quality (60). In other words, a mapped read was counted as 1 only if its mapping quality was 60 and it could be counted as 0.5 if, for instance, its mapping quality was 30. We then normalized the resulting values by raw library size.

### Inspection of candidate reads from alternative transposition events

We collected mapped HiFi reads from the *met1* individuals that overlapped the VANDAL21 hypermutable region, and the up- and down- stream 20 kb flanking regions. We extracted reads with supplementary alignments, *i.e.*, reads that were “chimeric” (SAM flag ‘2048’) and had deletions ≥100 bp according to their CIGAR string. We then inspected the reads in IGV and classified them according to the structural variation that they supported. When one portion of a “chimeric” read mapped outside the hypermutable region or its flanking 20 kb regions, we recorded its break-end (BND) coordinate.

### Insertion preferences

#### Genomic features

We intersected the final curated list of confirmed somatic insertions, VANDAL21 BND coordinates and 10,000 randomly generated insertion sites with the annotations of genes, TEs and repeats, after TEs overlapping repeats and genes overlapping TEs had been removed. Insertions not overlapping with any of these features were labeled as “intergenic”. When an insertion occurred within a gene, we determined whether this gene was present in a list of lethal genes (Lloyd *et al*. 2015).

We used a bootstrap approach to randomly resample the features and essentiality of genes overlapping insertion sites to estimate the true distribution of such insertions (https://github.com/aerilli/Somatic-transposition_met1).

We also determined the feature that is the closest downstream and upstream (gene, TE), or just closest (repeat), to any somatic or random insertion site, or to any VANDAL21 BND.

#### Methylation bias

To evaluate residual methylation of sites in *met1* mutant where TEs somatically inserted, we assessed the distribution of CG, CHG, CGG (BS-seq) or CG (HiFi reads) methylation values at confirmed somatic insertion sites, and up- and down- stream 25 bp.

#### ATAC-Seq analysis

To evaluate the accessibility of sites where TEs somatically inserted, we assessed the distribution of the average of three replicates of “Treatment” pile-up values at confirmed somatic insertion sites, and up- and down- stream 25 bp. Outliers were excluded from plotted distributions.

#### RNA-Seq analysis

We evaluated the expression (TPM) of genes overlapping with the position of the confirmed somatic insertions – *i.e.,* the genes that were somatically disrupted by a TE insertion – and of 10,000 randomly generated insertion sites. Outliers were excluded from plotted distributions.

### Circular DNA detection

The aerial parts of one Tsu-0 *met1* mutant (line 2, (Srikant *et al*. 2022)) were finely ground with mortar and pestle in liquid nitrogen. DNA extraction was carried out with a standard CTAB protocol, including one hour incubation in CTAB at 65°C followed by chloroform:IAA (24:1) precipitation. We performed genotyping via PCR amplification of *MET1* exon 7 (Srikant *et al*. 2022) followed by digestion with Hpy188I (NEB), which selectively recognizes the “T” insertion of the mutant allele. We used the DNA of the mutant as input for rolling circle amplification (RCA) as published (dx.doi.org/10.17504/protocols.io.bz8np9ve), with EquiPhi29 DNA Polymerase (ThermoFisher) and a reaction temperature of 42°C. We detected circular DNA by PCR amplification with outward facing primers (Supplementary Table 2) using either *met1* genomic DNA or the RCA reaction as input. We cloned the PCR bands in p-GEM-T Easy vectors (Promega) and confirmed their identity via Sanger-sequencing.

### Softwares, visualization and statistics

Statistical analyses and their visualization used RStudio Pro 2024.04.2 with R v4.4.0 and jupyterlab v4.0.7 with Python v3.10.13. Read mappings were visualized in IGV v2.16. Multiple sequence alignments were visualized in AliView v1.28.

## Supplementary Information

**Additional file 1: Fig. S1-S12, Table S1-S2**.

**Additional file 2: Dataset S1.** List of curated transposable element insertions and excisions in ten Tsu-0 *met1* mutants.

**Additional file 3: Dataset S2**. List of curated ‘out-region’ BND coordinates at Tsu-0 hypermutable VANDAL21 region.

**Additional file 4.** Visual inspection of somatic insertion and excision events. Visual inspection of syntenic TEs.

## Acknowledgements

We acknowledge Haijun Liu and Magnus Nordborg for contributing to the initial idea of the project. We thank for their valuable feedback Pierre Baduel, Zhigui Bao, Ilja Bezrukov, Michael Borg, Mireia Bueno-Merino, Adrián Contreras-Garrido, Rory J. Craig, Erica Dinatale, Hajk-Georg Drost, Alejandra Duque-Jaramillo, Li He, Miriam Lucke, Arturo Marí-Ordoñez, Kevin Murray, Gal Ofir, Luisa Pallares, Leandro Quadrana, Sheila Roitman, Rebecca Schwab, Gautam Shirsekar, Thanvi Srikant, Yueqi Tao, Luisa Teasdale, Marja Timmermans, and Wenfei Xian. A.M is a member of the International Max Planck Research School ’From Molecules to Organisms’. This study was supported by the Novozymes Prize of the Novo Nordisk Foundation and the Max Planck Society (D.W.).

## Authors’ contributions

A.M. and D.W conceived the project. A.M. performed all the experimental work and bioinformatic analyses, except for HiFi reads generation and genome assembly (F.A.R.), and additional TE annotation (S.S.). A.M. and D.W. interpreted the data and wrote the manuscript. All authors read and approved the final manuscript.

## Availability of data and materials

The PacBio HiFi reads generated for this study are available in the European Nucleotide Archive (ENA, https://www.ebi.ac.uk/ena/browser/home) under project name PRJEB85646. Additional files and scripts are available in GitHub at https://github.com/aerilli/Somatic-transposition_met1. Reads from ATAC-Seq, RNA-Seq and BS-Seq experiments were obtained for *met1* mutant Line 2 (“Tsu-0_P2”) from PRJEB53354 (Srikant *et al*. 2022; https://www.ebi.ac.uk/ena/browser/view/PRJEB54034), PRJEB54034 (Srikant *et al*. 2022; https://www.ebi.ac.uk/ena/browser/view/PRJEB53354) and PRJEB54036 (Srikant *et al*. 2022; https://www.ebi.ac.uk/ena/browser/view/PRJEB54036) respectively.

## Ethics approval and consent to participate

Not applicable.

## References

1. Alonge M., L. Lebeigle, M. Kirsche, K. Jenike, S. Ou, et al., 2022 Automated assembly scaffolding using RagTag elevates a new tomato system for high-throughput genome editing. Genome Biol. 23: 258.

2. Baduel P., B. Leduque, A. Ignace, I. Gy, J. Gil Jr, et al., 2021 Genetic and environmental modulation of transposition shapes the evolutionary potential of Arabidopsis thaliana. Genome Biol. 22: 138.

3. Baid G., D. E. Cook, K. Shafin, T. Yun, F. Llinares-López, et al., 2023 DeepConsensus improves the accuracy of sequences with a gap-aware sequence transformer. Nat. Biotechnol. 41: 232–238.

4. Baillie J. K., M. W. Barnett, K. R. Upton, D. J. Gerhardt, T. A. Richmond, et al., 2011 Somatic retrotransposition alters the genetic landscape of the human brain. Nature 479: 534–537.

5. Berger F., and D. Twell, 2011 Germline specification and function in plants. Annu. Rev. Plant Biol. 62: 461–484.

6. Borges F., J.-S. Parent, F. van Ex, P. Wolff, G. Martínez, et al., 2018 Transposon-derived small RNAs triggered by miR845 mediate genome dosage response in Arabidopsis. Nat. Genet. 50: 186–192.

7. Bourque G., K. H. Burns, M. Gehring, V. Gorbunova, A. Seluanov, et al., 2018 Ten things you should know about transposable elements. Genome Biol. 19: 199.

8. Bowman J. L., K. Sakakibara, C. Furumizu, and T. Dierschke, 2016 Evolution in the cycles of life. Annu. Rev. Genet. 50: 133–154.

9. Cameron D. L., L. Di Stefano, and A. T. Papenfuss, 2019 Comprehensive evaluation and characterisation of short read general-purpose structural variant calling software. Nat. Commun. 10: 3240.

10. Catoni M., T. Jonesman, E. Cerruti, and J. Paszkowski, 2019 Mobilization of Pack-CACTA transposons in Arabidopsis suggests the mechanism of gene shuffling. Nucleic Acids Res. 47: 1311–1320.

11. Chang C., D. J. Pagano, D. D. Lowe, and S. Kennedy, 2022 DNA repair strategy sets transposon mobilization rates in Caenorhabditis elegans. bioRxiv 2022.08.01.502254.

12. Cheng H., G. T. Concepcion, X. Feng, H. Zhang, and H. Li, 2021 Haplotype-resolved de novo assembly using phased assembly graphs with hifiasm. Nat. Methods 18: 170–175.

13. Coleman-Derr D., and D. Zilberman, 2012 Deposition of histone variant H2A.Z within gene bodies regulates responsive genes. PLoS Genet. 8: e1002988.

14. Creasey K. M., J. Zhai, F. Borges, F. Van Ex, M. Regulski, et al., 2014 miRNAs trigger widespread epigenetically activated siRNAs from transposons in Arabidopsis. Nature 508: 411–415.

15. D’Costa A. V., and J. T. Simpson, 2023 Somrit: The somatic retrotransposon insertion toolkit. bioRxiv.

16. Deleris A., H. Stroud, Y. Bernatavichute, E. Johnson, G. Klein, et al., 2012 Loss of the DNA methyltransferase MET1 Induces H3K9 hypermethylation at PcG target genes and redistribution of H3K27 trimethylation to transposons in Arabidopsis thaliana. PLoS Genet. 8: e1003062.

17. Déléris A., F. Berger, and S. Duharcourt, 2021 Role of Polycomb in the control of transposable elements. Trends Genet. 37: 882–889.

18. Döring H.-P., and P. Starlinger, 1984 Barbara McClintock’s controlling elements: Now at the DNA level. Cell 39: 253–259.

19. Drost H.-G., and D. H. Sanchez, 2019 Becoming a selfish clan: Recombination associated to reverse-transcription in LTR retrotransposons. Genome Biol. Evol. 11: 3382–3392.

20. Dubin M. J., P. Zhang, D. Meng, M.-S. Remigereau, E. J. Osborne, et al., 2015 DNA methylation in Arabidopsis has a genetic basis and shows evidence of local adaptation. Elife 4: e05255.

21. Feng S., S. J. Cokus, V. Schubert, J. Zhai, M. Pellegrini, et al., 2014 Genome-wide Hi-C analyses in wild-type and mutants reveal high-resolution chromatin interactions in Arabidopsis. Mol. Cell 55: 694–707.

22. Fransz P., G. Linc, C.-R. Lee, S. A. Aflitos, J. R. Lasky, et al., 2016 Molecular, genetic and evolutionary analysis of a paracentric inversion in Arabidopsis thaliana. Plant J. 88: 159–178.

23. Fu Y., A. Kawabe, M. Etcheverry, T. Ito, A. Toyoda, et al., 2013 Mobilization of a plant transposon by expression of the transposon-encoded anti-silencing factor. EMBO J. 32: 2407–2417.

24. Gerdes P., S. M. Lim, A. D. Ewing, M. R. Larcombe, D. Chan, et al., 2022 Retrotransposon instability dominates the acquired mutation landscape of mouse induced pluripotent stem cells. Nat. Commun. 13: 7470.

25. Gisby J. S., and M. Catoni, 2022 The widespread nature of Pack-TYPE transposons reveals their importance for plant genome evolution. PLoS Genet. 18: e1010078.

26. Gómez-Zambrano Á., W. Merini, and M. Calonje, 2019 The repressive role of Arabidopsis H2A.Z in transcriptional regulation depends on AtBMI1 activity. Nat. Commun. 10: 2828.

27. Gonzalez N., S. De Bodt, R. Sulpice, Y. Jikumaru, E. Chae, et al., 2010 Increased leaf size: different means to an end. Plant Physiol. 153: 1261–1279.

28. Goubert C., R. J. Craig, A. F. Bilat, V. Peona, A. A. Vogan, et al., 2022 A beginner’s guide to manual curation of transposable elements. Mob. DNA 13: 7.

29. Gray Y. H., 2000 It takes two transposons to tango: transposable-element-mediated chromosomal rearrangements. Trends Genet. 16: 461–468.

30. Haig D., 2016 Transposable elements: Self-seekers of the germline, team-players of the soma. Bioessays 38: 1158–1166.

31. He L., H. Huang, M. Bradai, C. Zhao, Y. You, et al., 2022 DNA methylation-free Arabidopsis reveals crucial roles of DNA methylation in regulating gene expression and development. Nat. Commun. 13: 1335.

32. Hure V., F. Piron-Prunier, T. Yehouessi, C. Vitte, A. E. Kornienko, et al., 2025 Alternative silencing states of transposable elements in Arabidopsis associated with H3K27me3. Genome Biol. 26: 11.

33. Jain C., A. Rhie, N. F. Hansen, S. Koren, and A. M. Phillippy, 2022 Long-read mapping to repetitive reference sequences using Winnowmap2. Nat. Methods 19: 705–710.

34. Jiang N., Z. Bao, X. Zhang, S. R. Eddy, and S. R. Wessler, 2004 Pack-MULE transposable elements mediate gene evolution in plants. Nature 431: 569–573.

35. Jiang N., A. A. Ferguson, R. K. Slotkin, and D. Lisch, 2011 Pack-Mutator-like transposable elements (Pack-MULEs) induce directional modification of genes through biased insertion and DNA acquisition. Proc. Natl. Acad. Sci. U. S. A. 108: 1537–1542.

36. Jiang H., R. Lei, S.-W. Ding, and S. Zhu, 2014 Skewer: a fast and accurate adapter trimmer for next-generation sequencing paired-end reads. BMC Bioinformatics 15: 182.

37. Kato M., K. Takashima, and T. Kakutani, 2004 Epigenetic control of CACTA transposon mobility in Arabidopsis thaliana. Genetics 168: 961–969.

38. Katoh K., and D. M. Standley, 2013 MAFFT multiple sequence alignment software version 7: improvements in performance and usability. Mol. Biol. Evol. 30: 772–780.

39. Kazazian H. H. Jr, 2011 Mobile DNA transposition in somatic cells. BMC Biol. 9: 62. Krueger F., and S. R. Andrews, 2011 Bismark: a flexible aligner and methylation caller for Bisulfite-Seq applications. Bioinformatics 27: 1571–1572.

40. Langmead B., and S. L. Salzberg, 2012 Fast gapped-read alignment with Bowtie 2. Nat. Methods 9: 357–359.

41. Lee S. C., E. Ernst, B. Berube, F. Borges, J.-S. Parent, et al., 2020 Arabidopsis retrotransposon virus-like particles and their regulation by epigenetically activated small RNA. Genome Res. 30: 576–588.

42. Lee S. C., D. W. Adams, J. J. Ipsaro, J. Cahn, J. Lynn, et al., 2023 Chromatin remodeling of histone H3 variants by DDM1 underlies epigenetic inheritance of DNA methylation. Cell 186: 4100–4116.e15.

43. Li W., L. Prazak, N. Chatterjee, S. Grüninger, L. Krug, et al., 2013 Activation of transposable elements during aging and neuronal decline in Drosophila. Nat. Neurosci. 16: 529–531.

44. Li H., 2018 Minimap2: pairwise alignment for nucleotide sequences. Bioinformatics 34: 3094–3100.

45. Liang W., J. Li, L. Sun, Y. Liu, Z. Lan, et al., 2022 Deciphering the synergistic and redundant roles of CG and non-CG DNA methylation in plant development and transposable element silencing. New Phytol. 233: 722–737.

46. Lippman Z., B. May, C. Yordan, T. Singer, and R. Martienssen, 2003 Distinct mechanisms determine transposon inheritance and methylation via small interfering RNA and histone modification. PLoS Biol. 1: e67.

47. Lloyd J. P., A. E. Seddon, G. D. Moghe, M. C. Simenc, and S.-H. Shiu, 2015 Characteristics of plant essential genes allow for within- and between-species prediction of lethal mutant phenotypes. Plant Cell 27: 2133–2147.

48. López-Cortegano E., R. J. Craig, J. Chebib, E. J. Balogun, and P. D. Keightley, 2023 Rates and spectra of de novo structural mutations in Chlamydomonas reinhardtii. Genome Res. 33: 45–60.

49. Manni M., M. R. Berkeley, M. Seppey, F. A. Simão, and E. M. Zdobnov, 2021 BUSCO update: Novel and streamlined workflows along with broader and deeper phylogenetic coverage for scoring of eukaryotic, prokaryotic, and viral genomes. Mol. Biol. Evol. 38: 4647–4654.

50. Marí-Ordóñez A., A. Marchais, M. Etcheverry, A. Martin, V. Colot, et al., 2013 Reconstructing de novo silencing of an active plant retrotransposon. Nat. Genet. 45: 1029–1039.

51. Mathieu O., J. Reinders, M. Caikovski, C. Smathajitt, and J. Paszkowski, 2007 Transgenerational stability of the Arabidopsis epigenome is coordinated by CG methylation. Cell 130: 851–862.

52. McClintock B., 1950 The origin and behavior of mutable loci in maize. Proc. Natl. Acad. Sci. U. S. A. 36: 344–355.

53. Merkulov P., E. Egorova, and I. Kirov, 2023 Composition and structure of Arabidopsis thaliana extrachromosomal circular DNAs revealed by nanopore sequencing. Plants 12: 2178.

54. Mikheenko A., V. Saveliev, P. Hirsch, and A. Gurevich, 2023 WebQUAST: online evaluation of genome assemblies. Nucleic Acids Res. 51: W601–W606.

55. Minh B. Q., H. A. Schmidt, O. Chernomor, D. Schrempf, M. D. Woodhams, et al., 2020 IQ-TREE 2: New models and efficient methods for phylogenetic inference in the genomic era. Mol. Biol. Evol. 37: 1530–1534.

56. Mirouze M., J. Reinders, E. Bucher, T. Nishimura, K. Schneeberger, et al., 2009 Selective epigenetic control of retrotransposition in Arabidopsis. Nature 461: 427–430.

57. Miura A., S. Yonebayashi, K. Watanabe, T. Toyama, H. Shimada, et al., 2001 Mobilization of transposons by a mutation abolishing full DNA methylation in Arabidopsis. Nature 411: 212–214.

58. Muotri A. R., V. T. Chu, M. C. N. Marchetto, W. Deng, J. V. Moran, et al., 2005 Somatic mosaicism in neuronal precursor cells mediated by L1 retrotransposition. Nature 435: 903–910.

59. Naish M., M. Alonge, P. Wlodzimierz, A. J. Tock, B. W. Abramson, et al., 2021 The genetic and epigenetic landscape of the Arabidopsis centromeres. Science 374: eabi7489.

60. Neumann P., P. Novák, N. Hoštáková, and J. Macas, 2019 Systematic survey of plant LTR-retrotransposons elucidates phylogenetic relationships of their polyprotein domains and provides a reference for element classification. Mob. DNA 10: 1.

61. Oberlin S., R. Rajeswaran, M. Trasser, V. Barragán-Borrero, M. A. Schon, et al., 2022 Innate, translation-dependent silencing of an invasive transposon in Arabidopsis. EMBO Rep. 23: e53400.

62. Ono Y., M. Hamada, and K. Asai, 2022 PBSIM3: a simulator for all types of PacBio and ONT long reads. NAR Genom. Bioinform. 4: lqac092.

63. Osakabe A., B. Jamge, E. Axelsson, S. A. Montgomery, S. Akimcheva, et al., 2021 The chromatin remodeler DDM1 prevents transposon mobility through deposition of histone variant H2A.W. Nat. Cell Biol. 23: 391–400.

64. Osakabe A., Y. Takizawa, N. Horikoshi, S. Hatazawa, L. Negishi, et al., 2024 Molecular and structural basis of the chromatin remodeling activity by Arabidopsis DDM1. Nat. Commun. 15: 5187.

65. Ou S., W. Su, Y. Liao, K. Chougule, J. R. A. Agda, et al., 2019 Benchmarking transposable element annotation methods for creation of a streamlined, comprehensive pipeline. Genome Biol. 20: 275.

66. Ou S., W. Su, Y. Liao, K. Chougule, J. R. A. Agda, et al., 2022 Author Correction: Benchmarking transposable element annotation methods for creation of a streamlined, comprehensive pipeline. Genome Biol. 23: 76.

67. Pertea G., and M. Pertea, 2020 GFF utilities: GffRead and GffCompare. F1000Res. 9: 304.

68. Qian J., H. Xue, S. Ou, J. Storer, L. Fürtauer, et al., 2024 TEtrimmer: a novel tool to automate the manual curation of transposable elements. bioRxiv 2024.06.27.600963.

69. Quadrana L., M. Etcheverry, A. Gilly, E. Caillieux, M.-A. Madoui, et al., 2019 Transposition favors the generation of large effect mutations that may facilitate rapid adaption. Nat. Commun. 10: 3421.

70. Quesneville H., 2020 Twenty years of transposable element analysis in the Arabidopsis thaliana genome. Mob. DNA 11: 28.

71. Rabanal F. A., M. Gräff, C. Lanz, K. Fritschi, V. Llaca, et al., 2022 Pushing the limits of HiFi assemblies reveals centromere diversity between two Arabidopsis thaliana genomes. Nucleic Acids Res. 50: 12309–12327.

72. Reinders J., B. B. H. Wulff, M. Mirouze, A. Marí-Ordóñez, M. Dapp, et al., 2009 Compromised stability of DNA methylation and transposon immobilization in mosaic Arabidopsis epigenomes. Genes Dev. 23: 939–950.

73. Rigal M., C. Becker, T. Pélissier, R. Pogorelcnik, J. Devos, et al., 2016 Epigenome confrontation triggers immediate reprogramming of DNA methylation and transposon silencing in Arabidopsis thaliana F1 epihybrids. Proc. Natl. Acad. Sci. U. S. A. 113: E2083–92.

74. Sasaki T., K. Ro, E. Caillieux, R. Manabe, G. Bohl-Viallefond, et al., 2022a Fast co-evolution of anti-silencing systems shapes the invasiveness of Mu-like DNA transposons in eudicots. EMBO J. 41: e110070.

75. Sasaki E., J. Gunis, I. Reichardt-Gomez, V. Nizhynska, and M. Nordborg, 2022b Conditional GWAS of non-CG transposon methylation in Arabidopsis thaliana reveals major polymorphisms in five genes. PLoS Genet. 18: e1010345.

76. Sharma S. P., and T. Peterson, 2023 Complex chromosomal rearrangements induced by transposons in maize. Genetics 223: iyac124.

77. Shen W., B. Sipos, and L. Zhao, 2024 SeqKit2: A Swiss army knife for sequence and alignment processing. Imeta 3: e191.

78. Shumate A., and S. L. Salzberg, 2021 Liftoff: accurate mapping of gene annotations. Bioinformatics 37: 1639–1643.

79. Sierra P., and R. Durbin, 2024 Identification of transposable element families from pangenome polymorphisms. Mob. DNA 15: 13.

80. Singer T., C. Yordan, and R. A. Martienssen, 2001 Robertson’s Mutator transposons in A. thaliana are regulated by the chromatin-remodeling gene Decrease in DNA Methylation (DDM1). Genes Dev. 15: 591–602.

81. Siudeja K., M. van den Beek, N. Riddiford, B. Boumard, A. Wurmser, et al., 2021 Unraveling the features of somatic transposition in the *Drosophila* intestine. EMBO J. 40: e106388.

82. Smit A. F. A., R. Hubley, and P. Green, 2013-2015 RepeatMasker Open-4.0. http://www.repeatmasker.org.

83. Srikant T., W. Yuan, K. W. Berendzen, A. Contreras-Garrido, H. Drost, et al., 2022; https://www.ebi.ac.uk/ena/browser/view/PRJEB54034 ATAC-Seq data generated for Srikant et al., 2022. Eur Nucleotide Arch.

84. Srikant T., W. Yuan, K. W. Berendzen, A. Contreras-Garrido, H. Drost, et al., 2022; https://www.ebi.ac.uk/ena/browser/view/PRJEB53354 RNA-Seq data generated for Srikant et al., 2022. Eur Nucleotide Arch.

85. Srikant T., W. Yuan, K. W. Berendzen, A. Contreras-Garrido, H. Drost, et al., 2022; https://www.ebi.ac.uk/ena/browser/view/PRJEB54036 Bisulfite-Seq data generated for Srikant et al., 2022. Eur Nucleotide Arch.

86. Srikant T., W. Yuan, K. W. Berendzen, A. Contreras-Garrido, H.-G. Drost, et al., 2022 Canalization of genome-wide transcriptional activity in Arabidopsis thaliana accessions by MET1-dependent CG methylation. Genome Biol. 23: 263.

87. Stroud H., M. V. C. Greenberg, S. Feng, Y. V. Bernatavichute, and S. E. Jacobsen, 2013 Comprehensive analysis of silencing mutants reveals complex regulation of the Arabidopsis methylome. Cell 152: 352–364.

88. Su W., S. P. Sharma, and T. Peterson, 2018 Evolutionary Impacts of Alternative Transposition, pp. 113–130 in Origin and Evolution of Biodiversity, Springer International Publishing, Cham.

89. Su W., T. Zuo, and T. Peterson, 2020 Ectopic expression of a maize gene is induced by composite insertions generated through alternative transposition. Genetics 216: 1039–1049.

90. To T. K., C. Yamasaki, S. Oda, S. Tominaga, A. Kobayashi, et al., 2022 Local and global crosstalk among heterochromatin marks drives DNA methylome patterning in Arabidopsis. Nat. Commun. 13: 861.

91. Tsukahara S., A. Kobayashi, A. Kawabe, O. Mathieu, A. Miura, et al., 2009 Bursts of retrotransposition reproduced in Arabidopsis. Nature 461: 423–426.

92. Vendrell-Mir P., B. Leduque, and L. Quadrana, 2024 Ultra-sensitive detection of transposon insertions across multiple families by transposable element display sequencing. bioRxiv 2024.08.21.608910.

93. Wang D., C. Yu, T. Zuo, J. Zhang, D. F. Weber, et al., 2015 Alternative transposition generates new chimeric genes and segmental duplications at the maize p1 locus. Genetics 201: 925–935.

94. Wells J. N., and C. Feschotte, 2020 A Field Guide to Eukaryotic Transposable Elements. Annu. Rev. Genet. 54: 539–561.

95. Wicker T., F. Sabot, A. Hua-Van, J. L. Bennetzen, P. Capy, et al., 2007 A unified classification system for eukaryotic transposable elements. Nat. Rev. Genet. 8: 973–982.

96. Zhang J., and T. Peterson, 2004 Transposition of reversed Ac element ends generates chromosome rearrangements in maize. Genetics 167: 1929–1937.

97. Zhang J., and T. Peterson, 2005 A segmental deletion series generated by sister-chromatid transposition of Ac transposable elements in maize. Genetics 171: 333–344.

98. Zhang Y., T. Liu, C. A. Meyer, J. Eeckhoute, D. S. Johnson, et al., 2008 Model-based analysis of ChIP-Seq (MACS). Genome Biol. 9: R137.

99. Zhang J., C. Yu, V. Pulletikurti, J. Lamb, T. Danilova, et al., 2009 Alternative Ac/Ds transposition induces major chromosomal rearrangements in maize. Genes Dev. 23: 755–765.

100. Zhang J., T. Zuo, D. Wang, and T. Peterson, 2014 Transposition-mediated DNA re-replication in maize. Elife 3: e03724.

101. Zhang H., Z. Lang, and J.-K. Zhu, 2018 Dynamics and function of DNA methylation in plants. Nat. Rev. Mol. Cell Biol. 19: 489–506.

102. Zhang R.-G., G.-Y. Li, X.-L. Wang, J. Dainat, Z.-X. Wang, et al., 2022 TEsorter: an accurate and fast method to classify LTR-retrotransposons in plant genomes. Hortic. Res. 9.

103. Zhang P., A. Mbodj, A. Soundiramourtty, C. Llauro, A. Ghesquière, et al., 2023 Extrachromosomal circular DNA and structural variants highlight genome instability in Arabidopsis epigenetic mutants. Nat. Commun. 14: 5236.

104. Zhong Z., S. Feng, S. H. Duttke, M. E. Potok, Y. Zhang, et al., 2021 DNA methylation-linked chromatin accessibility affects genomic architecture in Arabidopsis. Proc. Natl. Acad. Sci. U. S. A. 118: e2023347118.

