## Additional File 1 for "Long-read detection of transposable element mobilization in the soma of hypomethylated *Arabidopsis thaliana* individuals"

a)

| scaffold sum (bp) | N50 (bp) | L50 | N90 (bp) | L90 | Gaps | contigs sum (bp) |
|---|---|---|---|---|---|---|
| 134463716 | 26159155 | 3 | 22155302 | 5 | 4 | 146349704 |

| n contigs | average contig length (bp) | largest contig (bp) | contig N50 | contig L50 | contig N90 | contig L90 |
|---|---|---|---|---|---|---|
| 192 | 762238.04 | 32857596 | 22873944 | 3 | 4380539 | 8 |

b)

| Busco completeness | S | D | F | I | M | N |
|---|---|---|---|---|---|---|
| **Assembly** | 99.06%, 421 | 0.71%, 3 | 0.00%, 0 | 0.00%, 0 | 0.24%, 1 | 425 |
| **Gene annotation** | 65.65%, 279 | 33.8%, 144 | 0.24%, 1 | - | 0.24%, 1 | 425 |

**lineage:viridiplantae_odb10**

**Supplementary Table 1**. **Tsu-0 genome assembly and gene annotation. a** Statistics of genome assembly and reference-based scaffolds. **b** Busco completeness of genome assembly and lift-over gene annotation.

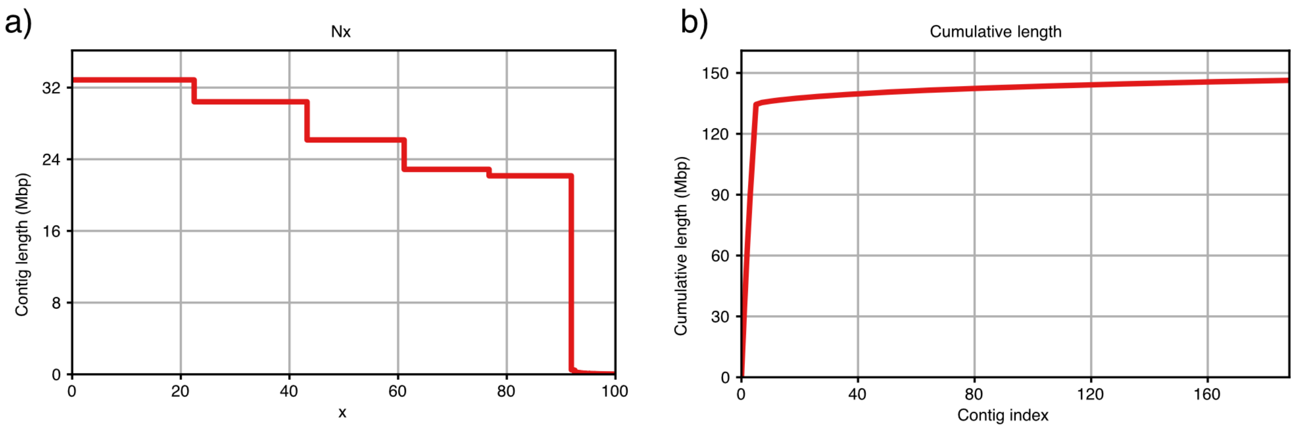


**Supplementary Figure 1**. **Contiguity of Tsu-0 genome assembly. a** Contiguity (Nx) as a function of contig length. **b** Cumulative length of contigs. Output from QUAST (https://github.com/ablab/quast).

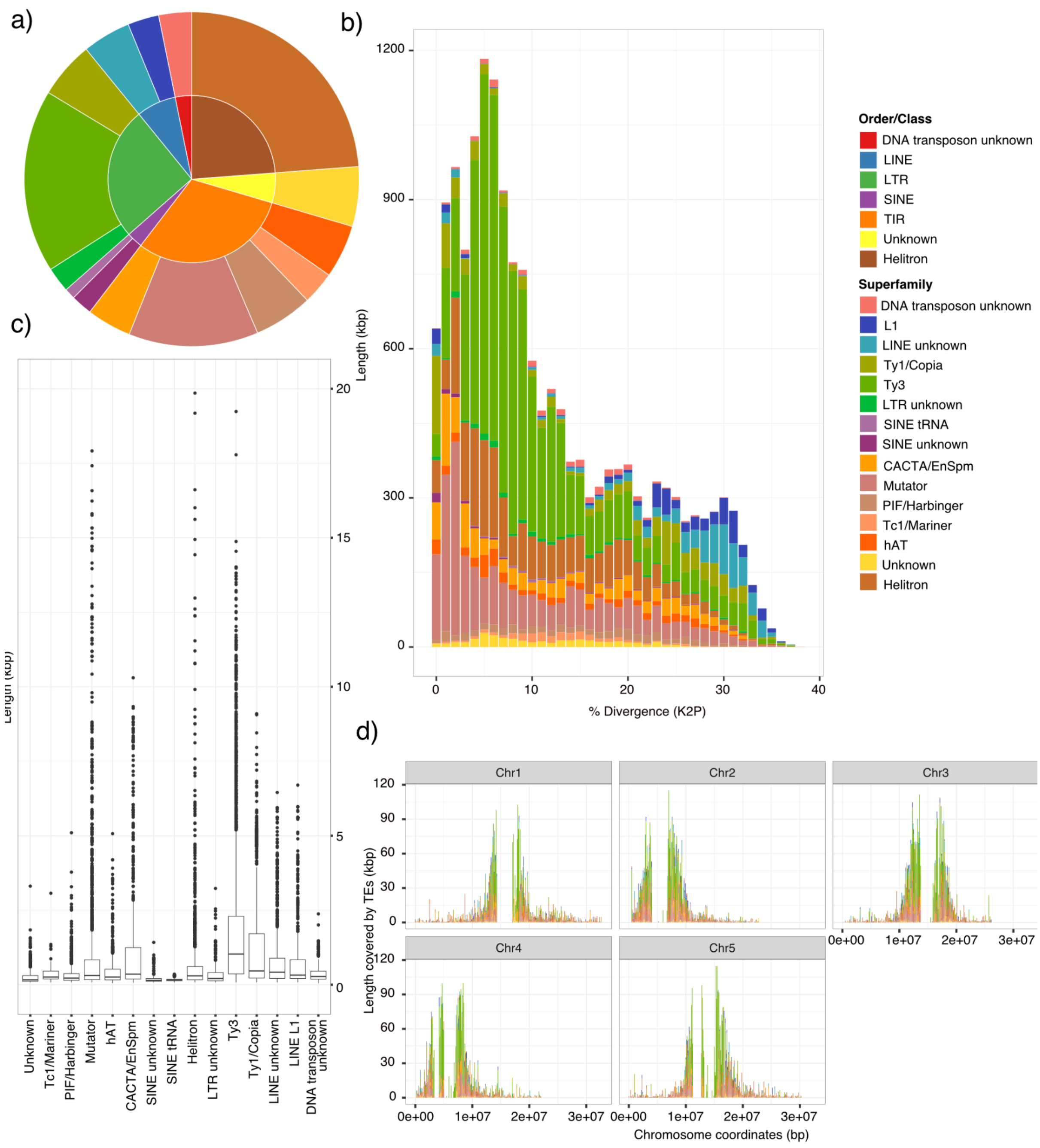


**Supplementary Figure 2**. **TE annotation of Tsu-0 genome. a** Mobilome composition of the Tsu-0 genome. **b** Divergence (% Kimura-2-Parameter, K2P) of TE copies within the genome, colored by superfamily. **d** Length distribution of TE insertions of each type. **c** Distribution of TE copies along chromosomes.

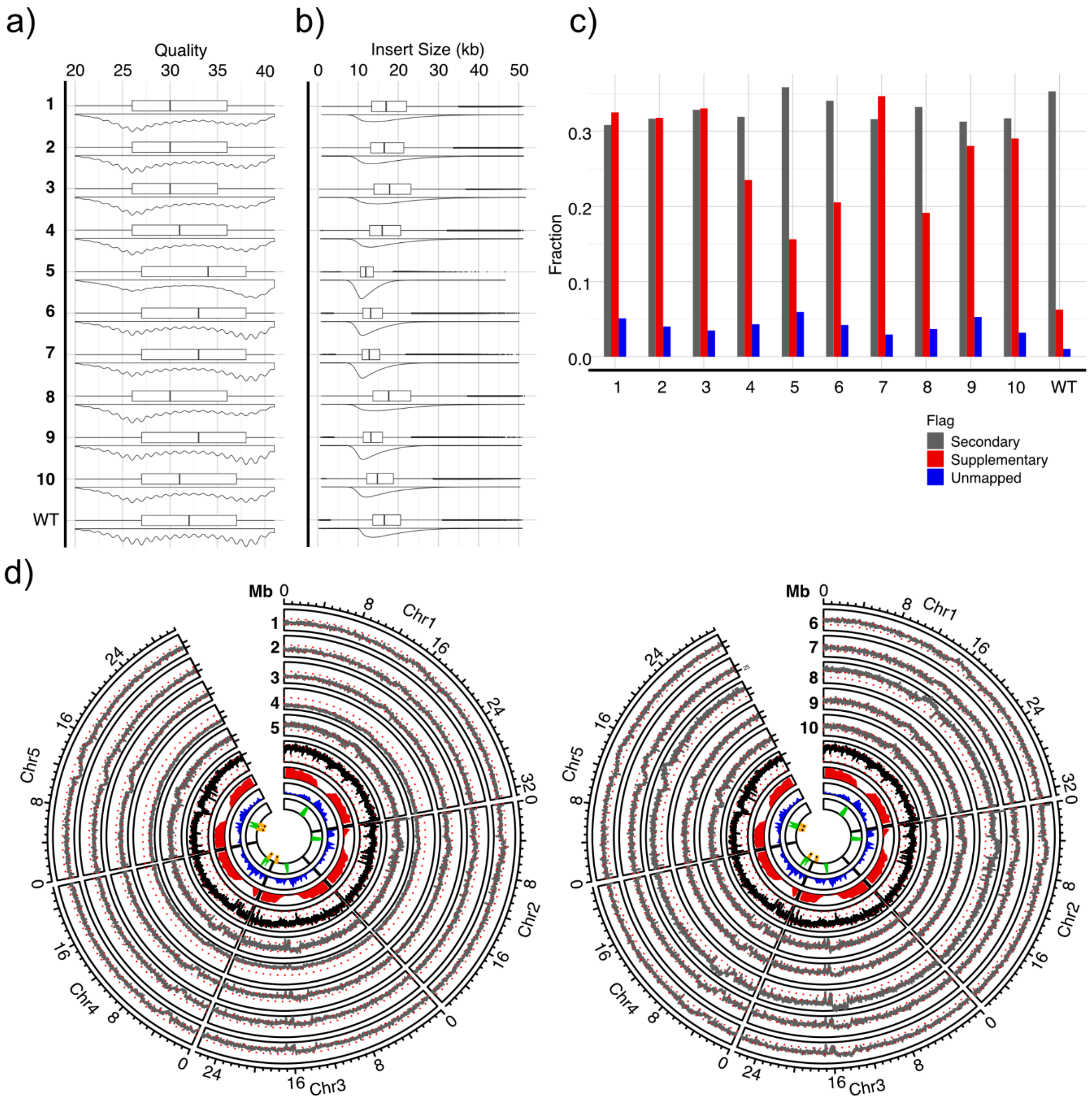


**Supplementary Figure 3**. **Sequencing statistics of *met1* individuals. a** Sequencing quality scores and **b** insert sizes of Tsu-0 wild-type pool and Tsu-0 *met1* individuals (1-10). **c** Fraction of alignment flags for reads from *met1* individuals (1-10) and wild-type pool. **d** Circos plots showing the coverage of reads from *met1* individuals aligned to the Tsu-0 wild-type genome (left: *met1_01-05*; right: *met1_06-10*) (grey), and of reads from wild-type pool (black), gene annotation (red), TE annotation (blue), repeat annotation (rDNA, centromere, telomere, nuclear insertions of organellar DNA) (green) and scaffold gaps (yellow). Red dotted lines indicate coverages of 25x, 50x, 100x.

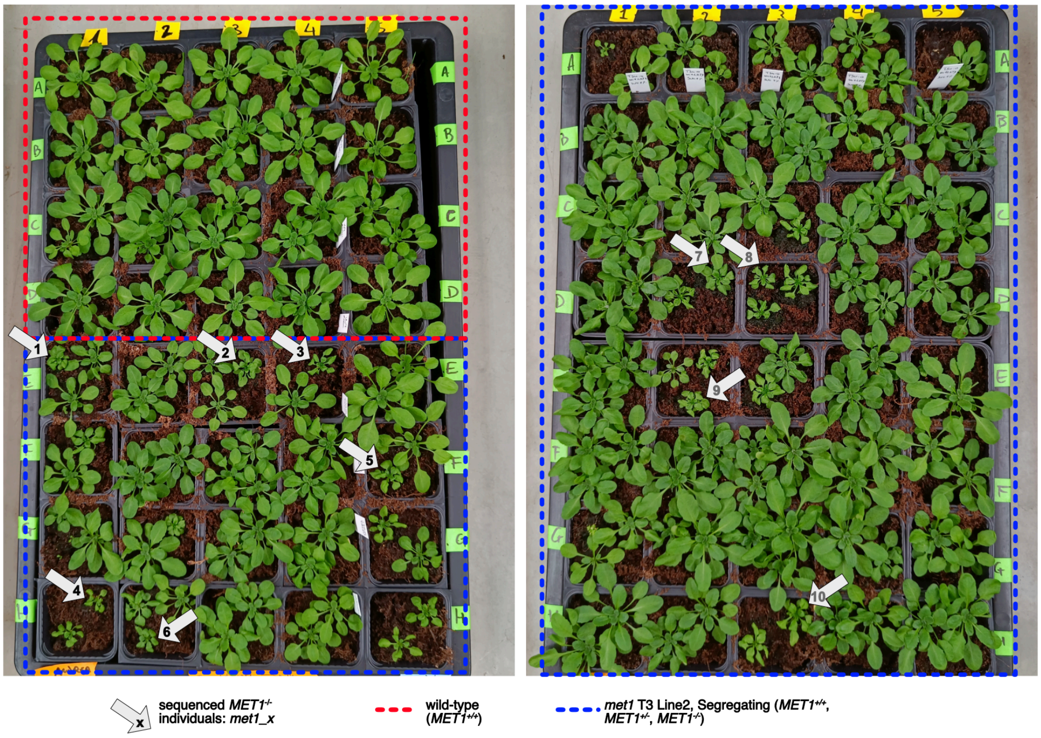


**Supplementary Figure 4**. **Morphology of *met1* individuals at the time of harvest.** A pure population of wild types (red) and a population of segregating wild-type, *met1* heterozygous and *met1* homozygous individuals (blue). Sampled *met1* homozygotes are indicated with a grey arrow.

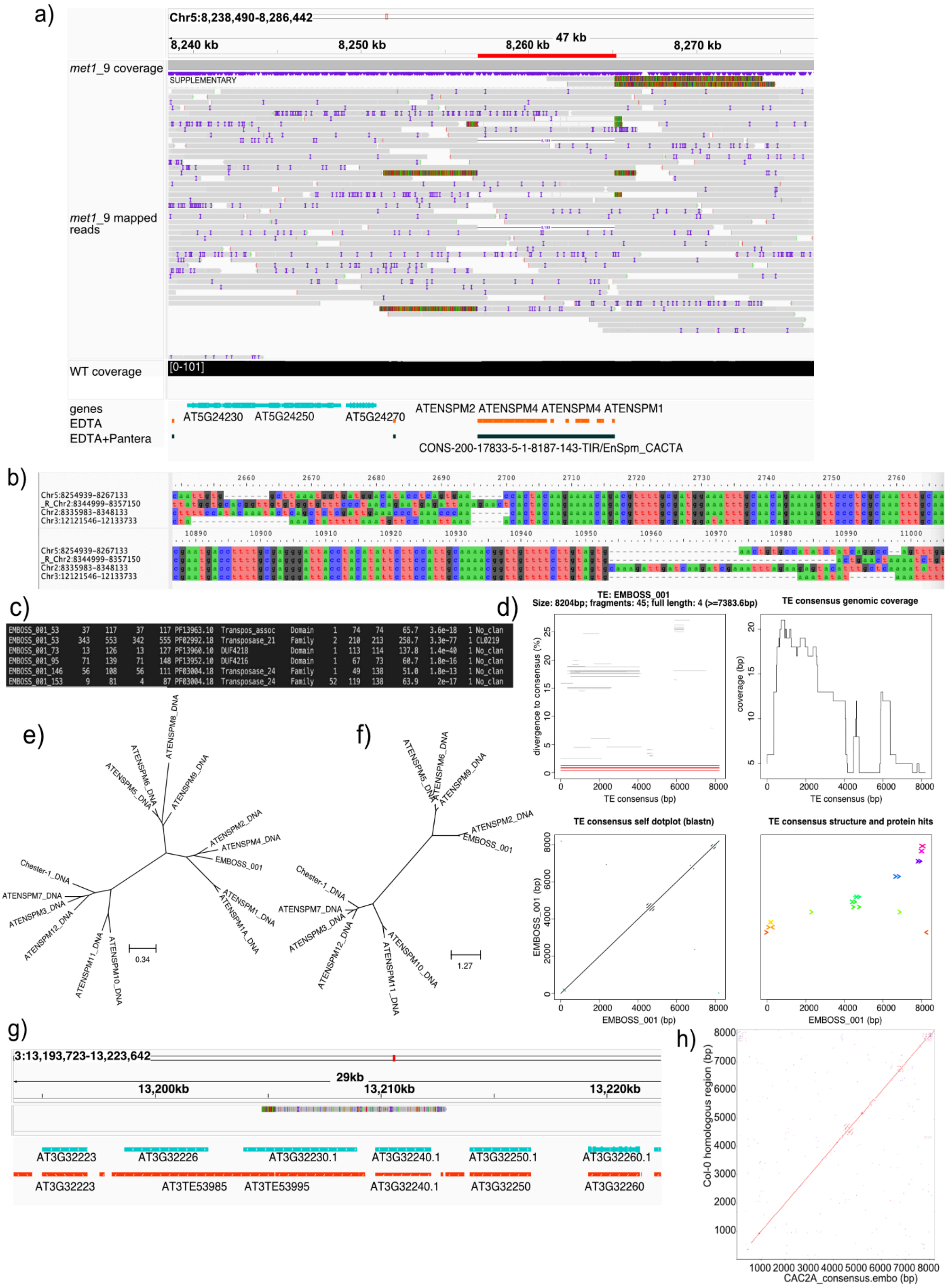


**Supplementary Figure 5**. **The CAC_2A TE family in the Tsu-0 genome. a** IGV screenshot of CAC2A_2. Mapped HiFi reads of the *met1_09* individual are shown in the first track. Alignments of high-quality reads (grey) showed both a somatic excision (read with solid black line connecting two aligned grey parts) and several somatic insertions (reads with multi-colored sections – when the reads mapped somewhere else and stop mapping they are colored by their nucleotide composition). The EDTA TE annotation was fragmented (red)

but the PANTERA annotation (black) showed a contiguous annotation for the element. **b** Multiple-sequence alignment of the borders of CAC2A copies, visualized in AliView. **c** Screenshot of Pfam domain annotation of CAC2A *consensus* (EMBOSS_001). **d** TE-Aid screenshot of CAC2A consensus. **e** Tree of *consensus* sequences of ATENSPMs from the Col-CC TE library (https://github.com/oushujun/TAIR12-TE) and CAC2A *consensus*. **f** Tree of translated Tnp domains of *consensus* sequences of ATENSPMs from Col-CC library and CAC2A *consensus* (EMBOSS_001). **g** IGV screenshot of CAC2A *consensus* aligned to TAIR10 Col-0 genome. Gene annotation (cyan), TE annotation (red). **h** Dotplot of CAC2A *consensus* against the region in the Col-0 genome where the CAC2A *consensus* mapped, as seen in Supplementary Fig. 5g, generated with re-DOT-able.

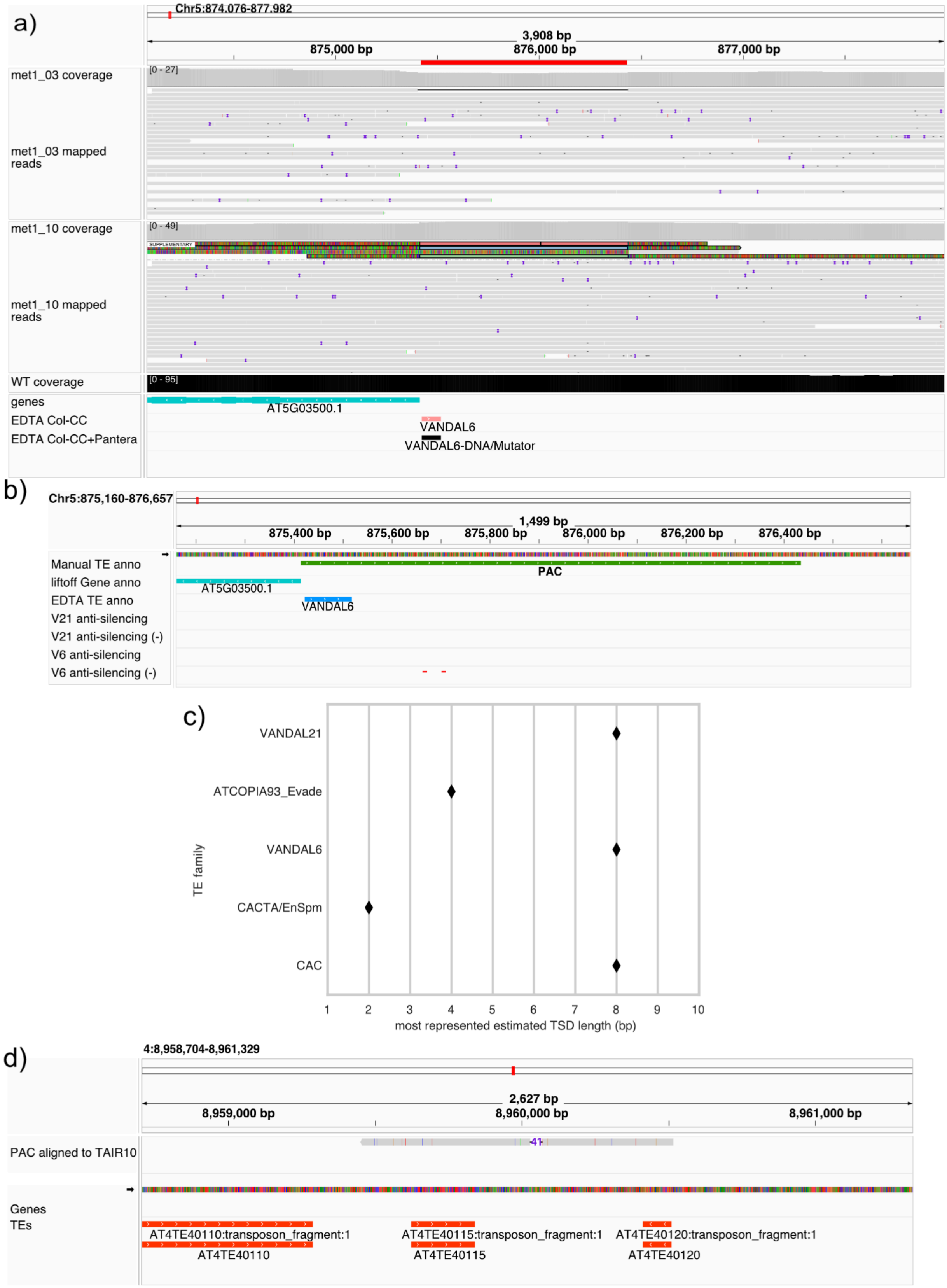


**Supplementary Figure 6**. **PAC non-autonomous element. a** IGV screenshot showing an apparent somatic insertion in the *met1_03* individual (read with solid black line connecting two aligned grey parts) and somatic insertions in the *met1_10* individual (reads with supplementary alignments stacked on top of the track, colored in red, blue and green). EDTA and Pantera annotated only a small section of this element (red and black) as VANDAL6. **b** IGV screenshot showing VANDAL6 anti-silencing motifs (Sasaki et al. 2022) (bottom tracks, red) in the PAC sequence. **c** Modes of estimated TSD lengths in different classifications of somatically mobile TEs. **d** IGV screenshot of PAC Tsu-0 sequence mapped to the TAIR10 Col-0 genome.

a)

| Name/Position TE | Predicted ORF; codon | Abbreviation |
|---|---|---|
| Chr1_11941106-11946436_279 | [4773 - 559] (REVERSE); 1405 | **EVD_1** |
| Chr3_8698969-8699375 | 406bp FRAGMENT; -- | **Chr3:86** |
| Chr3_11698155-11703548_102 | [1098 - 4796]; 1233 | **Chr3:11** |
| Chr3_19688293-19693627_104 | [3038 - 4777]; 580 | **Chr3:19** |
| Chr4_3014412-3019742_279 | [4773 -559] (REVERSE); 1405 | **EVD_3** |
| Chr5_21419693-21425022_279 | [4772 - 558] (REVERSE); 1405 | **EVD_2** |
| ATR_TAIR10_104 | [558 - 4772]; 1405 | **ATR** |
| EVD_TAIR10_105 | [558 - 4772]; 1405 | **EVD** |

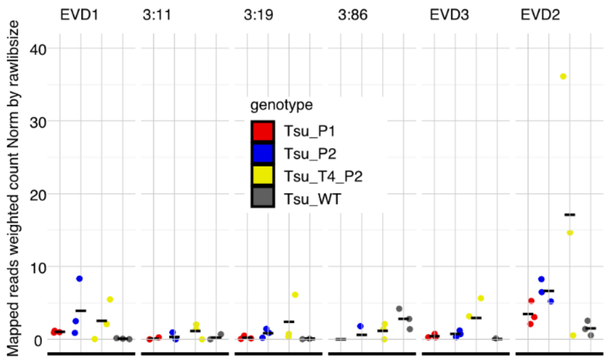


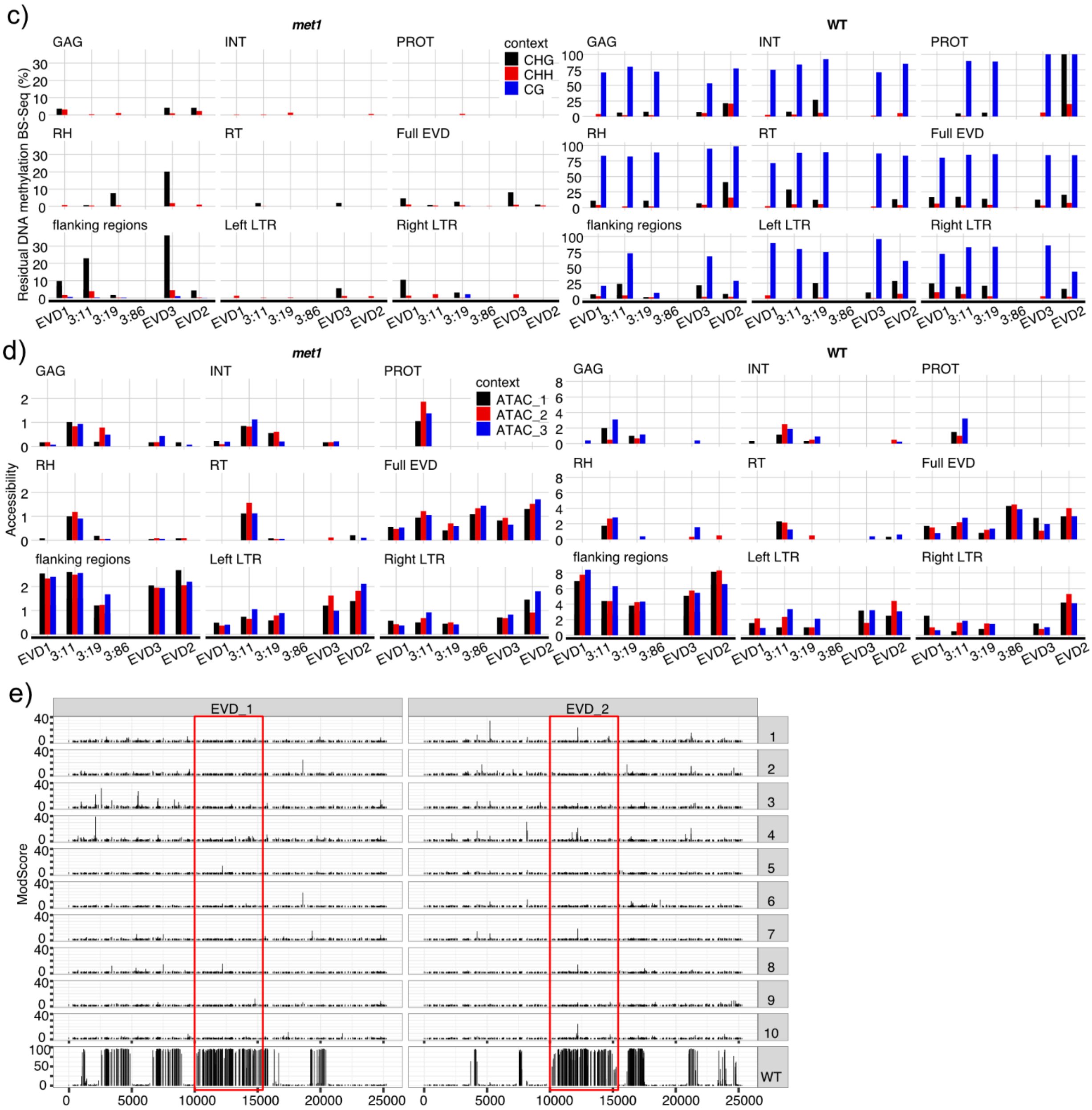


**Supplementary Figure 7**. **Properties of EVD elements in Tsu-0. a** Length (bp) of predicted ORFs of Tsu-0 EVDs (EVD1-3, Chr3:86, Chr3:11, Chr3:19) compared to the intact ÉVADÉ subfamily elements EVD and ATR in Col-0. **b** Expression of Tsu-0 EVDs in *met1* line 1 (Tsu_P1, red), *met1* line 2 (Tsu_P2, blue), second-generation *met1* line 2 (Tsu_T4_P2, yellow), and wild type (Tsu_WT, grey). **c** Methylation over Tsu-0 EVDs as determined by

BS-Seq data in *met1* (left) and wild-type (right). **d** ATAC-Seq accessibility of Tsu-0 EVDs in *met1* (left) and wild-type (right). **e** CG methylation probability scores from CCS reads over EVDs mobile in Tsu-0 (EVD_1,2, red line) and flanking sequences in *met1* individuals (1-10) and the wild-type pool.

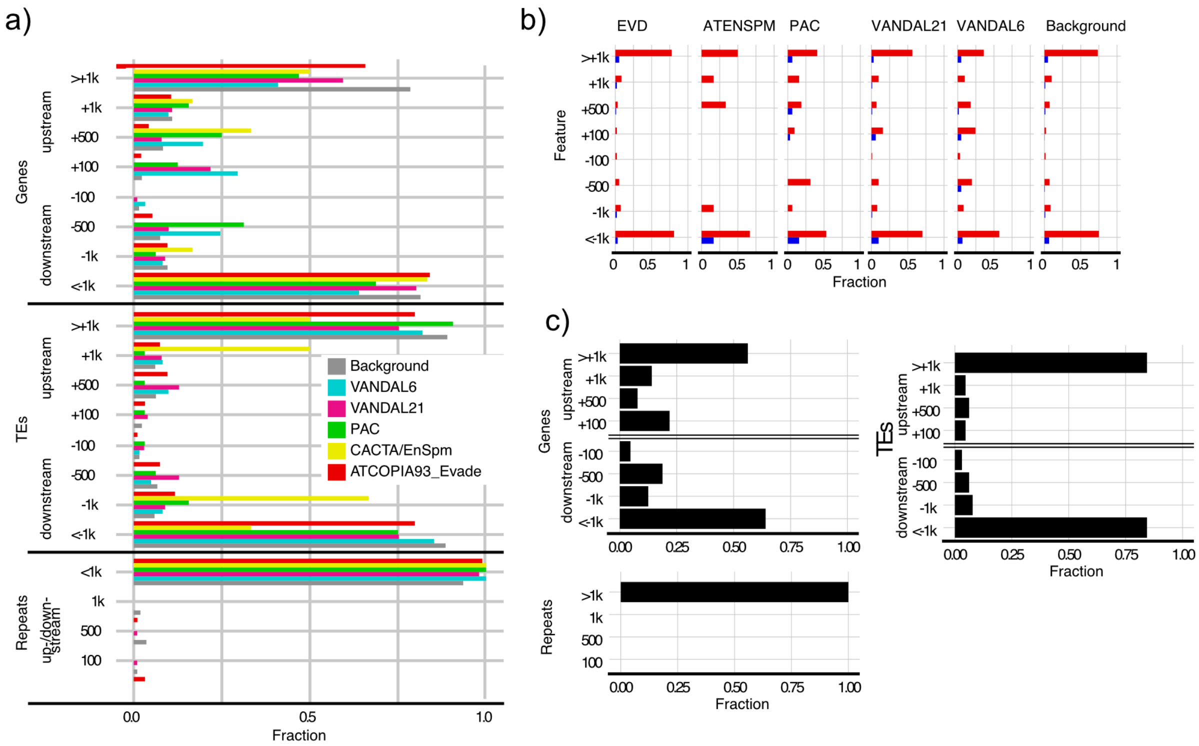


**Supplementary Figure 8**. **Features close to somatic insertion sites and breakends of VANDAL21 hypermutable region. a** Fractions of somatic insertion sites and randomly generated positions (Background), plotted by distance from feature (bp) and colored by category. For genes and TEs, insertions are split as inserted upstream or downstream of the respective feature. **b** Fractions of genic somatic insertion sites plotted by distance from feature (bp) and colored by "essentiality" of the disrupted gene (non-essential, red; essential, blue). **c** Fractions of breakend coordinates of VANDAL21 hypermutable region, plotted by distance from feature (bp). For genes and TEs, breakend coordinates are split as inserted upstream or downstream of the respective feature.

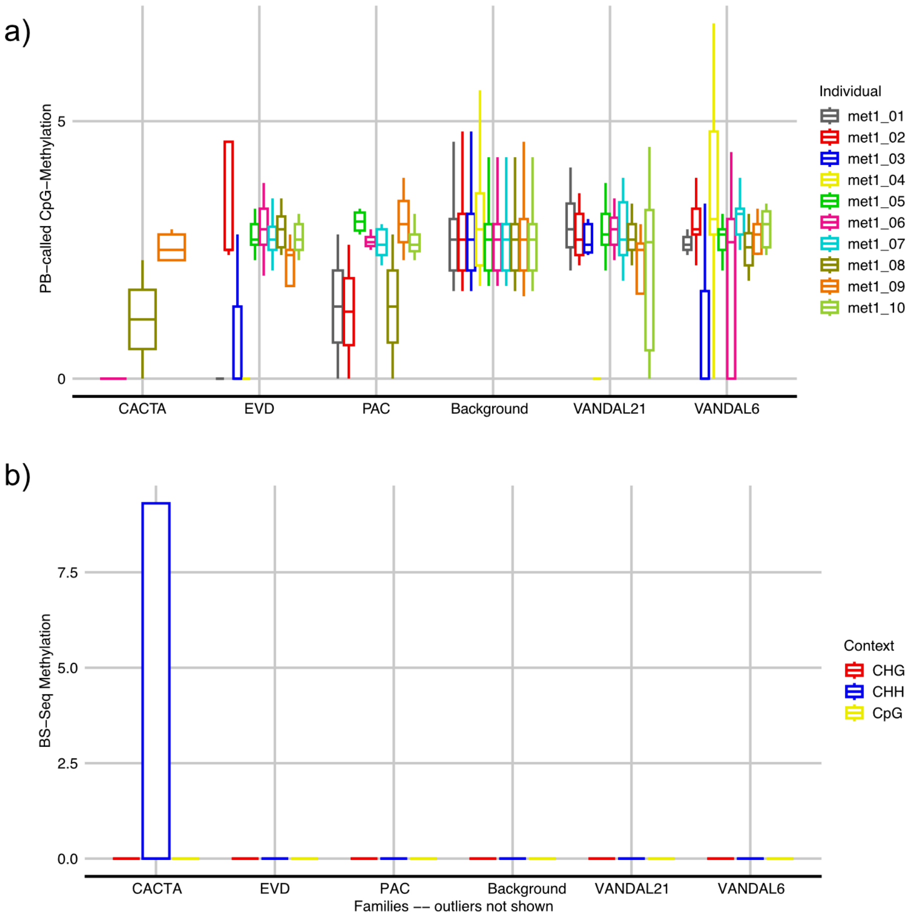


**Supplementary Figure 9**. **Residual methylation at somatic insertion sites. a** Distribution of residual CG methylation per TE category, called from CCS data and colored by *met1* individual. **b** Distribution of residual CG, CHH, CHG methylation per TE category, called from BS-Seq data and colored by context.

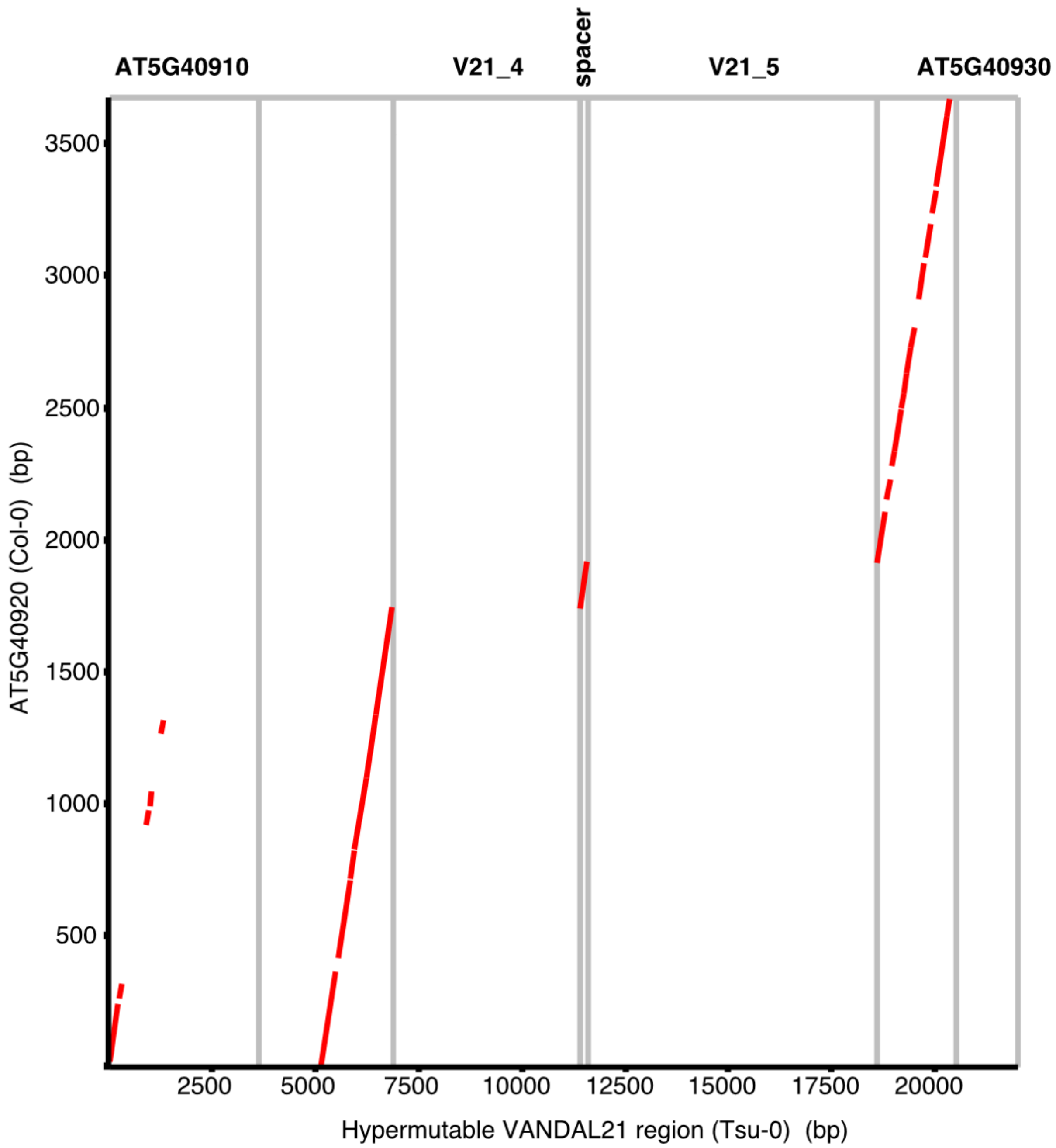


**Supplementary Figure 10. VANDAL21 elements from the hypermutable region disrupting a TIR-NLR derived sequence.** Dot plot indicating sequence similarity between TAIR10 AT5G40920 and Tsu-0 hypermutable VANDAL21 region, comprising V21_4, spacer, V21_5, and upstream and downstream flanking sequences. Visualized with re-DOT-able.

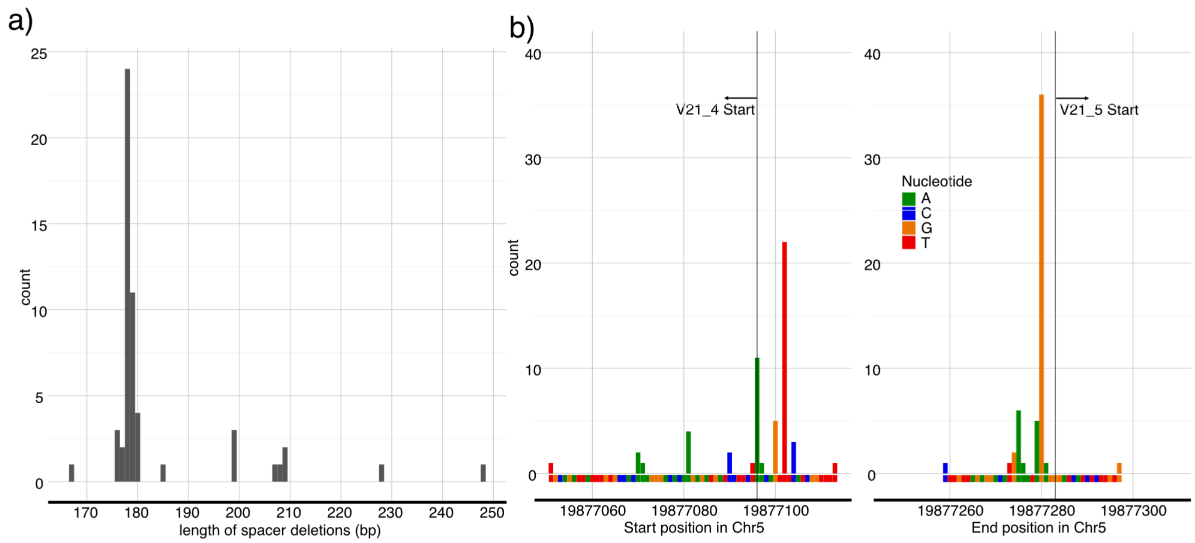


**Supplementary Figure 11**. **Spacer DNA deletions. a** Number of spacer deletions by length. **b** Start (left) and end (right) positions of spacer deletions. The start and direction of VANDAL21s is indicated with arrows.

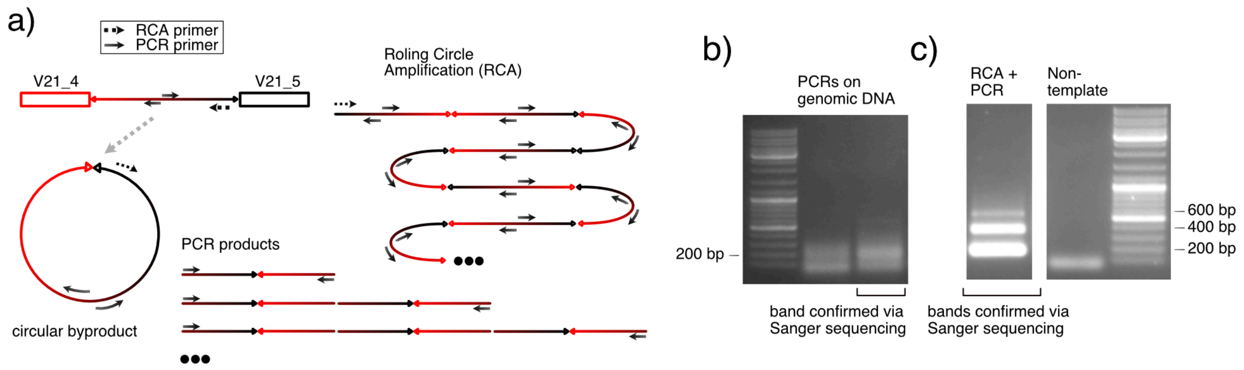


**Supplementary Figure 12**. **Detection of circular forms of the spacer DNA via RCA and PCR. a** Schematic representation of the circular DNA formed as a byproduct of alternative transposition of VANDAL21 TEs (V21_4 and V21_5). The circular spacer is amplified with one primer (dotted arrow) via rolling circle amplification (RCA) and used as input DNA for a PCR with two primers (solid arrows). Different PCR products of incrementally-doubled sizes are expected **b** DNA ladder and two technical replicates of PCR product from genomic DNA input in a 1% agarose gel. **c** PCR products from RCA input, non-template control and DNA ladder in a 1% agarose gel.

| Name | Sequence |
| --- | --- |
| spacer_DNA_F | AACGAGTCTTTGAAGGGATGC |
| spacer_DNA_R | TAGAGAACTCCCCGATCTTGG |
| spacer_RCA | GTTTTGCAATATTTGG |

**Supplementary Table 2. Primers table**
