## Additional File 4 for "Long-read detection of transposable element mobilization in the soma of hypomethylated *Arabidopsis thaliana* individuals": CIGAR_Excisions.pdf

#### Only deletions that match 99% with a TE

### Merged elements

```
"SatCEN" = satellite or centromeric rearrangements - not to be trusted
```

#### met1_01

> Chr2 6734994 6775798 ATMSAT1 met1_01

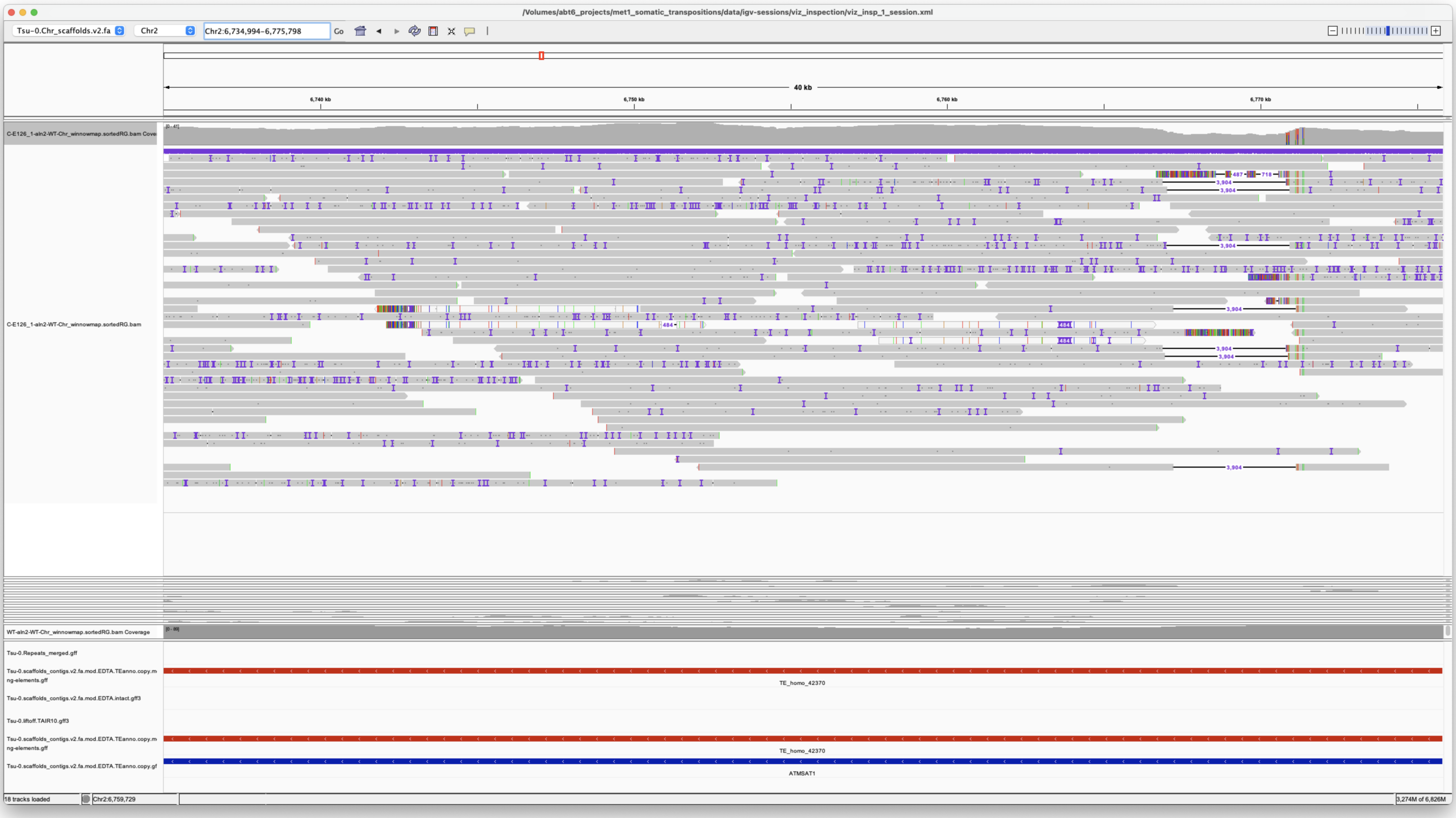


*SatCEN*

> Chr2 8591002 8595324 ATENSPM1 met1_01

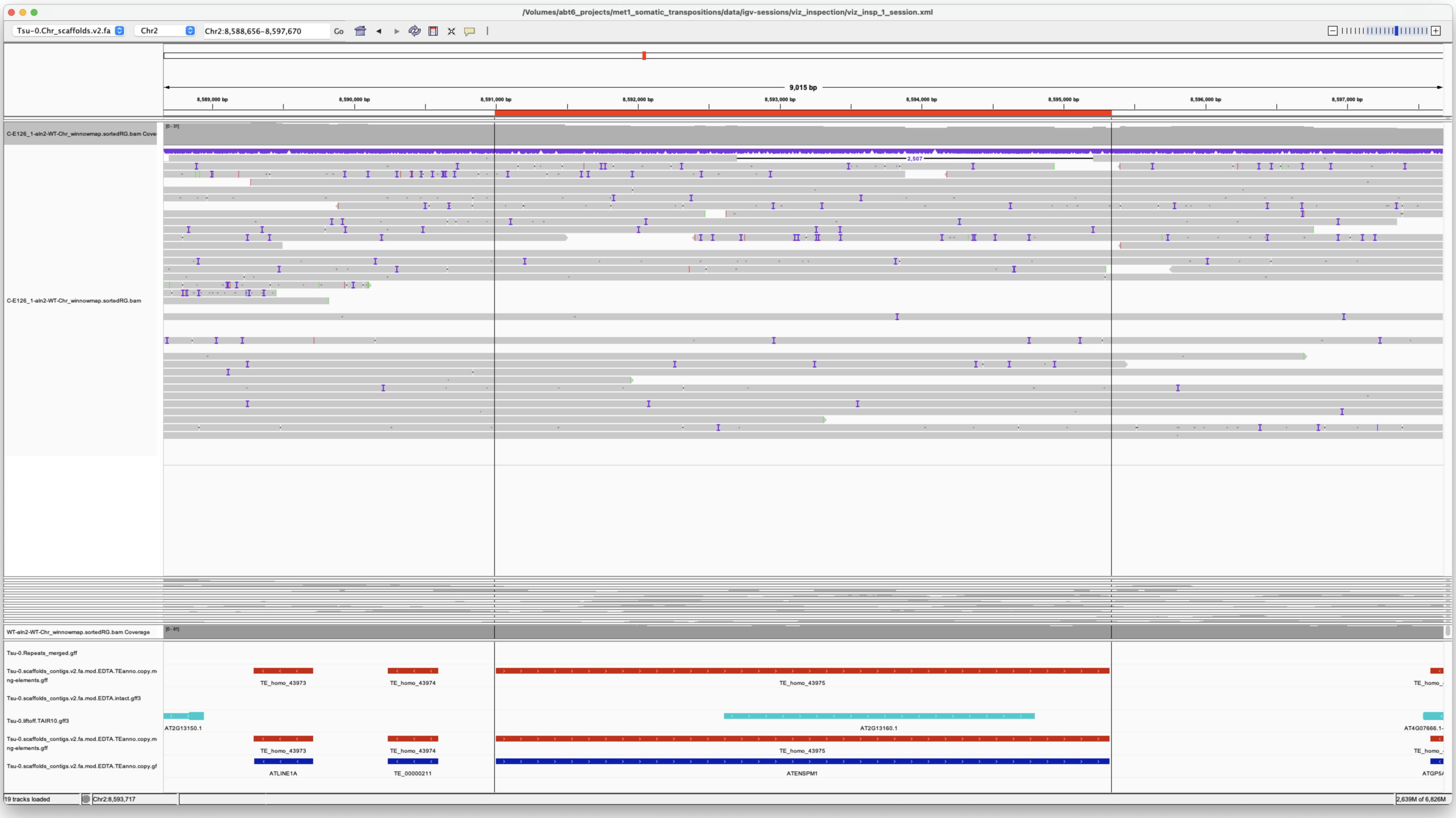


internal deletion not associated to insertions

```
There is also the possibility this is misannotation
```

*Rearrangement*

> Chr5 19152829 19160826 VANDAL21 met1_01

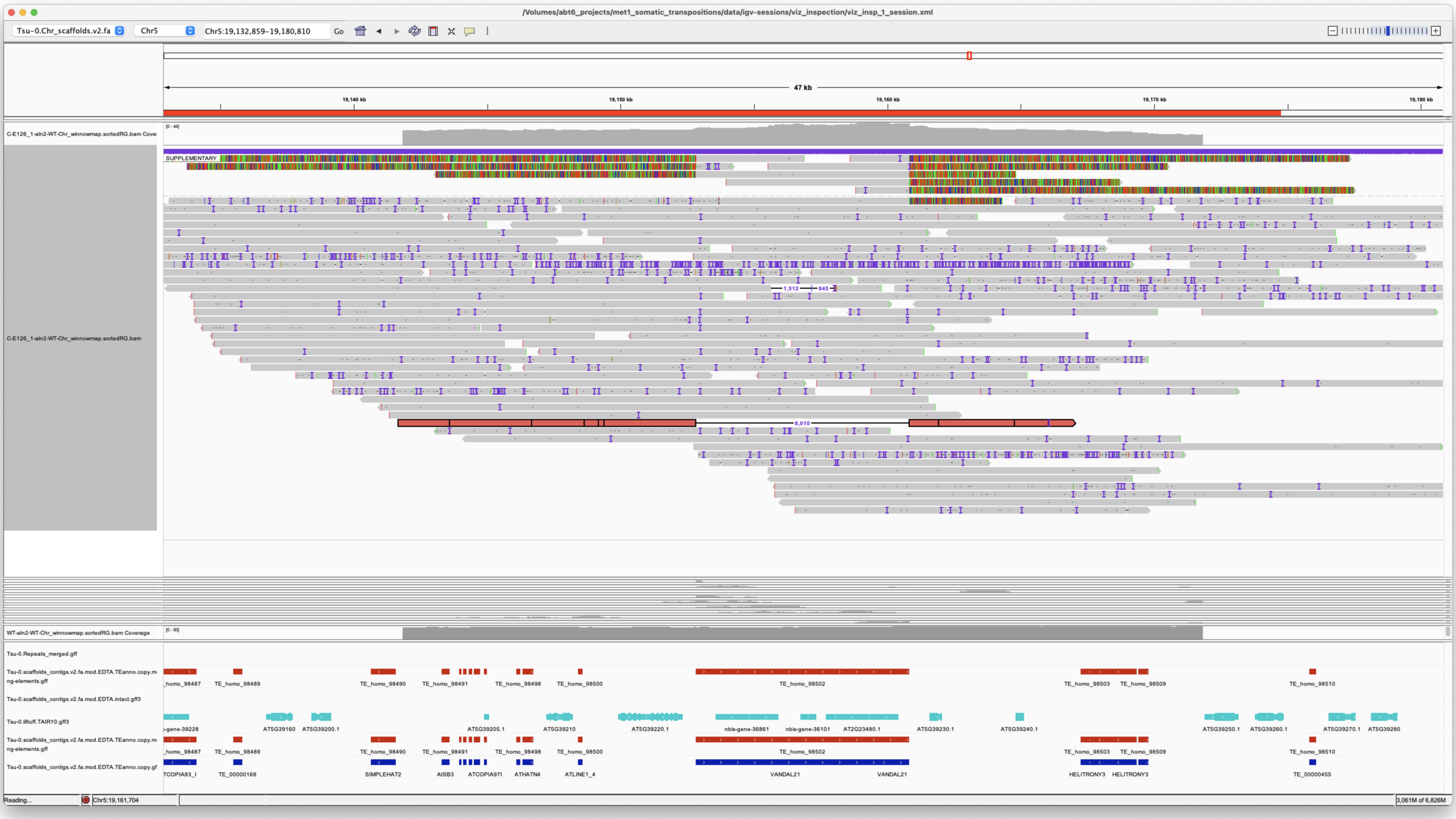


Associated with insertion

*Confirmed*

### met1_02

### met1_03

Chr5 875414 876434 PutativePackTypeCACTAMuDR met1_03

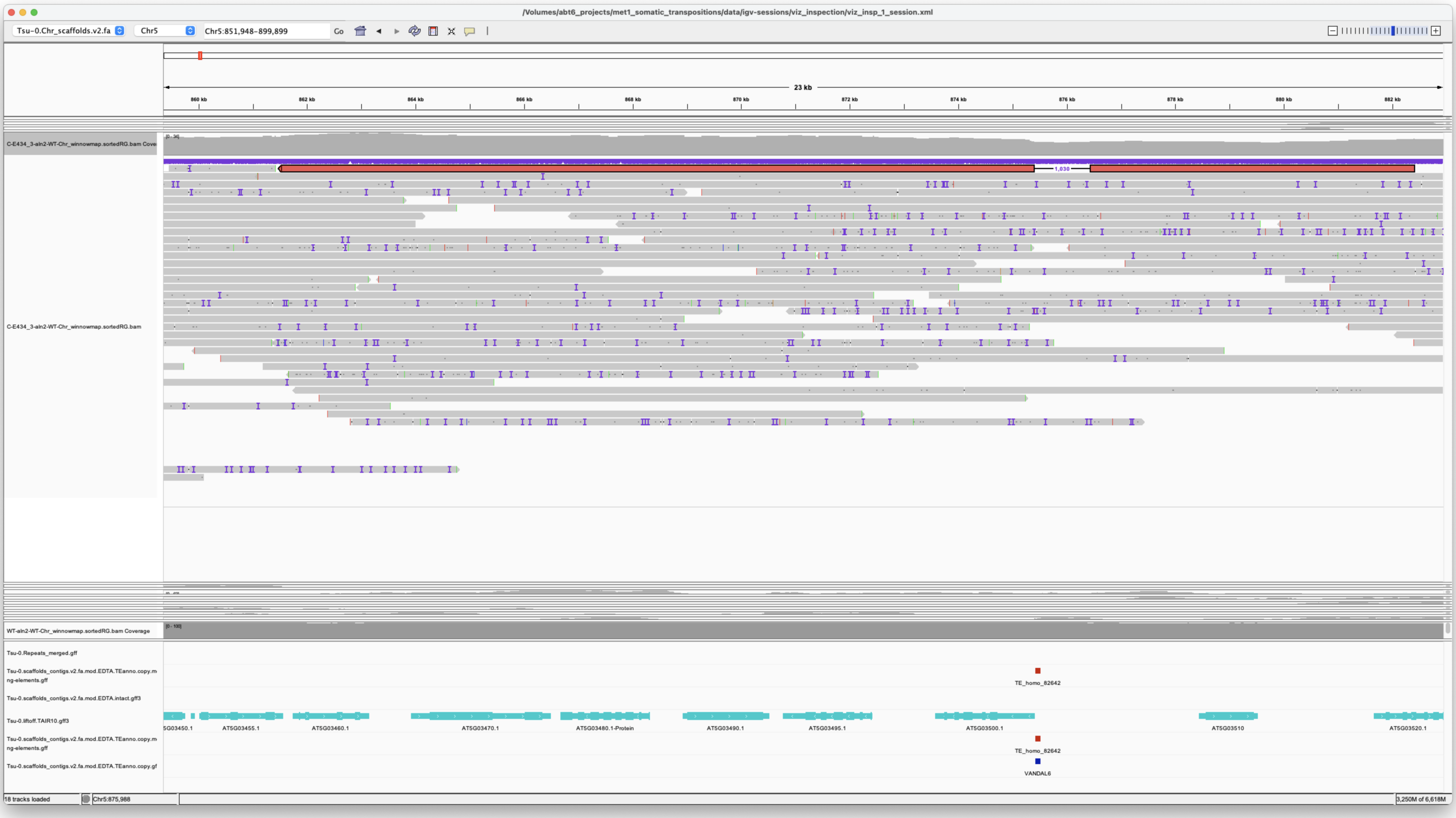


Annotation of putative Pack-Type missing from bottom tracks

*Confirmed*

### met1_04

Chr2 6734994 6775798 ATMSAT1 met1_04

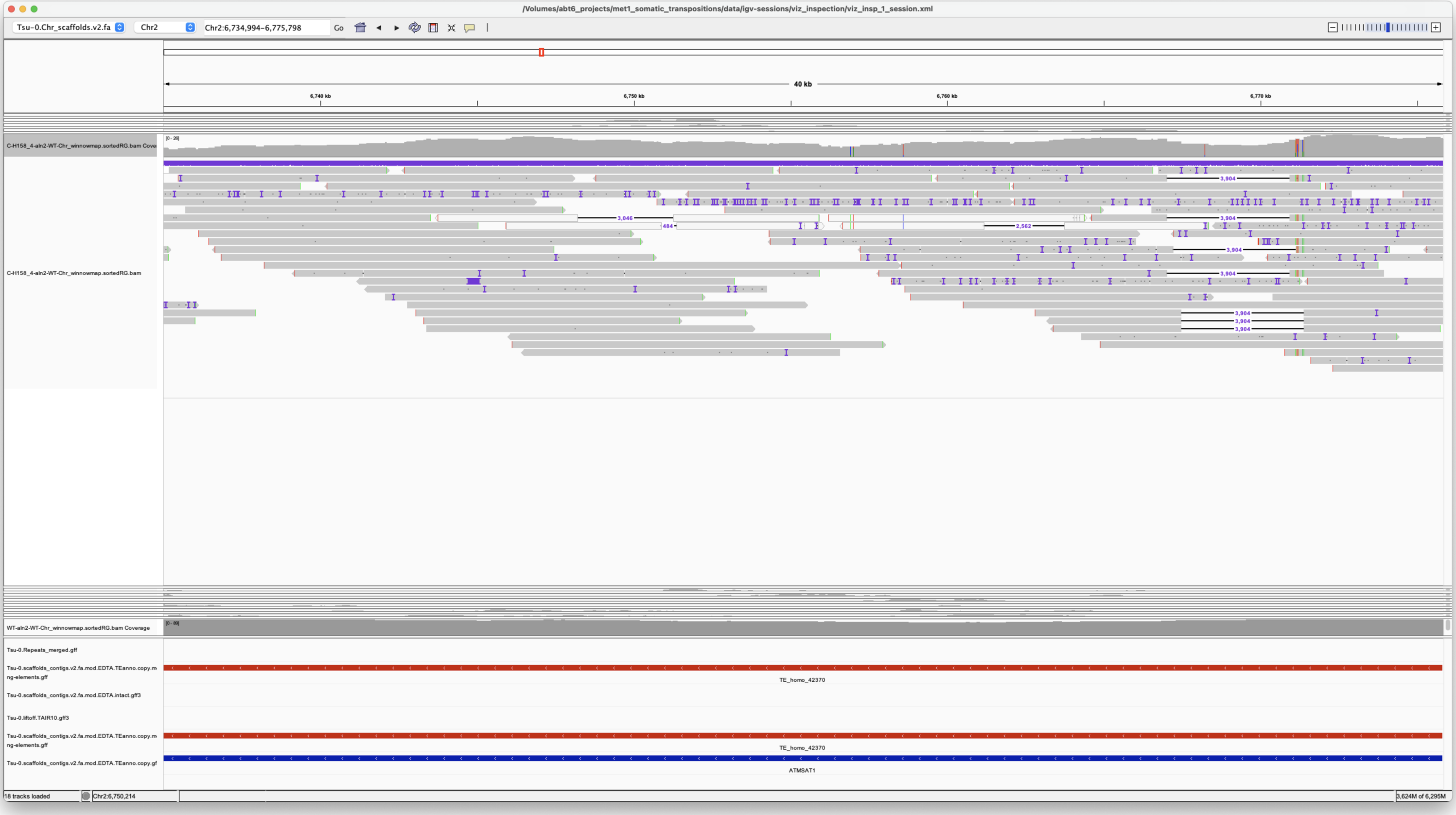


*SatCEN*

Chr5 19152829 19160826 VANDAL21 met1_04

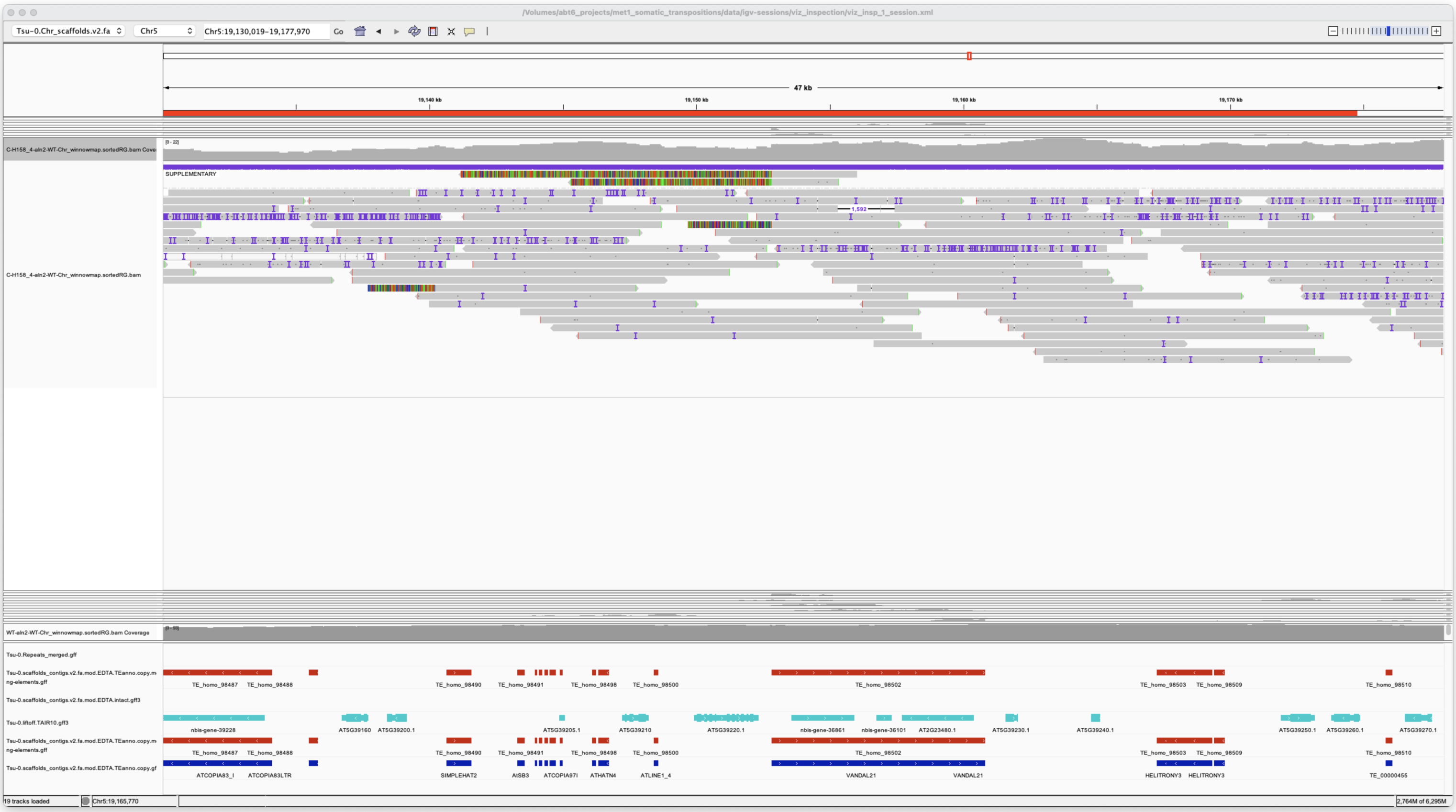


Internal deletion

### met1_05

Chr2 6734994 6775798 ATMSAT1 met1_05

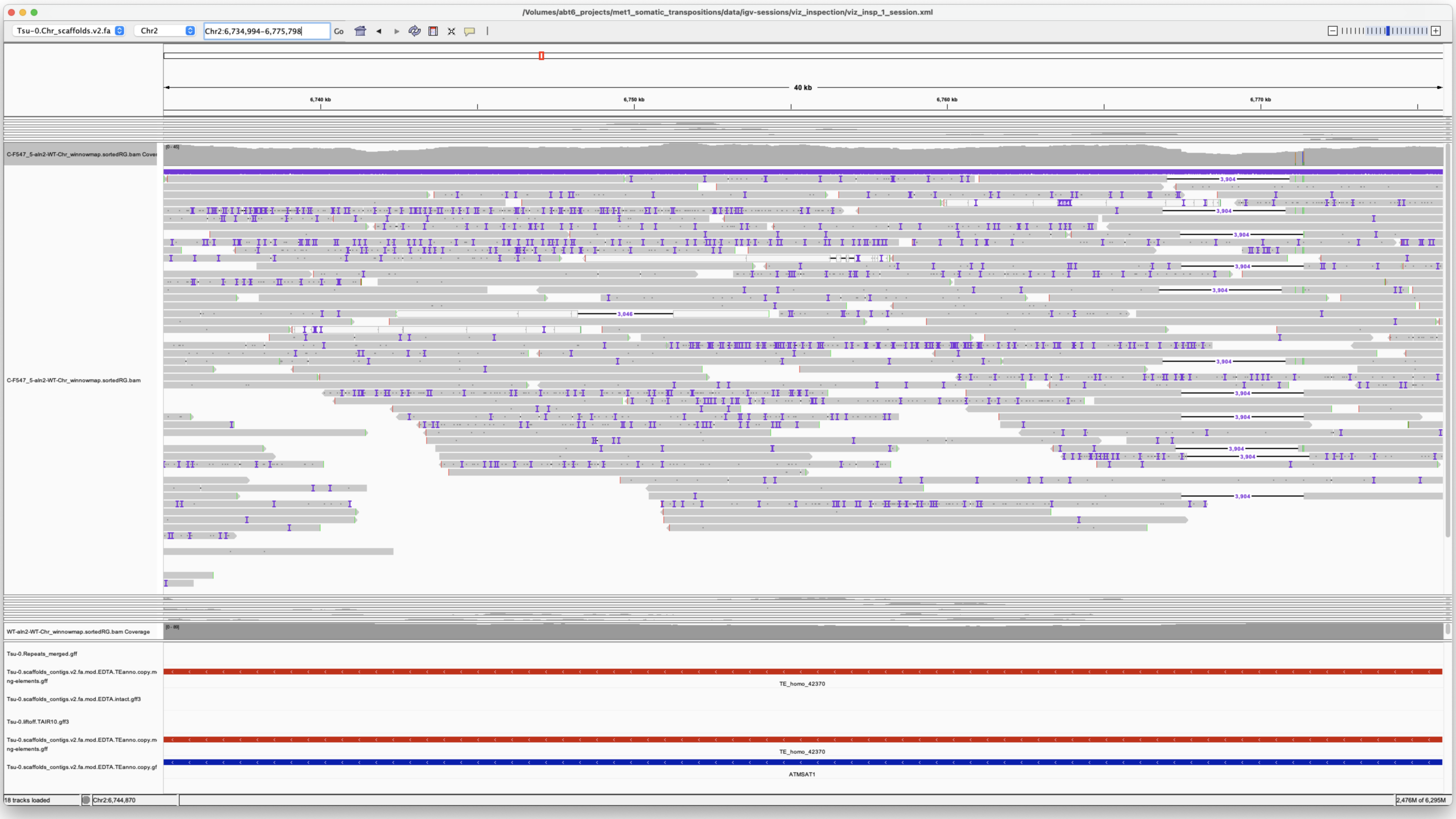


*SatCEN*

Chr5 19152829 19160826 VANDAL21 met1_05

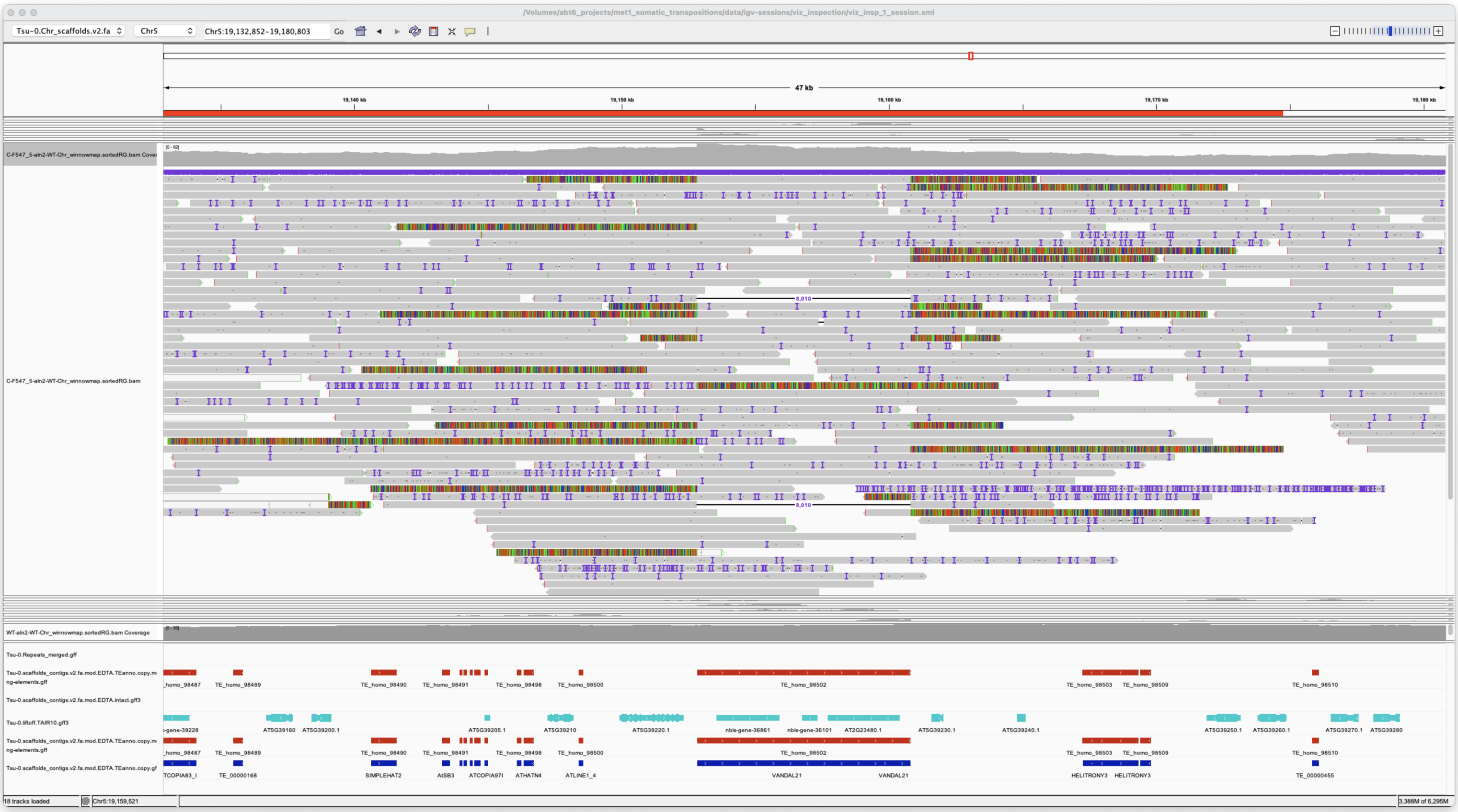


2x

*Confirmed*

---

### met1_06

Chr2 6734994 6775798 ATMSAT1 met1_06

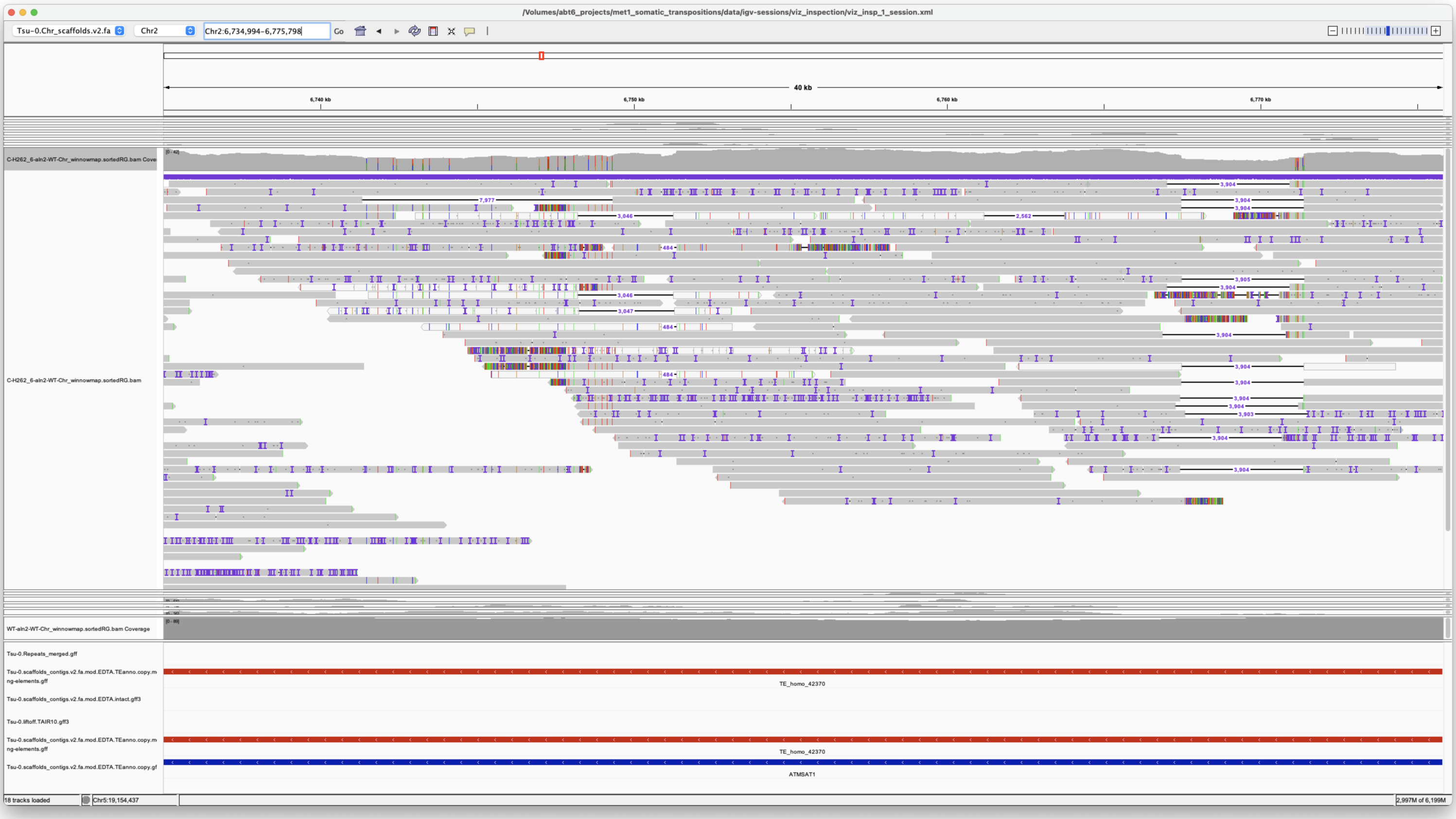

*SatCEN*

Chr3 16344522 16352497 VANDAL6 met1_06

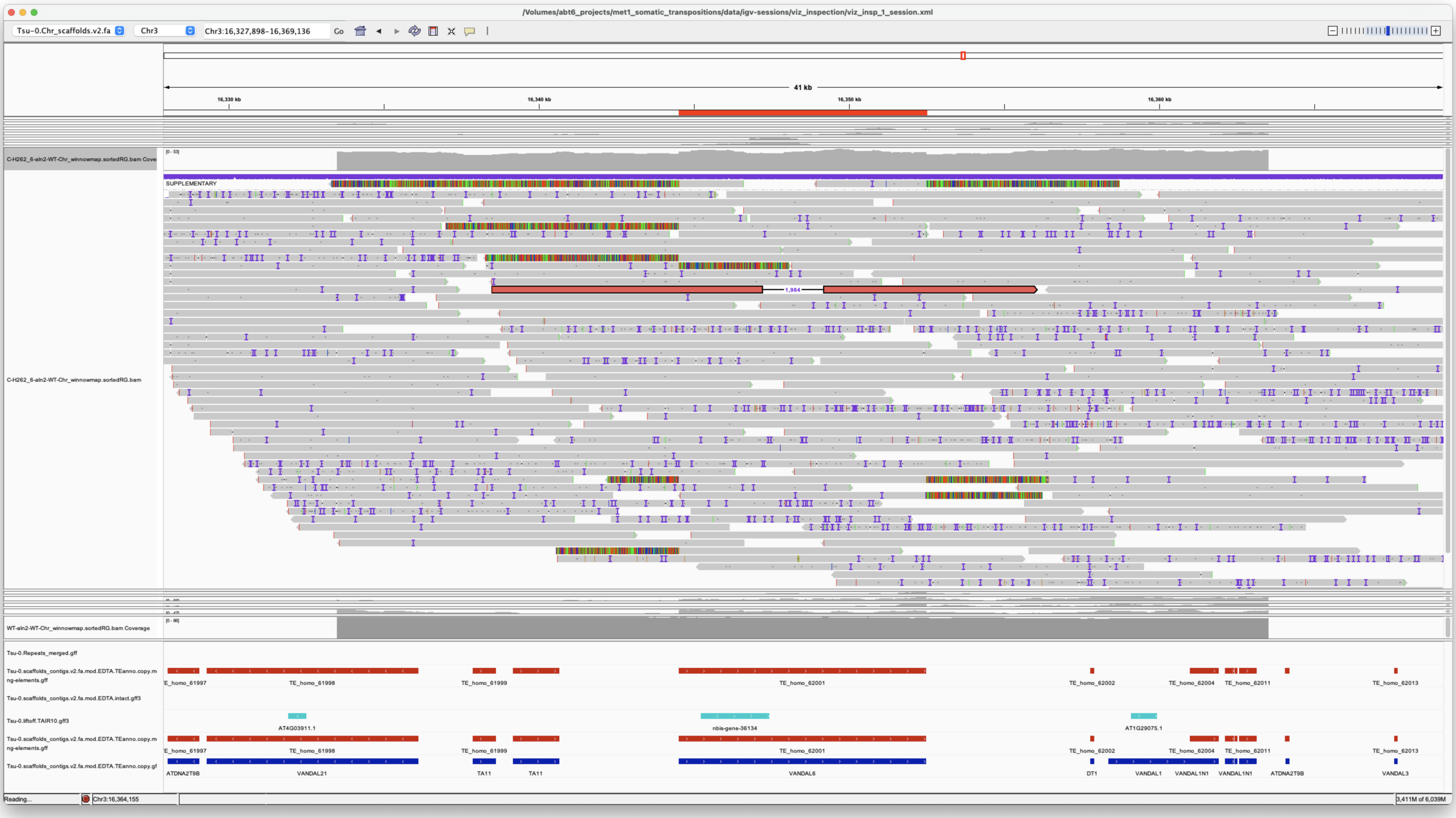

internal deletion

*Rearrangement*

Chr5 19152829 19160826 VANDAL21 met1_06

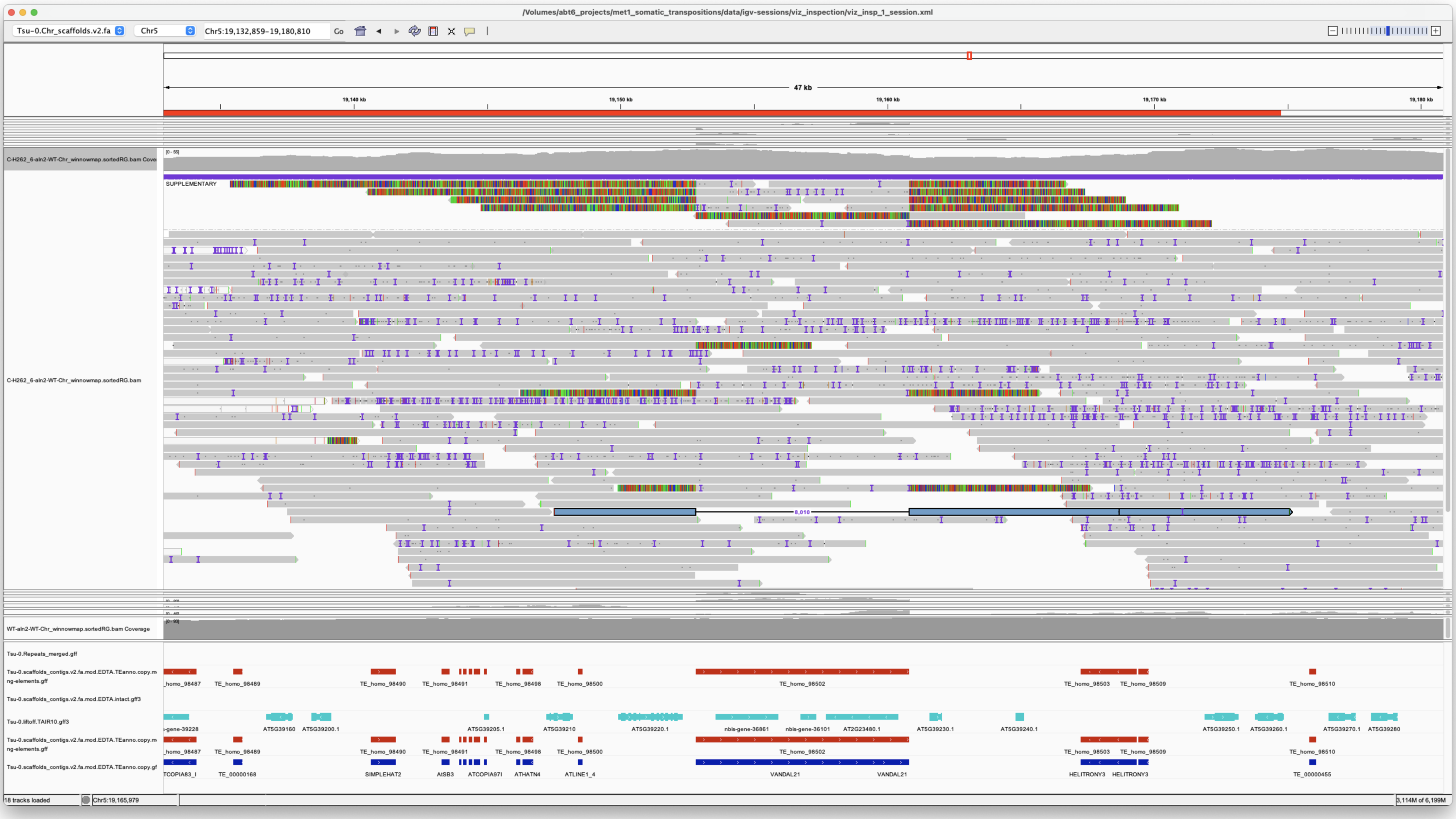


*Confirmed*

### met1_07

> Chr5 19152829 19160826 VANDAL21 met1_07

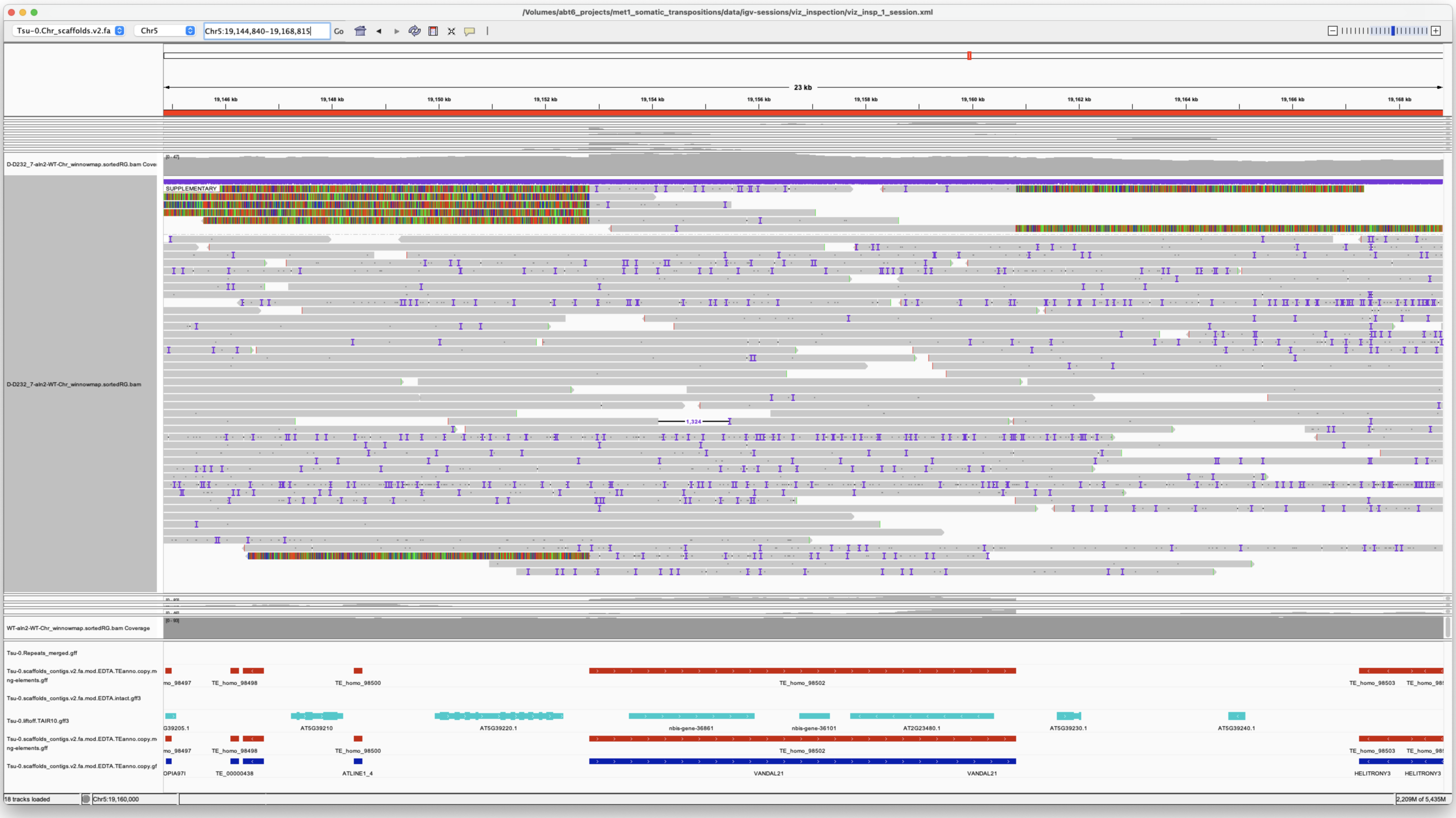


Internal deletion

*Rearrangement*

> Chr5 19872565 19877095 VANDAL21 met1_07

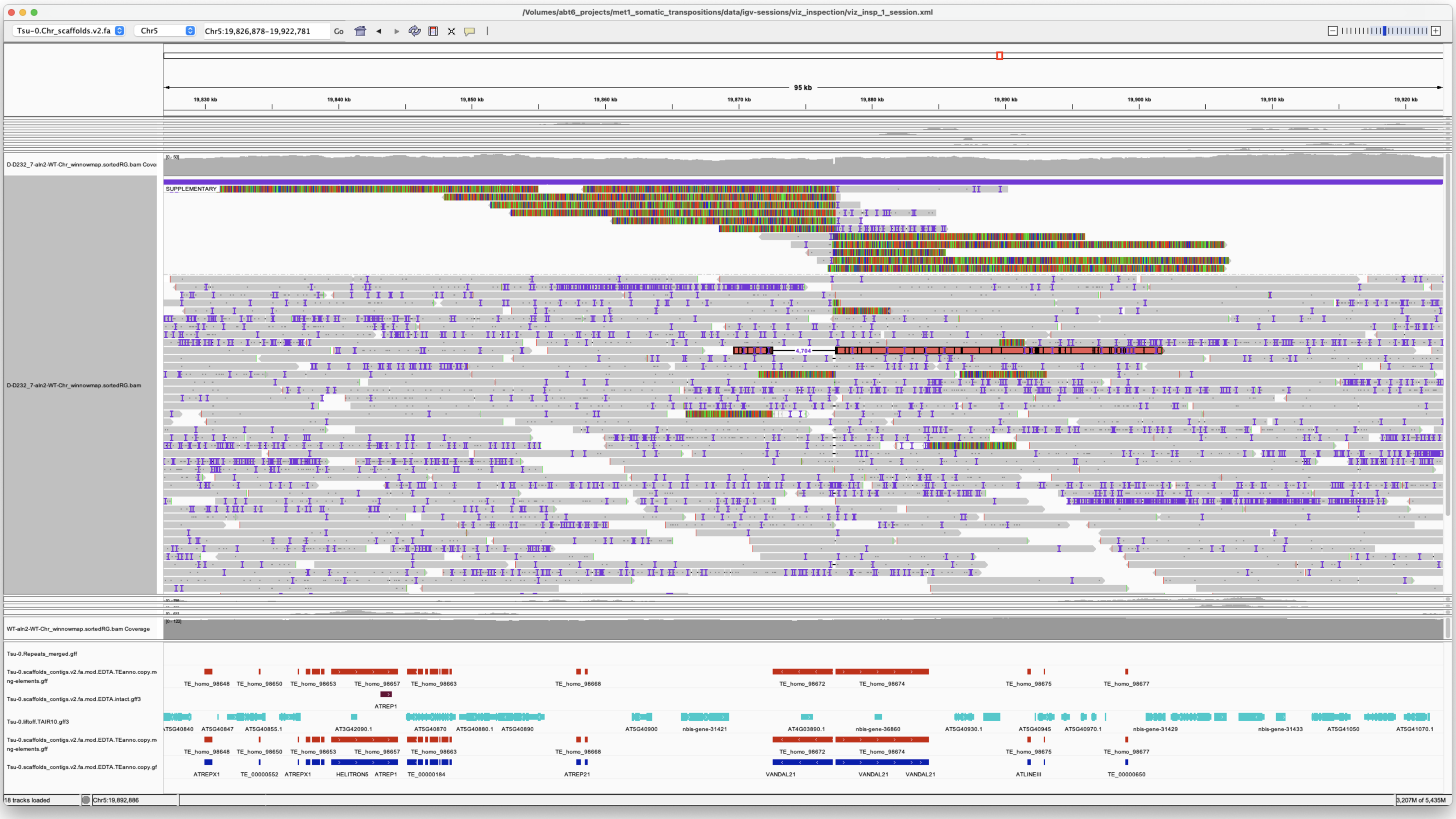

Hypermutable region, probable alternative transposition involved

Excluded

---

---

### met1_08

> Chr2 6734994 6775798 ATMSAT1 met1_08

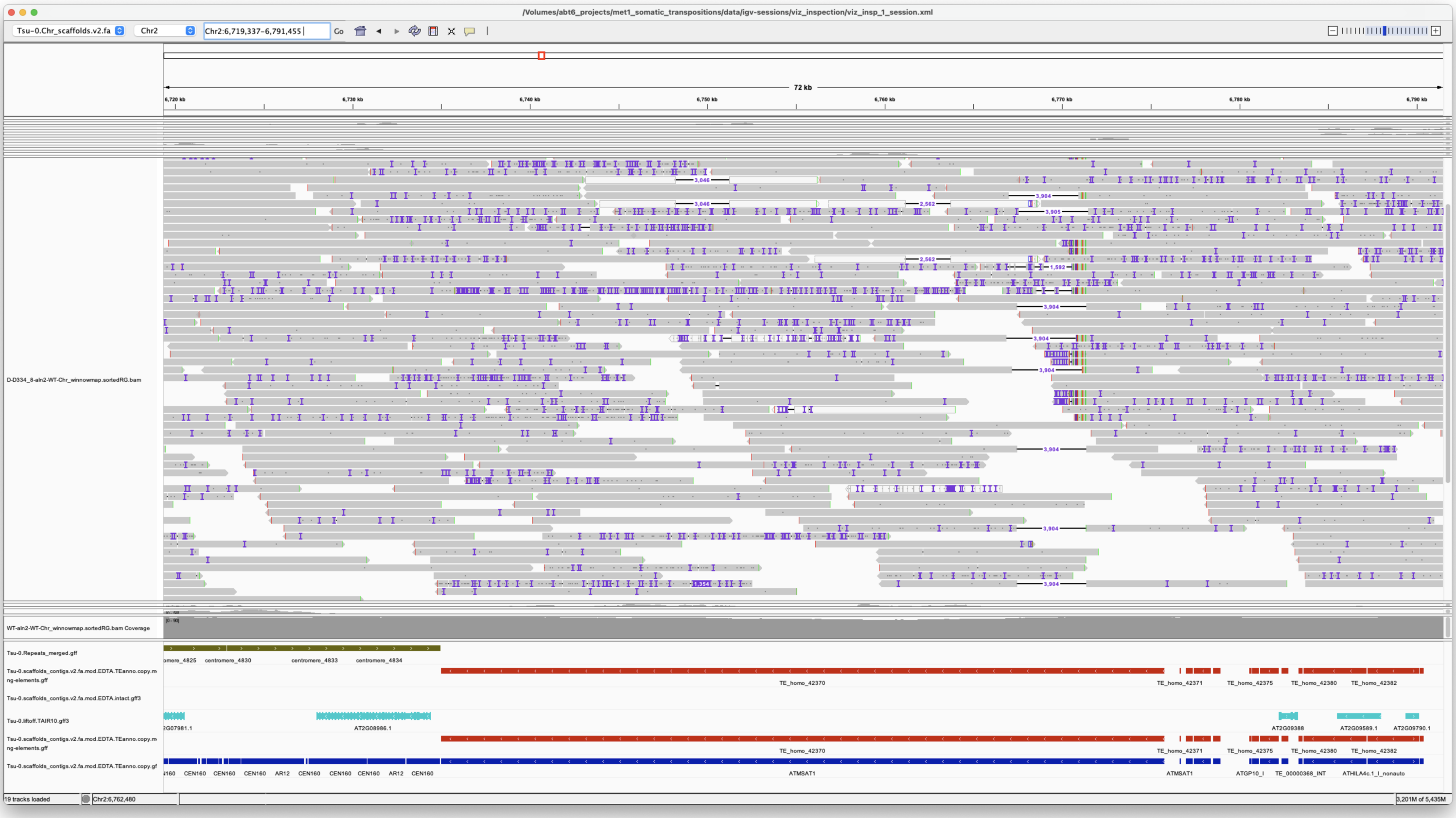

*SatCEN*

> Chr5 19152829 19160826 VANDAL21 met1_08

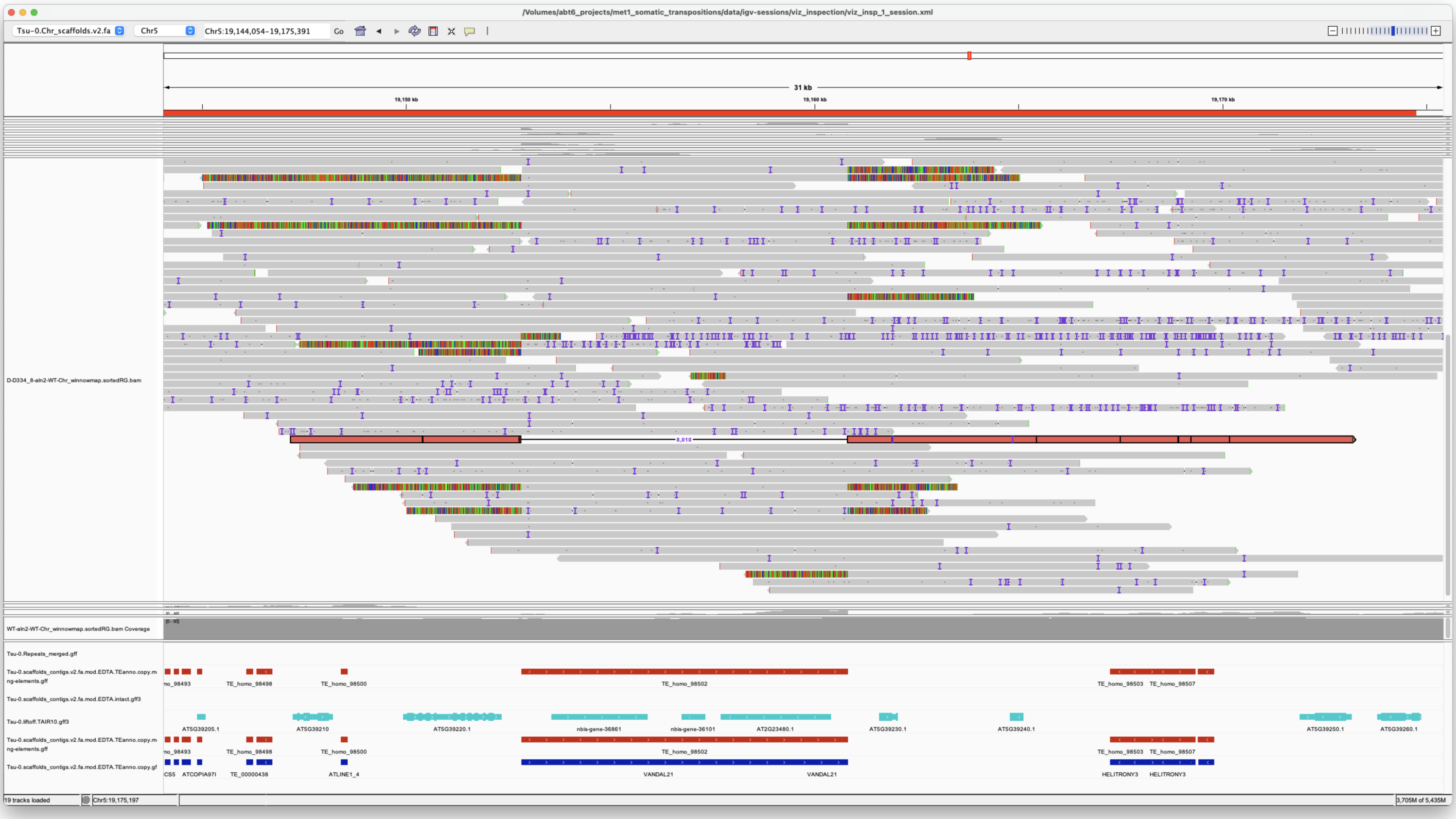


*Confirmed*

### met1_09

> Chr2 6734994 6775798 ATMSAT1 met1_09

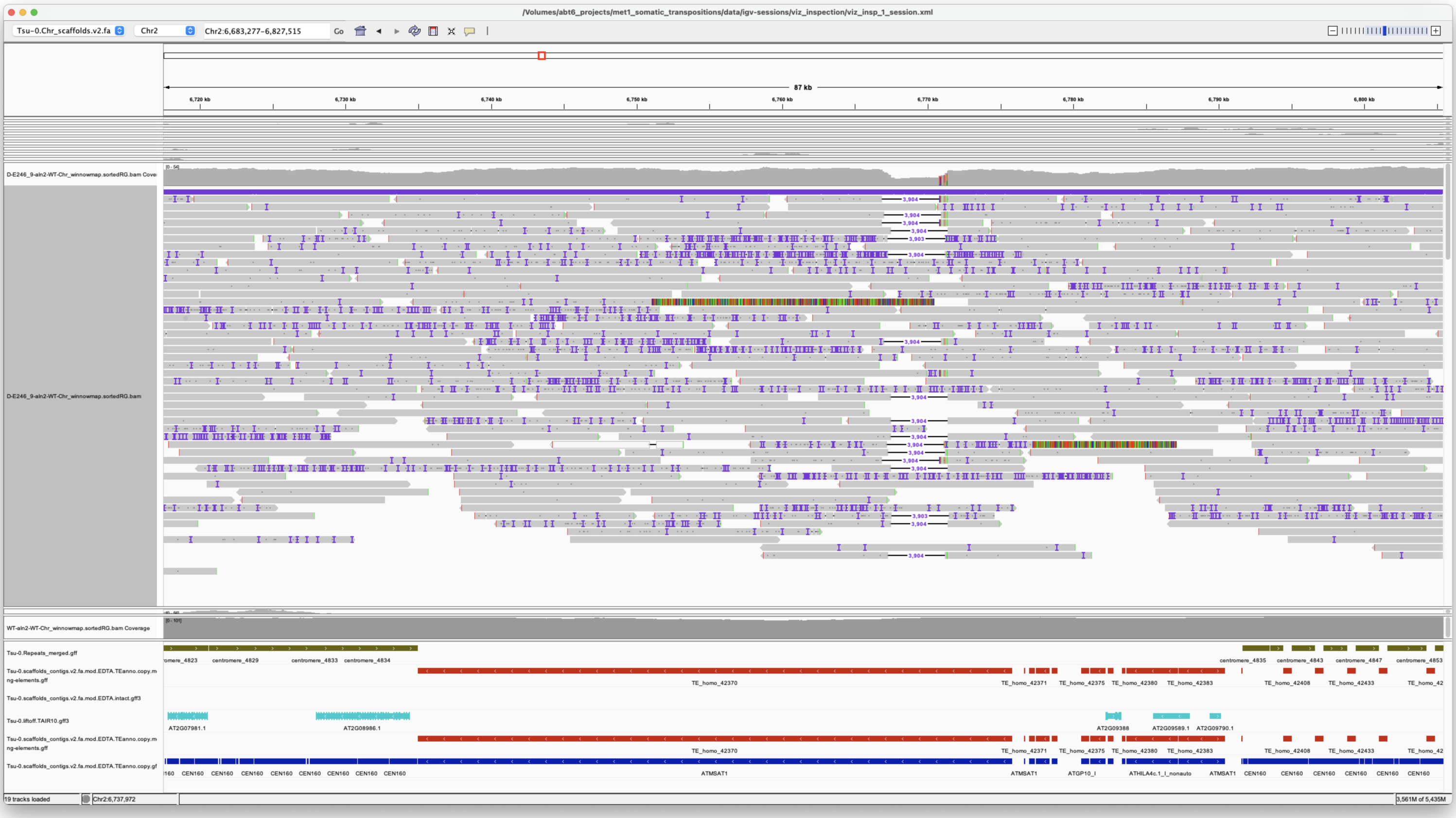


*SatCEN*

> Chr3 19893498 19897077 ATLINE1_12 met1_09

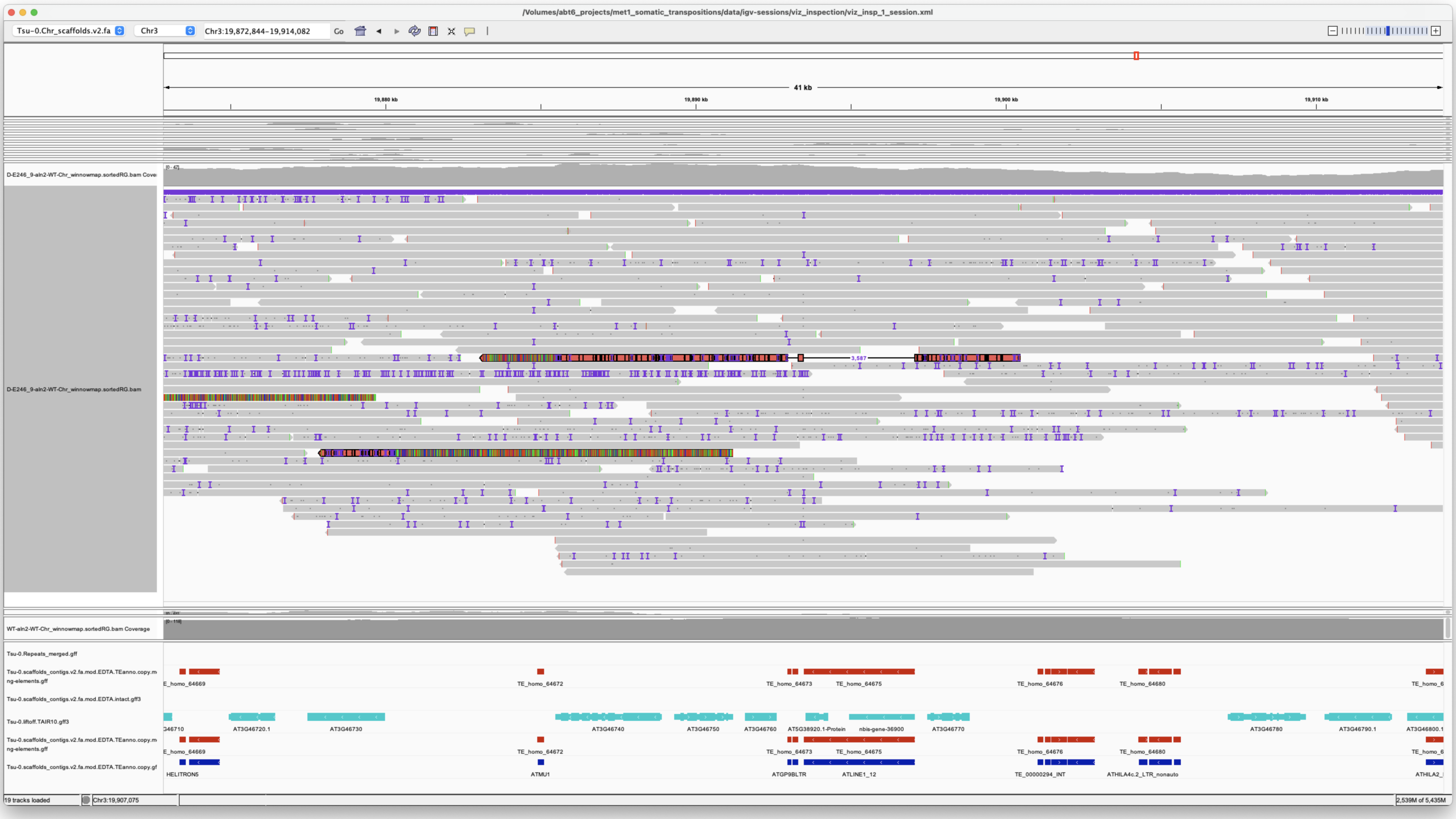


Weird, interesting Rearrangement

but confirmed that the deletion is overlapping perfectly with the annotated TE, which is a LINE and shouldnt be excided but..
probably something weird going on here OR ARTIFACT!!!
There is also a supplementary alignment on the left, THEREFORE:

*Not Confirmed*

> Chr5 4335532 4336888 ATDNAI27T9A met1_09

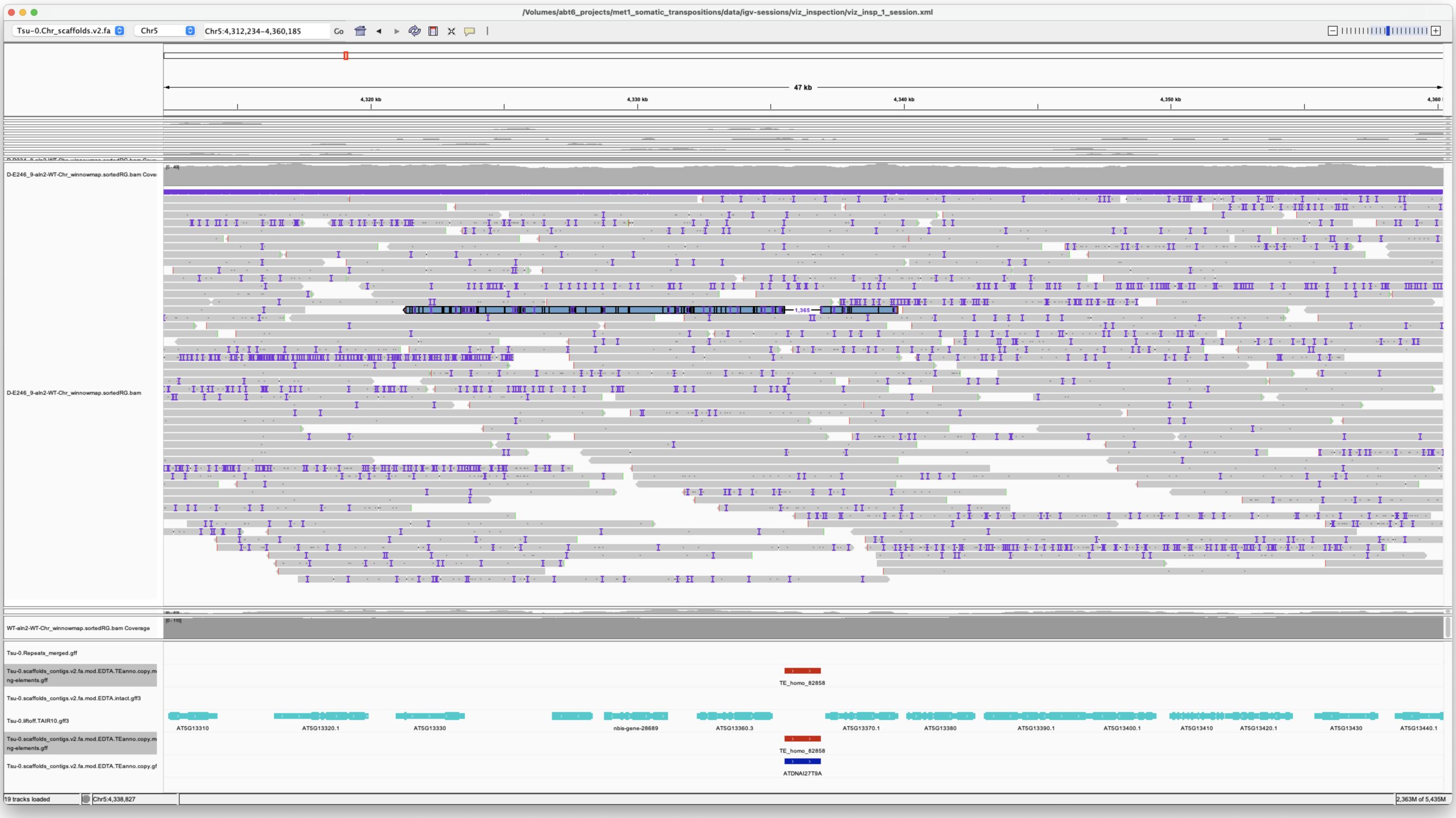


Seems real because it coinciding with the underlying annotation BUT there are no evidences of this element to insert in any met1 individual plant

> Chr5 8256939 8265133 ChimericCACTA met1_09

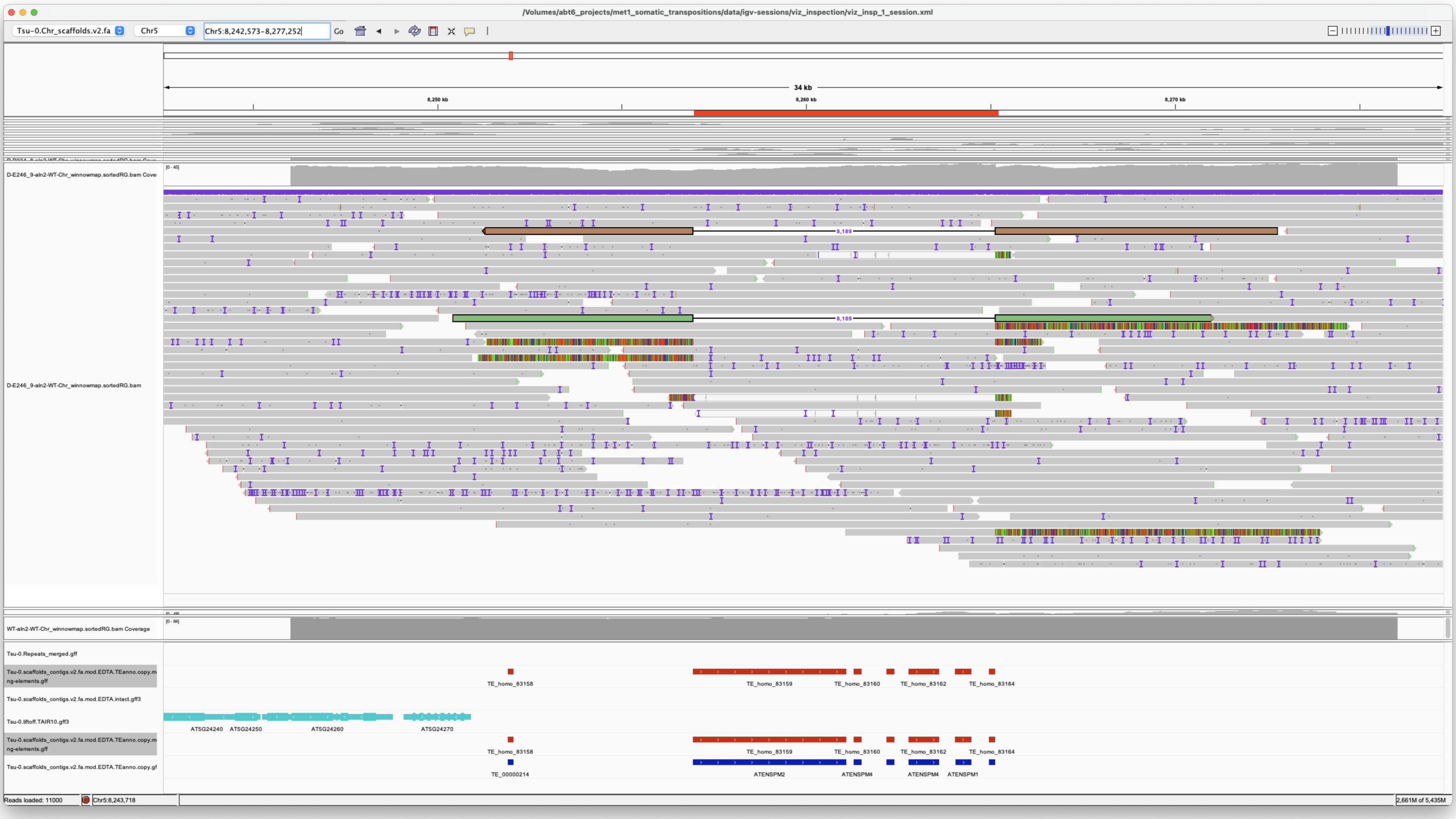

2x coverage

*Confirmed*

### met1_10

Chr2 6734994 6775798 ATMSAT1 met1_10

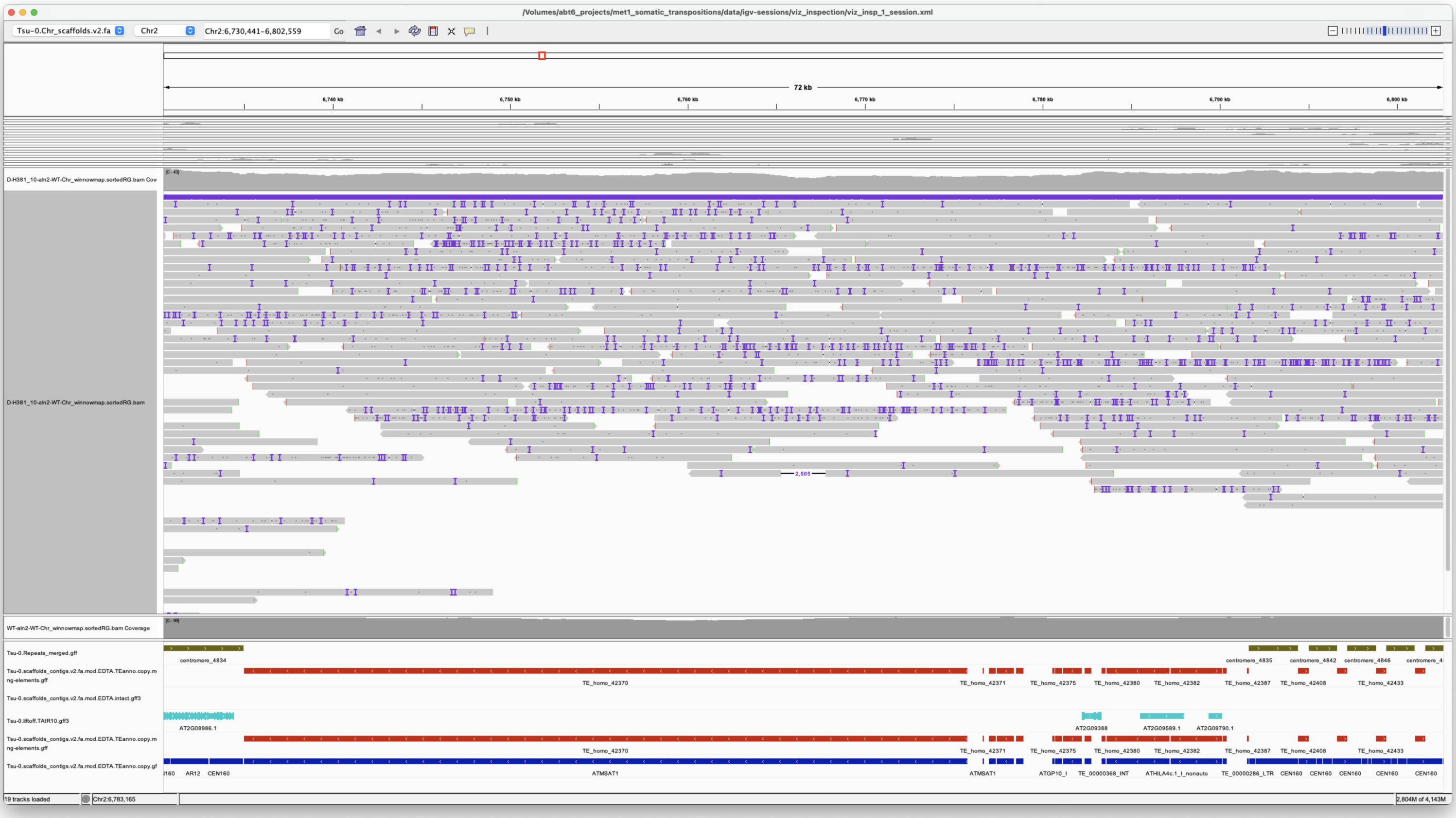

*SATELLITE*

Chr5 19872565 19877095 VANDAL21 met1_10

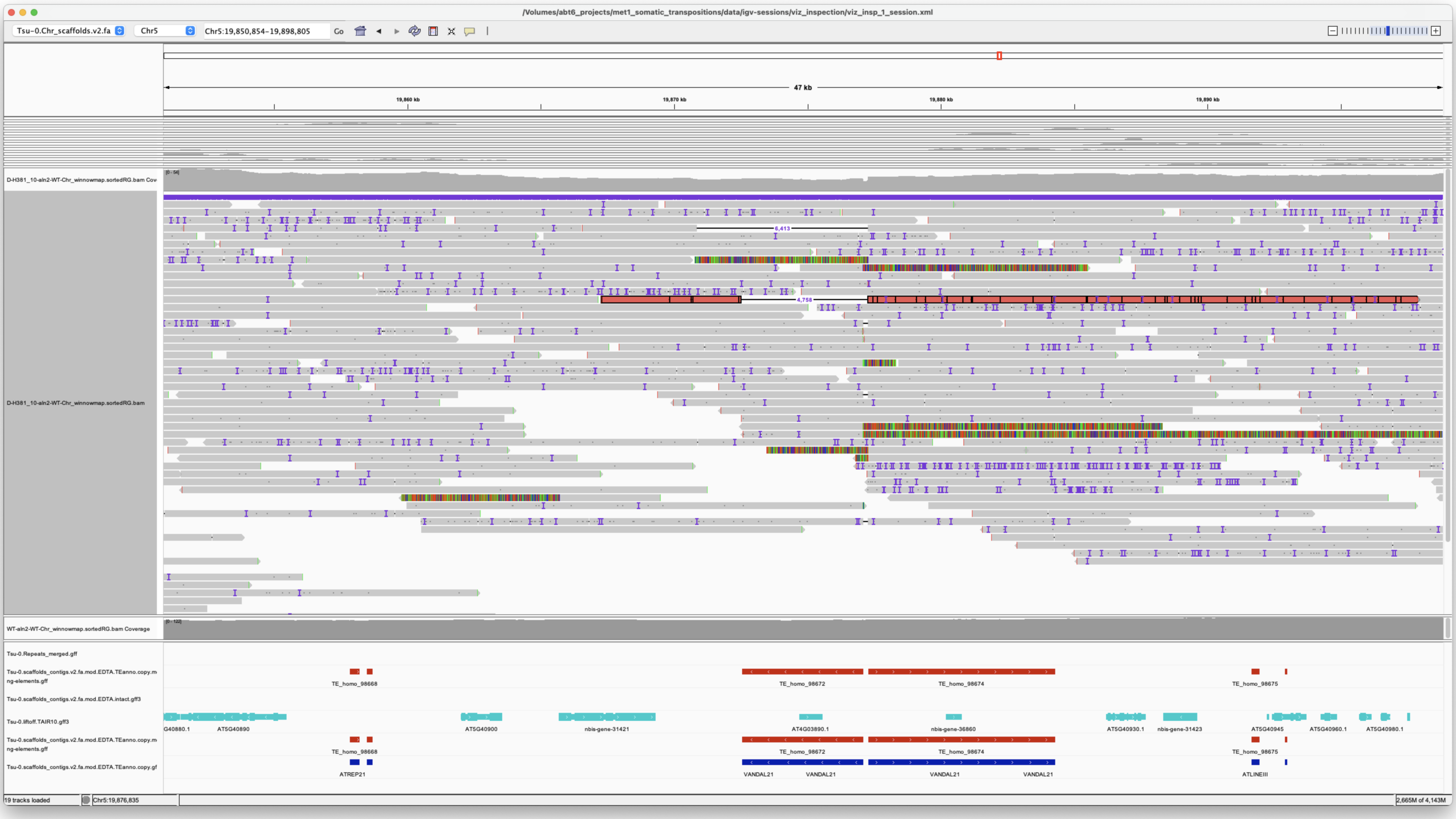


~200bp to the right of the element are also cut.

In this locus there are several rearrangements! this could be easily one of them

*Not Confirmed*

---

---

---

## WT

---

---

---
