## Additional File 4 for "Long-read detection of transposable element mobilization in the soma of hypomethylated *Arabidopsis thaliana* individuals": CIGAR_Insertions.pdf

### CIGAR_Insertions
### Merged elements

#### met1_01

> Chr1 32282511 32282511 m64079_221220_112036/81594785/ccs Chr5 875414 876434
> Chr5|875414|876433|ID=TE_MANUAL_02;Name=PutativePackTypeCACTAMuDR;classification=DNA/DTC;sequence_ontology=MANUAL;identity=MANUAL;method=MANUAL;ID=TE_MANUAL_02;sequence_ontology=MANUAL met1_01

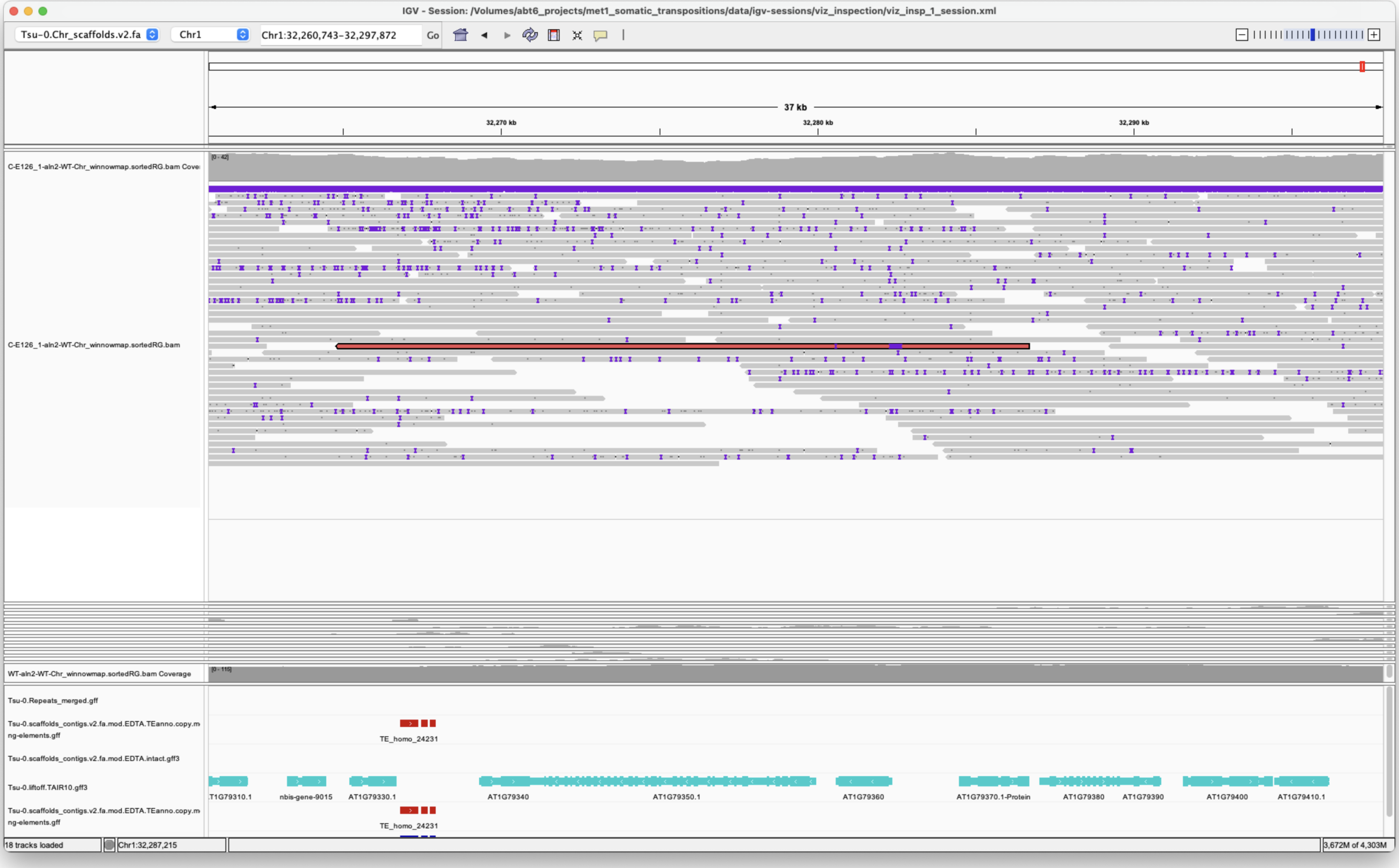


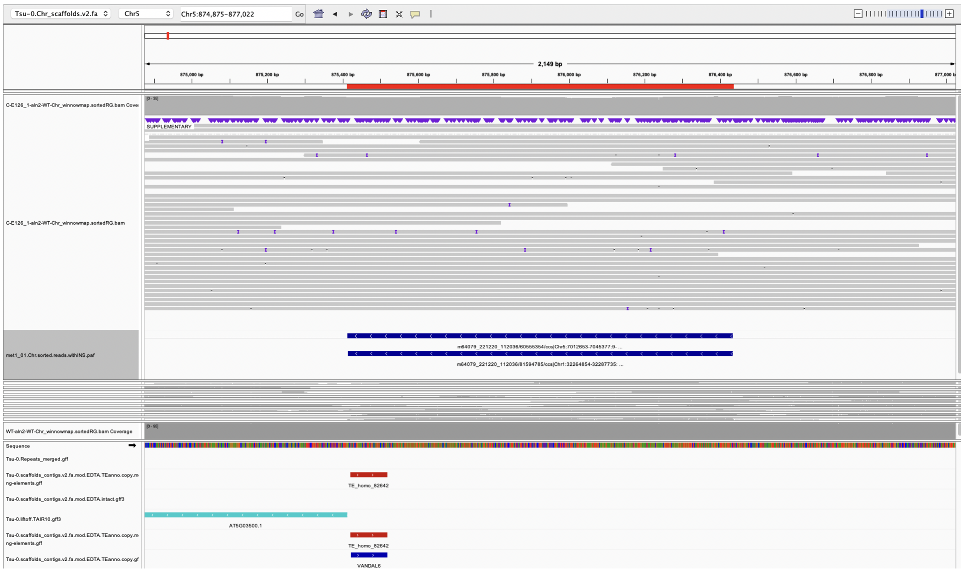


*Confirmed*

---

> Chr5 7026319 7026319 m64079_221220_112036/60555354/ccs Chr5 875413 876434
> Chr5|875414|876433|ID=TE_MANUAL_02;Name=PutativePackTypeCACTAMuDR;classification=DNA/DTC;sequence_ontology=MANUAL;identity=MANUAL;method=MANUAL;ID=TE_MANUAL_02;sequence_ontology=MANUAL met1_01

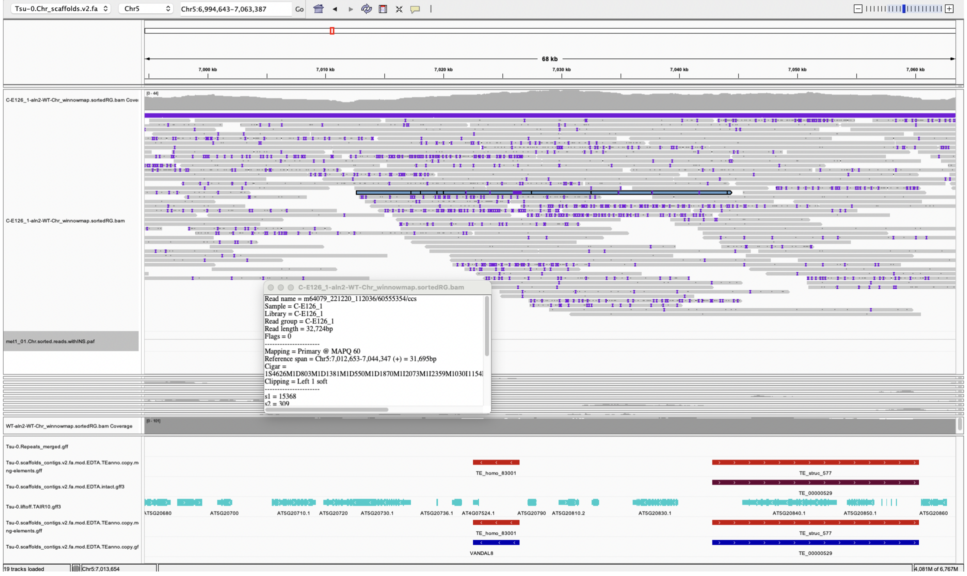


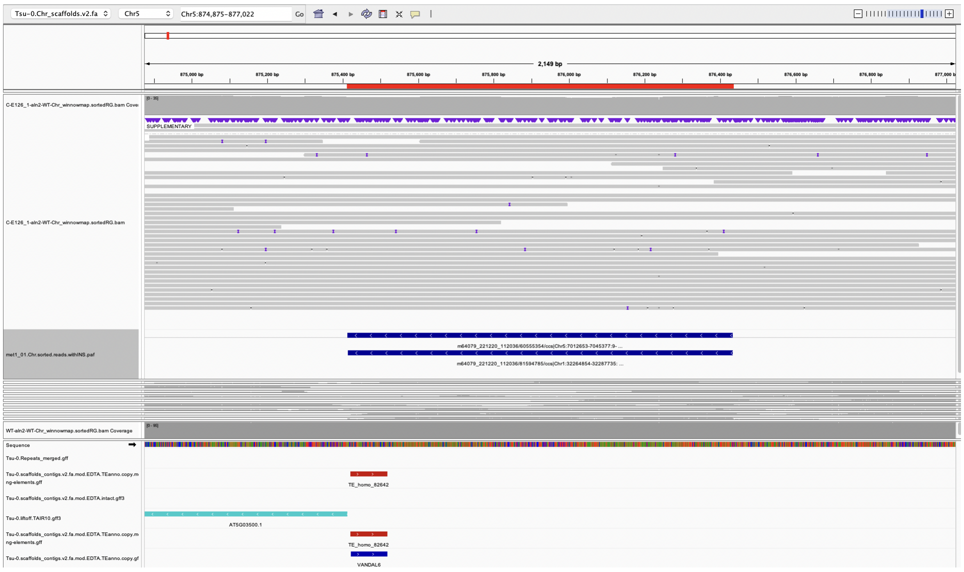


*Confirmed*

> Chr5 15902949 15902949 m64079_221220_112036/45091048/ccs Chr5 19152825 19160826 Chr5|19152829|19160825|ID=TE_homo_95640;Name=VANDAL21;classification=DNA/Mutator;sequence_ontology=SO:0002280;identity=0.976;method=homology;ID=TE_homo_98501;sequence_ontology=SO:0002280|ID=TE_homo_95641;Name=VANDAL21;classification=DNA/Mutator;sequence_ontology=SO:0002280;identity=0.966;method=homology;ID=TE_homo_98502;sequence_ontology=SO:0002280 met1_01

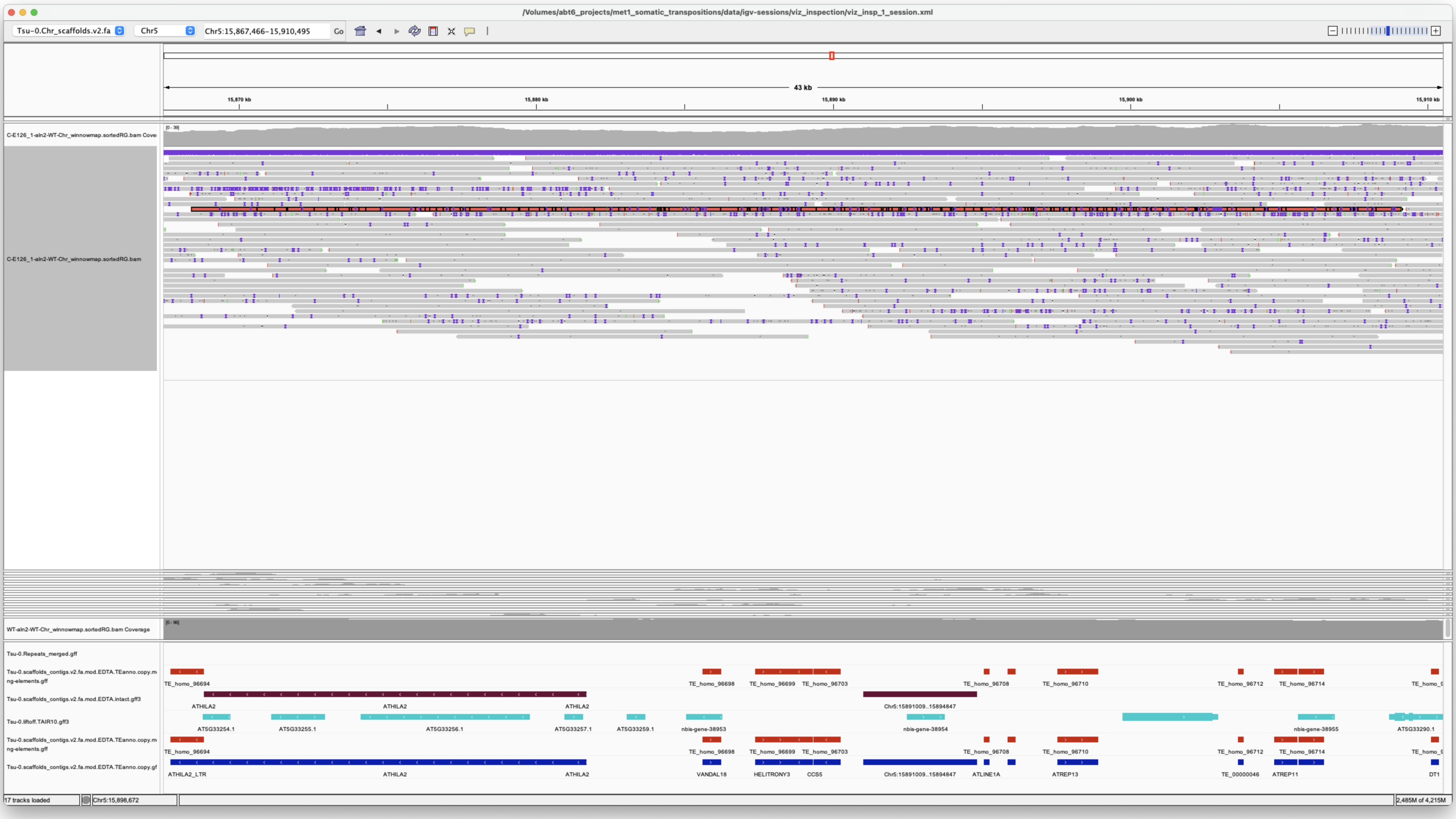

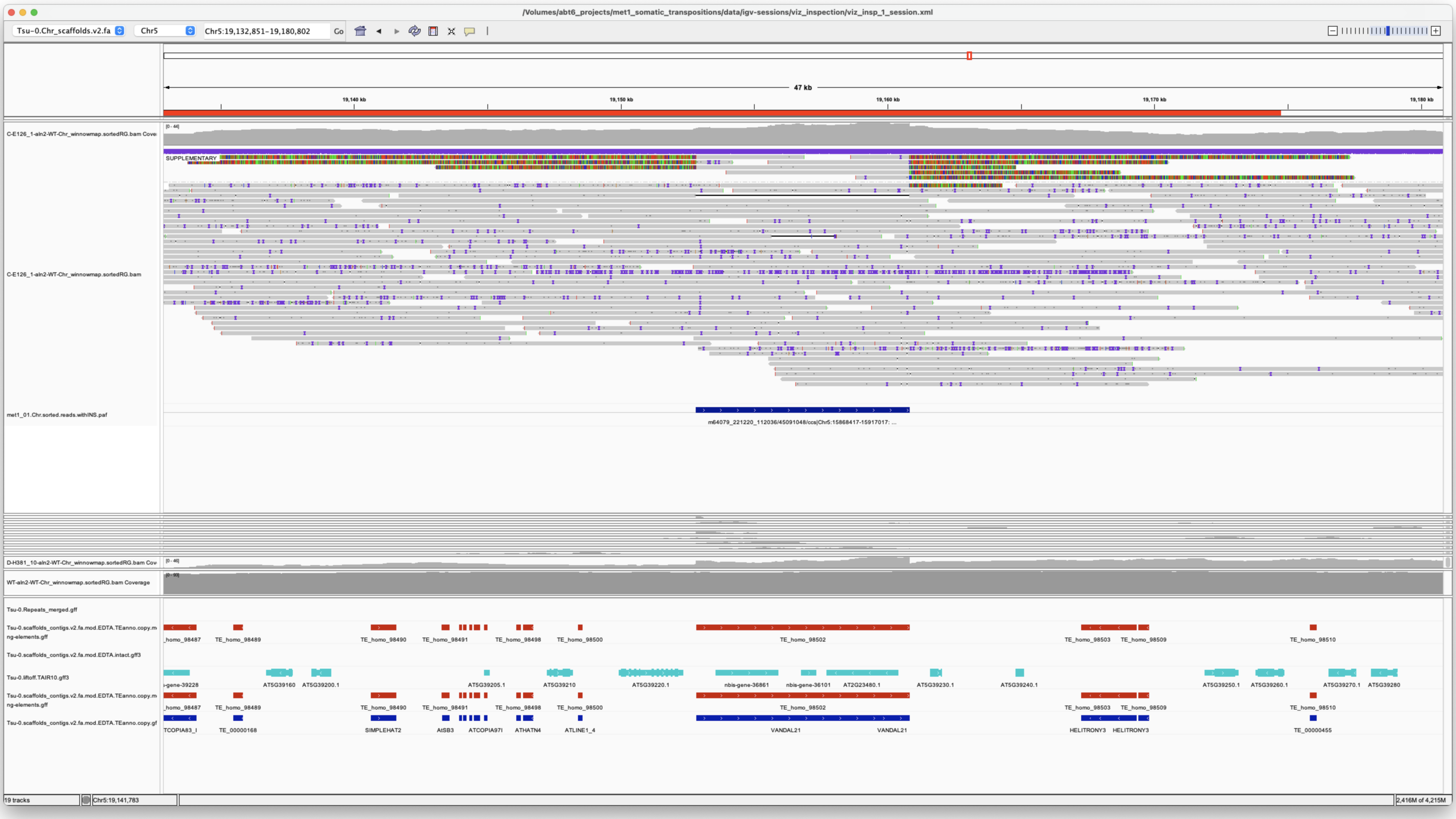


VANDAL21 insertion

*Confirmed*

---

---

---

#### met1_02

> Chr1 6039355 6039355 m64079_240212_113350/111674010/ccs Chr3 16344522 16352496 Chr3|16344522|16352496|ID=TE_homo_60420;Name=VANDAL6;classification=DNA/Mutator;sequence_ontology=SO:0002280;identity=0.969;method=homology;ID=TE_homo_62001;sequence_ontology=SO:0002280 met1_02

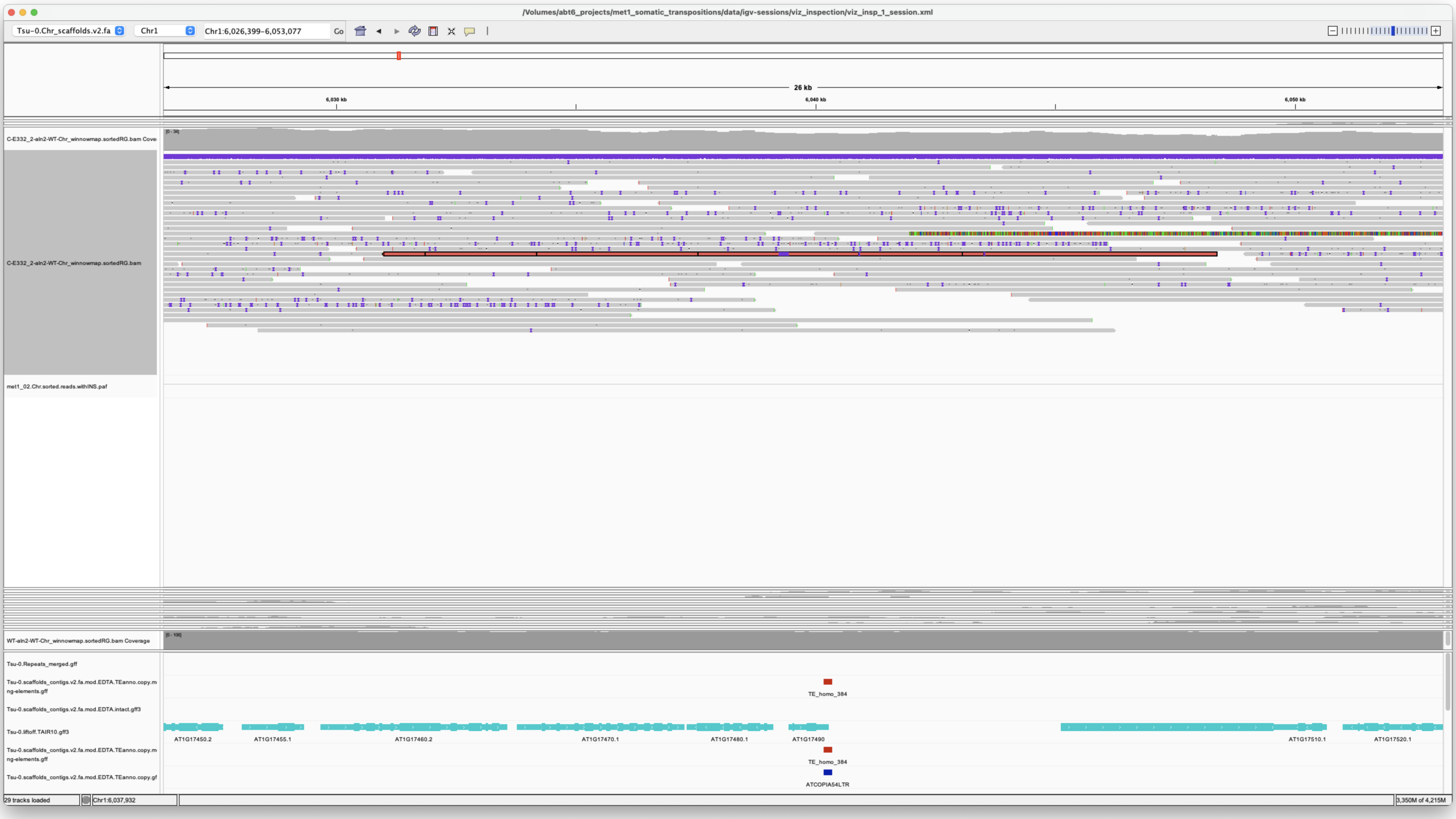


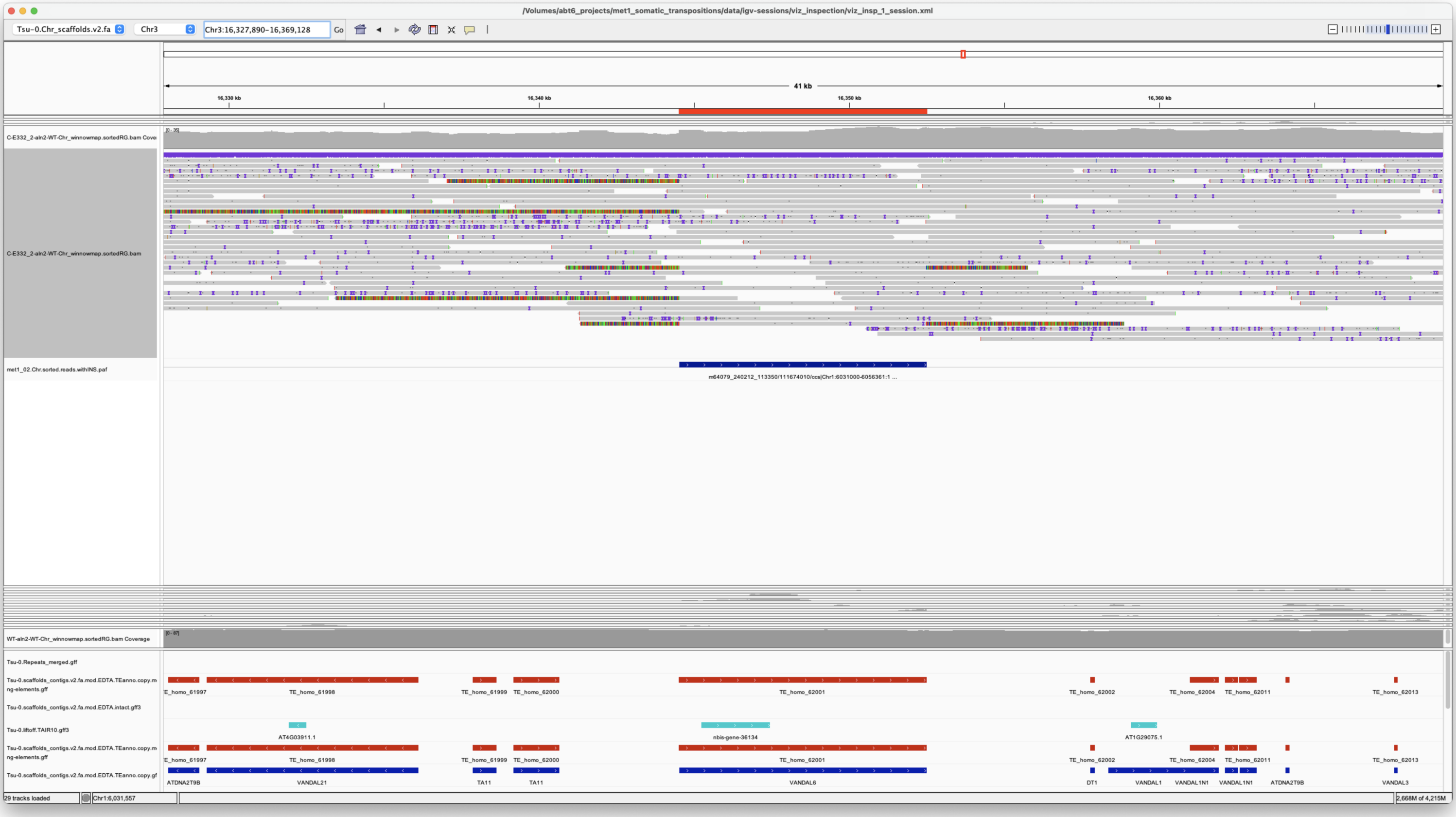


*Confirmed*

Chr1 20639051 20639051 m64079_221220_112036/15731028/ccs Chr5 875413 876434 Chr5|875414|876433|ID=TE_MANUAL_02;Name=PutativePackTypeCACTAMuDR;classification=DNA/DTC;sequence_ontology=MANUAL;identity=MANUAL;method=MANUAL;ID=TE_MANUAL_02;sequence_ontology=MANUAL met1_02

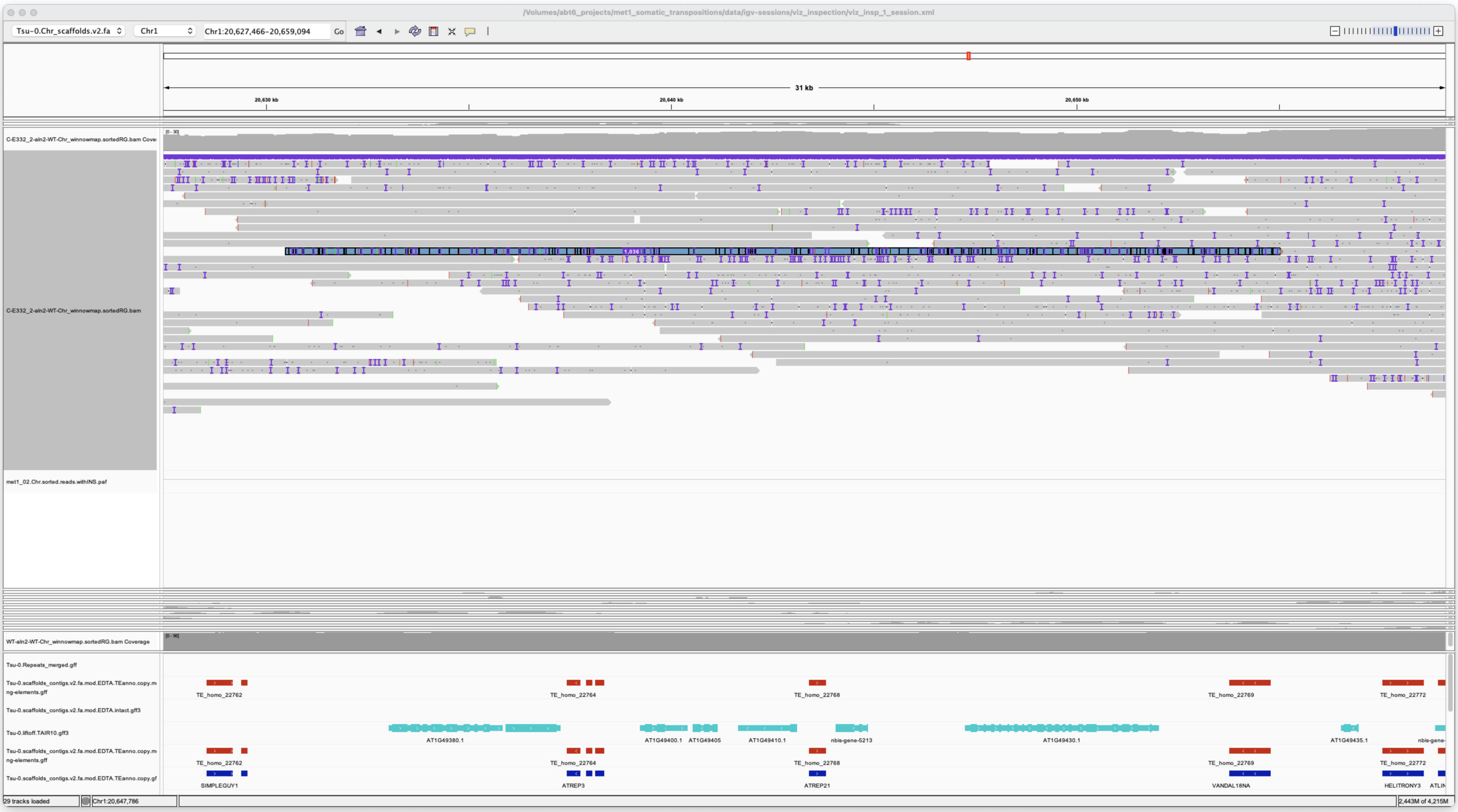


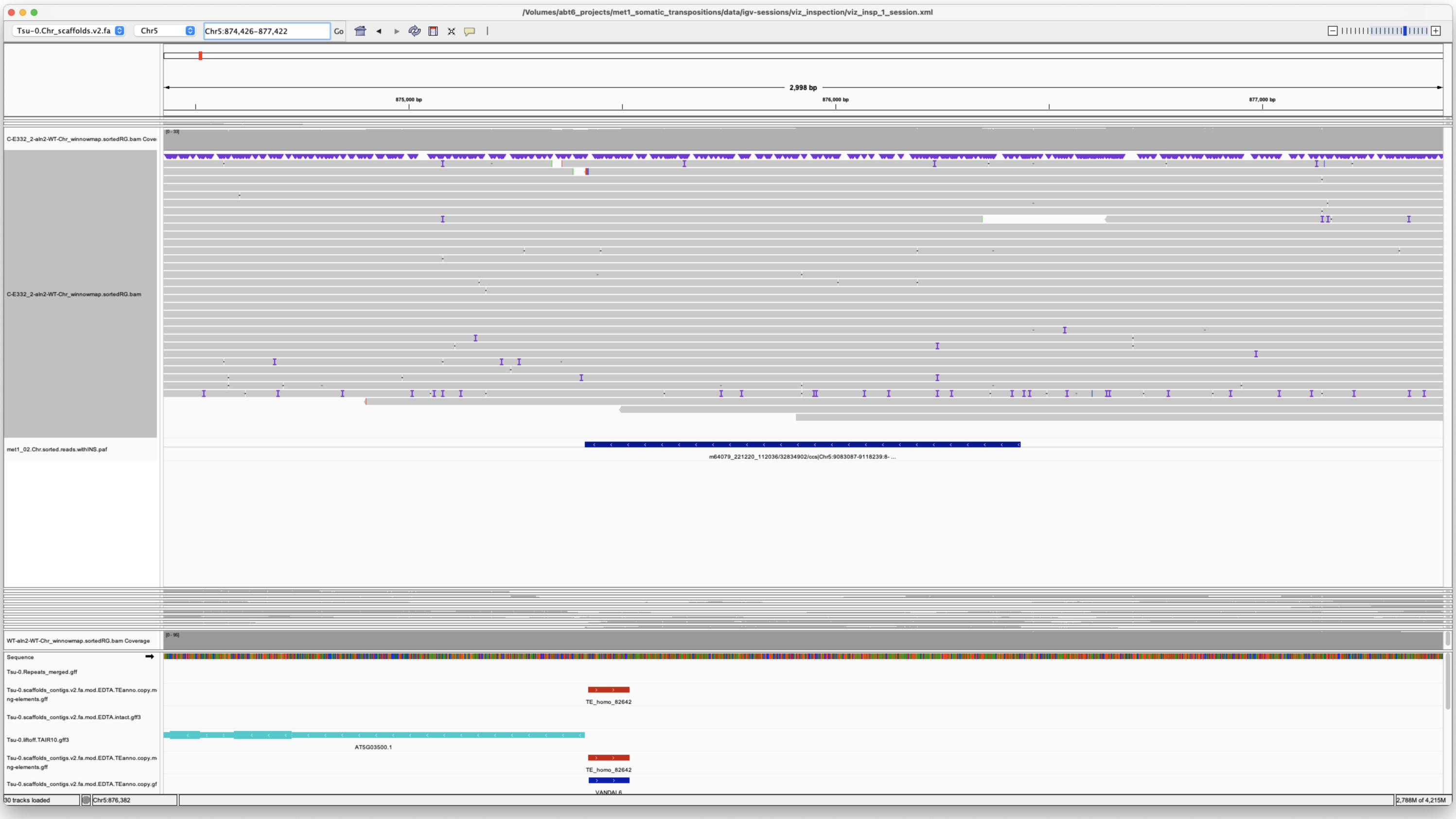


*Confirmed*

---

> Chr1 32269307 32269307 m64079_240212_113350/161613456/ccs Chr5 19152823 19160826 Chr5|19152829|19160825|ID=TE_homo_95640;Name=VANDAL21;classification=DNA/Mutator;sequence_ontology=SO:0002280;identity=0.976;method=homology;ID=TE_homo_98501;sequence_ontology=SO:0002280|ID=TE_homo_95641;Name=VANDAL21;classification=DNA/Mutator;sequence_ontology=SO:0002280;identity=0.966;method=homology;ID=TE_homo_98502;sequence_ontology=SO:0002280 met1_02

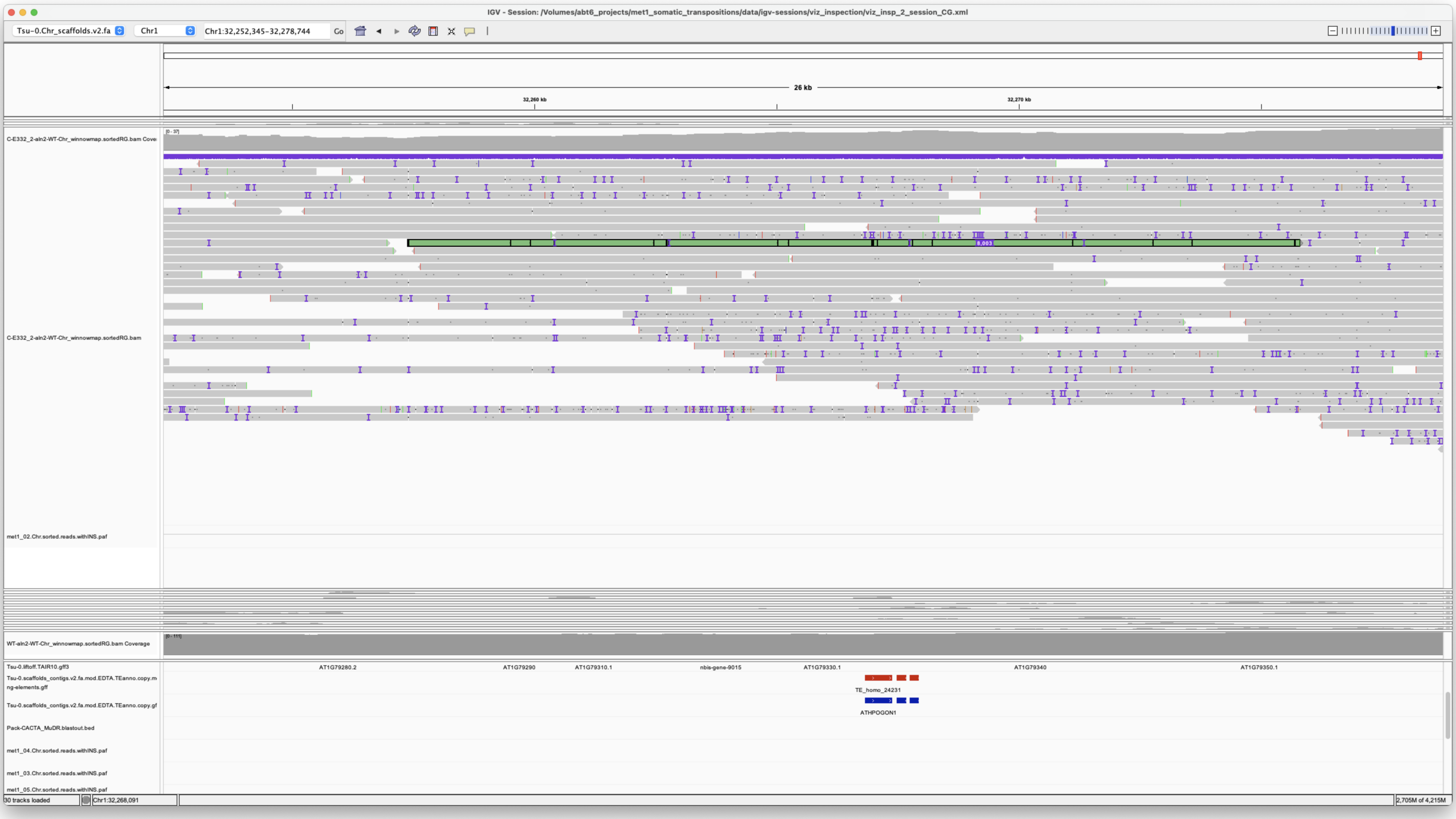


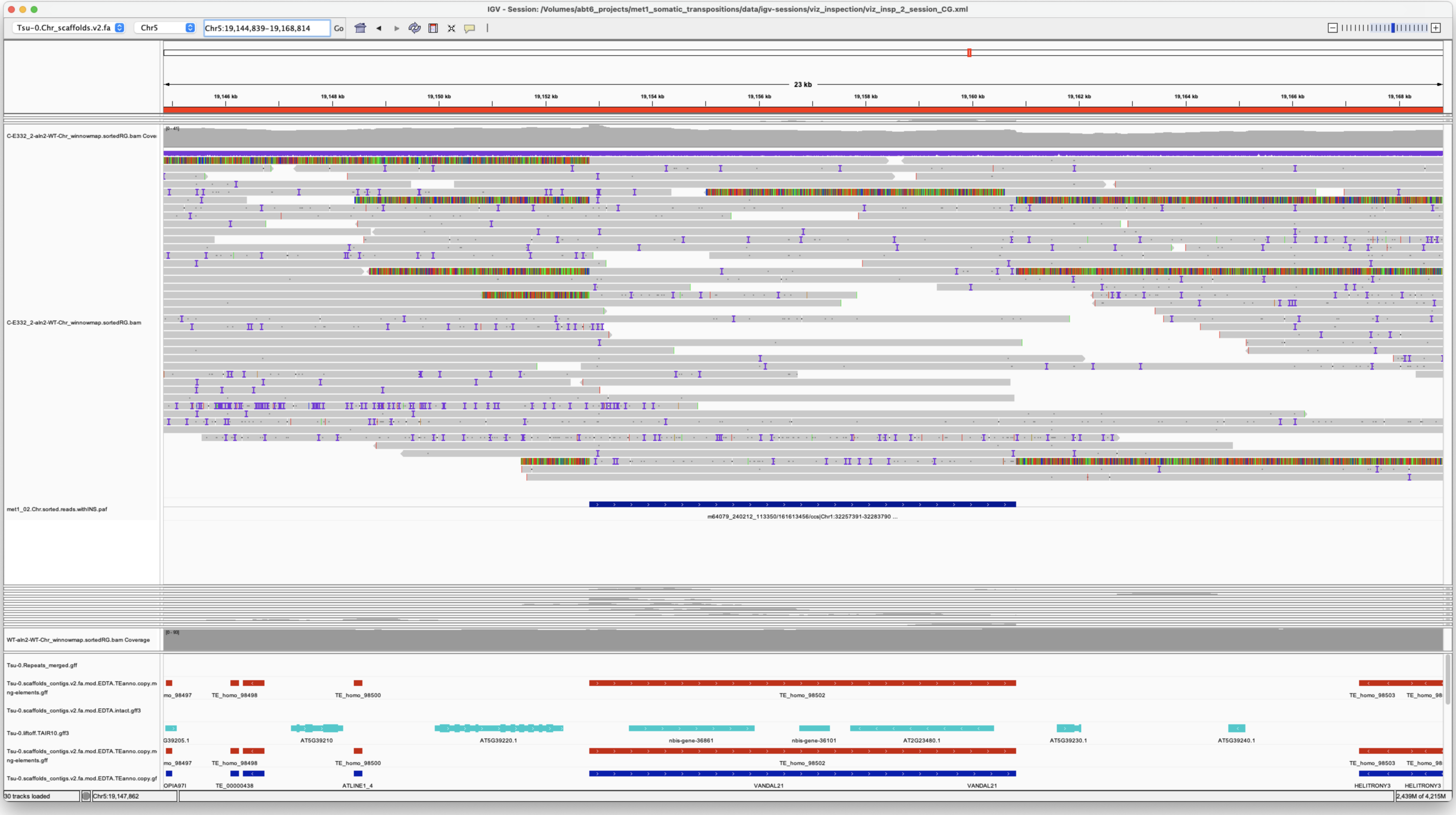


*Confirmed*

---

> Chr4 7023215 7023217
> m64079_221220_112036/29032572/ccs**m64079_221220_112036/41615485/ccs**m64079_221220_112036/42140263/ccs**m64079_221220_112036/60687065/ccs**m64079_240212_113350/120195183/ccs**m64079_240212_113350/15402943/ccs Chr4 7023215 7024283 .|-1|-2|.**.|-1|-2|.**.|-1|-2|.**.|-1|-2|.**.|-1|-2|.**.|-1|-2|. met1_02
> Chr4 7023225 7023225 m64079_221220_112036/5113584/ccs Chr4 7023224 7024291 .|-1|-2|. met1_02
> Chr4 7024209 7024209 m64079_221220_112036/47776303/ccs Chr4 7023140 7024207 .|-1|-2|. met1_02
> Chr4 7024279 7024279 m64079_221220_112036/10813820/ccs Chr4 7023210 7024277 .|-1|-2|. met1_02

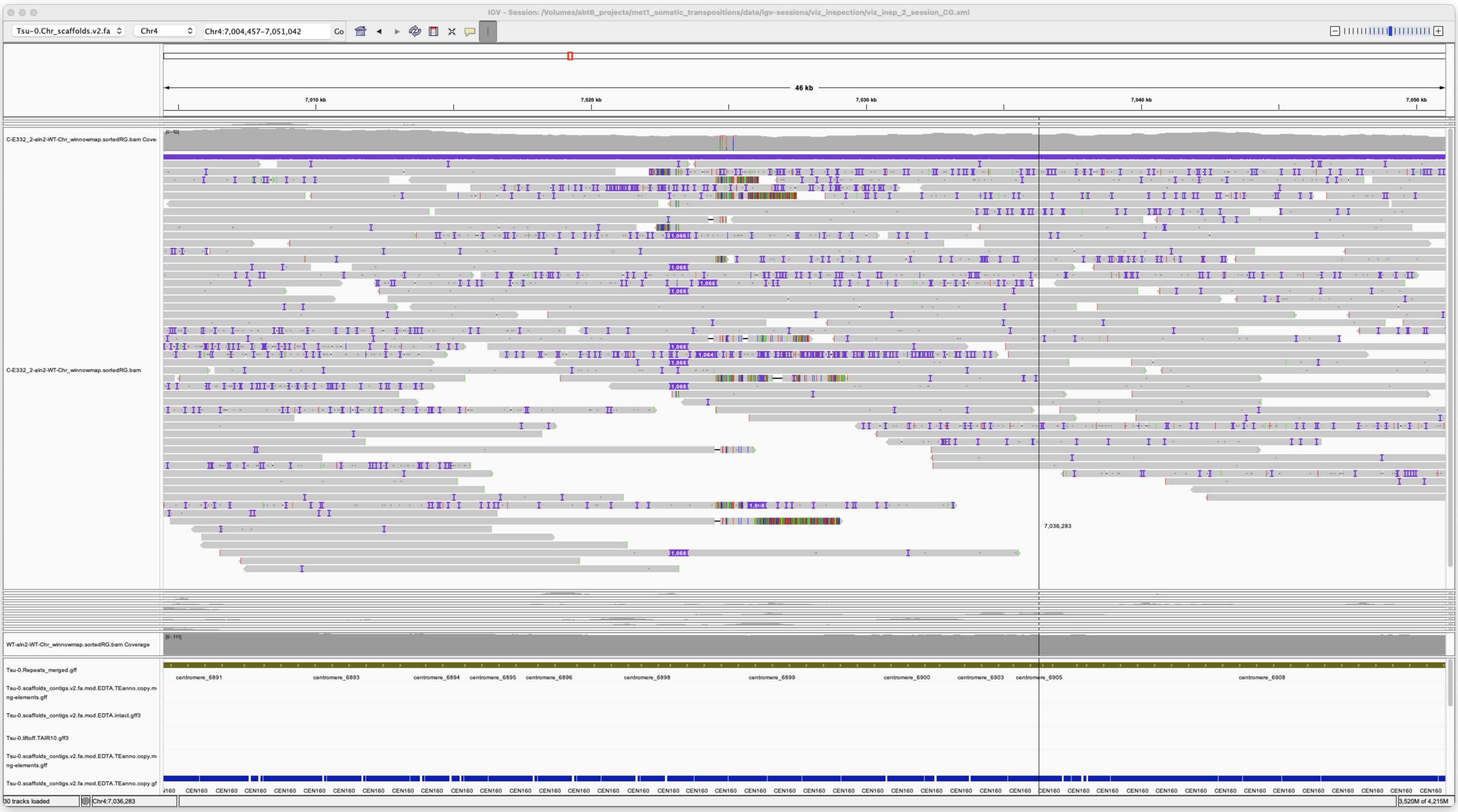

Centromeric rearrangements

*unsupported*

*Rearrangement*

---

> Chr5 9101100 9101100 m64079_221220_112036/32834902/ccs Chr5 875413 876434 Chr5|875414|876433|ID=TE_MANUAL_02;Name=PutativePackTypeCACTAMuDR;classification=DNA/DTC;sequence_ontology=MANUAL;identity=MANUAL;method=MANUAL;ID=TE_MANUAL_02;sequence_ontology=MANUAL met1_02

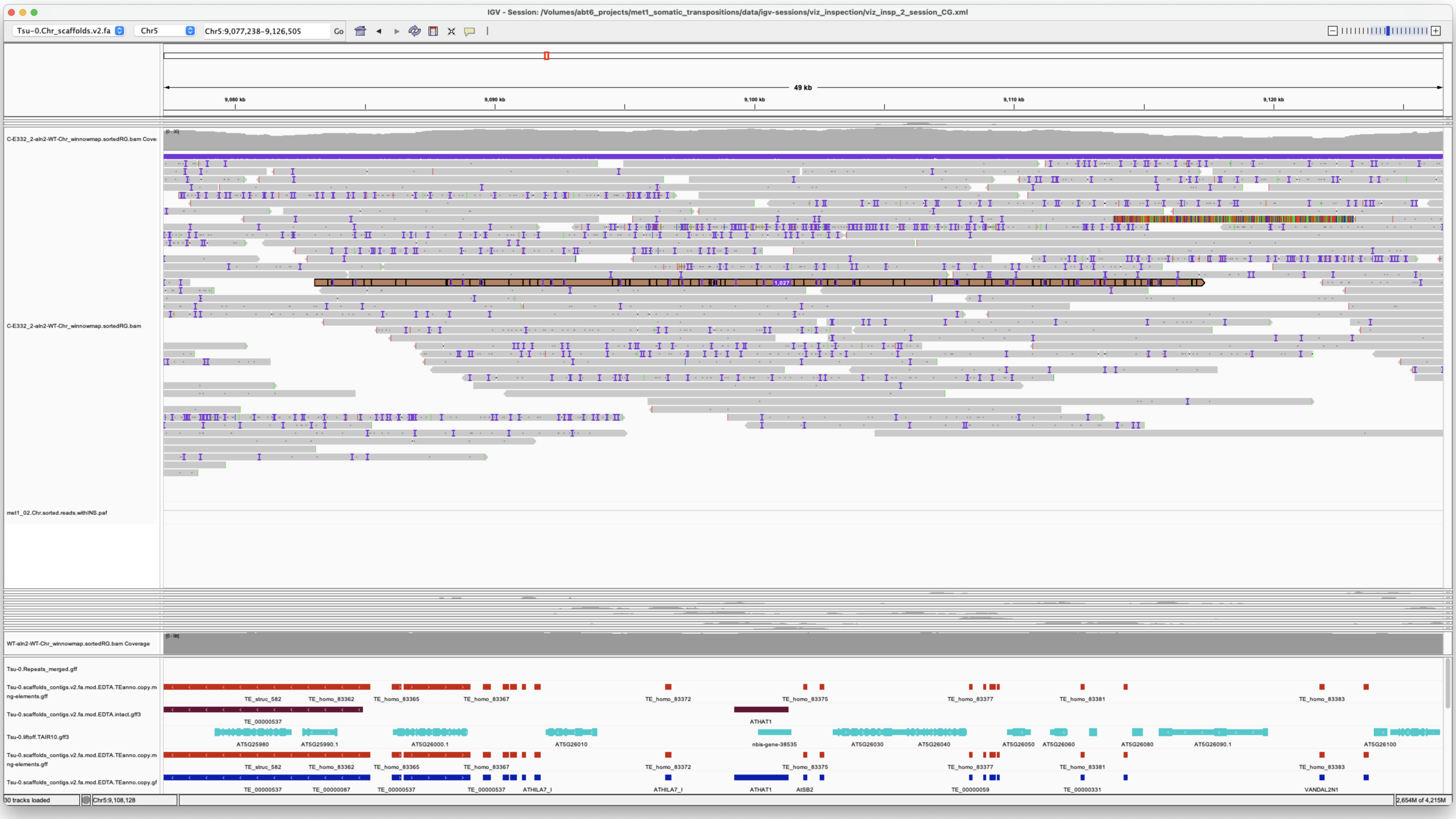

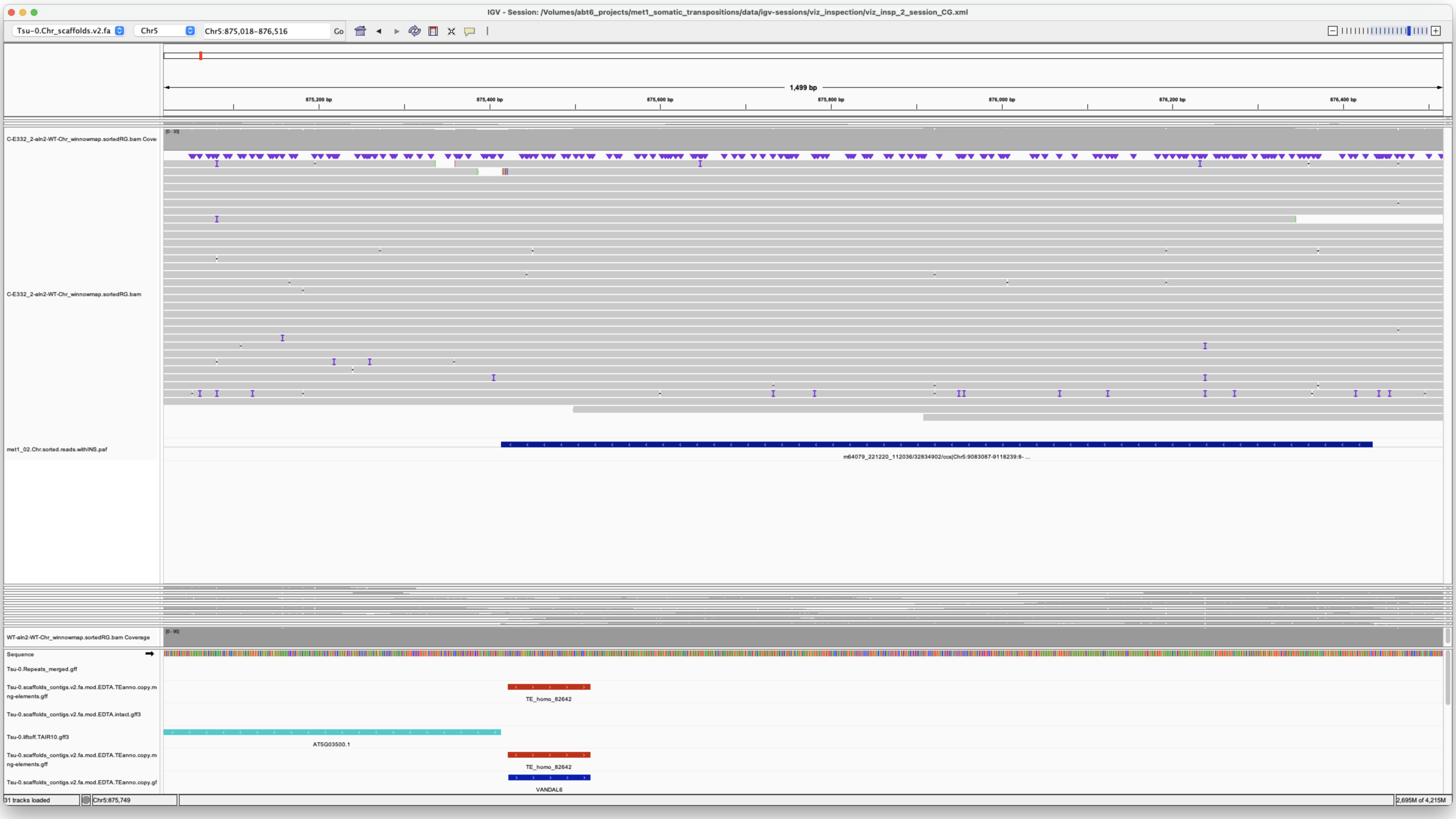

*Confirmed*

#### met1_03

> Chr2 6768455 6768455 m64079_221220_112036/112396489/ccs Chr2 6768454 6769622 Chr2|6734994|6775797|ID=TE_homo_41523;Name=ATMSAT1;classification=Satellite/Satellite;sequence_ontology=SO:0000005;identity=0.934;method=homology;ID=TE_homo_42370;sequence_ontology=SO:0000005 met1_03

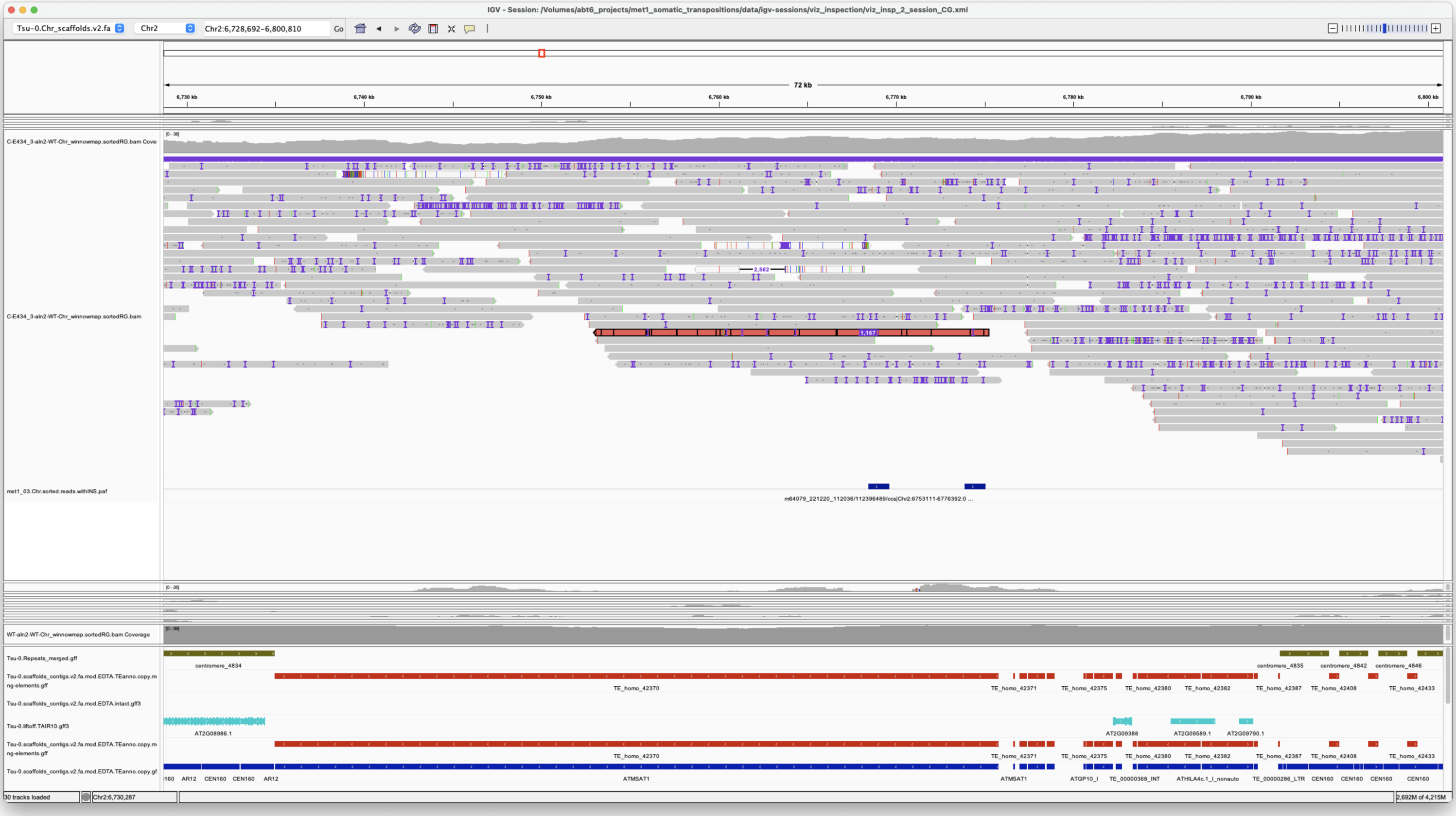

Satellite rearrangement

*unsupported*

*Rearrangement*

### met1_04

**NONE**

### met1_05

> Chr1 7025171 7025171 m64079_221220_112036/97912206/ccs Chr3 16344522 16352504 Chr3|16344522|16352496|ID=TE_homo_60420;Name=VANDAL6;classification=DNA/Mutator;sequence_ontology=SO:0002280;identity=0.969;method=homology;ID=TE_homo_62001;sequence_ontology=SO:0002280 met1_05

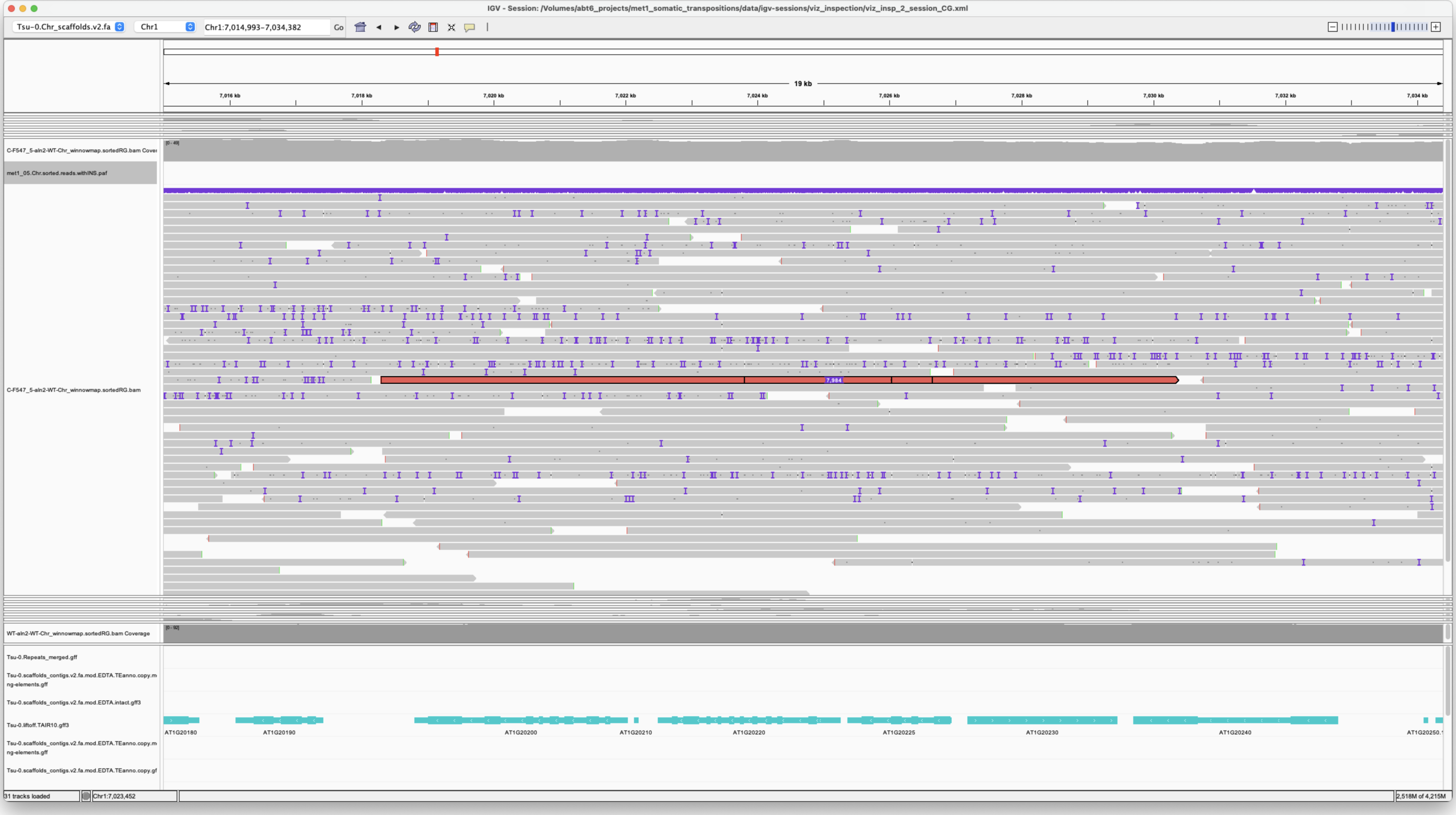


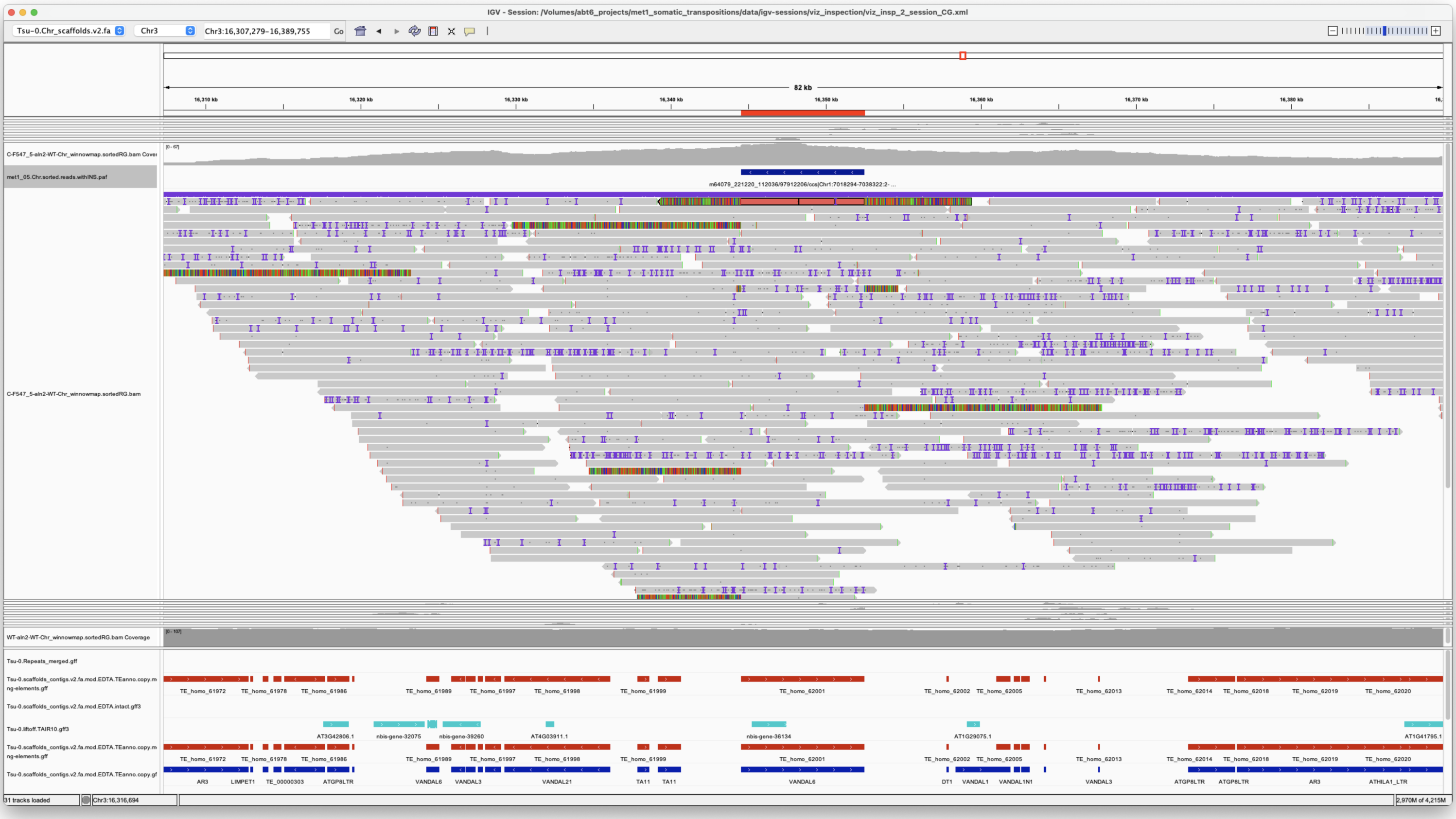

*Confirmed*

Chr2 11071303 11071303 m64079_240212_113350/170396218/ccs Chr1 11941106 11946436 Chr1|11941106|11946435|ID=LTRRT_5;Name=ATCOPIA93.2_Evade;Classification=LTR/Copia;Sequence_ontology=SO:0002264;ltr_identity=1.0000;Method=structural;motif=TACA;tsd=ATATG met1_05

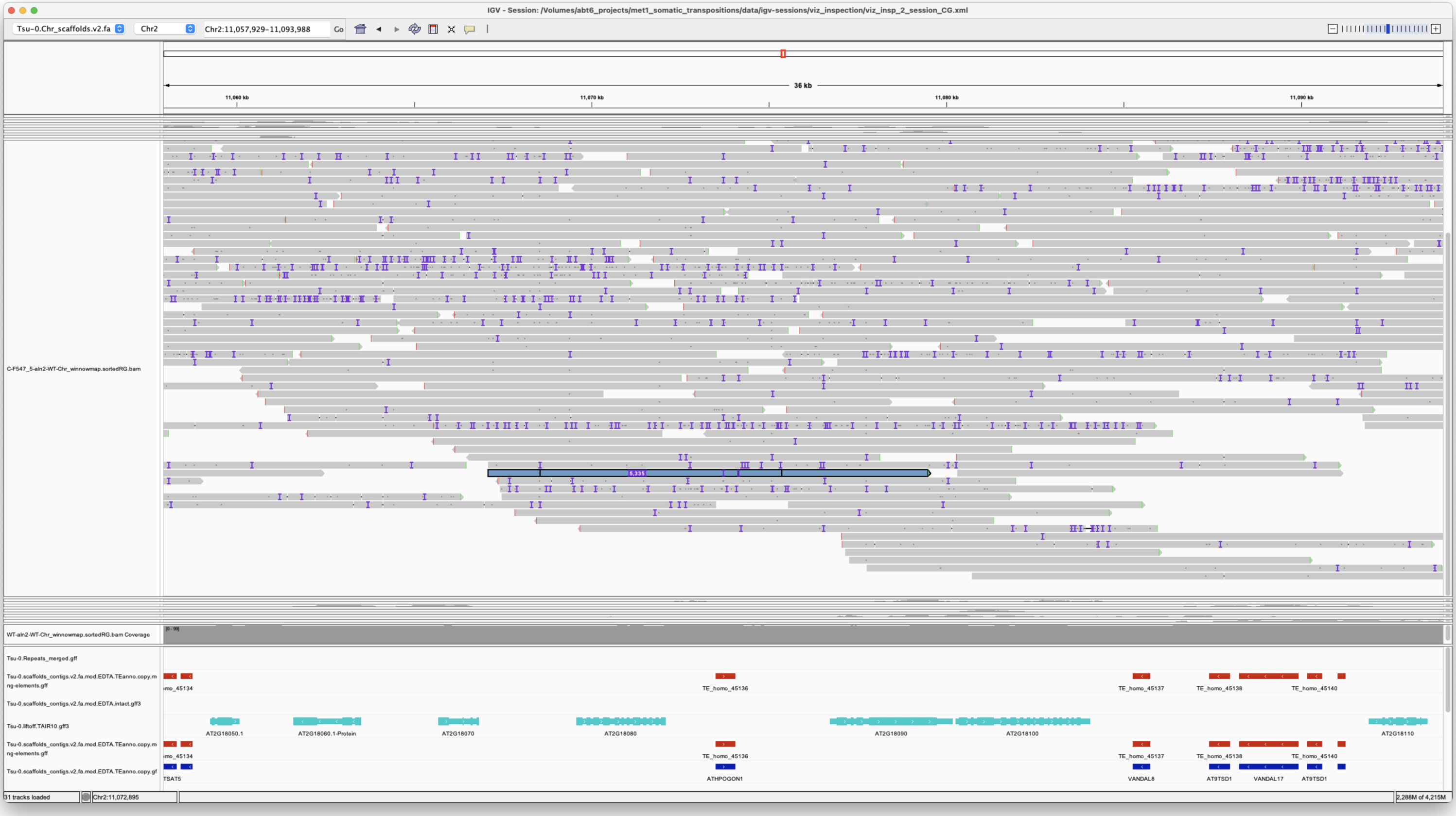


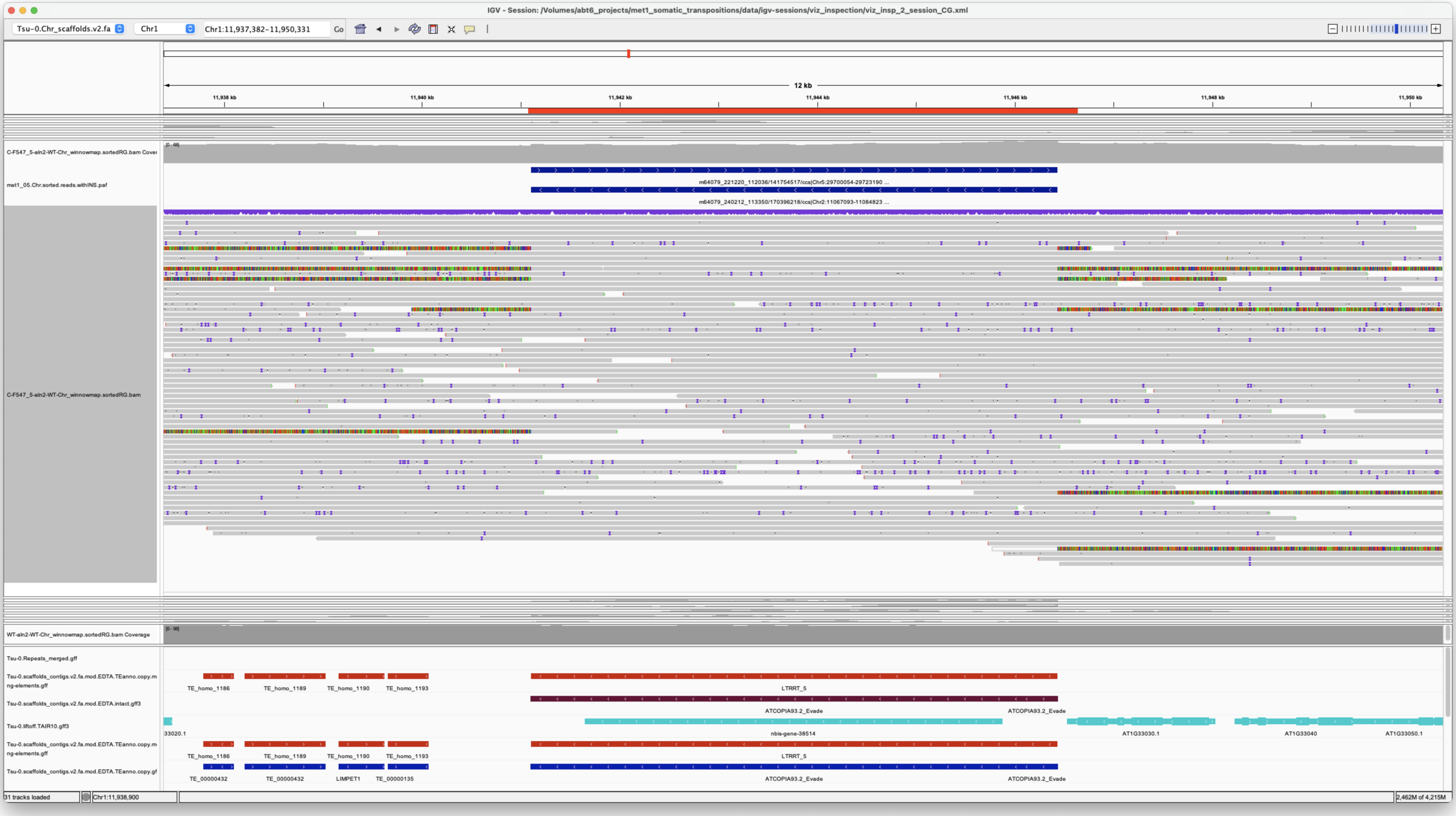


*Confirmed*

Chr3 2809693 2809693 m64079_240212_113350/9307420/ccs Chr5 875413 876434 Chr5|875414|876433|ID=TE_MANUAL_02;Name=PutativePackTypeCACTAMuDR;classification=DNA/DTC;sequence_ontology=MANUAL;identity=MANUAL;method=MANUAL;ID=TE_MANUAL_02;sequence_ontology=MANUAL met1_05

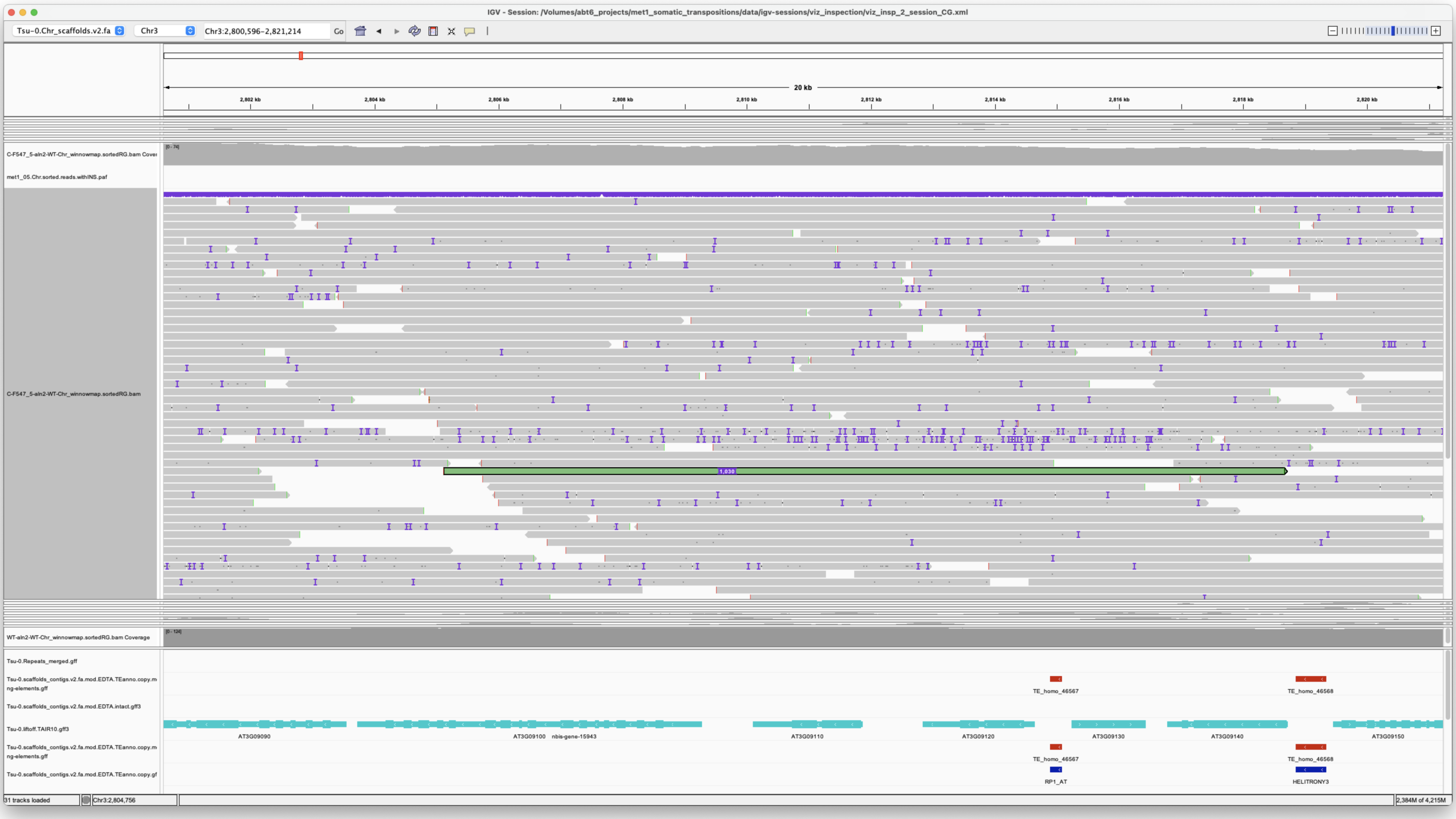


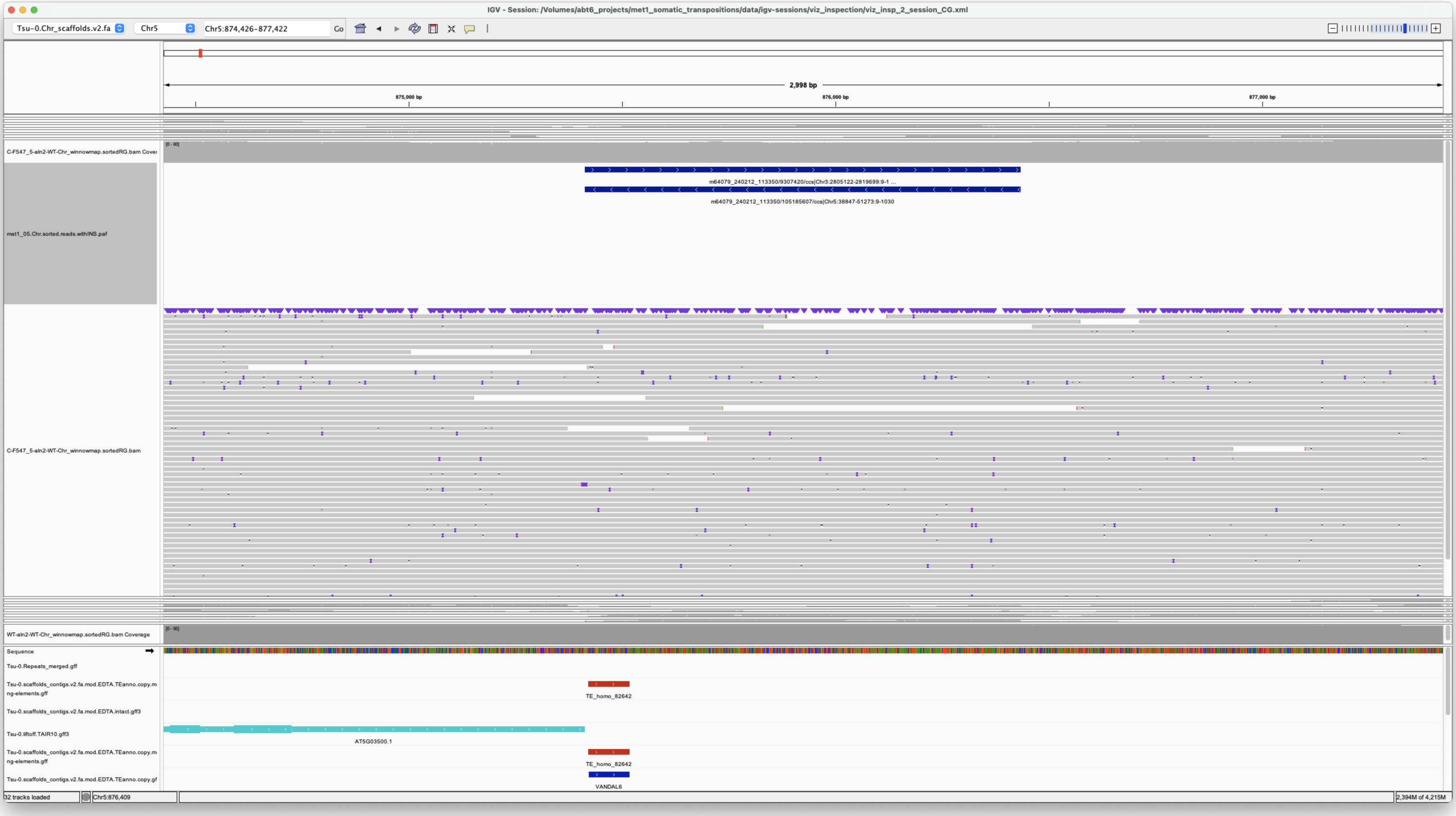


Pack-Type

*Confirmed*

---

> Chr5 49106 49106 m64079_240212_113350/105185607/ccs Chr5 875413 876434 Chr5|875414|876433|ID=TE_MANUAL_02;Name=PutativePackTypeCACTAMuDR;classification=DNA/DTC;sequence_ontology=MANUAL;identity=MANUAL;method=MANUAL;ID=TE_MANUAL_02;sequence_ontology=MANUAL met1_05

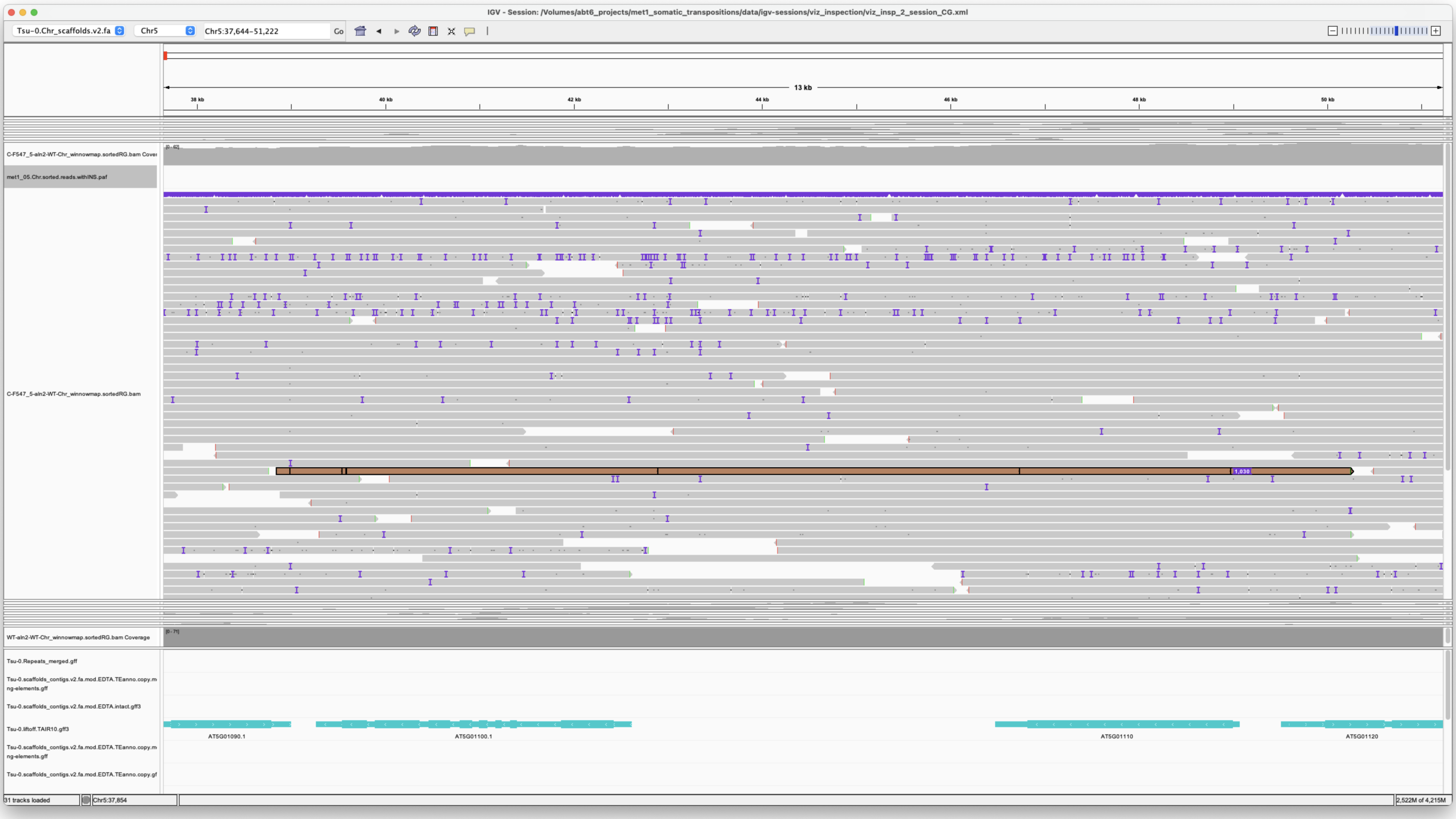


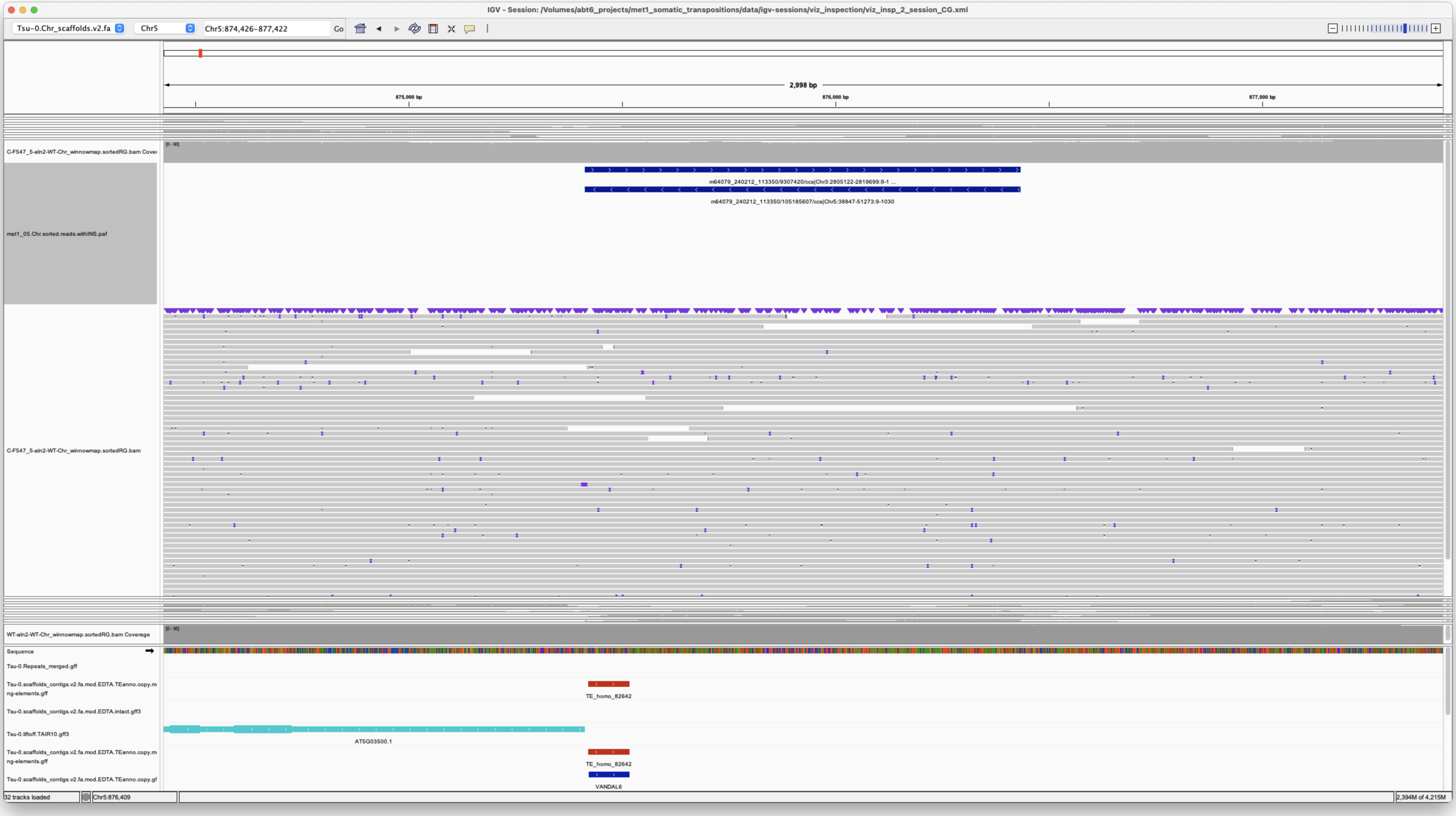


Pack-Type

*Confirmed*

---

> Chr5 28025714 28025714 m64079_221220_112036/166528667/ccs Chr5 19152825 19160826 Chr5|19152829|19160825|ID=TE_homo_95640;Name=VANDAL21;classification=DNA/Mutator;sequence_ontology=SO:0002280;identity=0.976;method=homology;ID=TE_homo_98501;sequence_ontology=SO:0002280|ID=TE_homo_95641;Name=VANDAL21;classification=DNA/Mutator;sequence_ontology=SO:0002280;identity=0.966;method=homology;ID=TE_homo_98502;sequence_ontology=SO:0002280 met1_05

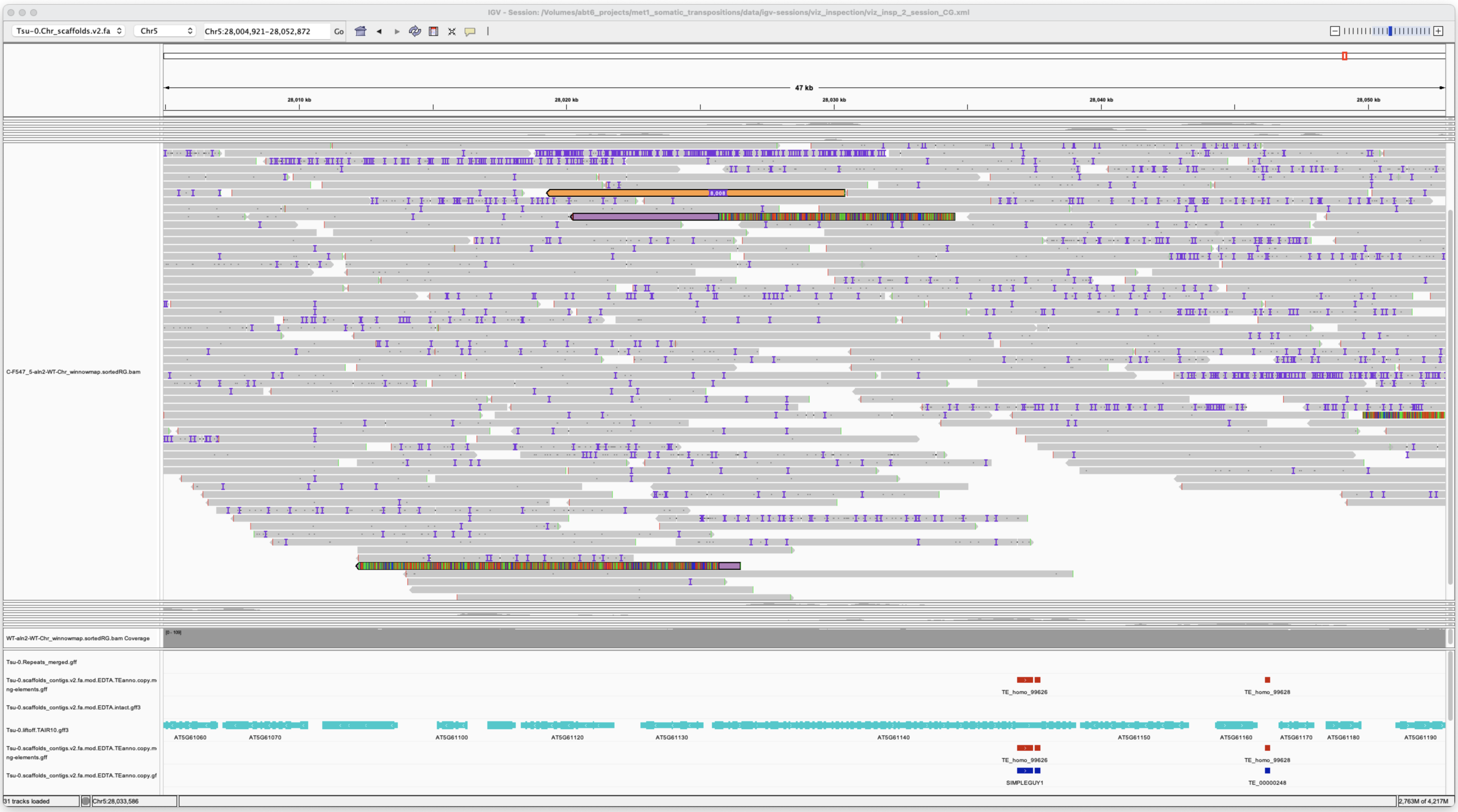

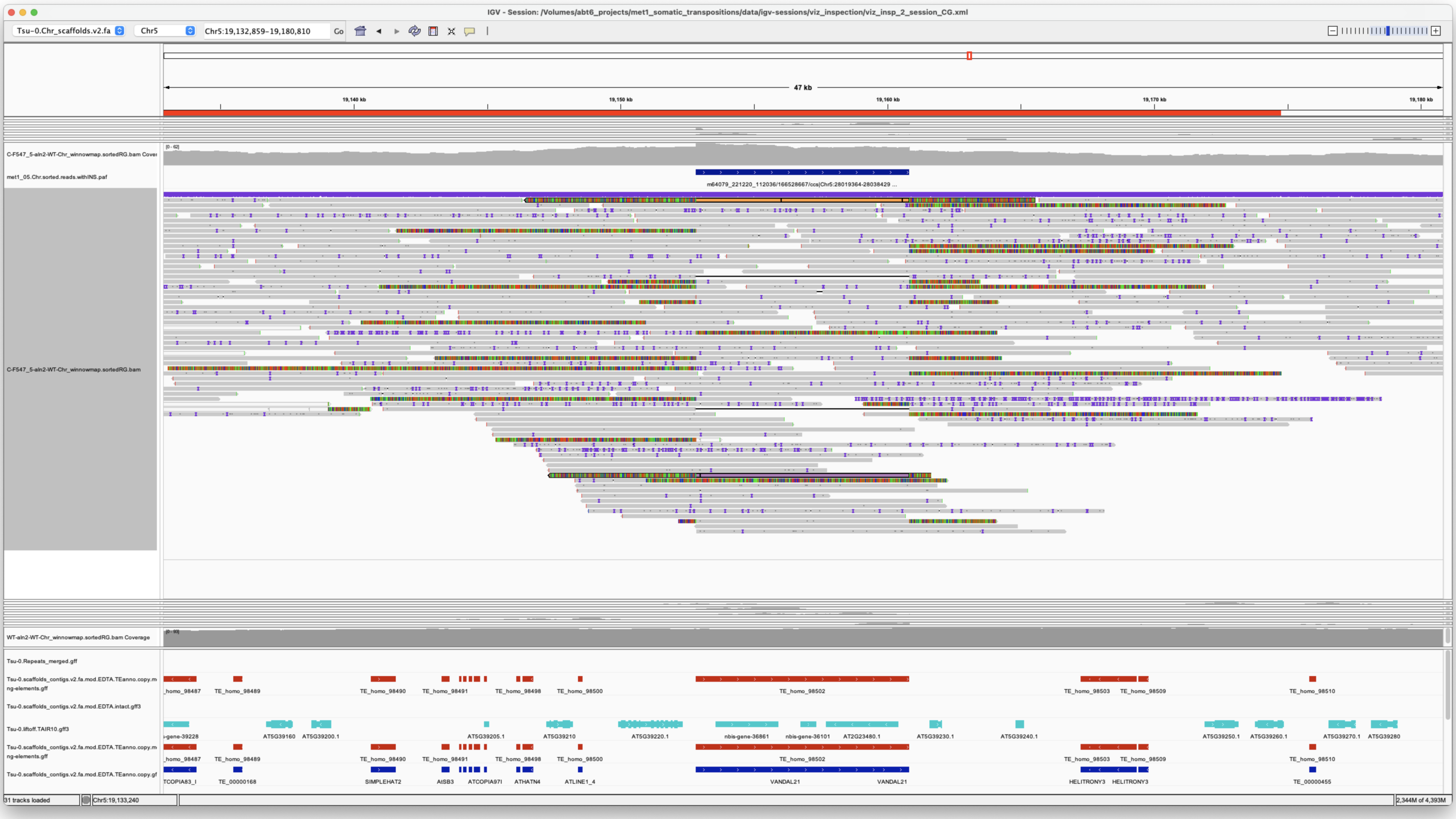

**2X coverage for insertion**

*Confirmed*

---

> Chr5 29702875 29702875 m64079_221220_112036/141754517/ccs Chr1 11941106 11946436
> Chr1|11941106|11946435|ID=LTRRT_5;Name=ATCOPIA93.2_Evade;Classification=LTR/Copia;Sequence_ontology=SO:0002264;ltr_identity=1.0000;Method=structural;motif=TACA;tsd=ATATG met1_05

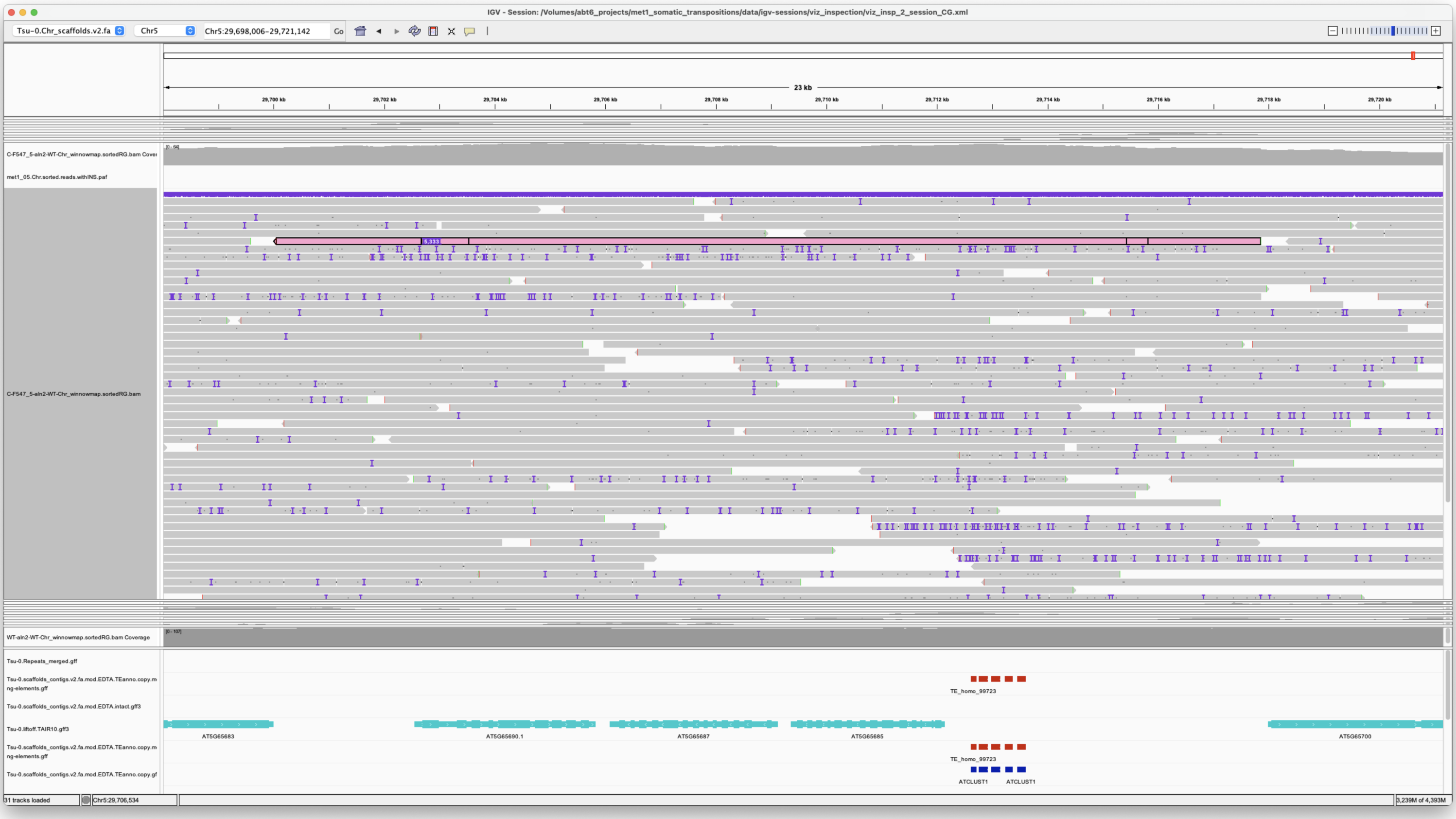


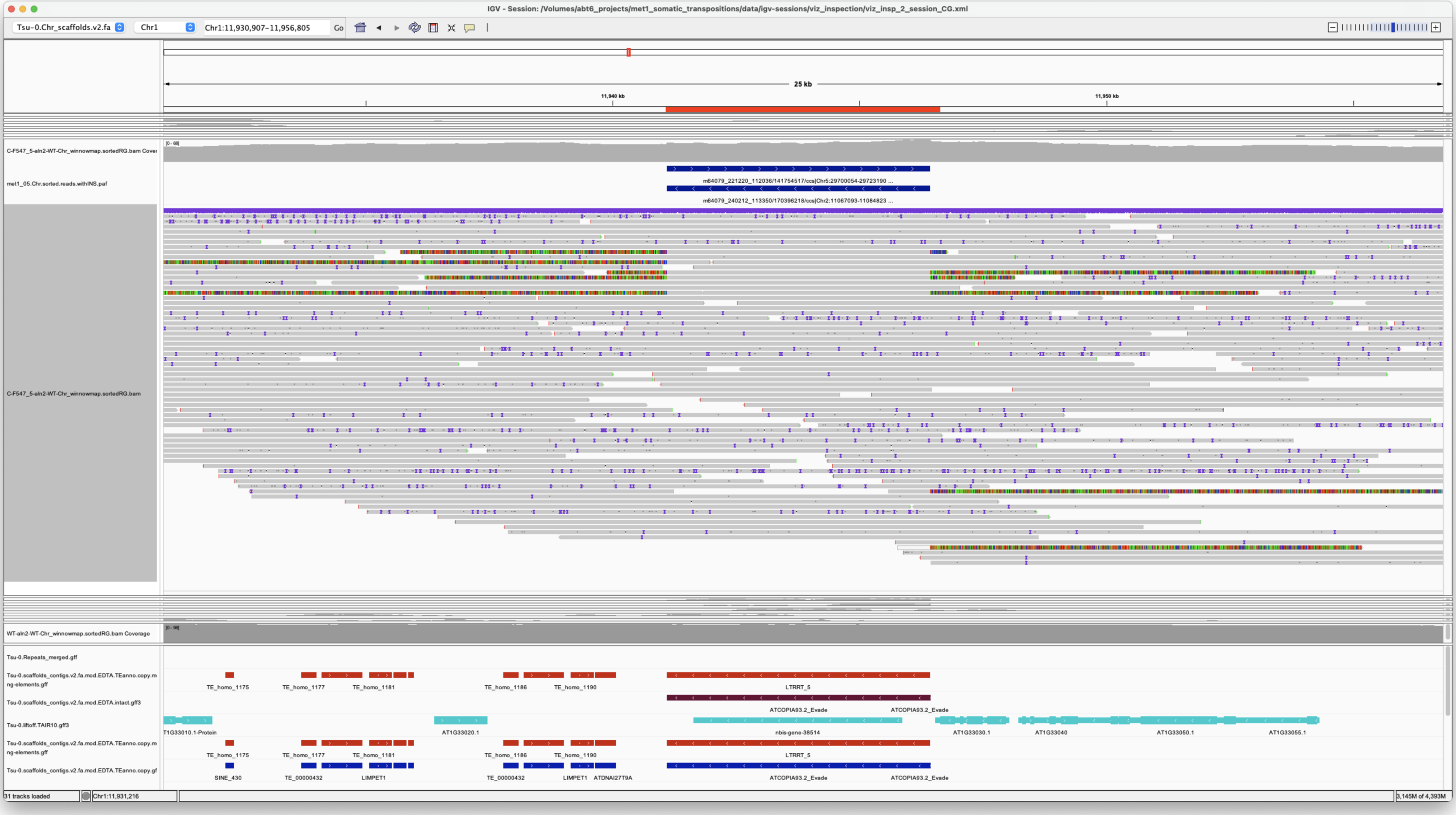


*Confirmed*

---

---

#### met1_06

---

> Chr1 17891752 17891752 m64079_221220_112036/12321302/ccs Chr5 19152826 19160834 Chr5|19152829|19160825|ID=TE_homo_95640;Name=VANDAL21;classification=DNA/Mutator;sequence_ontology=SO:0002280;identity=0.976;method=homology;ID=TE_homo_98501;sequence_ontology=SO:0002280|ID=TE_homo_95641;Name=VANDAL21;classification=DNA/Mutator;sequence_ontology=SO:0002280;identity=0.966;method=homology;ID=TE_homo_98502;sequence_ontology=SO:0002280 met1_06

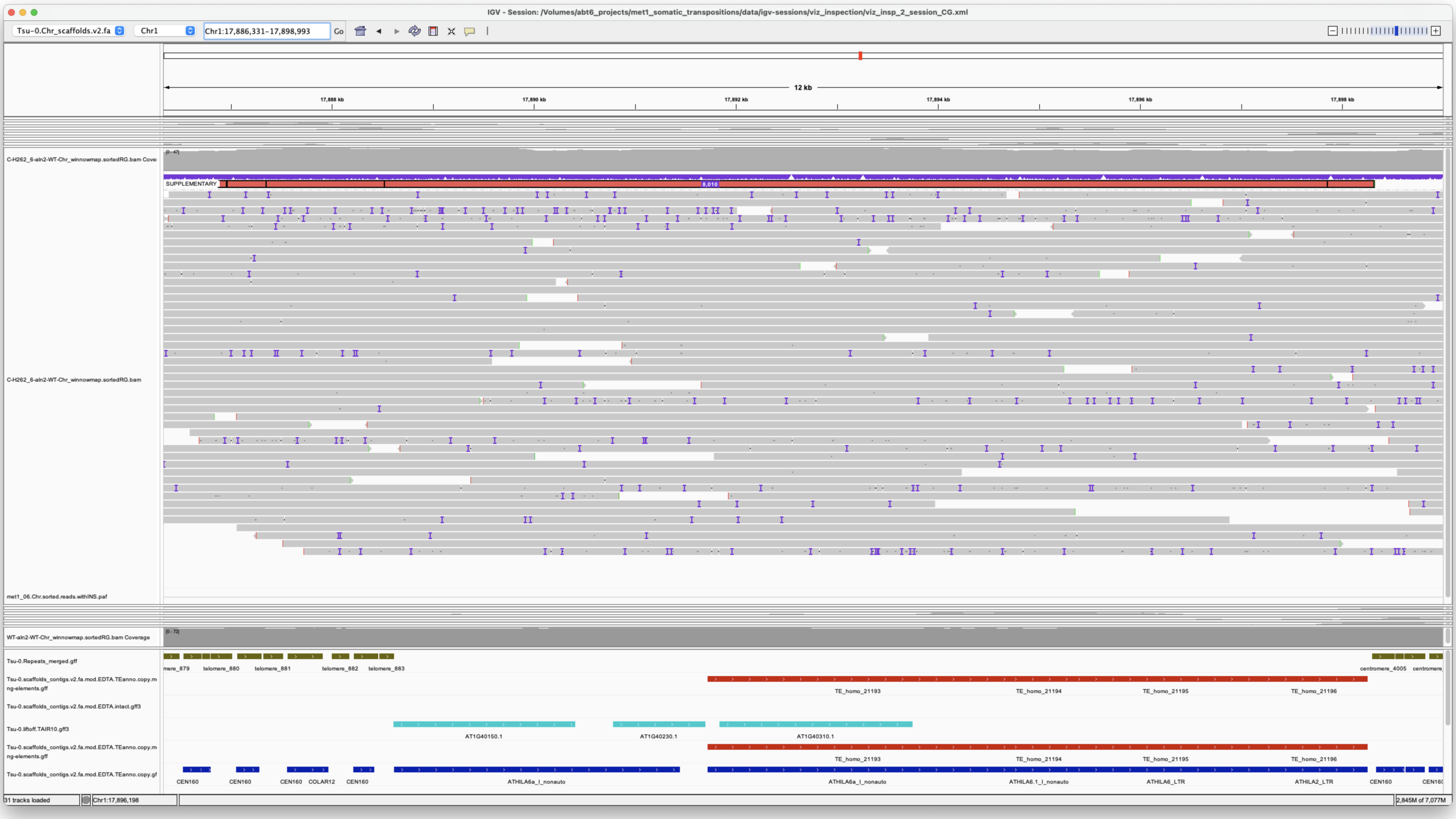


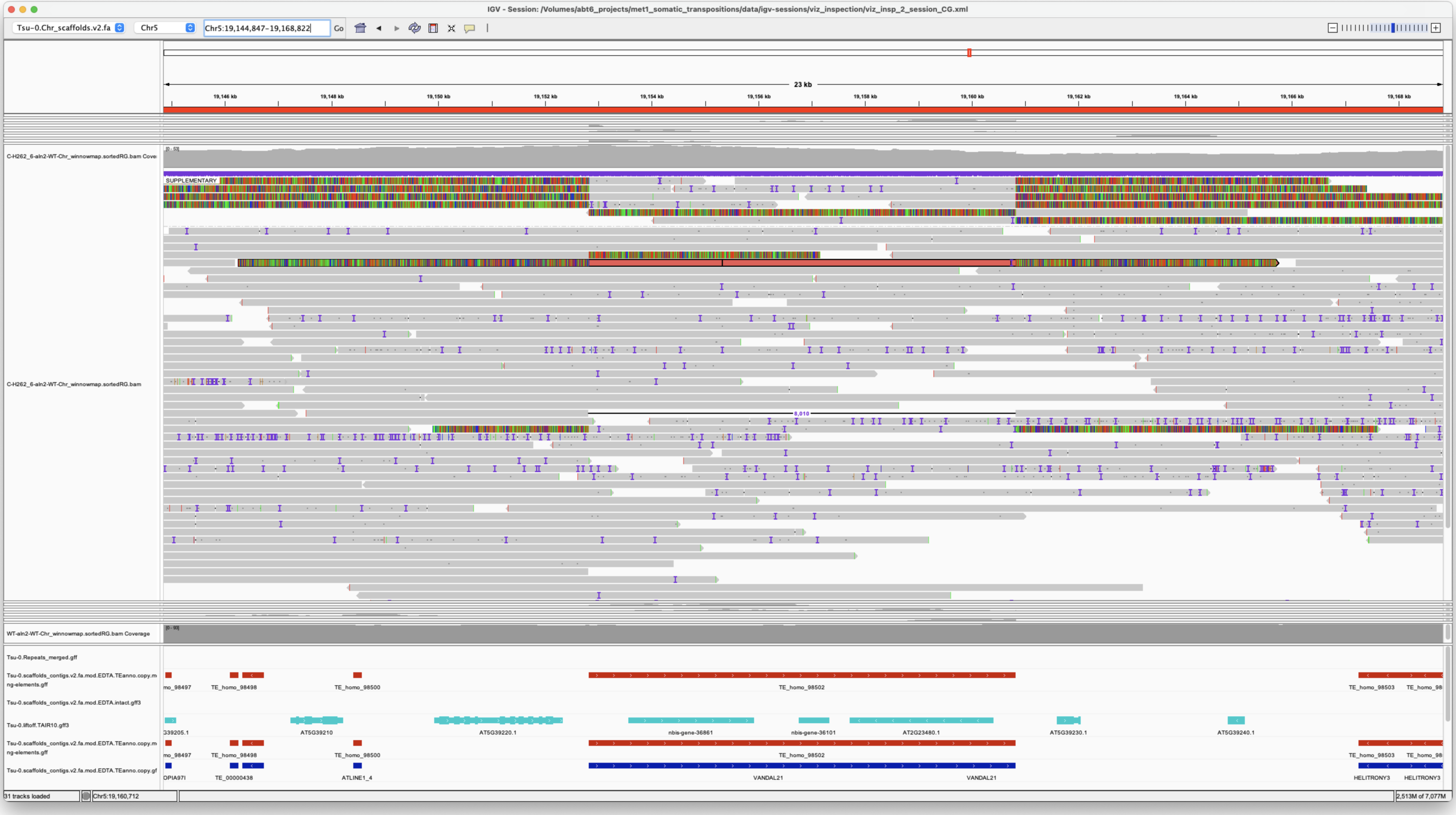


*Confirmed*

---

> Chr1 31010416 31010416 m64079_221220_112036/97584208/ccs Chr5 875412 876434
> Chr5|875414|876433|ID=TE_MANUAL_02;Name=PutativePackTypeCACTAMuDR;classification=DNA/DTC;sequence_ontology=MANUAL;identity=MANUAL;method=MANUAL;ID=TE_MANUAL_02;sequence_ontology=MANUAL met1_06

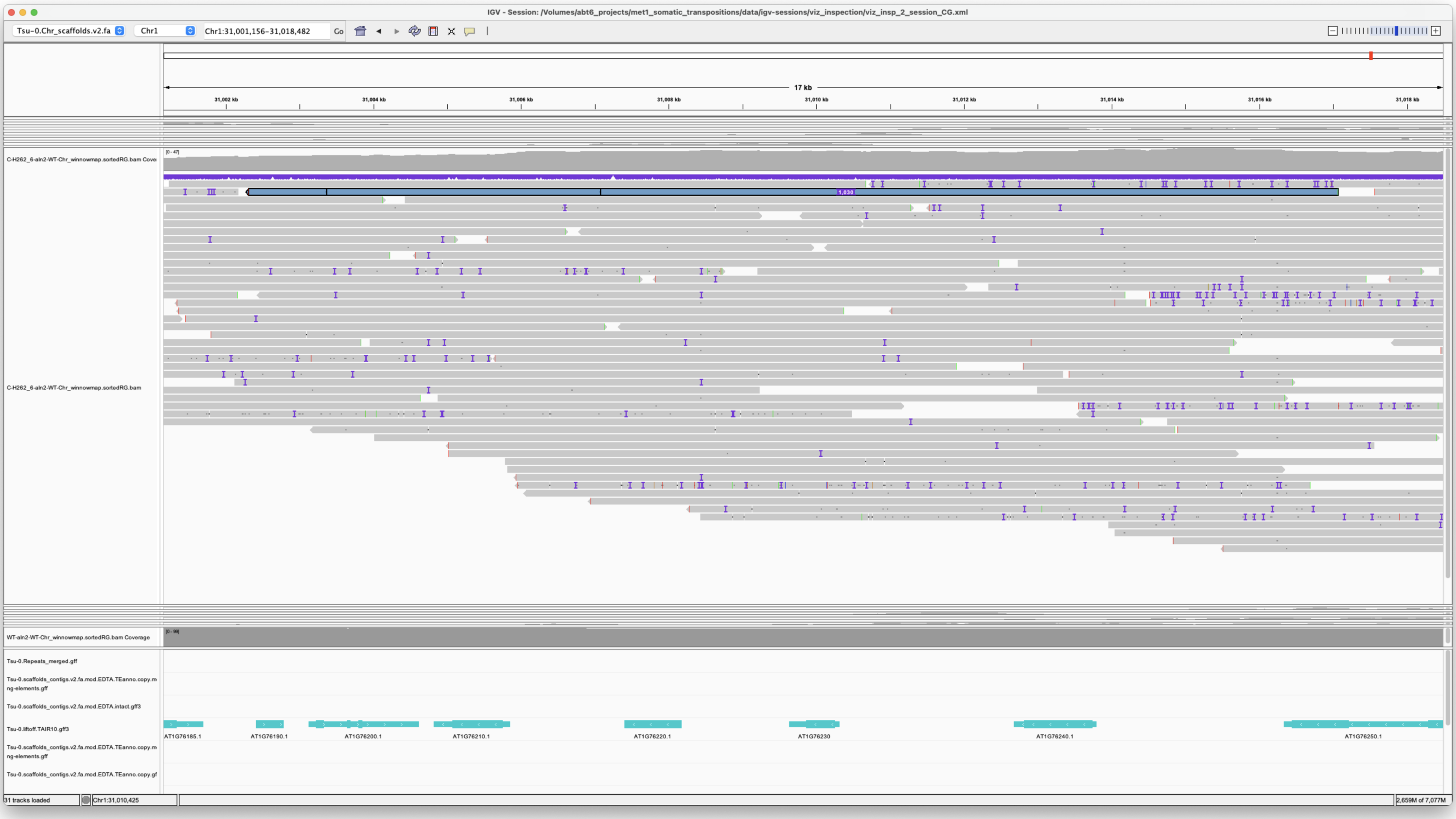

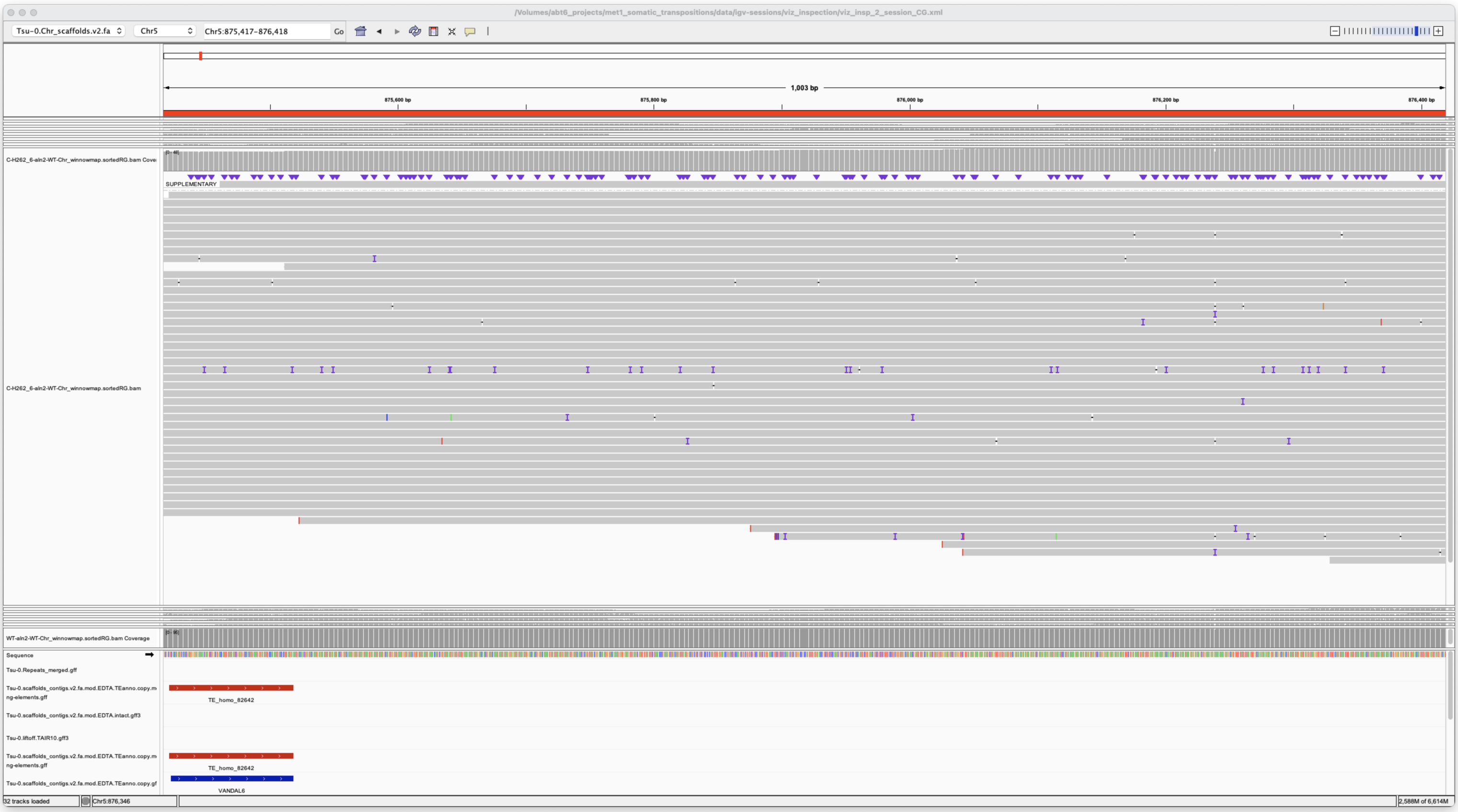

*Confirmed*

> Chr2 2187852 2187852 m64079_221220_112036/39389477/ccs Chr1 11941106 11946435
> Chr1|11941106|11946435|ID=LTRRT_5;Name=ATCOPIA93.2_Evade;Classification=LTR/Copia;Sequence_ontology=SO:0002264;ltr_identity=1.0000;Method=structural;motif=TACA;tsd=ATATG met1_06

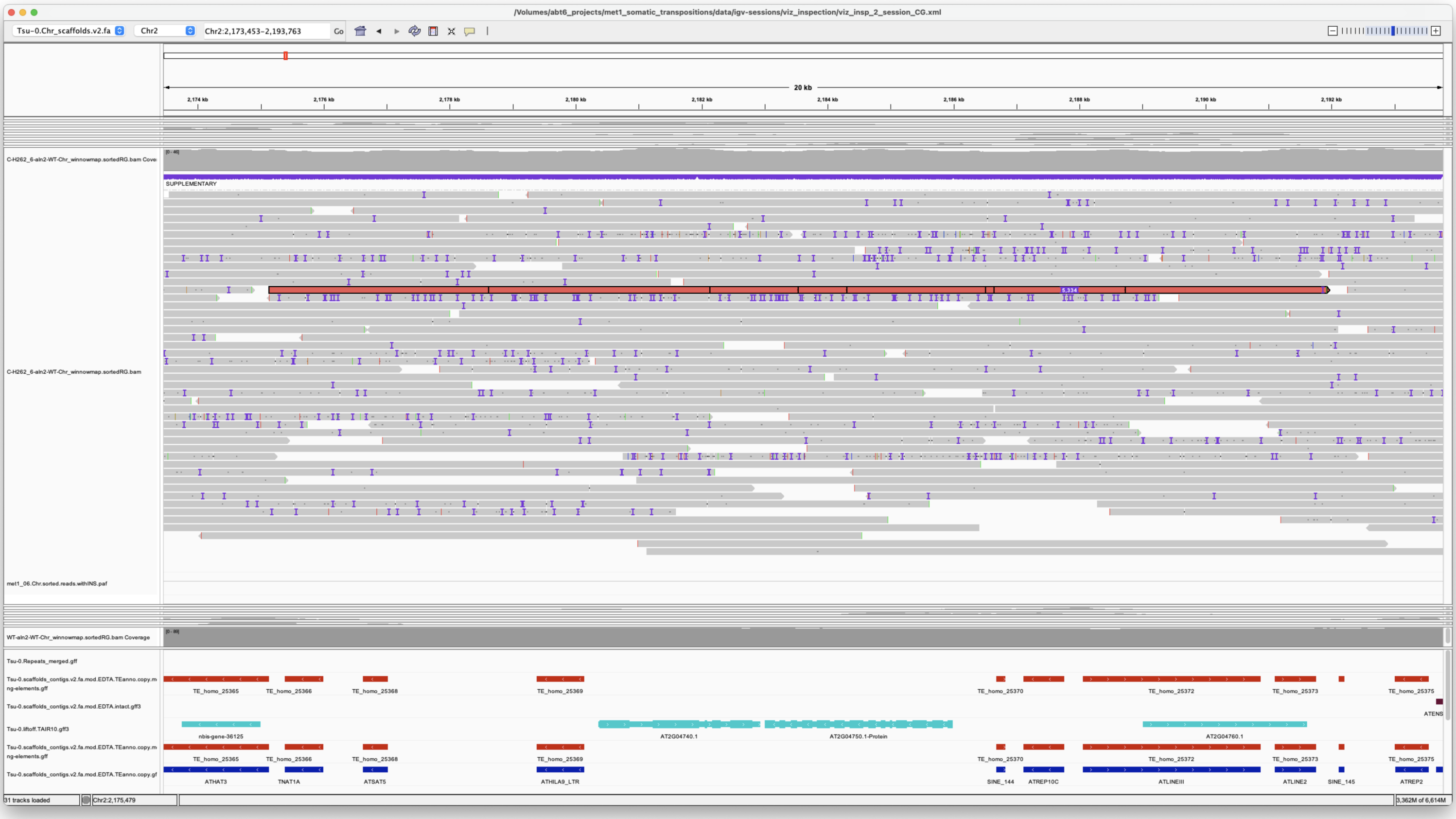


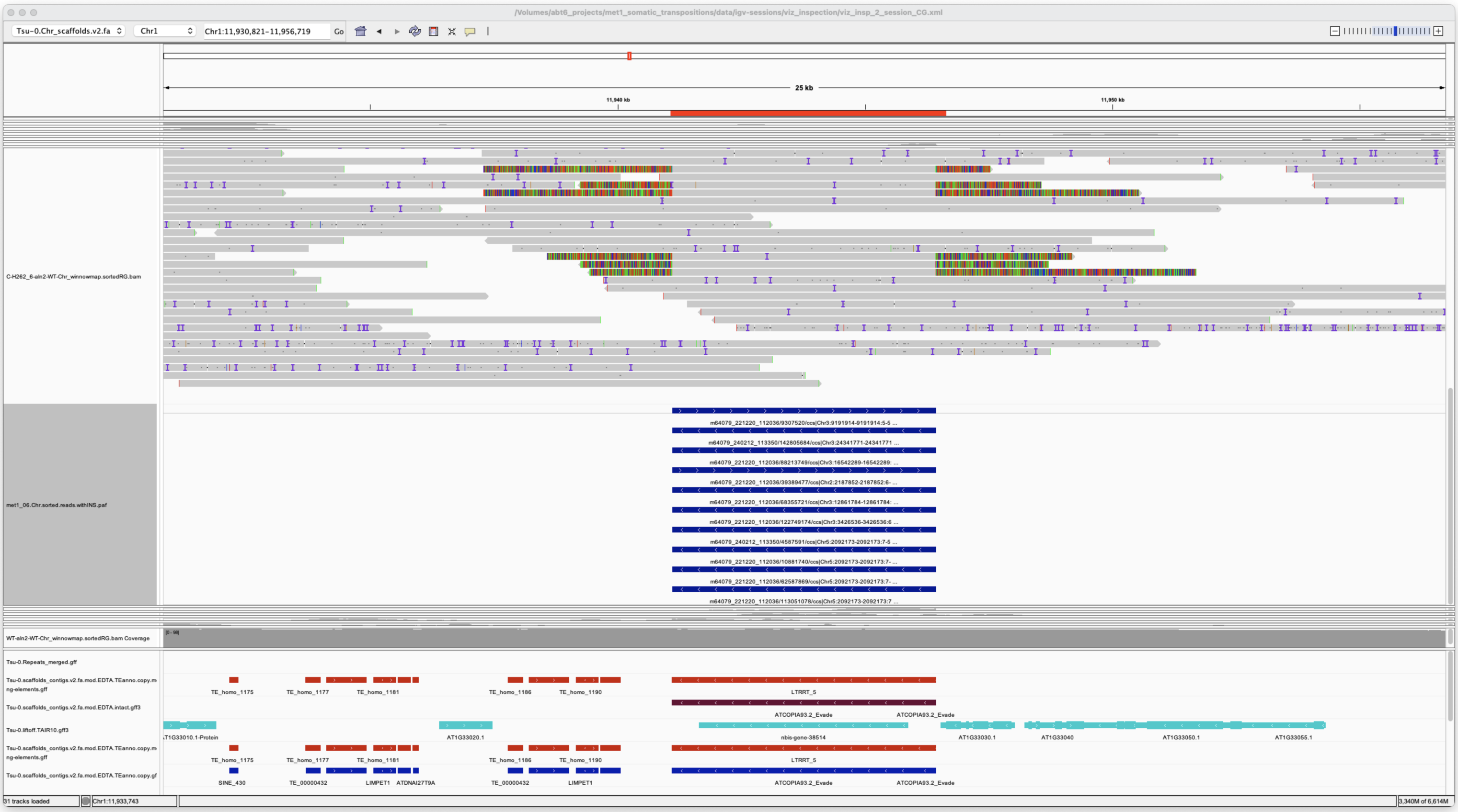


*Confirmed*

---

> Chr3 3426536 3426536 m64079_221220_112036/122749174/ccs Chr1 11941107 11946436
> Chr1|11941106|11946435|ID=LTRRT_5;Name=ATCOPIA93.2_Evade;Classification=LTR/Copia;Sequence_ontology=SO:0002264;ltr_identity=1.0000;Method=structural;motif=TACA;tsd=ATATG met1_06

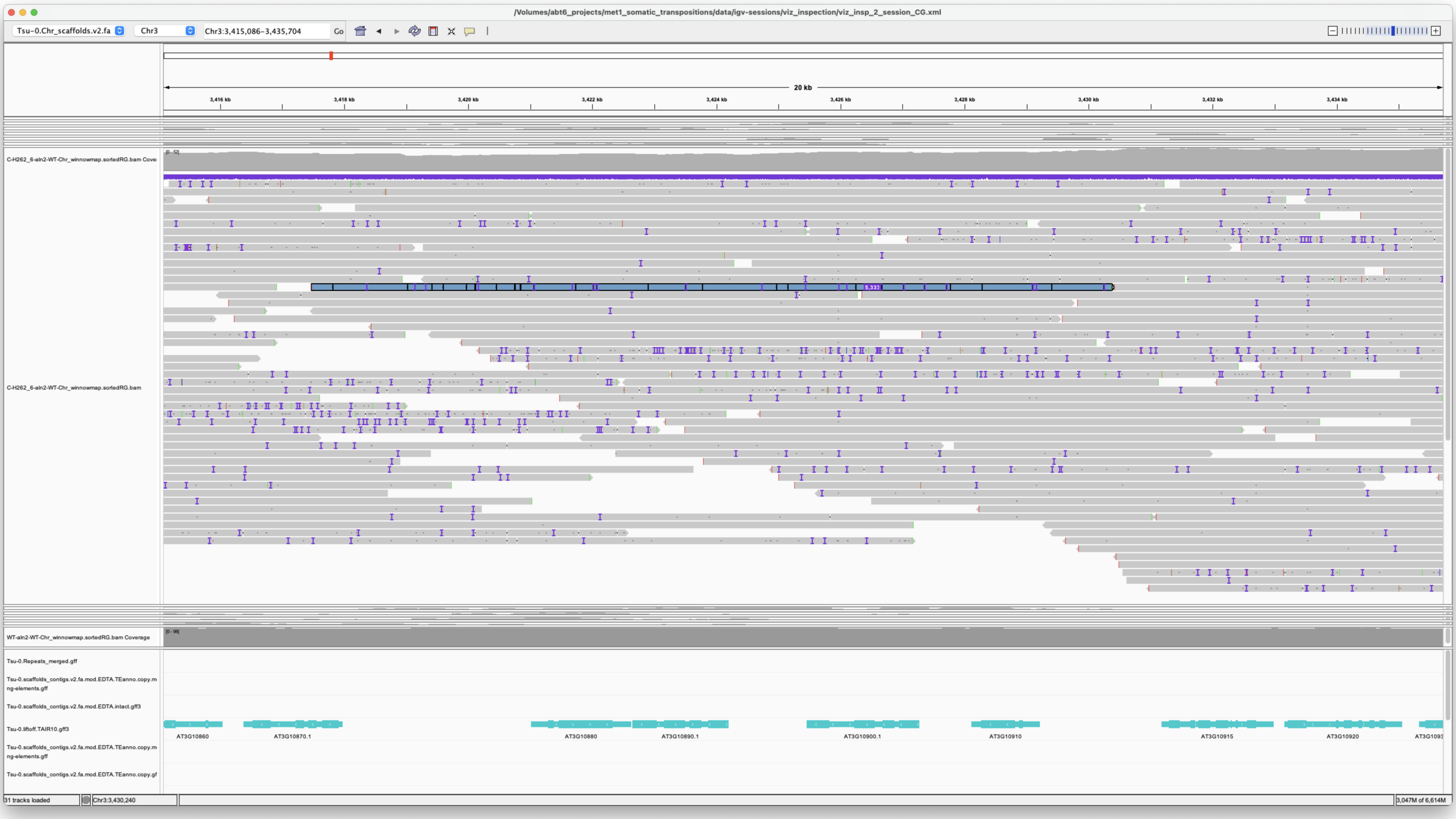


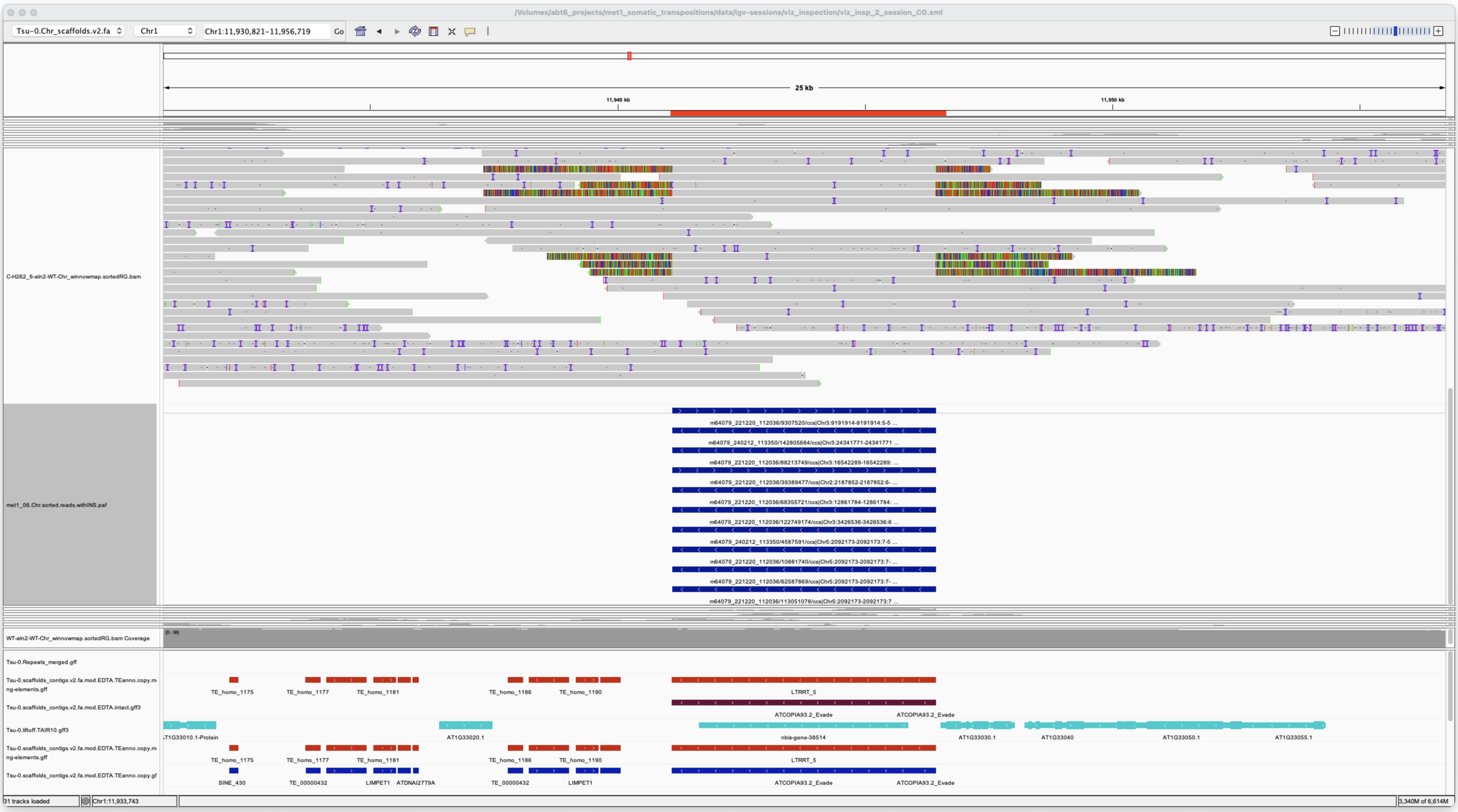


*Confirmed*

Chr3 9191914 9191914 m64079_221220_112036/9307520/ccs Chr1 11941106 11946436 Chr1|11941106|11946435|ID=LTRRT_5;Name=ATCOPIA93.2_Evade;Classification=LTR/Copia;Sequence_ontology=SO:0002264;ltr_identity=1.0000;Method=structural;motif=TACA;tsd=ATATG met1_06

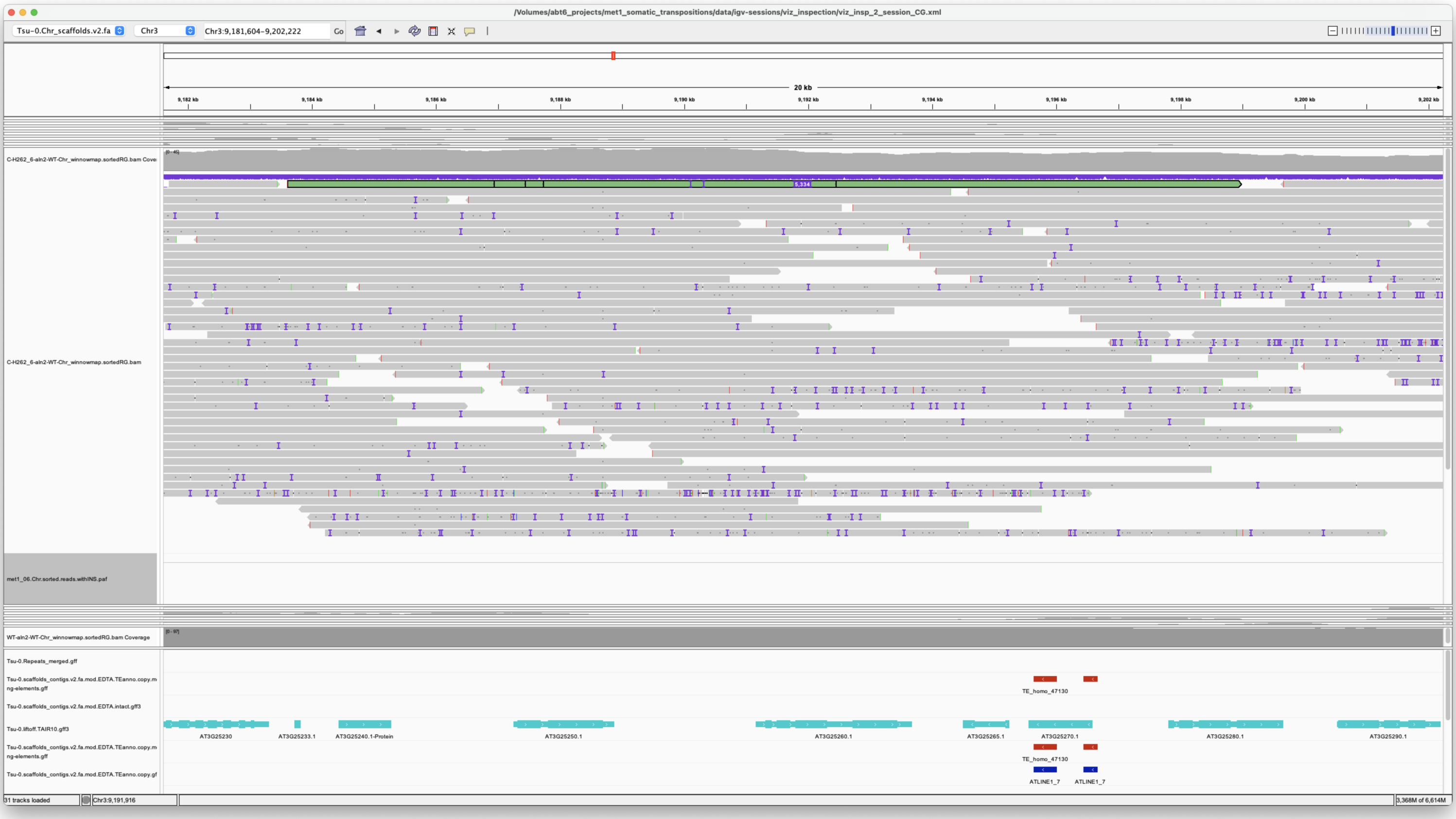


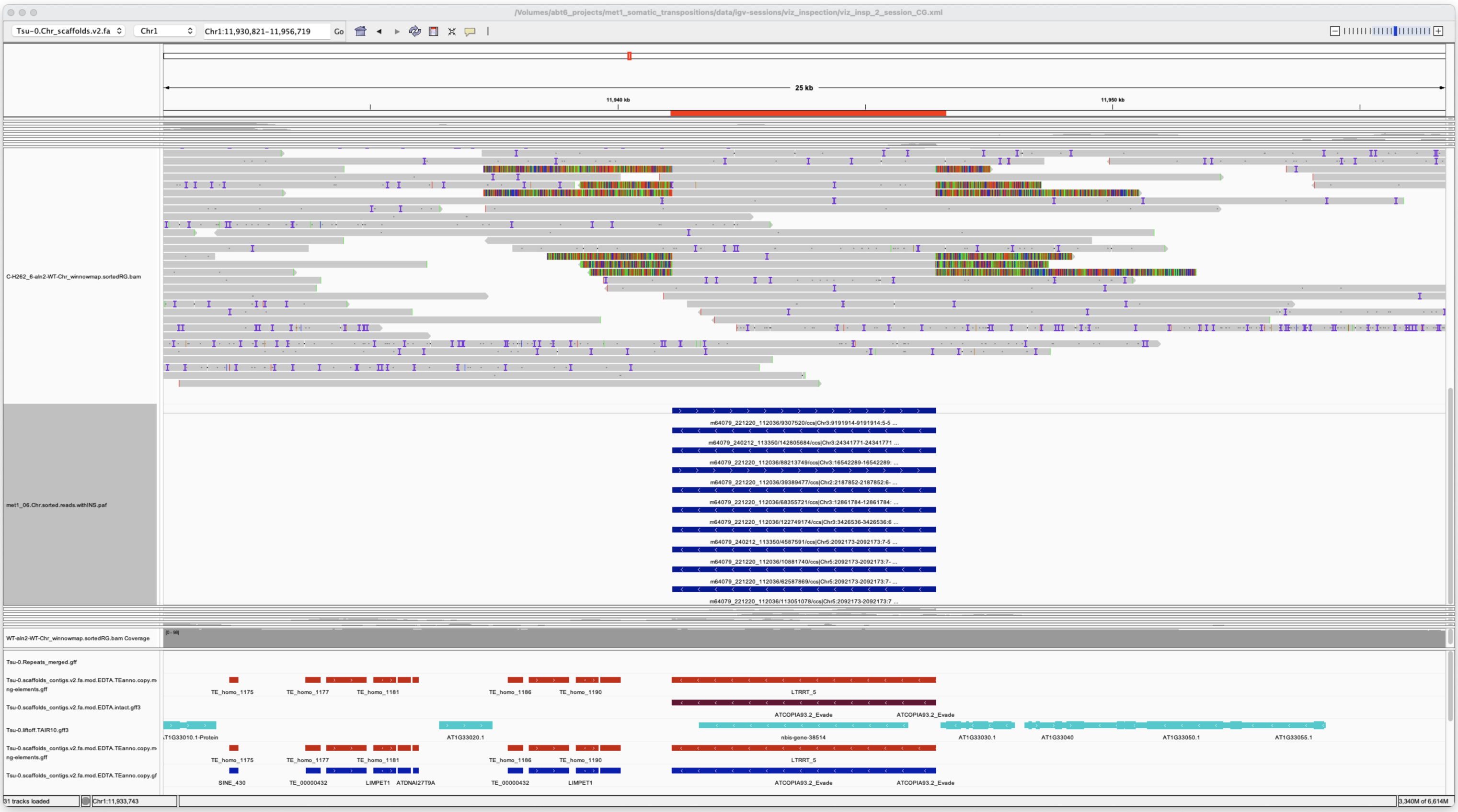


*Confirmed*

Chr3 12861784 12861784 m64079_221220_112036/68355721/ccs Chr1 11941107 11946437
Chr1|11941106|11946435|ID=LTRRT_5;Name=ATCOPIA93.2_Evade;Classification=LTR/Copia;Sequence_ontology=SO:0002264;ltr_identity=1.0000;Method=structural;motif=TACA;tsd=ATATG met1_06

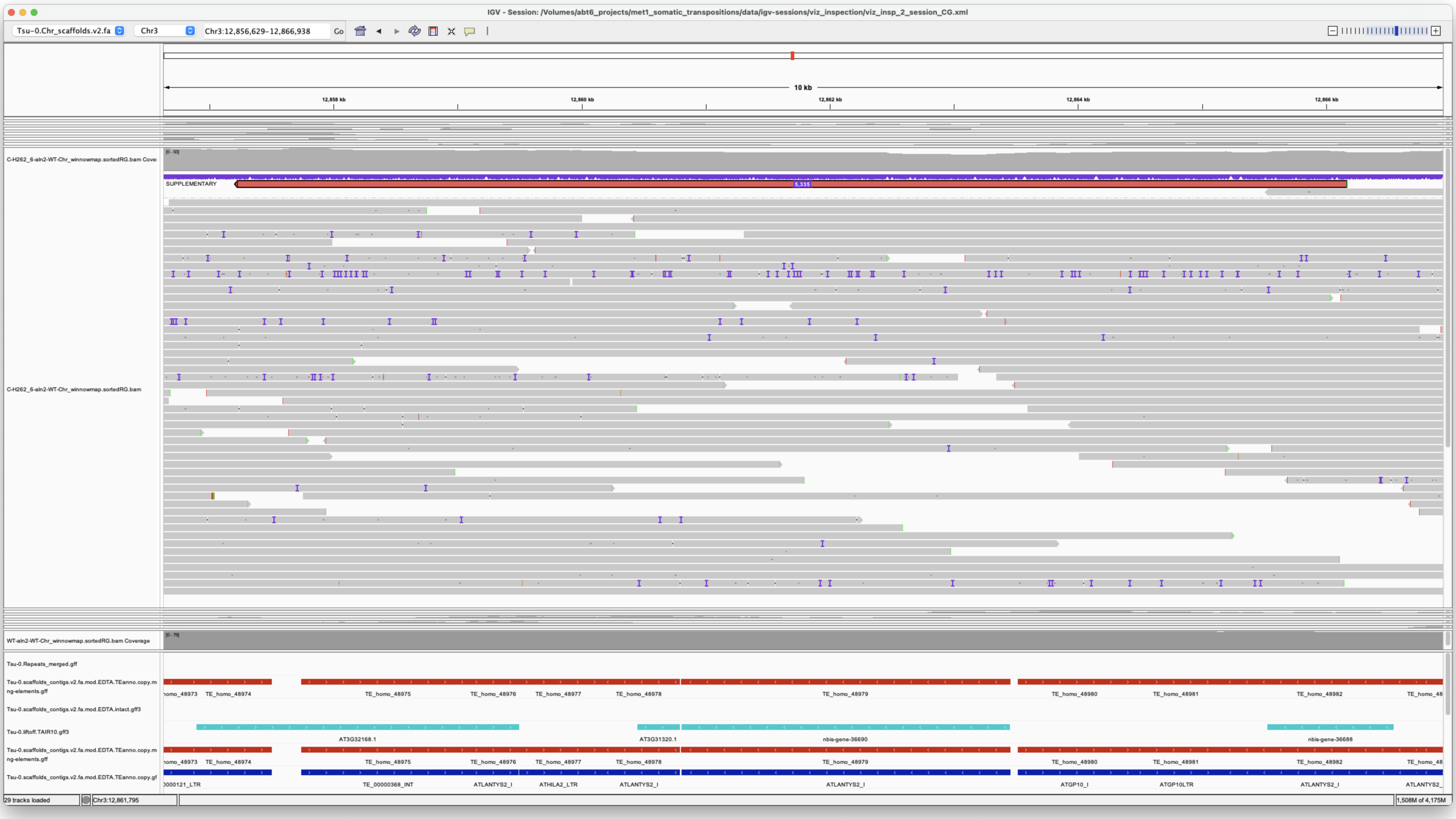


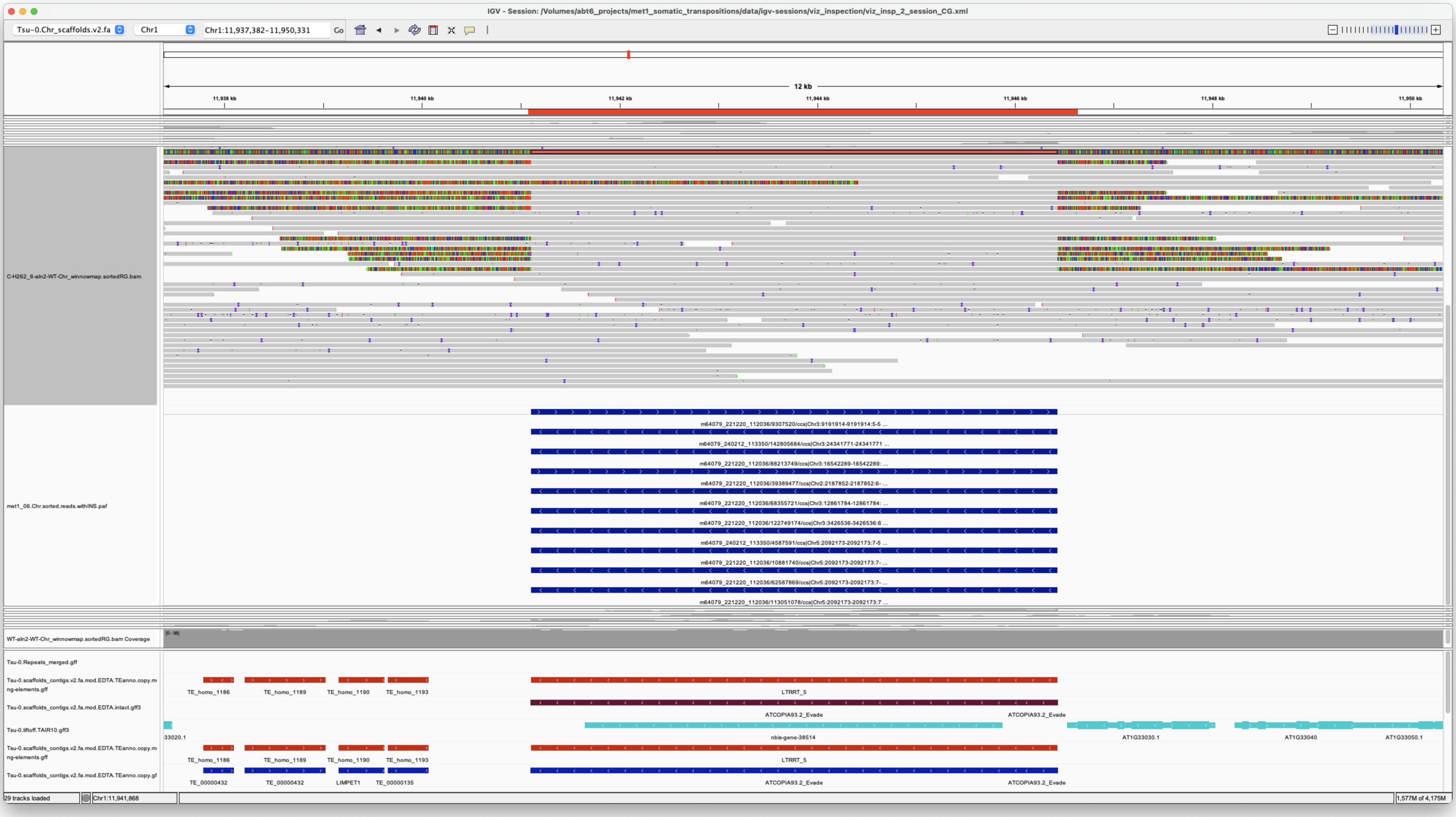


*Confirmed*

---

Chr3 16542289 16542289 m64079_221220_112036/88213749/ccs Chr1 11941106 11946436
Chr1|11941106|11946435|ID=LTRRT_5;Name=ATCOPIA93.2_Evade;Classification=LTR/Copia;Sequence_ontology=SO:0002264;ltr_identity=1.0000;Method=structural;motif=TACA;tsd=ATATG met1_06

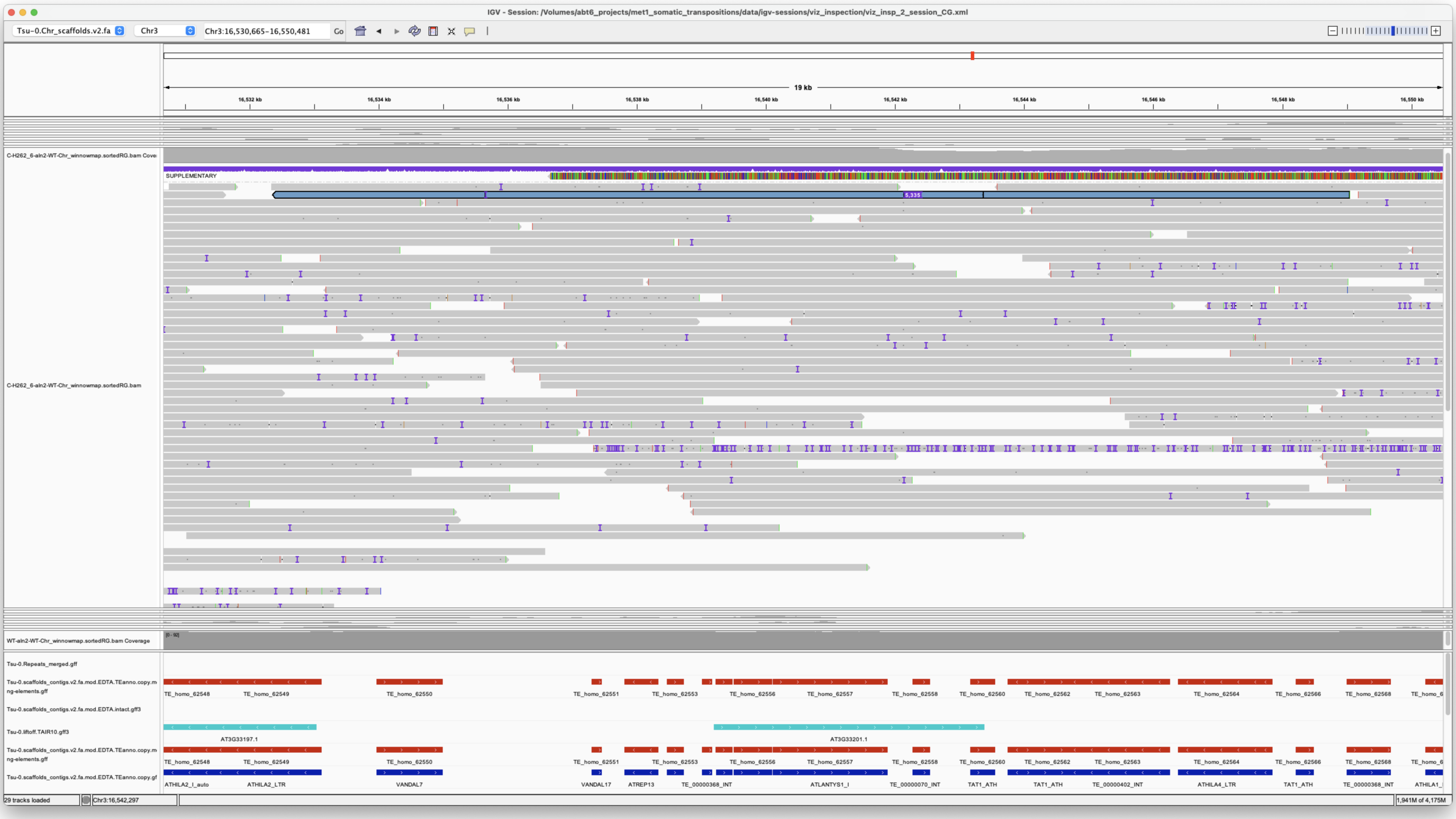


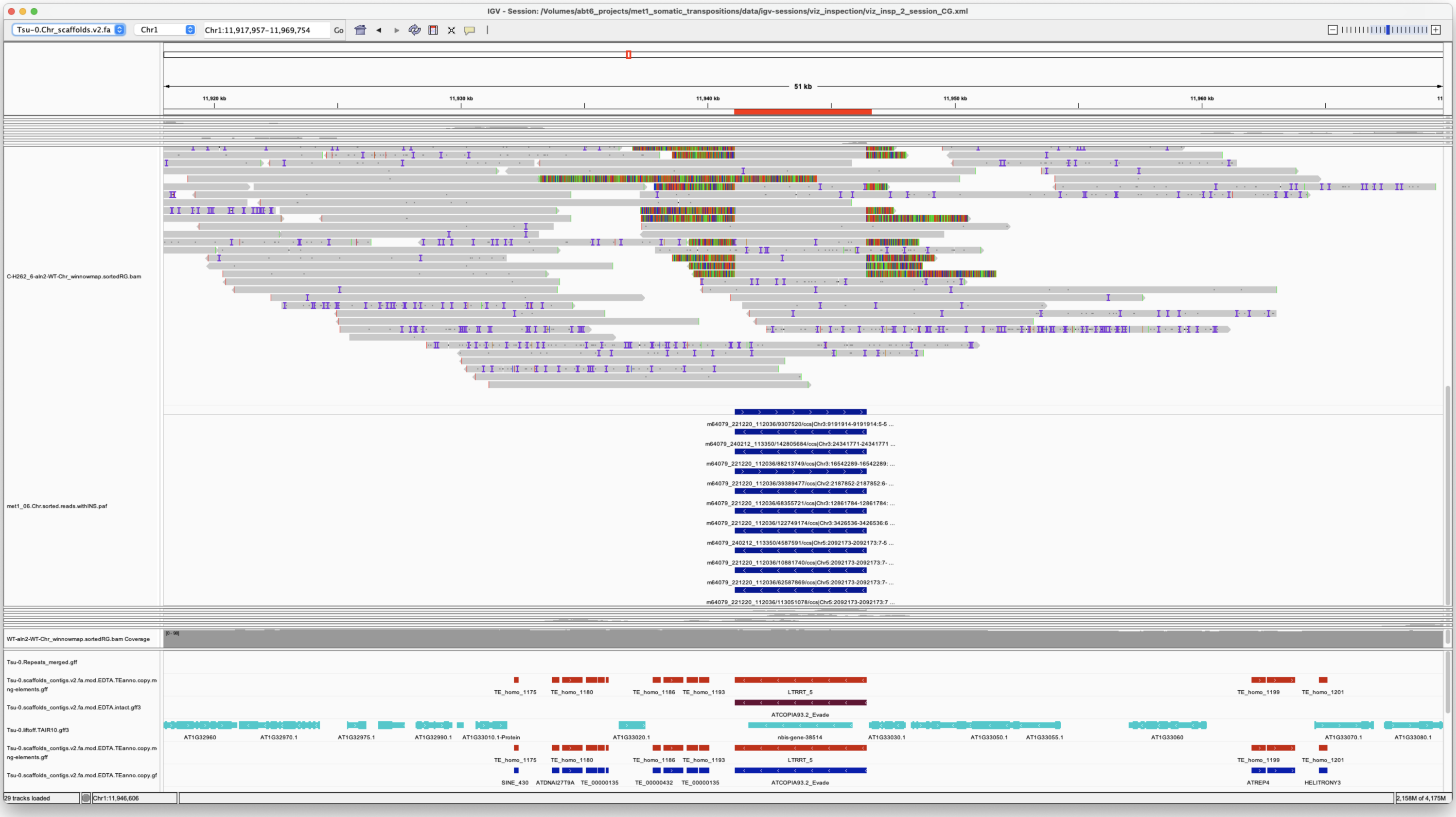


*Confirmed*

---

> Chr3 24341771 24341771 m64079_240212_113350/142805684/ccs Chr1 11941106 11946436
> Chr1|11941106|11946435|ID=LTRRT_5;Name=ATCOPIA93.2_Evade;Classification=LTR/Copia;Sequence_ontology=SO:0002264;ltr_identity=1.0000;Method=structural;motif=TACA;tsd=ATATG met1_06

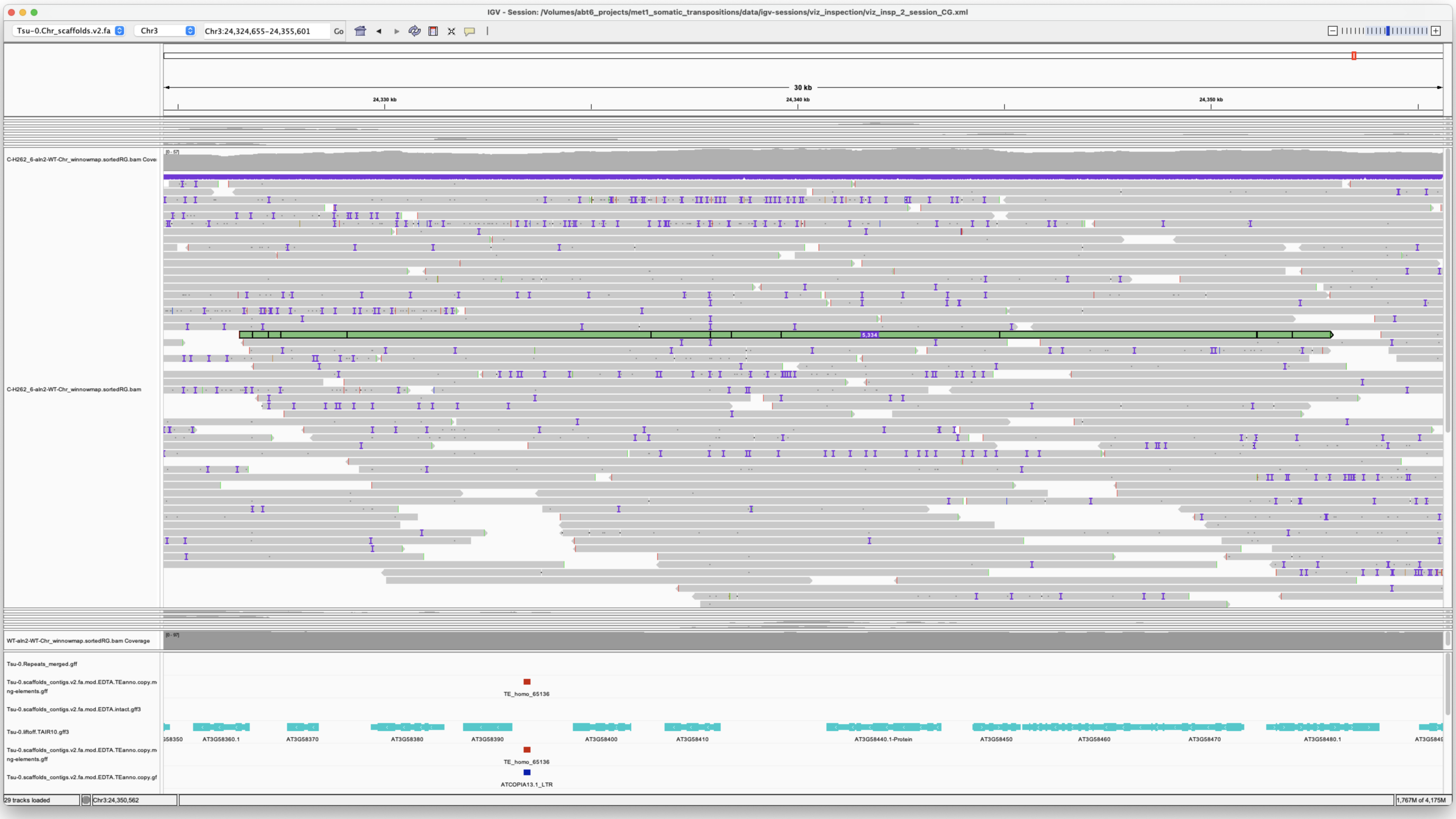

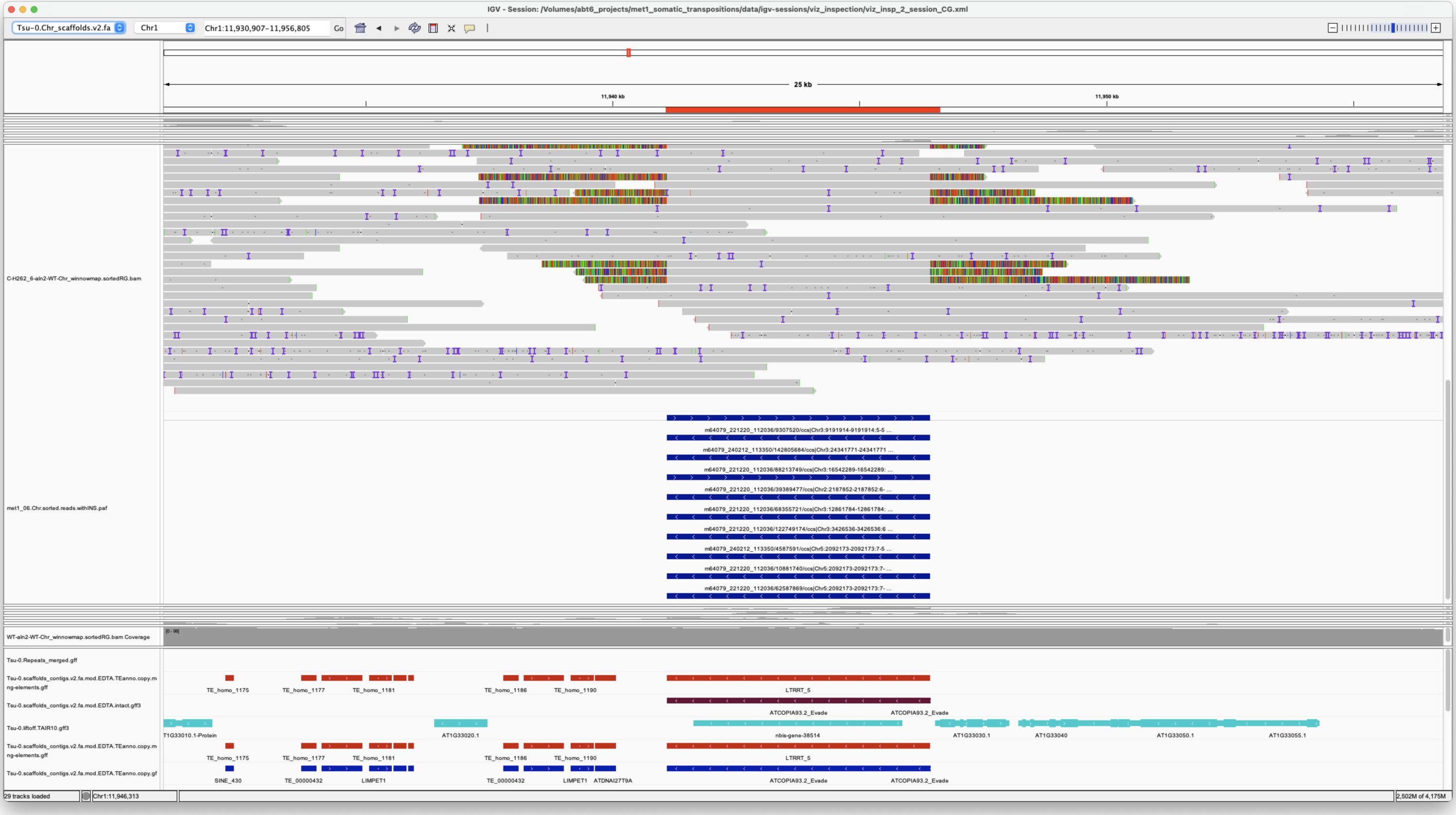

*Confirmed*

Chr4 6070355 6070355 m64079_221220_112036/121309221/ccs Chr4 6067864 6070355 .|-1|-2|. met1_06

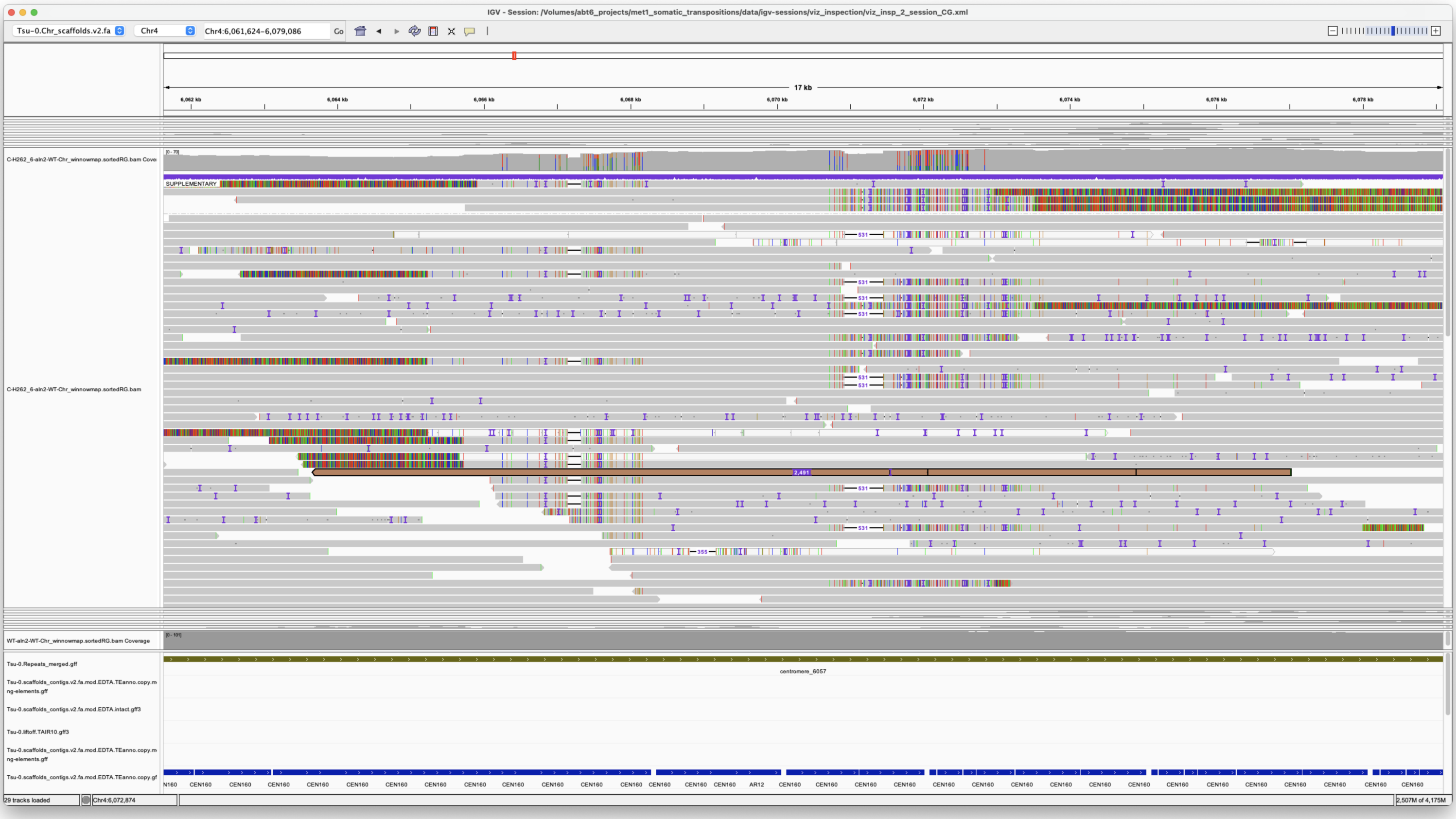

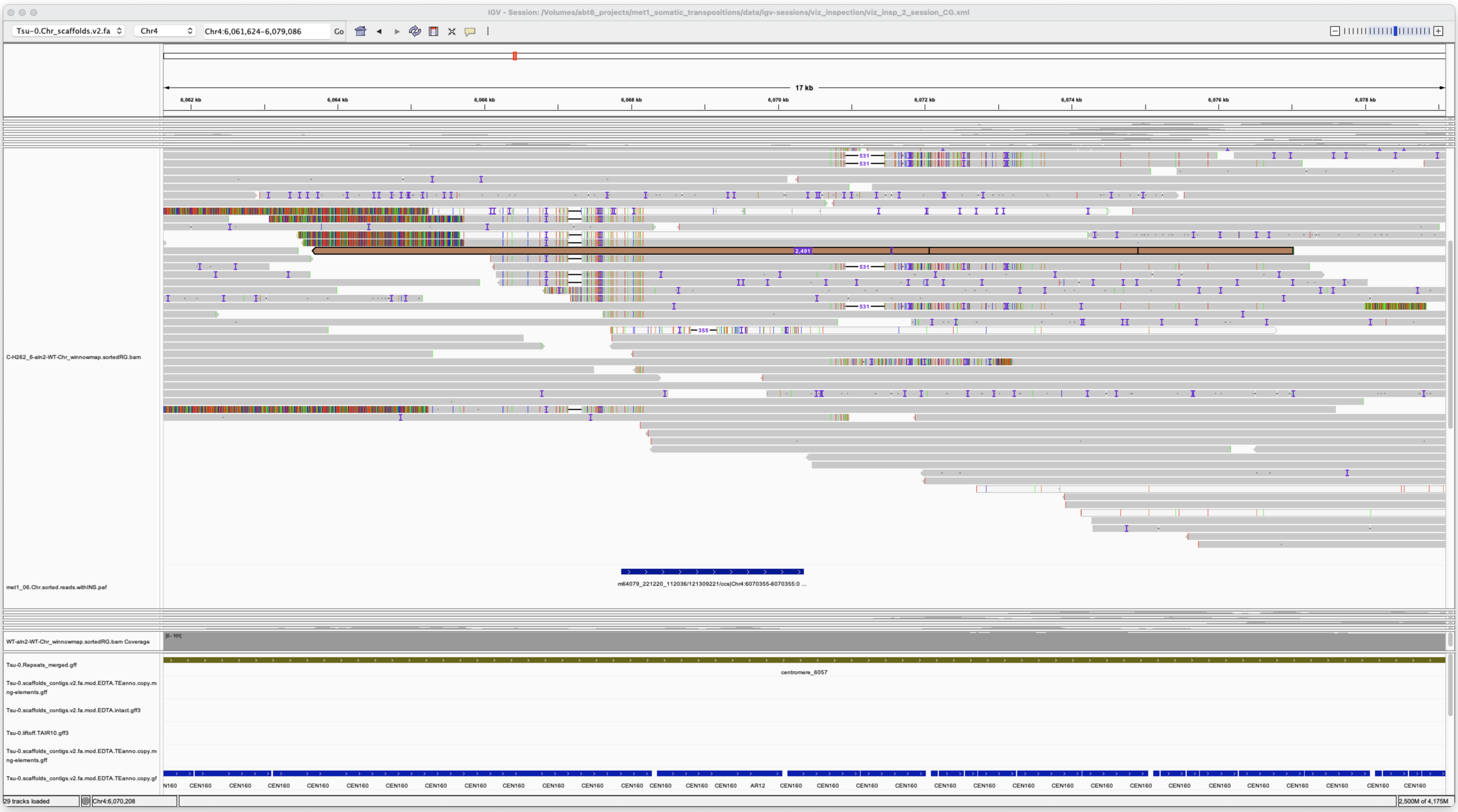

Centromeric rearrangement

*unsupported*

---

> Chr4 12391953 12391953 m64079_240212_113350/99025857/ccs Chr5 875413 876434 Chr5|875414|876433|ID=TE_MANUAL_02;Name=PutativePackTypeCACTAMuDR;classification=DNA/DTC;sequence_ontology=MANUAL;identity=MANUAL;method=MANUAL;ID=TE_MANUAL_02;sequence_ontology=MANUAL met1_06

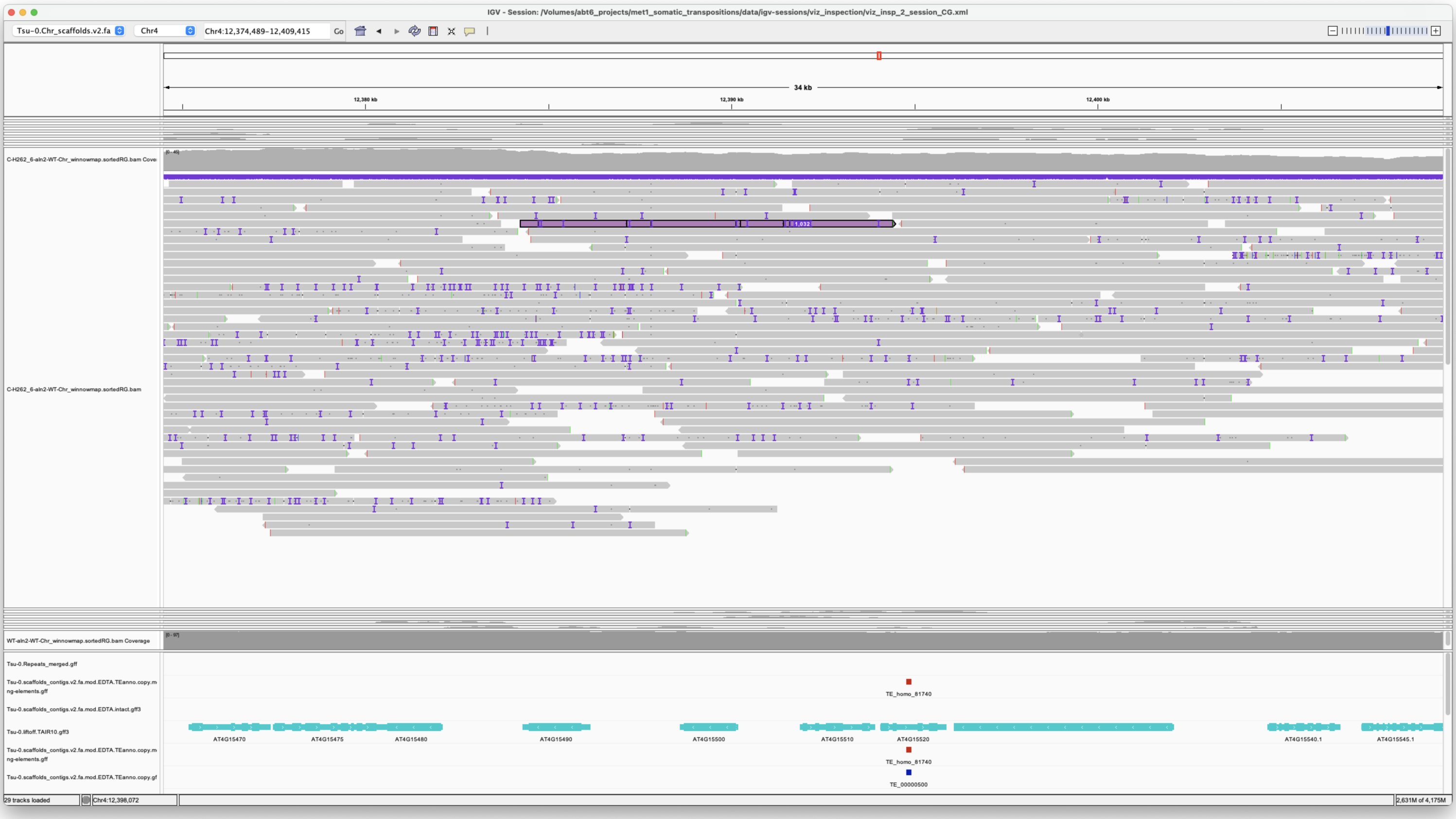


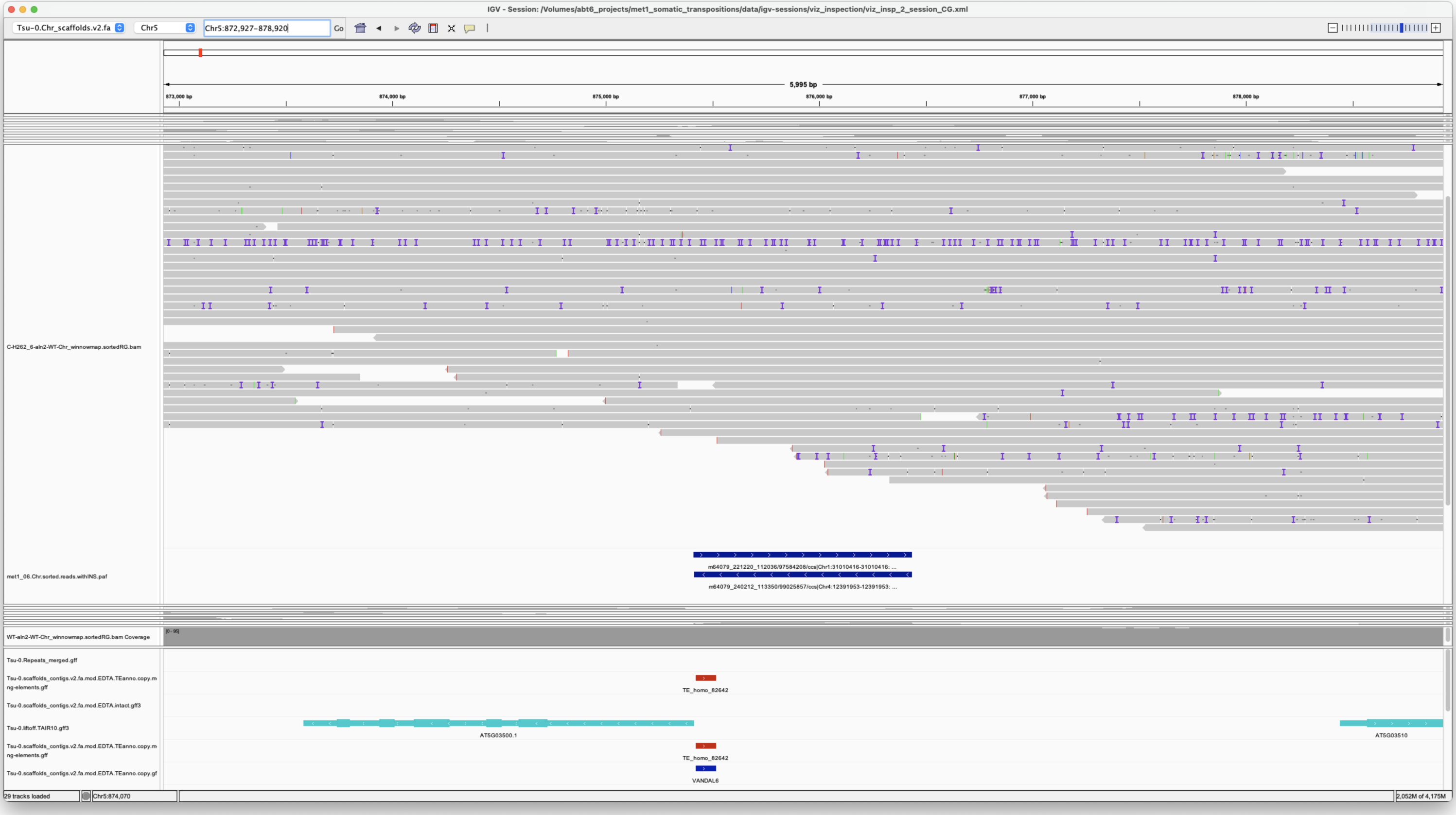


*Confirmed*

---

> Chr5 2092172 2092174 m64079_221220_112036/10881740/ccs**m64079_221220_112036/113051078/ccs**m64079_221220_112036/62587869/ccs**m64079_240212_113350/4587591/ccs Chr1 11941108 11946436 Chr1|11941106|11946435|ID=LTRRT_5;Name=ATCOPIA93.2_Evade;Classification=LTR/Copia;Sequence_ontology=SO:0002264;ltr_identity=1.0000;Method=structural;motif=TACA;tsd=ATATG**Chr1|11941106|11946435|ID=LTRRT_5;Name=ATCOPIA93.2_Evade;Classification=LTR/Copia;Sequence_ontology=SO:0002264;ltr_identity=1.0000;Method=structural;motif=TACA;tsd=ATATG**Chr1|11941106|11946435|ID=LTRRT_5;Name=ATCOPIA93.2_Evade;Classification=LTR/Copia;Sequence_ontology=SO:0002264;ltr_identity=1.0000;Method=structural;motif=TACA;tsd=ATATG**Chr1|11941106|11946435|ID=LTRRT_5;Name=ATCOPIA93.2_Evade;Classification=LTR/Copia;Sequence_ontology=SO:0002264;ltr_identity=1.0000;Method=structural;motif=TACA;tsd=ATATG met1_06

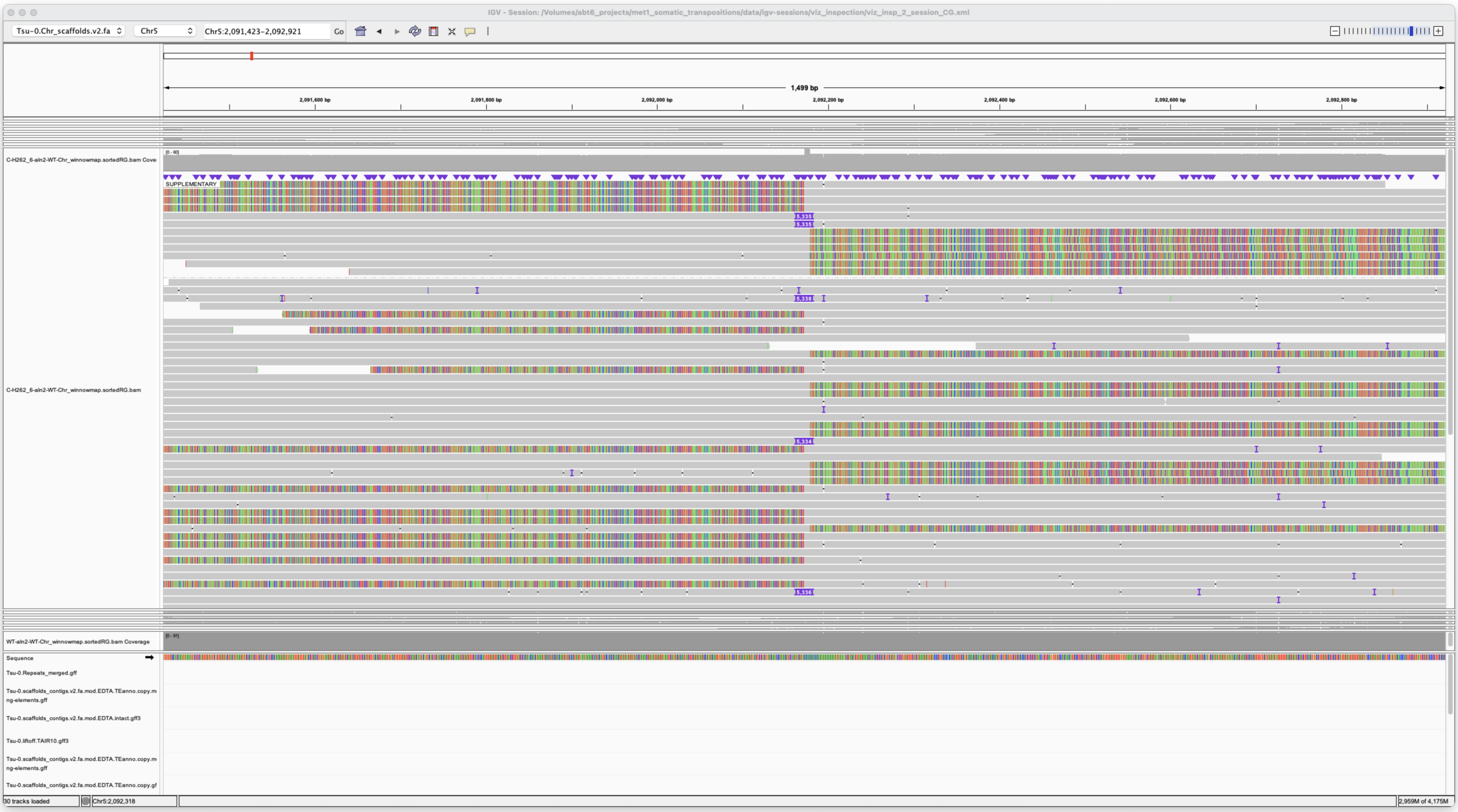


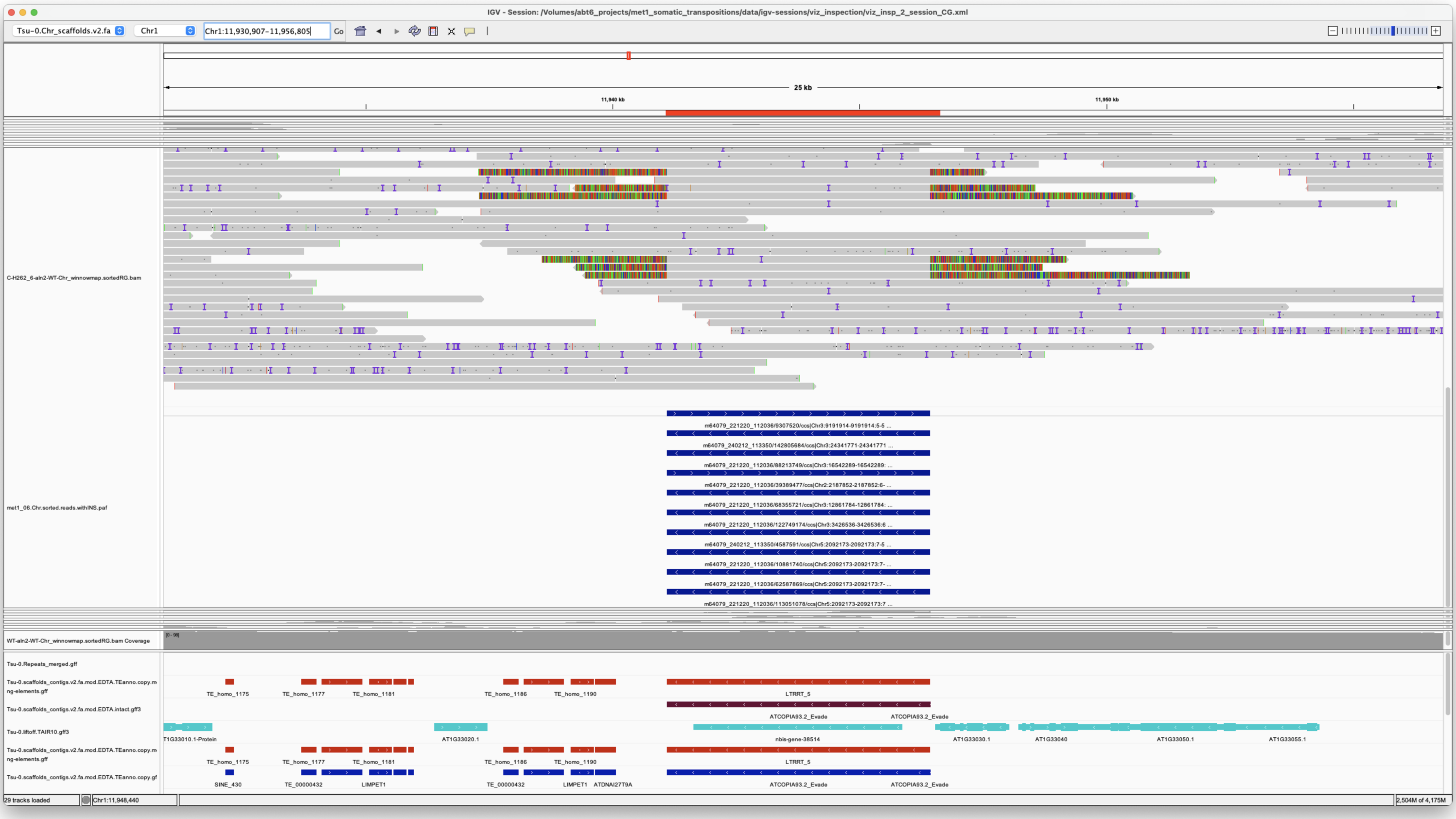


*Confirmed*

Chr5 23005790 23005790 m64079_221220_112036/79759803/ccs Chr5 8256941 8265133
Chr5|8256939|8265132|ID=TE_MANUAL_01;Name=ChimericCACTA;classification=DNA/DTC;sequence_ontology=MANUAL;identity=MANUAL;method=MANUAL;ID=TE_MANUAL_01;sequence_ontology=MANUAL met1_06

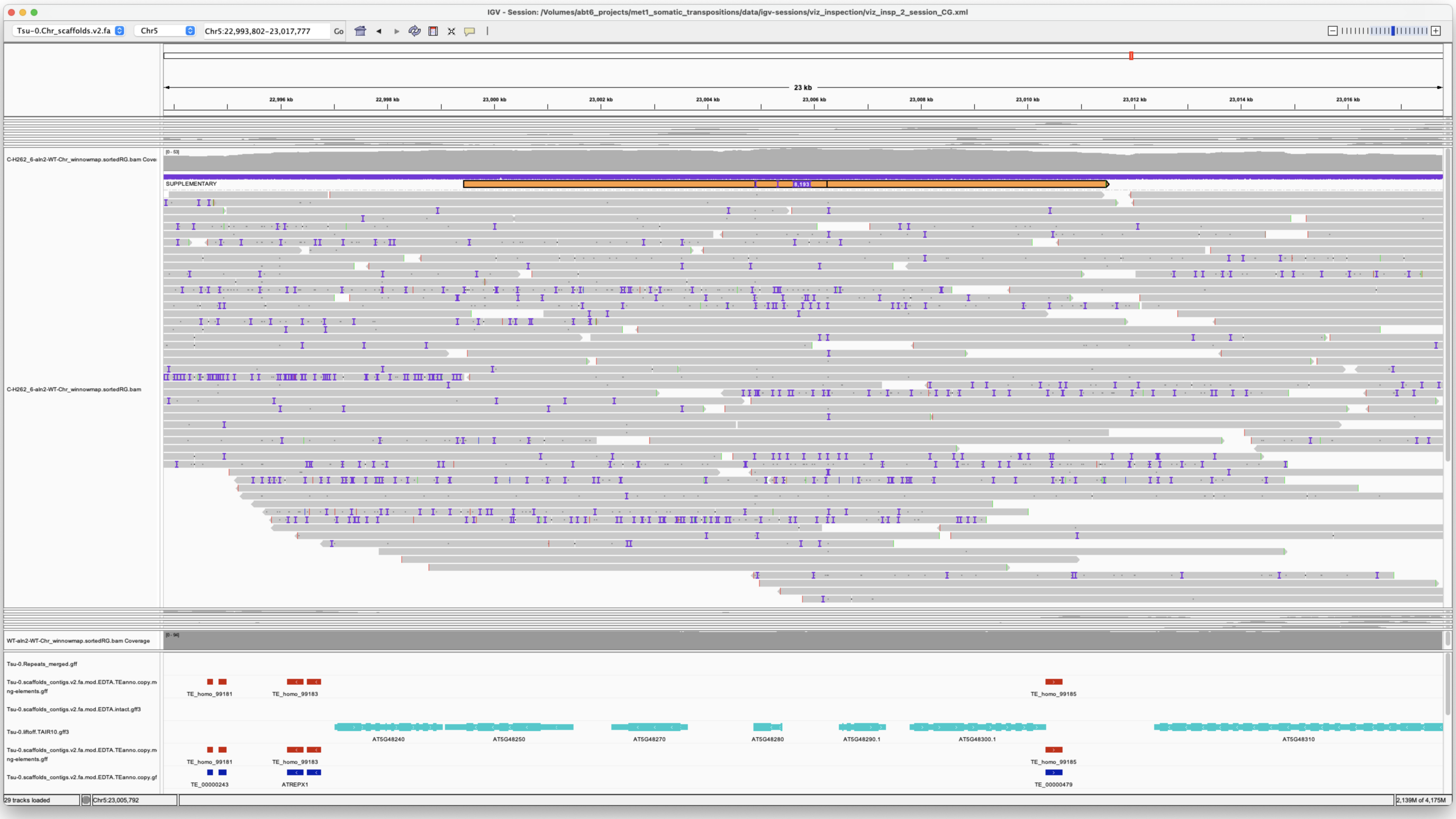


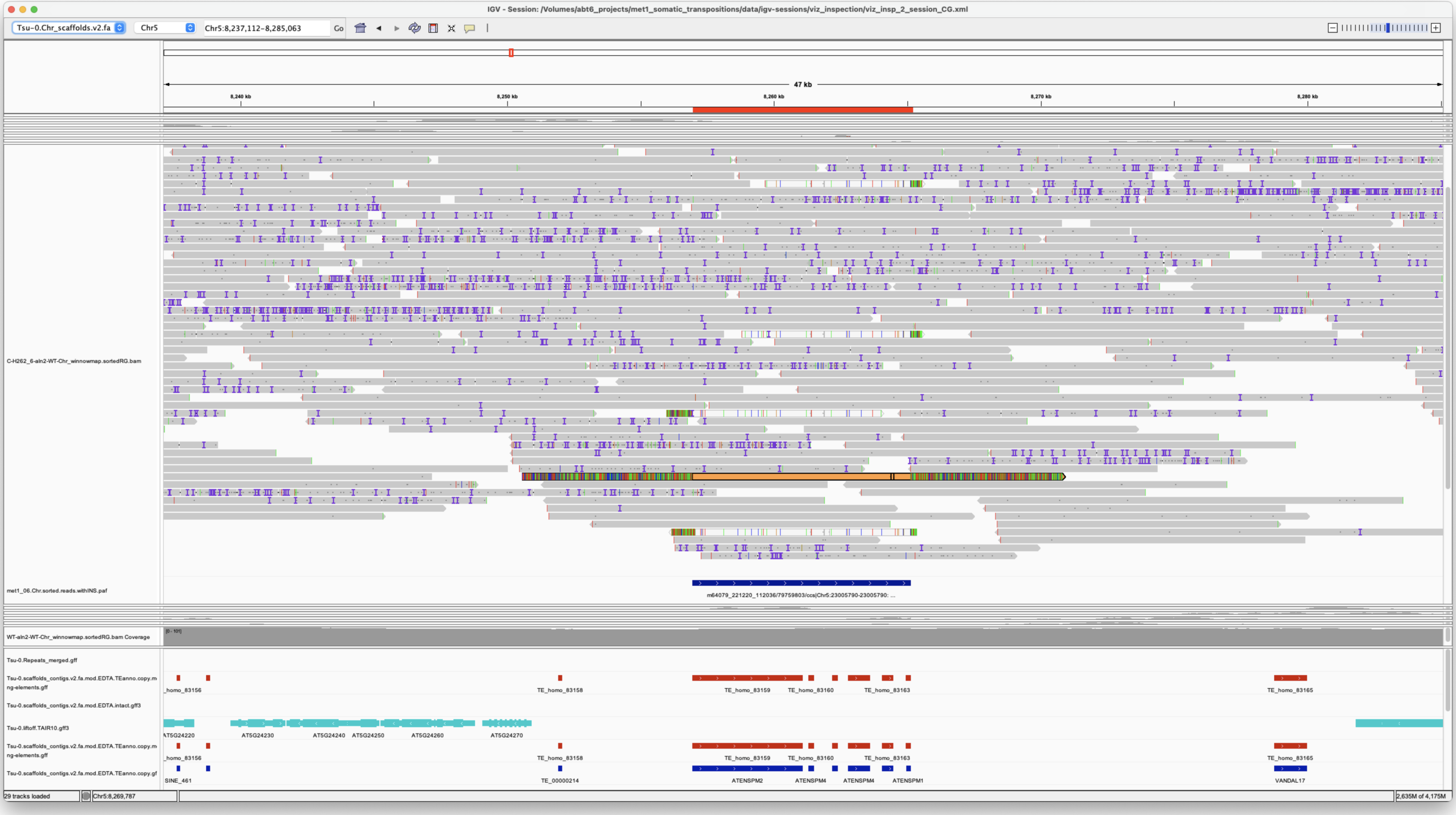


*Confirmed*

#### met1_07

Chr1 276643 276643 m64079_240212_113350/83624893/ccs Chr5 875415 876434 Chr5|875414|876433|ID=TE_MANUAL_02;Name=PutativePackTypeCACTAMuDR;classification=DNA/DTC;sequence_ontology=MANUAL;identity=MANUAL;method=MANUAL;ID=TE_MANUAL_02;sequence_ontology=MANUAL met1_07

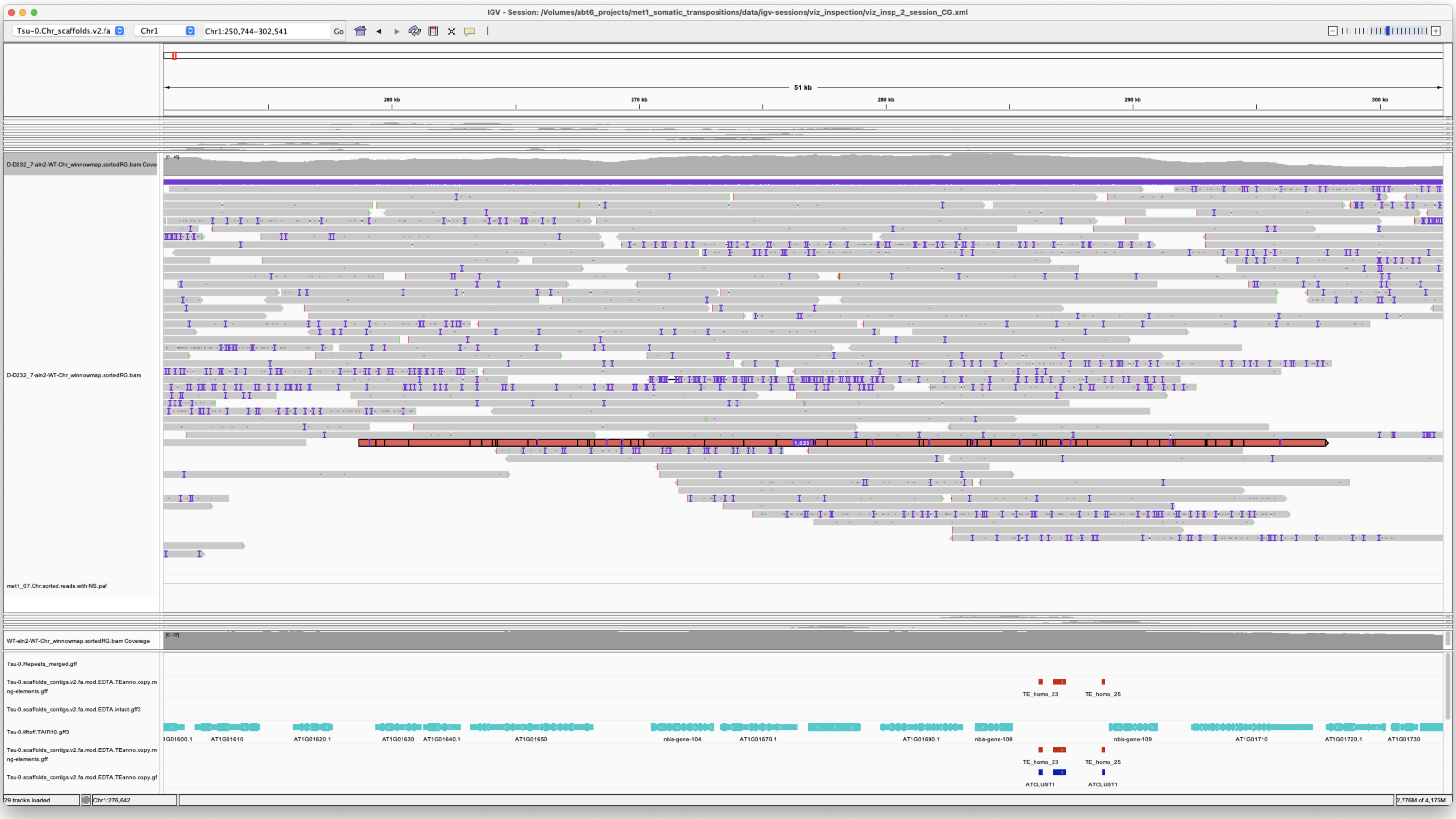


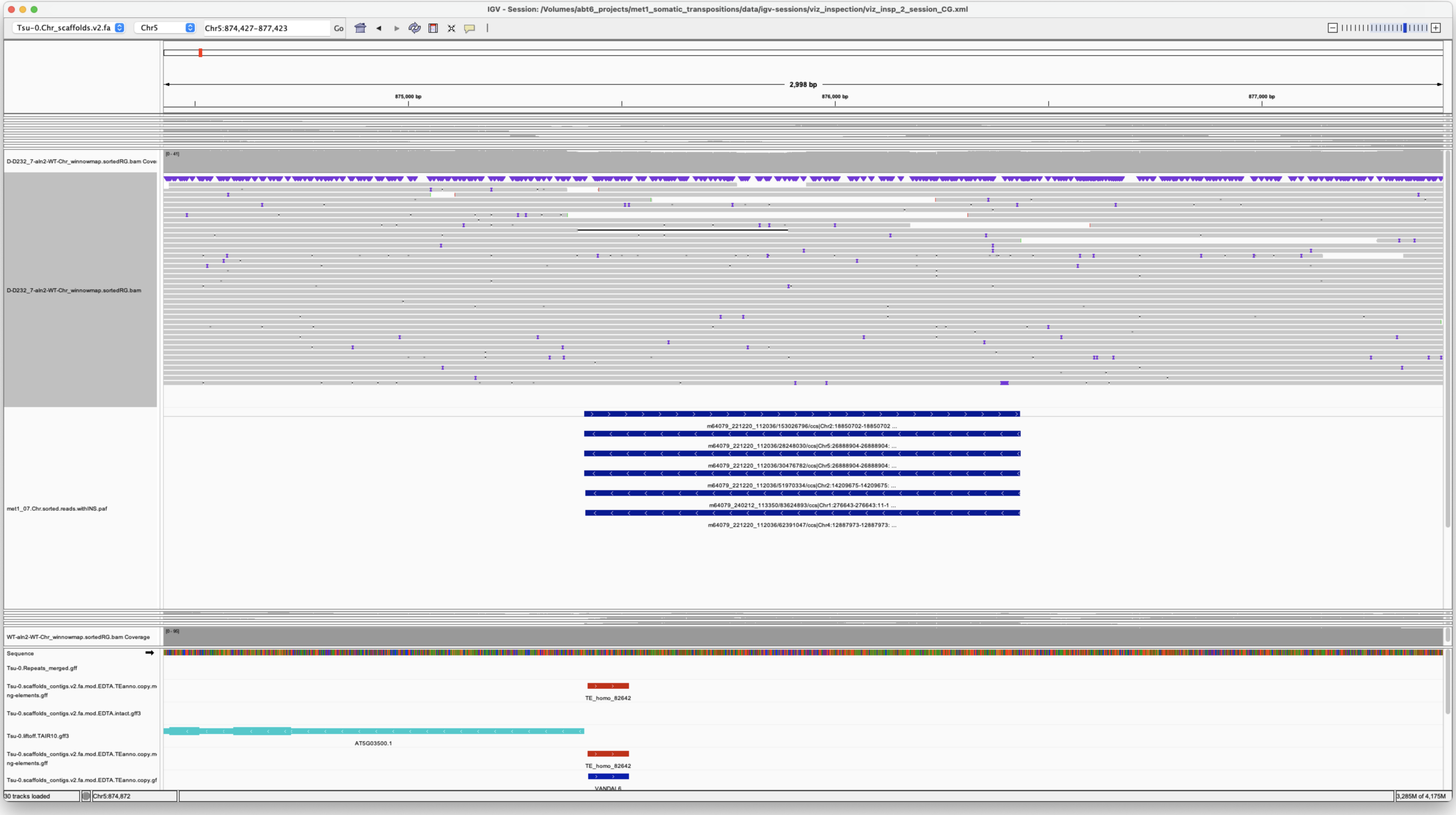


*Confirmed*

> Chr1 8551345 8551345 m64079_221220_112036/173541973/ccs Chr1 11941106 11946436 Chr1|11941106|11946435|ID=LTRRT_5;Name=ATCOPIA93.2_Evade;Classification=LTR/Copia;Sequence_ontology=SO:0002264;ltr_identity=1.0000;Method=structural;motif=TACA;tsd=ATATG met1_07

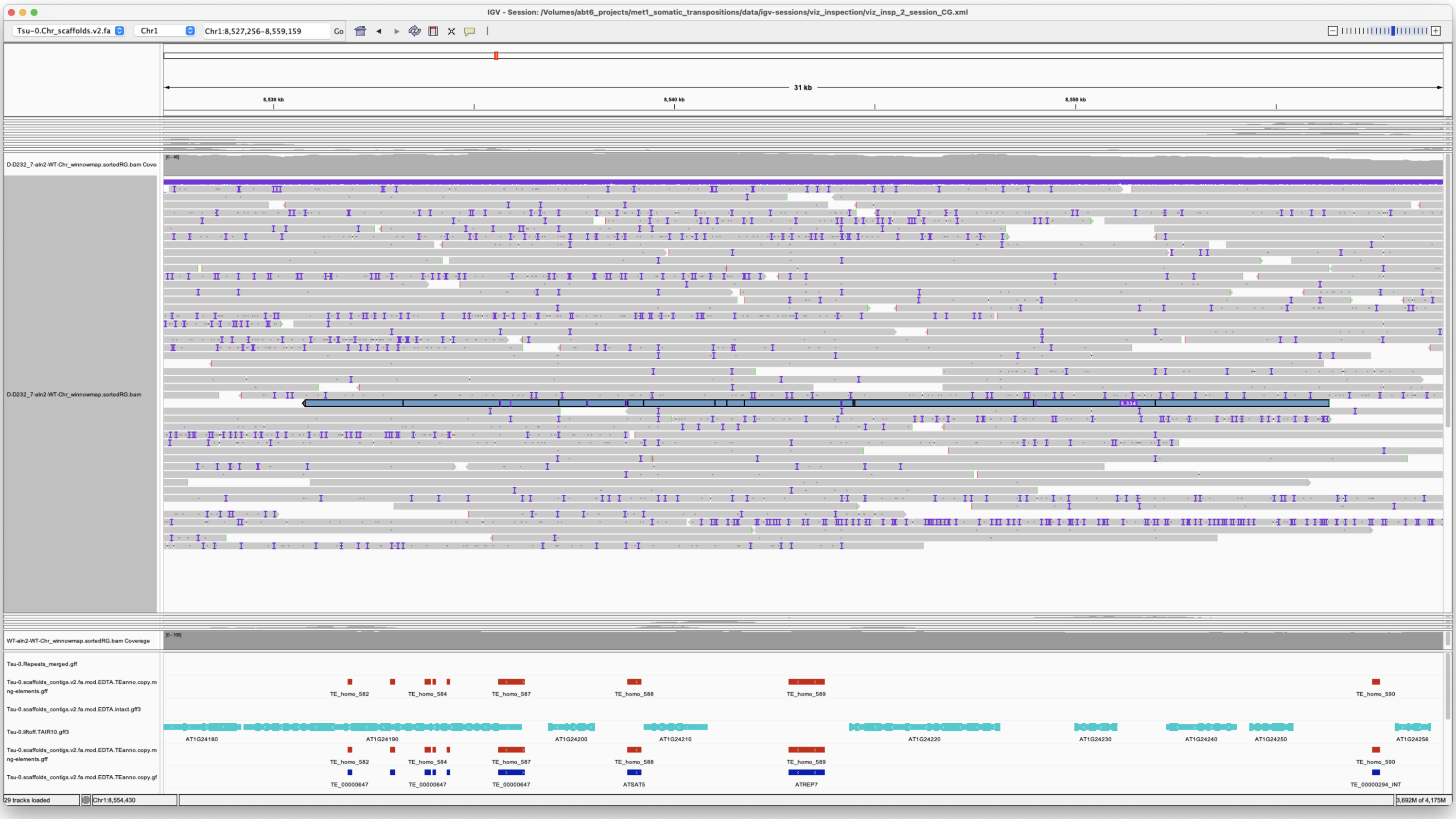


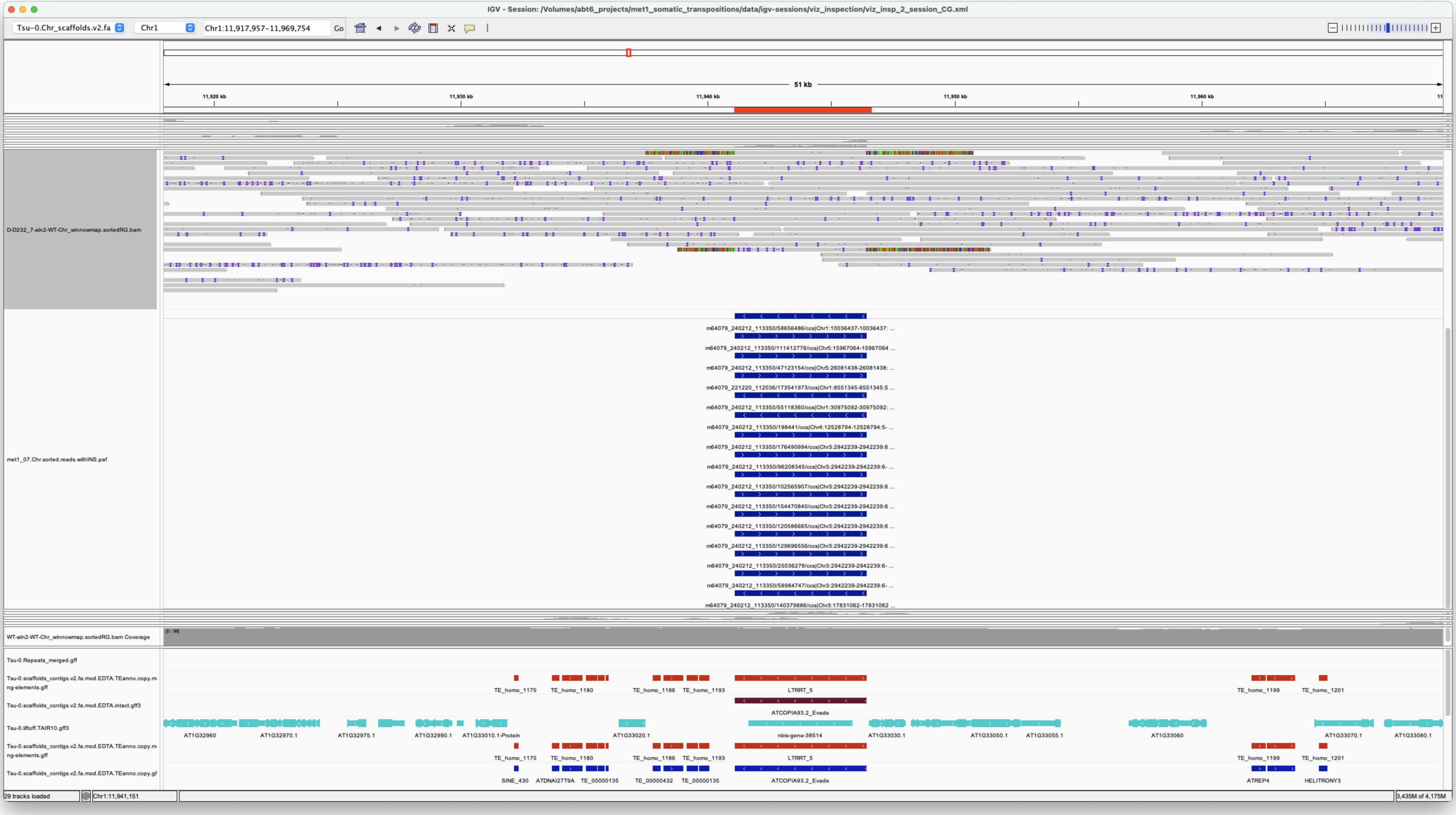


*Confirmed*

> Chr1 10036437 10036437 m64079_240212_113350/58656486/ccs Chr1 11941106 11946437 Chr1|11941106|11946435|ID=LTRRT_5;Name=ATCOPIA93.2_Evade;Classification=LTR/Copia;Sequence_ontology=SO:0002264;ltr_identity=1.0000;Method=structural;motif=TACA;tsd=ATATG met1_07

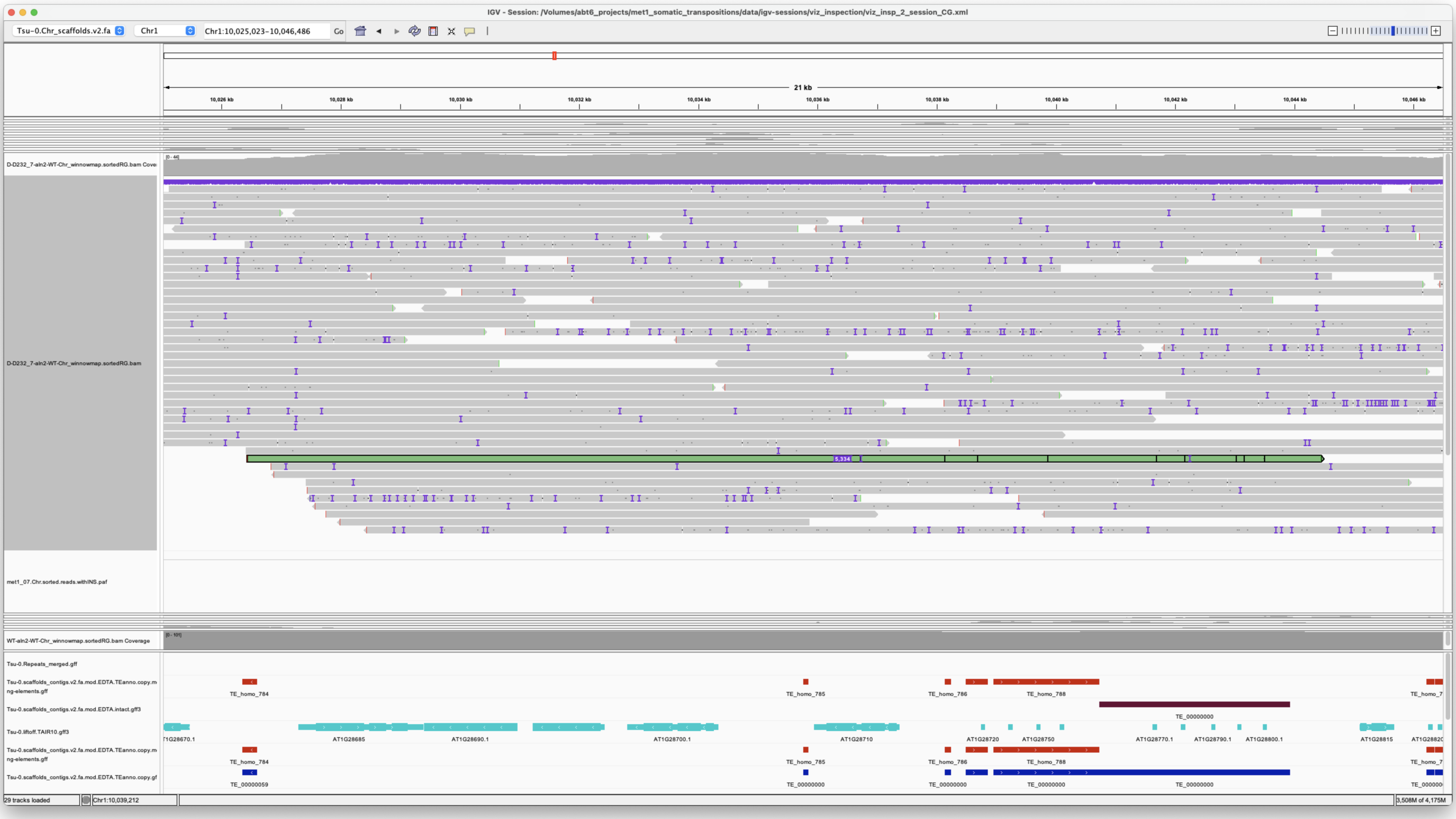


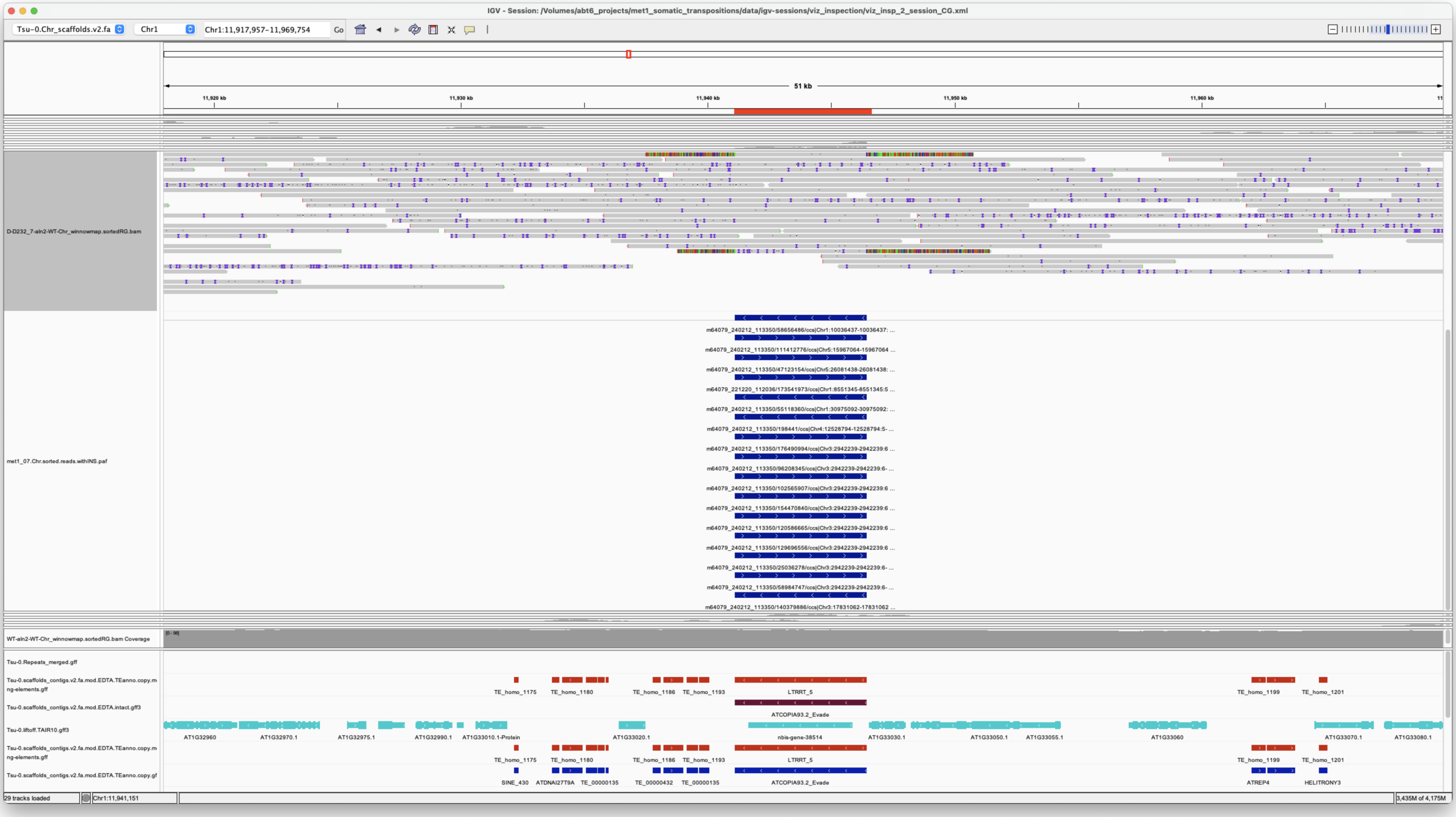


*Confirmed*

---

> Chr1 24744044 24744044 m64079_221220_112036/22151649/ccs Chr3 16344525 16352497
> Chr3|16344522|16352496|ID=TE_homo_60420;Name=VANDAL6;classification=DNA/Mutator;sequence_ontology=SO:0002280;identity=0.969;method=homology;ID=TE_homo_62001;sequence_ontology=SO:0002280 met1_07

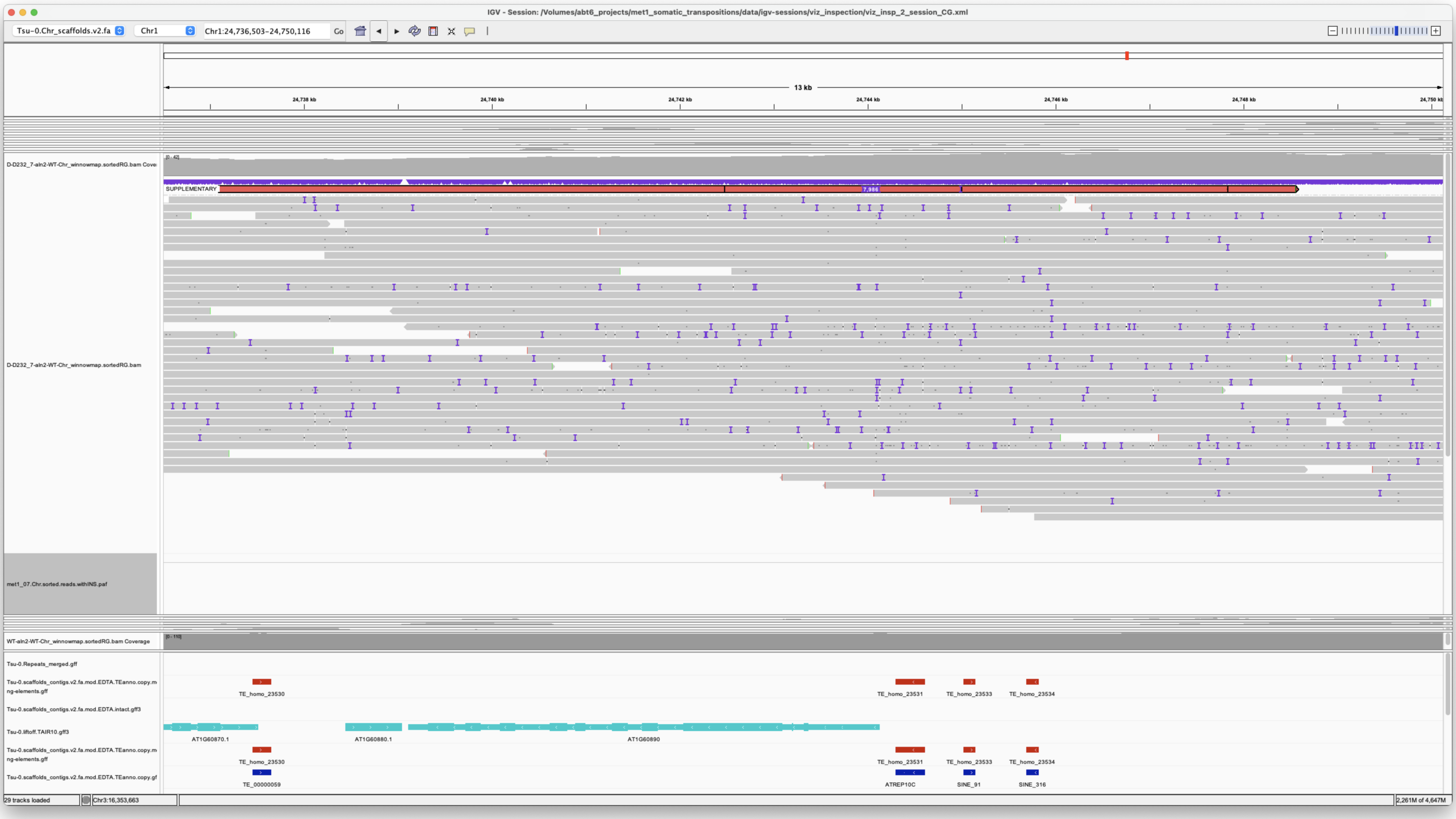


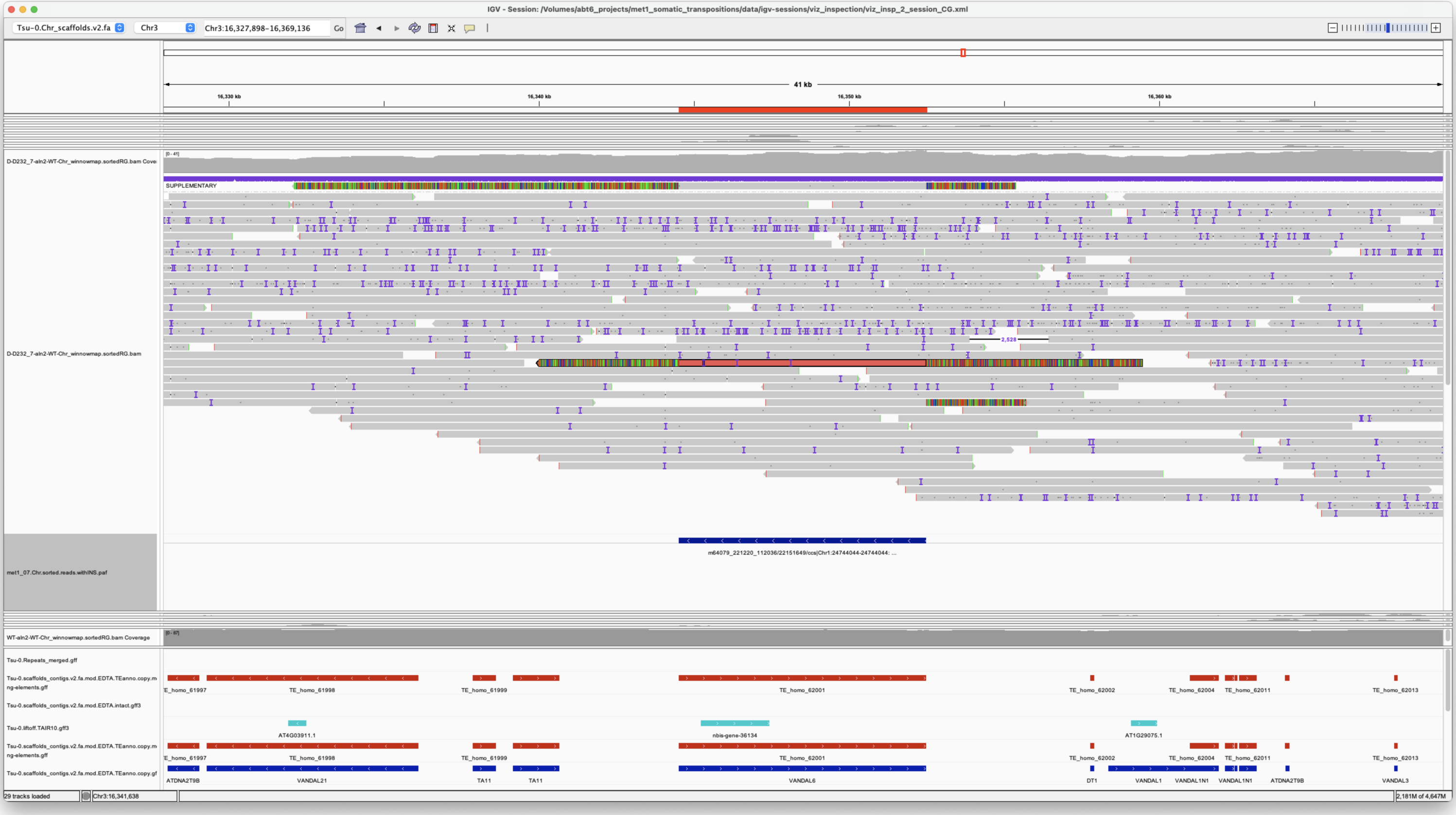


*Confirmed*

> Chr1 30975092 30975092 m64079_240212_113350/55118360/ccs Chr5 21419693 21425022
> Chr5|21419693|21425021|ID=LTRRT_299;Name=ATCOPIA93.2_Evade;Classification=LTR/Copia;Sequence_ontology=SO:0002264;ltr_identity=1.0000;Method=structural;motif=TACA;tsd=GGACA met1_07

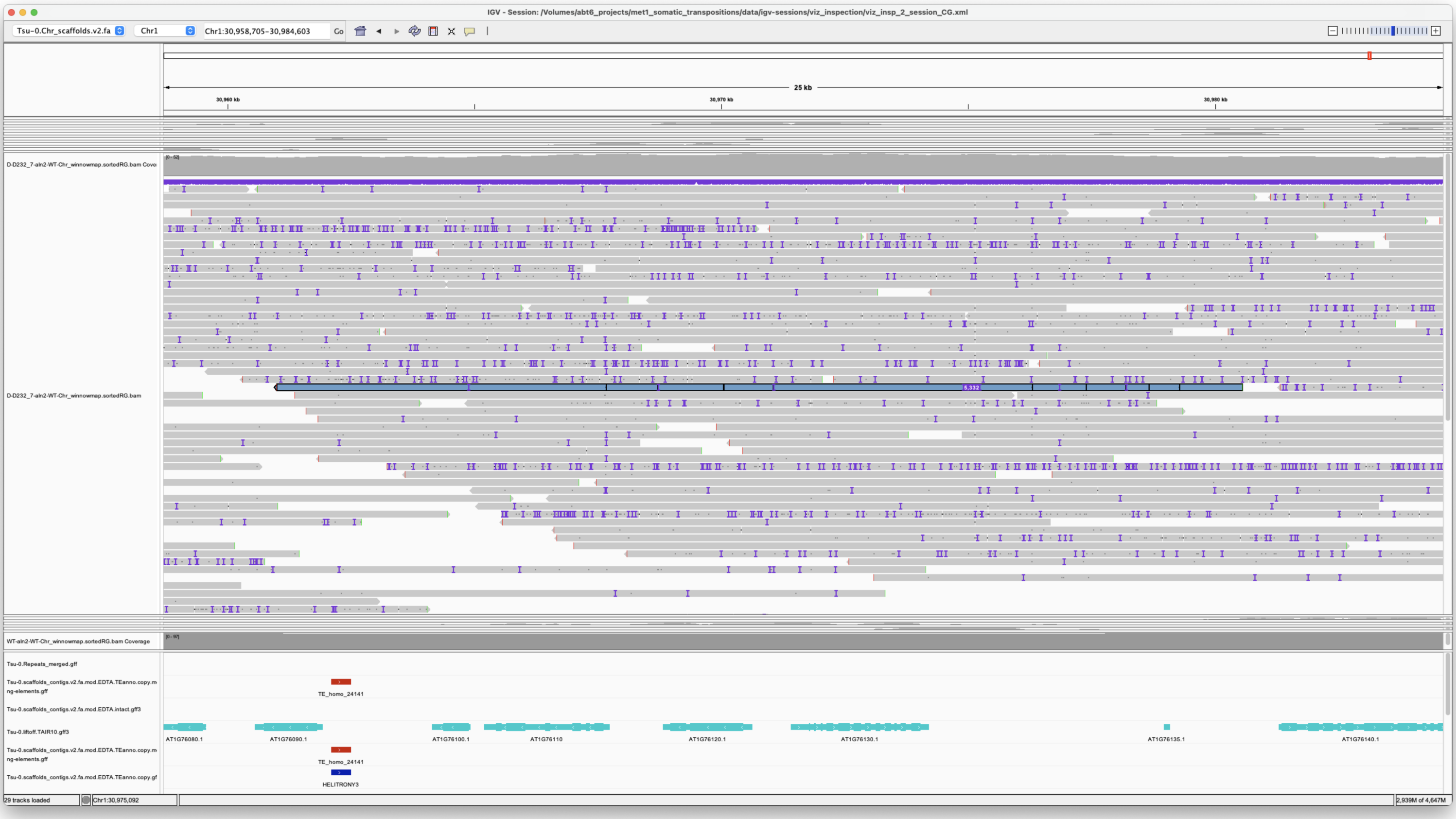


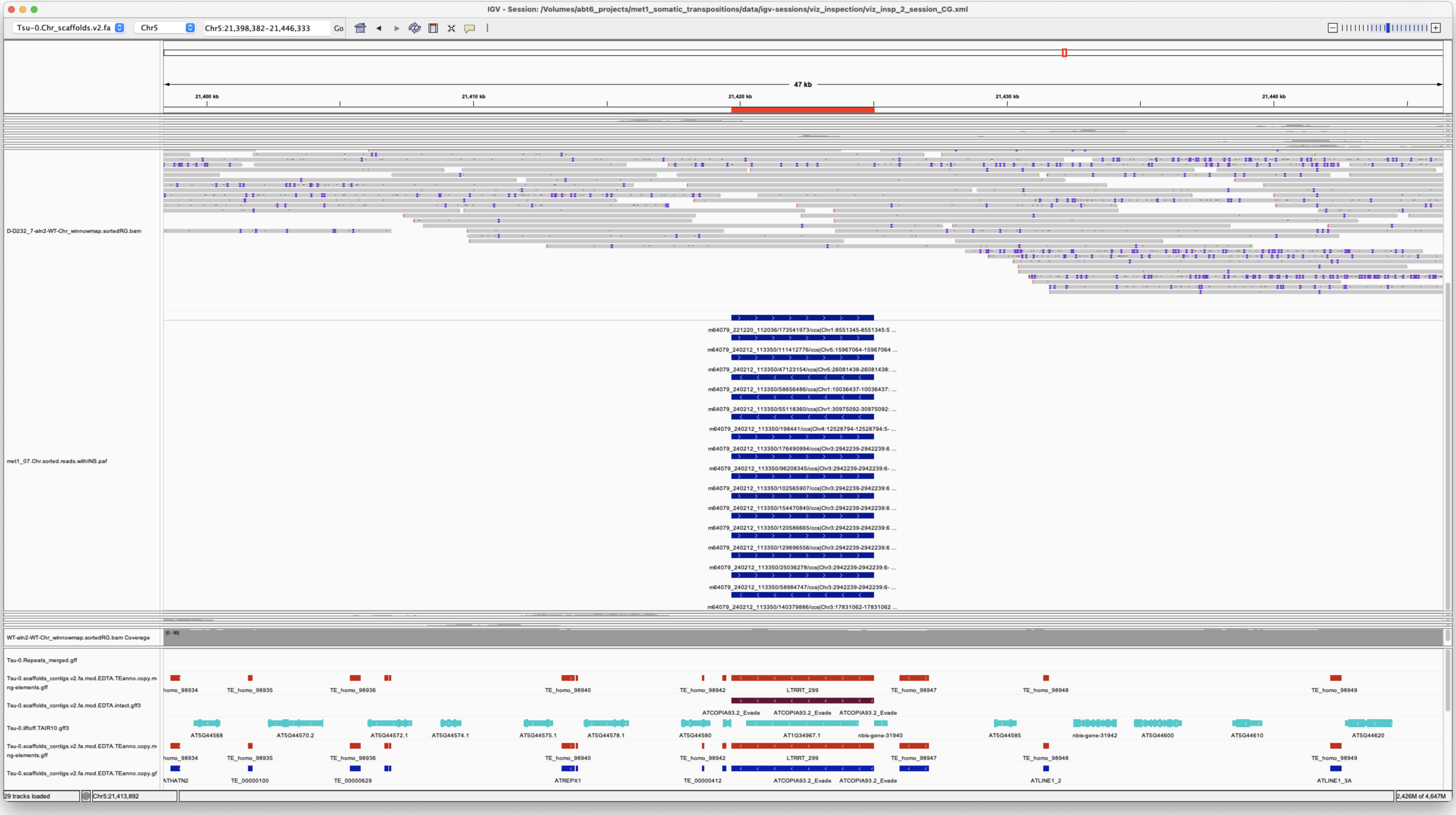


*Confirmed*

---

> Chr2 14209675 14209675 m64079_221220_112036/51970334/ccs Chr5 875413 876434 Chr5|875414|876433|ID=TE_MANUAL_02;Name=PutativePackTypeCACTAMuDR;classification=DNA/DTC;sequence_ontology=MANUAL;identity=MANUAL;method=MANUAL;ID=TE_MANUAL_02;sequence_ontology=MANUAL met1_07

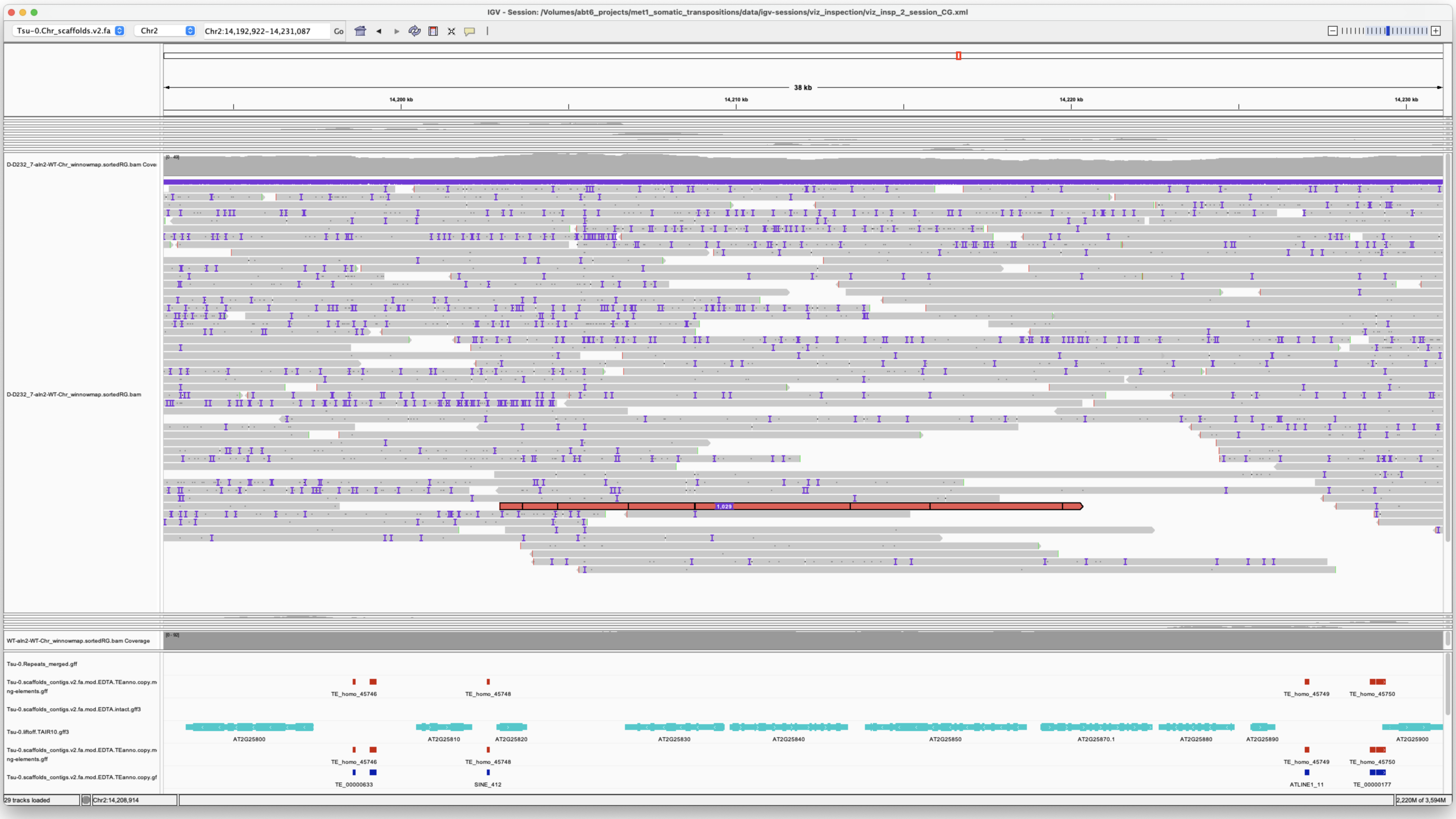


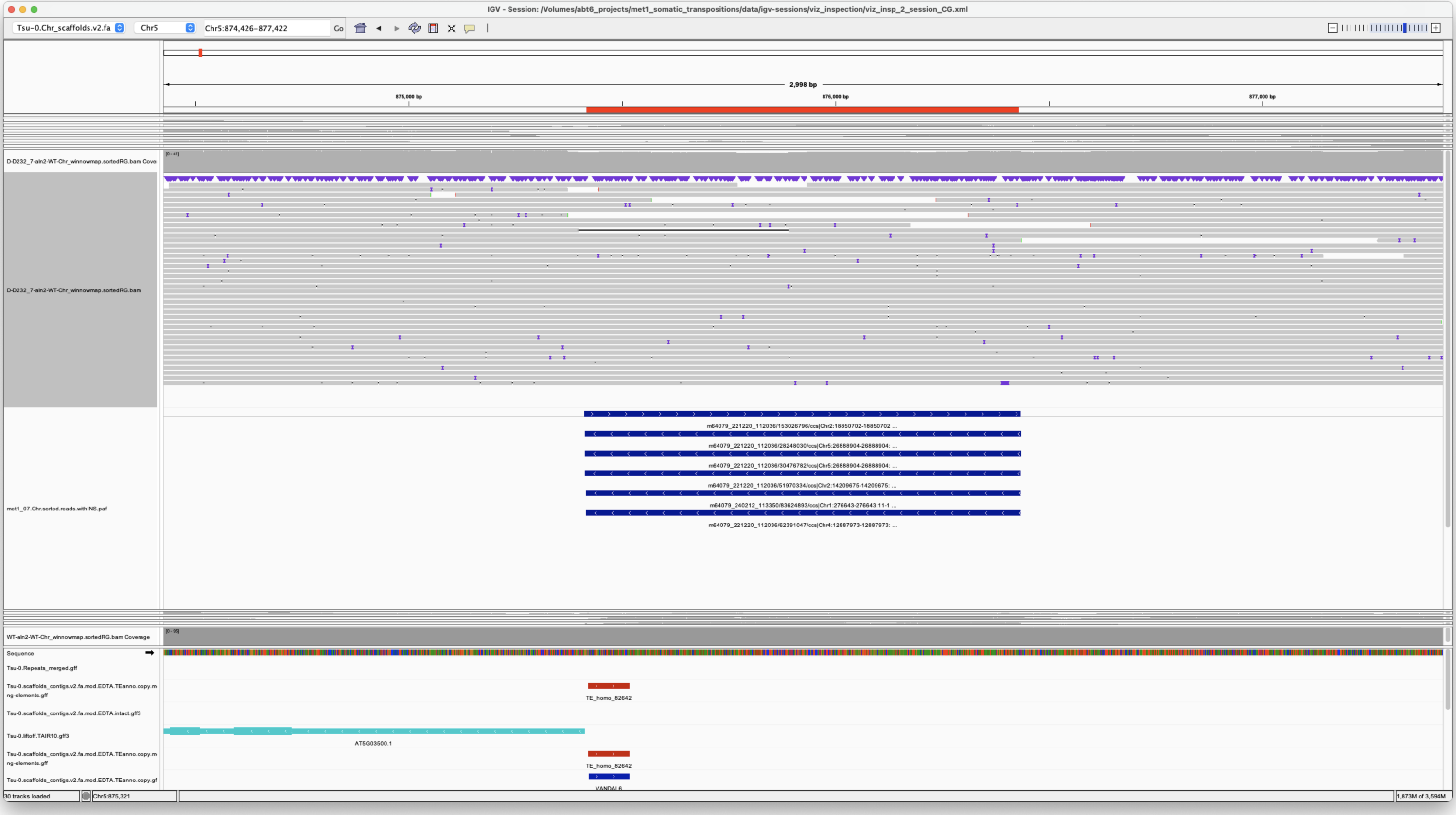


*Confirmed*

---

> Chr2 18850702 18850702 m64079_221220_112036/153026796/ccs Chr5 875412 876434 Chr5|875414|876433|ID=TE_MANUAL_02;Name=PutativePackTypeCACTAMuDR;classification=DNA/DTC;sequence_ontology=MANUAL;identity=MANUAL;method=MANUAL;ID=TE_MANUAL_02;sequence_ontology=MANUAL met1_07

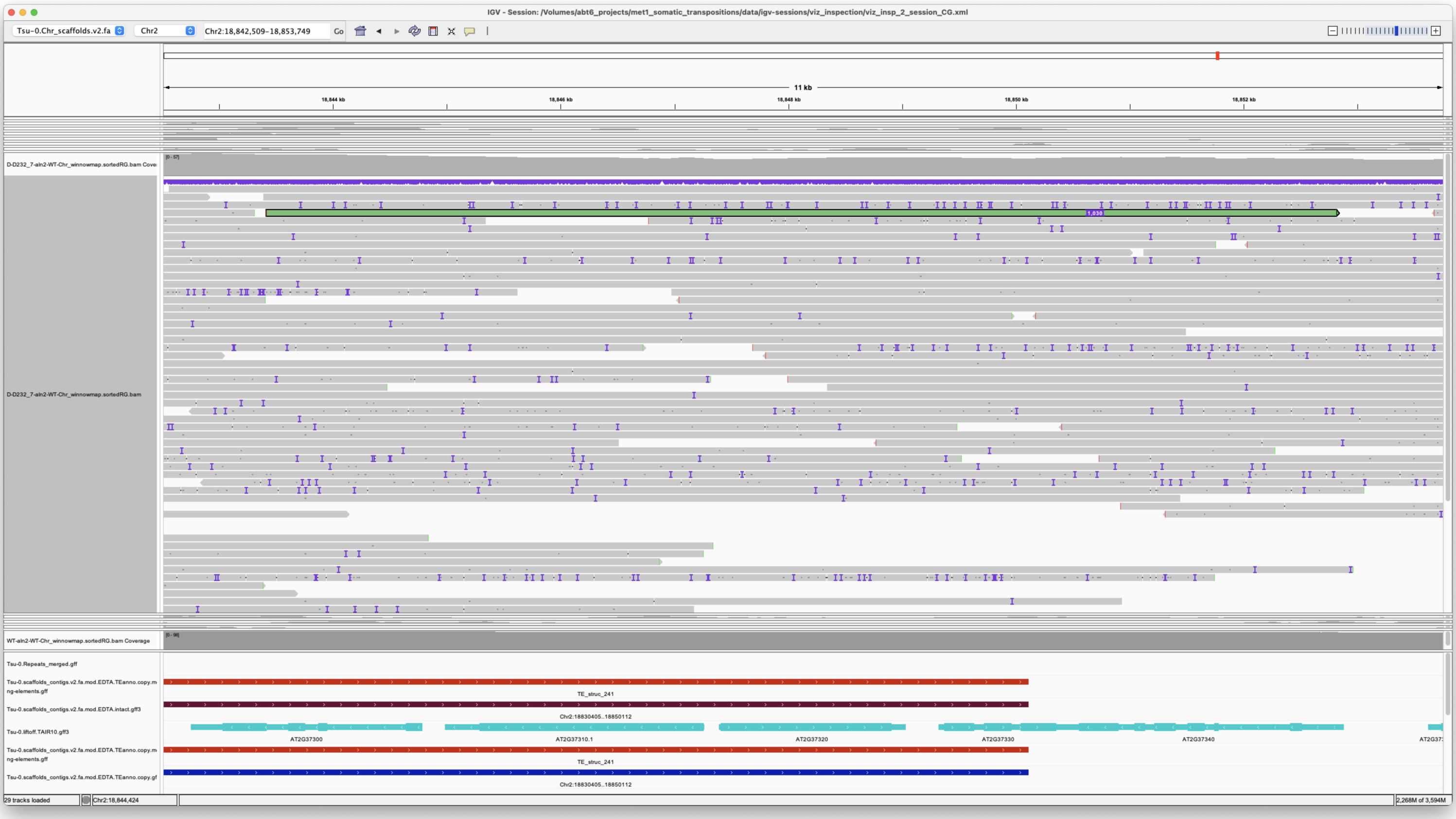


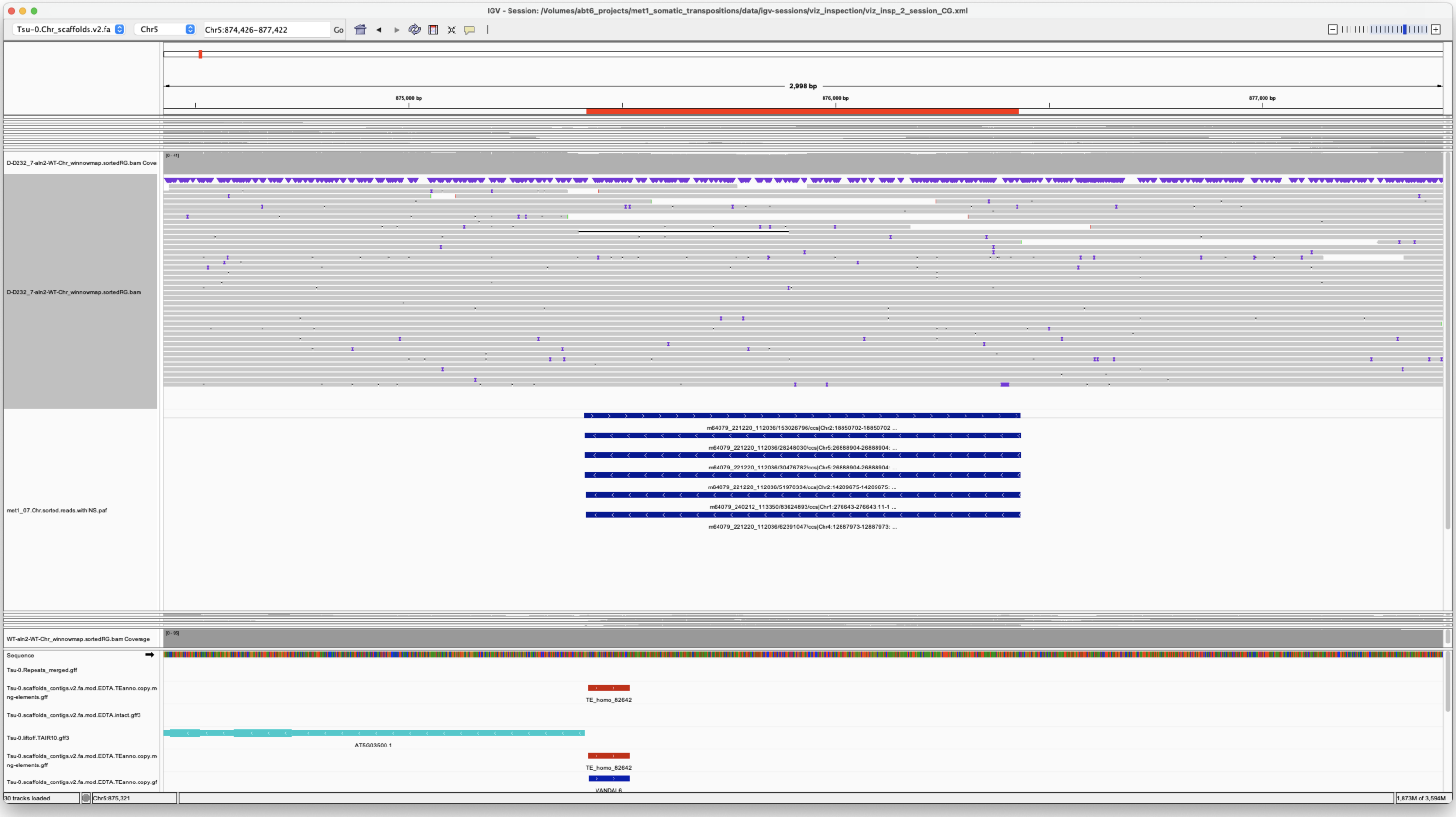


*Confirmed*

---

> Chr3 2942238 2942240
> m64079_240212_113350/102565907/ccs**m64079_240212_113350/120586665/ccs**m64079_240212_113350/129696556/ccs**m64079_240212_113350/154470840/ccs**m64079_240212_113350/176490994/ccs**m64079_240212_113350/25036278/ccs**m64079_240212_113350/58984747/ccs**m64079_240212_113350/96208345/ccs Chr1 11941106 11946435**
> **Chr1|11941106|11946435|ID=LTRRT_5;Name=ATCOPIA93.2_Evade;Classification=LTR/Copia;Sequence_ontology=SO:0002264;ltr_identity=1.0000;Method=structural;motif=TACA;tsd=ATATG**Chr1|11941106|11946435|ID=LTRRT_5;Name=ATCOPIA93.2_Evade;Classification=LTR/Copia;Sequence_ontology=SO:0002264;ltr_identity=1.0000;Method=structural;motif=TACA;tsd=ATATG**Chr1|11941106|11946435|ID=LTRRT_5;Name=ATCOPIA93.2_Evade;Classification=LTR/Copia;Sequence_ontology=SO:0002264;ltr_identity=1.0000;Method=structural;motif=TACA;tsd=ATATG**Chr1|11941106|11946435|ID=LTRRT_5;Name=ATCOPIA93.2_Evade;Classification=LTR/Copia;Sequence_ontology=SO:0002264;ltr_identity=1.0000;Method=structural;motif=TACA;tsd=ATATG**Chr1|11941106|11946435|ID=LTRRT_5;Name=ATCOPIA93.2_Evade;Classification=LTR/Copia;Sequence_ontology=SO:0002264;ltr_identity=1.0000;Method=structural;motif=TACA;tsd=ATATG**Chr1|11941106|11946435|ID=LTRRT_5;Name=ATCOPIA93.2_Evade;Classification=LTR/Copia;Sequence_ontology=SO:0002264;ltr_identity=1.0000;Method=structural;motif=TACA;tsd=ATATG**Chr1|11941106|11946435|ID=LTRRT_5;Name=ATCOPIA93.2_Evade;Classification=LTR/Copia;Sequence_ontology=SO:0002264;ltr_identity=1.0000;Method=structural;motif=TACA;tsd=ATATG**Chr1|11941106|11946435|ID=LTRRT_5;Name=ATCOPIA93.2_Evade;Classification=LTR/Copia;Sequence_ontology=SO:0002264;ltr_identity=1.0000;Method=structural;motif=TACA;tsd=ATATG met1_07

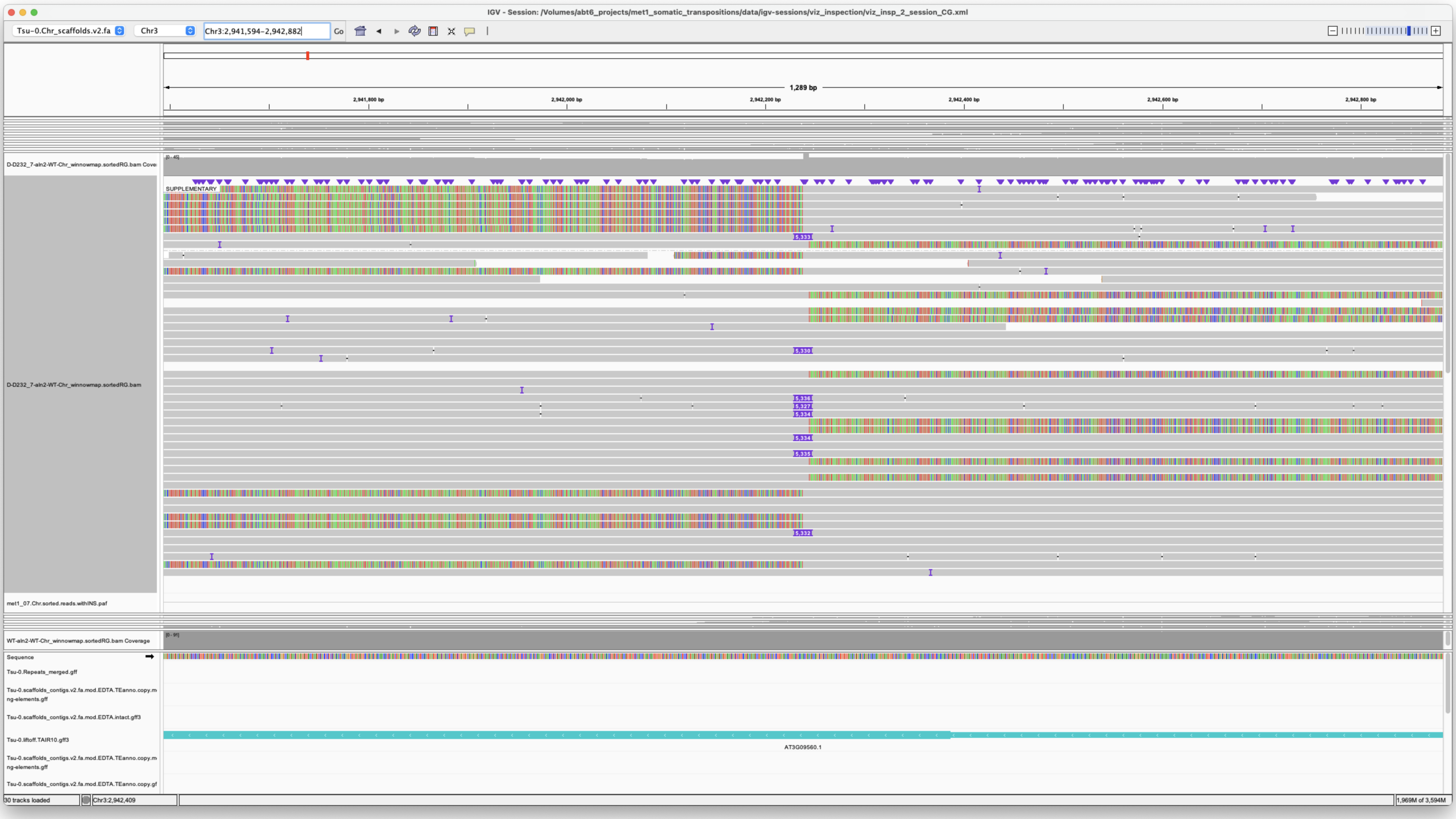


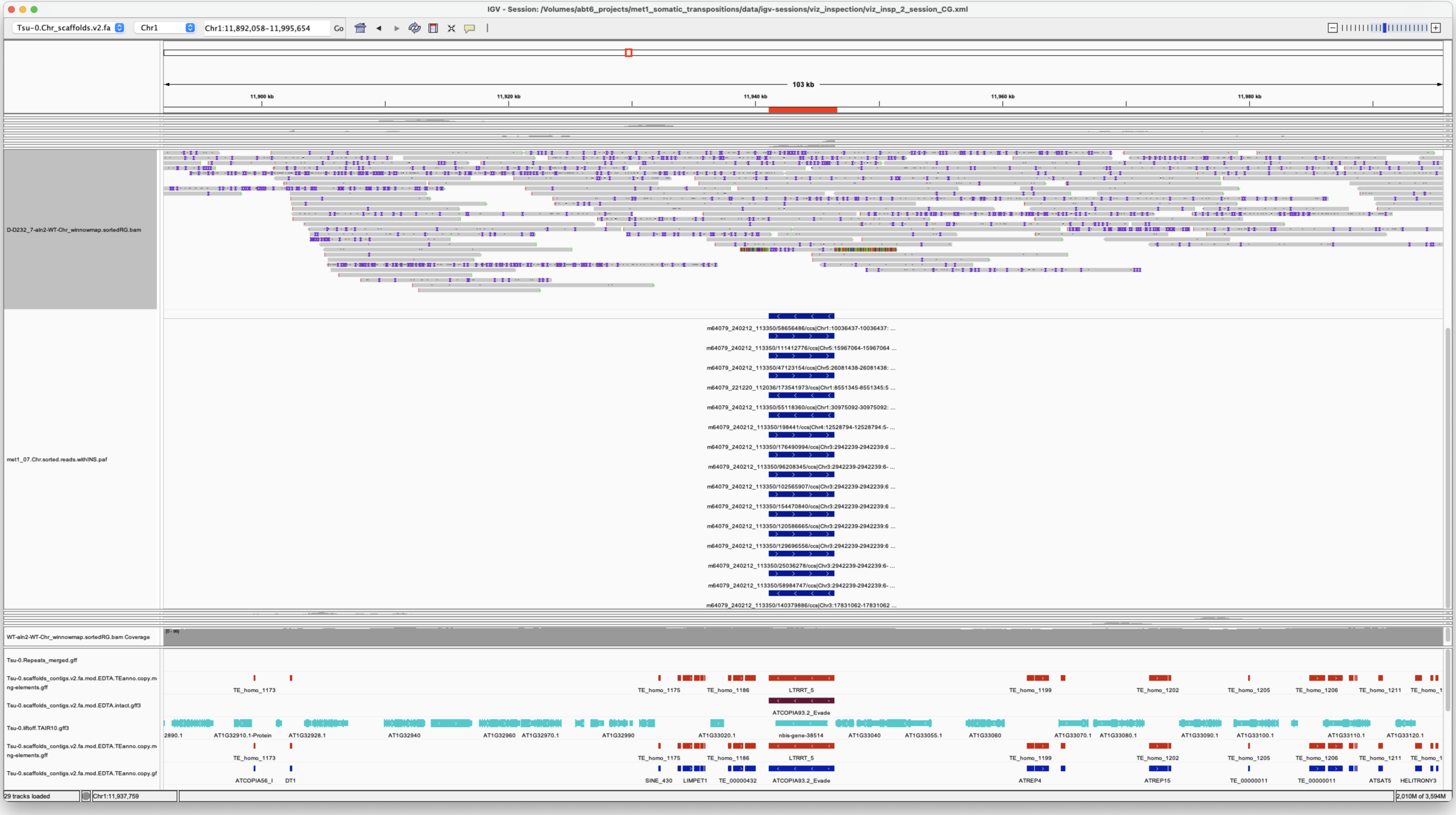


*Confirmed*

---

> Chr3 17831062 17831062 m64079_240212_113350/140379886/ccs Chr1 11941107 11946436
> Chr1|11941106|11946435|ID=LTRRT_5;Name=ATCOPIA93.2_Evade;Classification=LTR/Copia;Sequence_ontology=SO:0002264;ltr_identity=1.0000;Method=structural;motif=TACA;tsd=ATATG met1_07

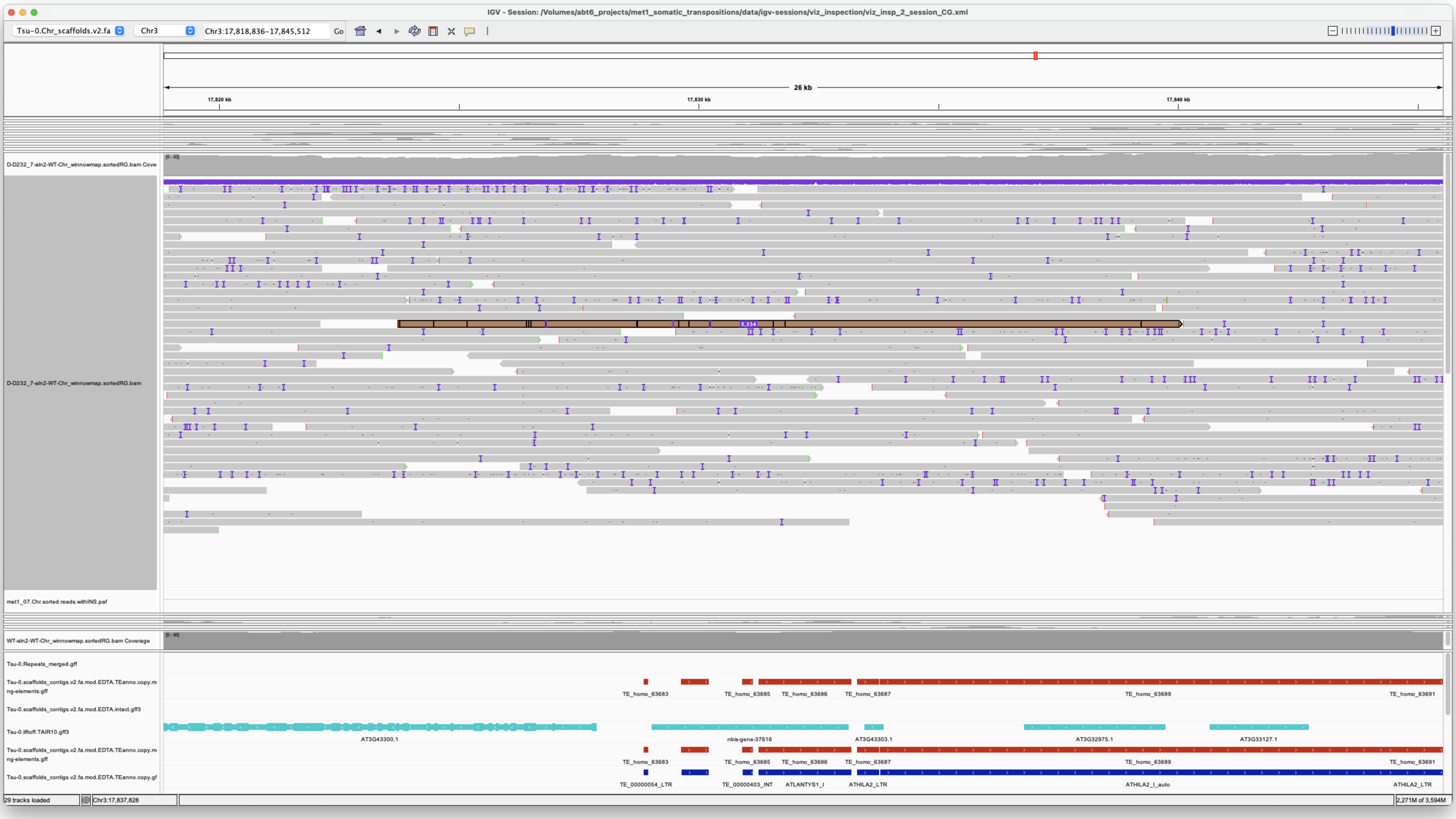


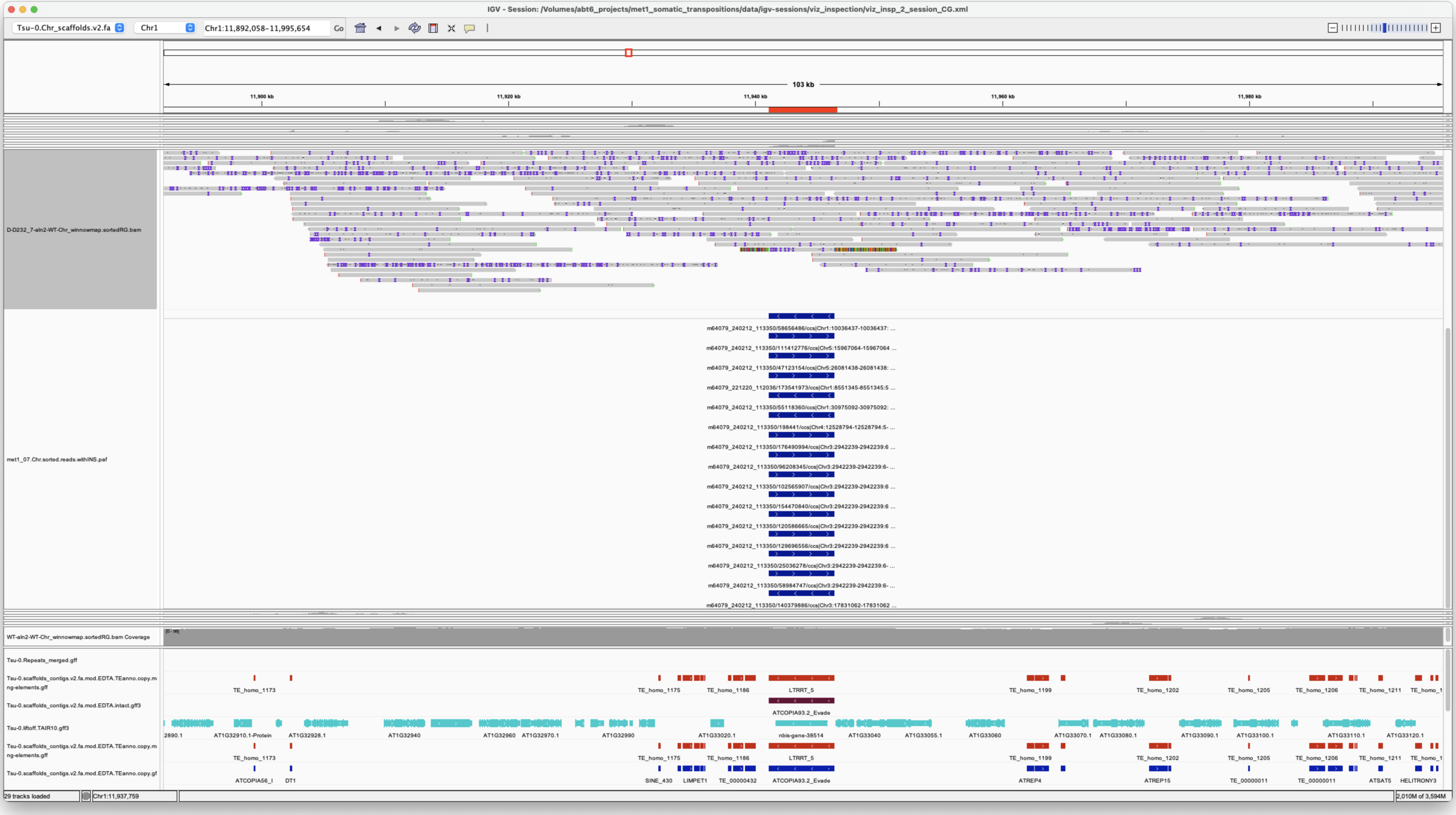


*Confirmed*

---

Chr4 12528794 12528794 m64079_240212_113350/198441/ccs Chr1 11941106 11946436 Chr1|11941106|11946435|ID=LTRRT_5;Name=ATCOPIA93.2_Evade;Classification=LTR/Copia;Sequence_ontology=SO:0002264;ltr_identity=1.0000;Method=structural;motif=TACA;tsd=ATATG met1_07

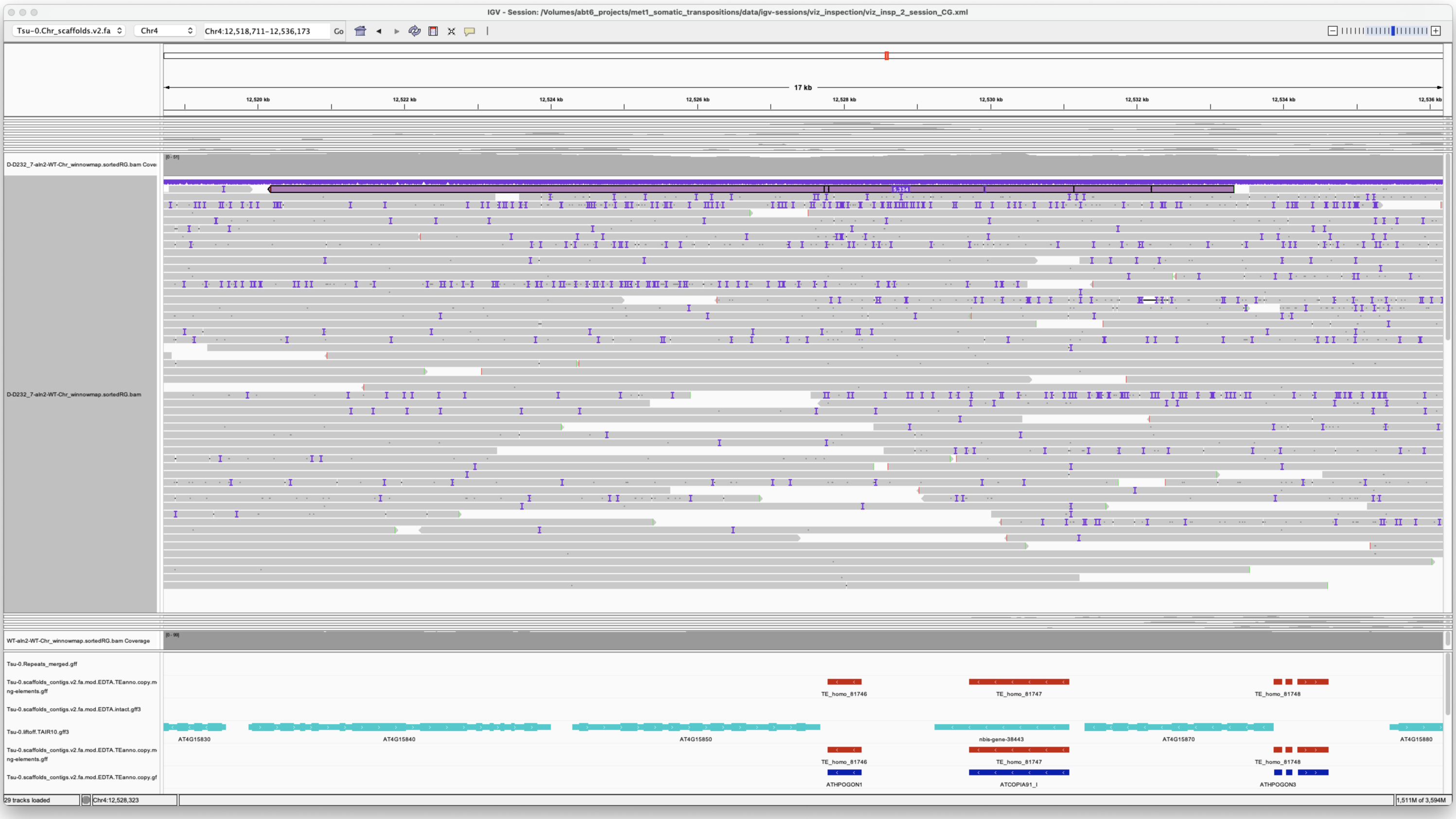

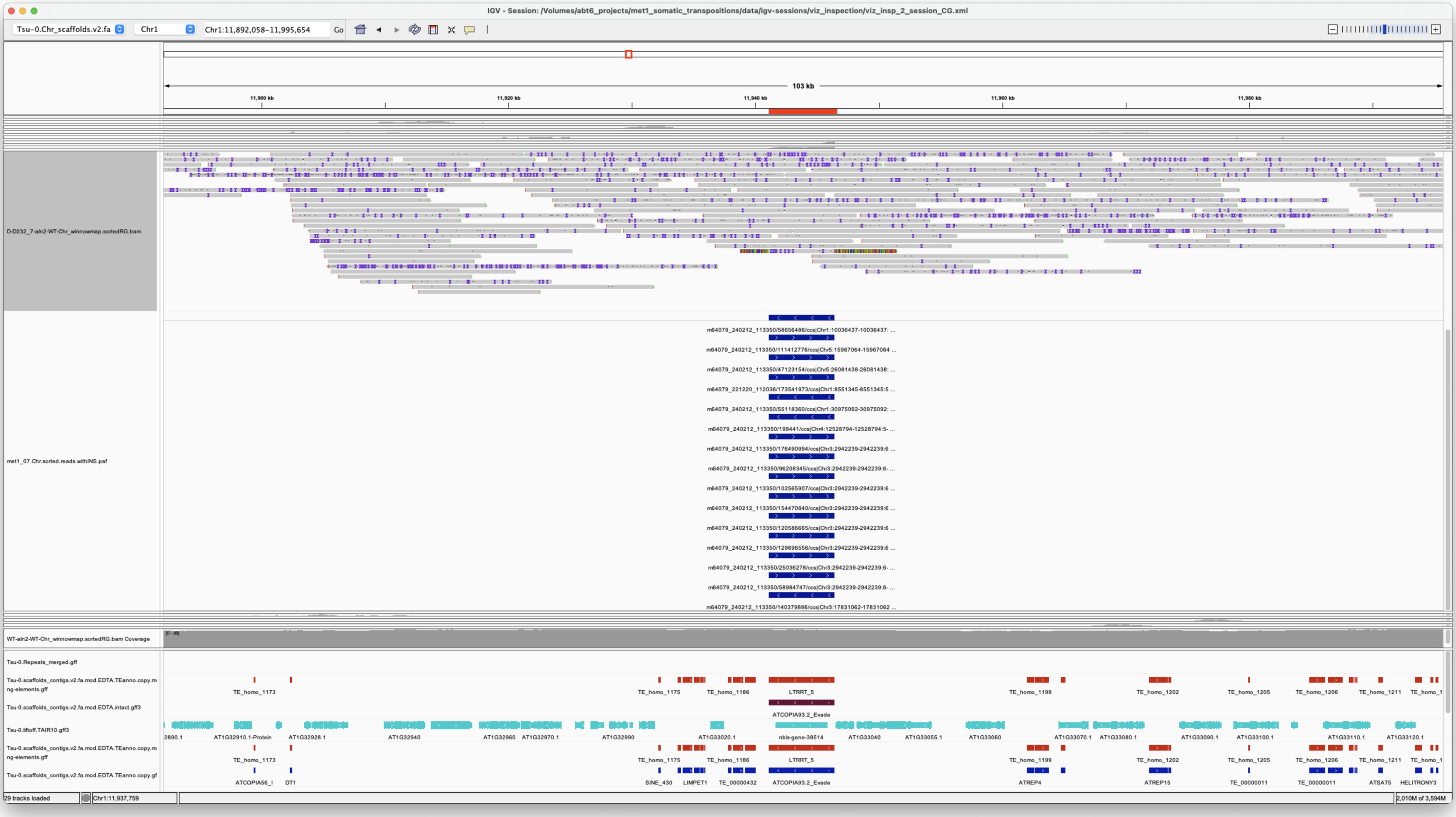

*Confirmed*

---

> Chr4 12887973 12887973 m64079_221220_112036/62391047/ccs Chr5 875415 876434
> Chr5|875414|876433|ID=TE_MANUAL_02;Name=PutativePackTypeCACTAMuDR;classification=DNA/DTC;sequence_ontology=MANUAL;identity=MANUAL;method=MANUAL;ID=TE_MANUAL_02;sequence_ontology=MANUAL met1_07

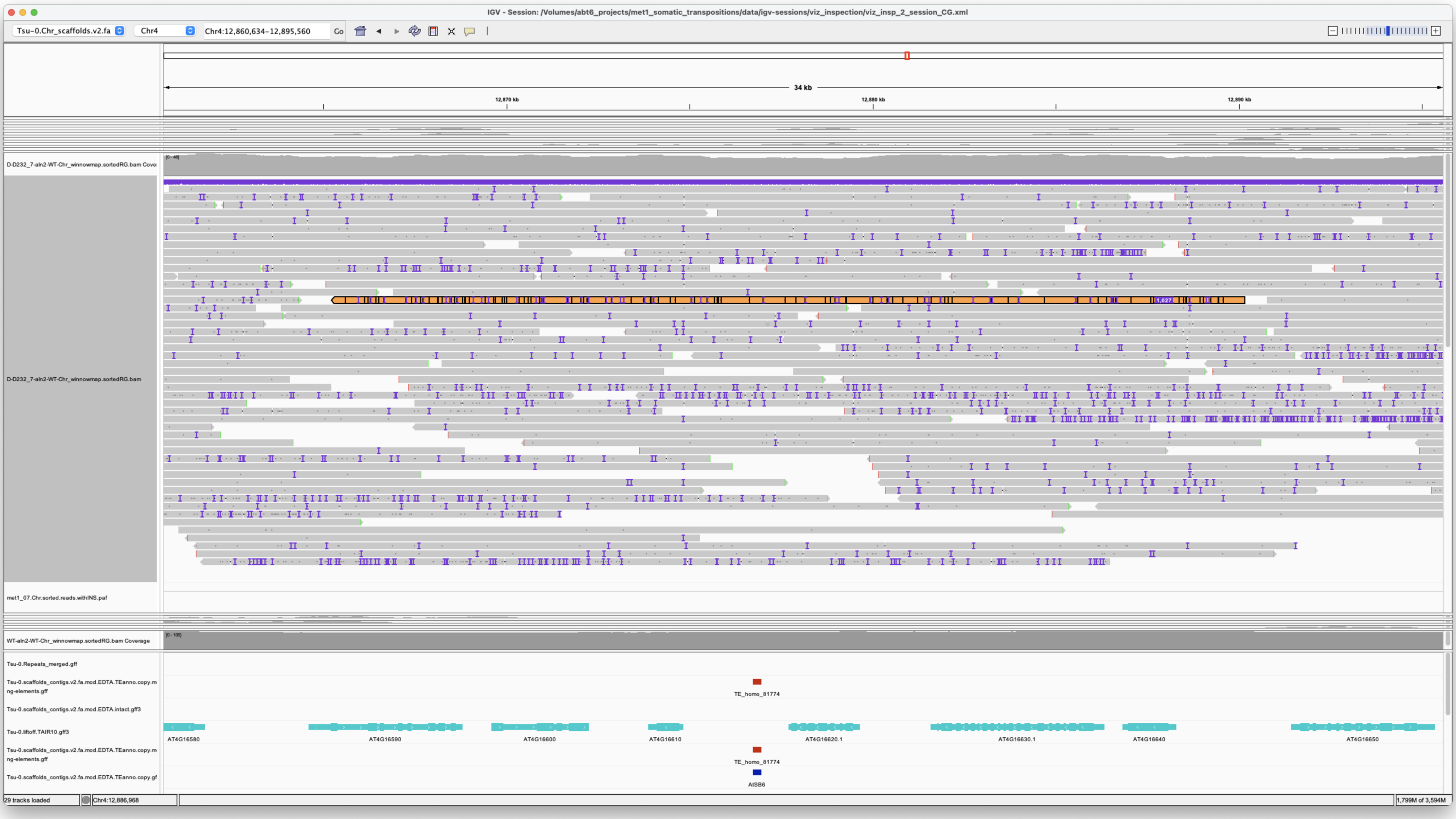


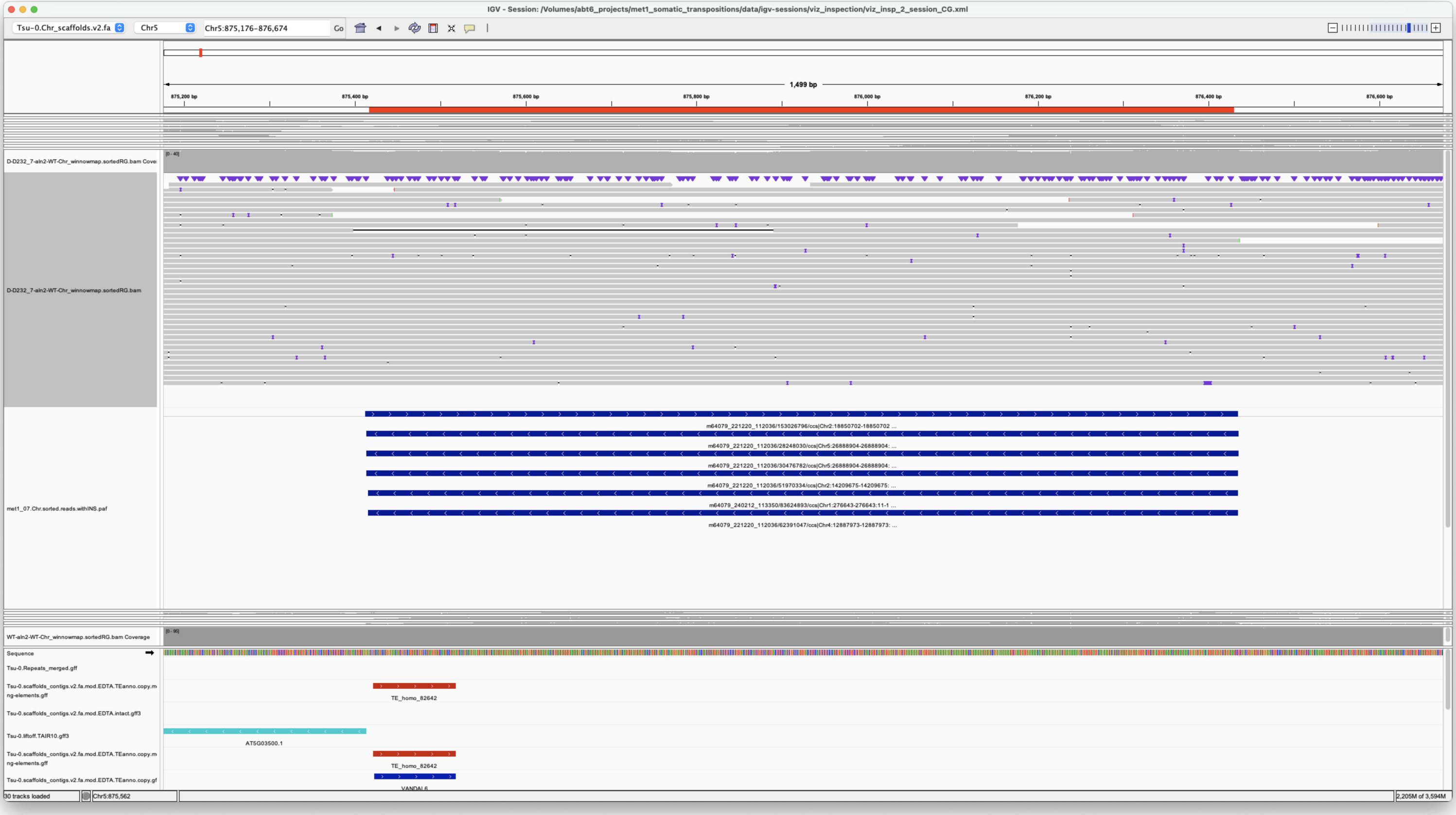


*Confirmed*

> Chr5 15967064 15967064 m64079_240212_113350/111412776/ccs Chr1 11941106 11946436
> Chr1|11941106|11946435|ID=LTRRT_5;Name=ATCOPIA93.2_Evade;Classification=LTR/Copia;Sequence_ontology=SO:0002264;ltr_identity=1.0000;Method=structural;motif=TACA;tsd=ATATG met1_07

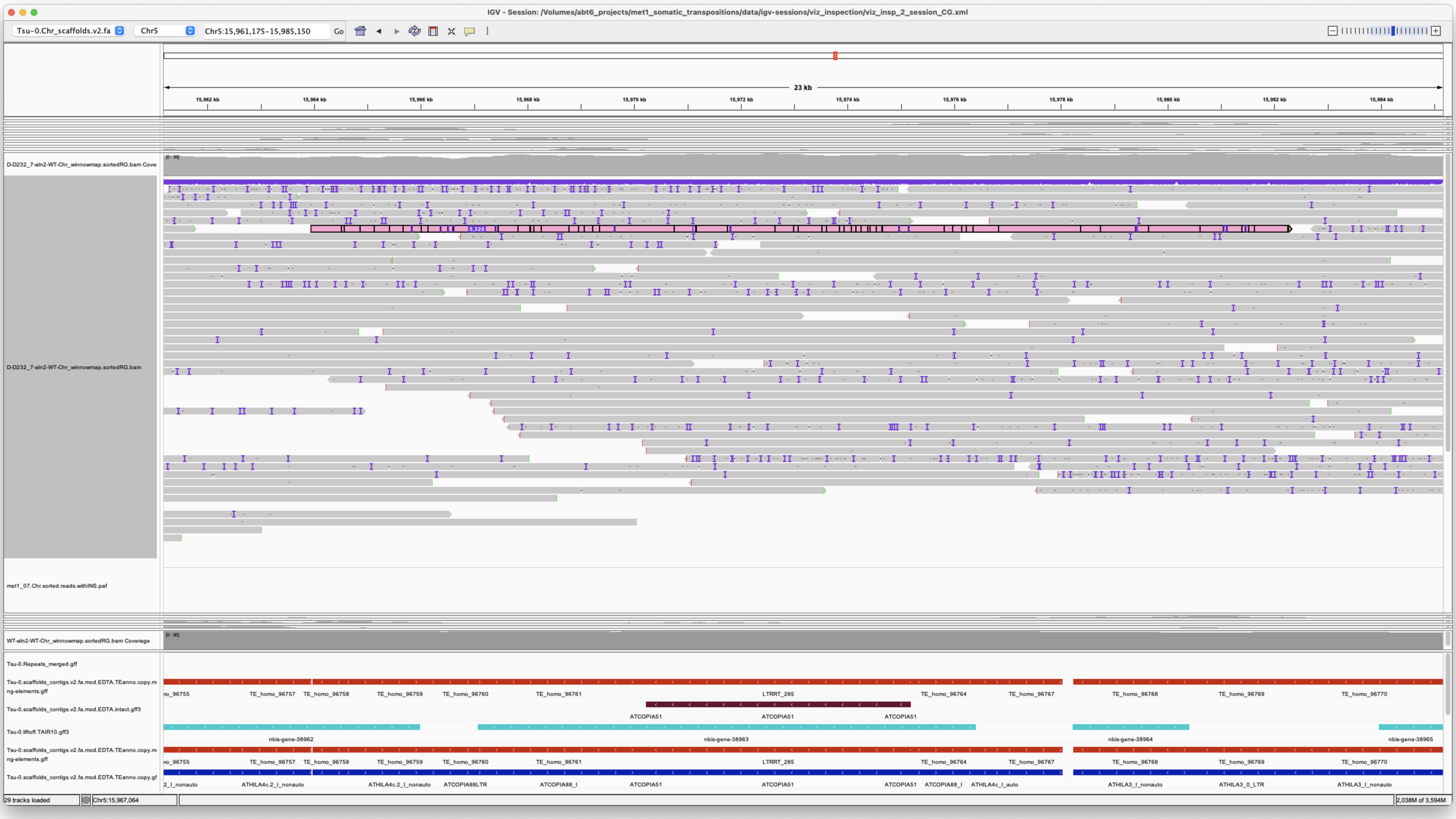

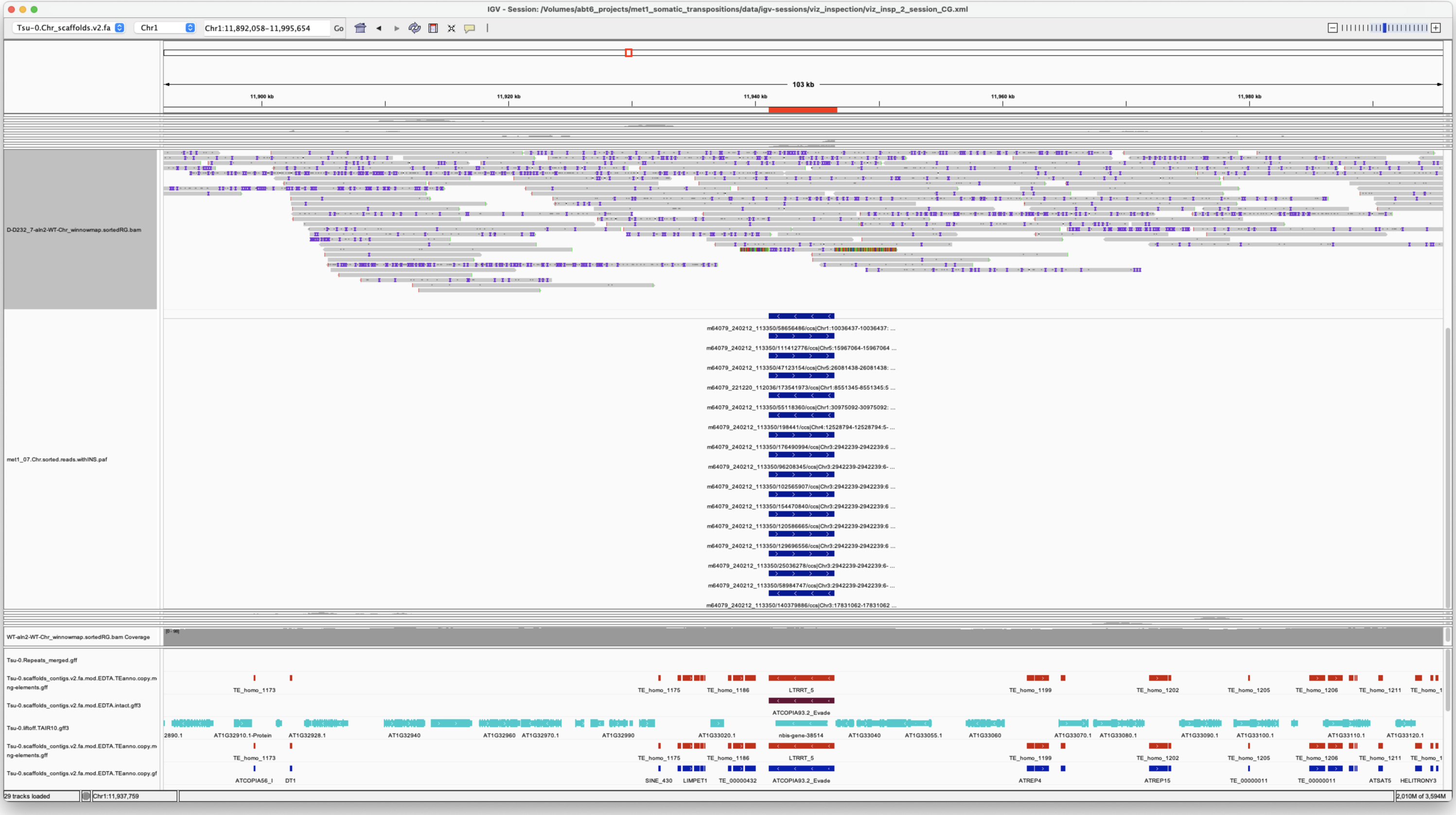

*Confirmed*

Chr5 26081438 26081438 m64079_240212_113350/47123154/ccs Chr1 11941106 11946436 Chr1|11941106|11946435|ID=LTRRT_5;Name=ATCOPIA93.2_Evade;Classification=LTR/Copia;Sequence_ontology=SO:0002264;ltr_identity=1.0000;Method=structural;motif=TACA;tsd=ATATG met1_07

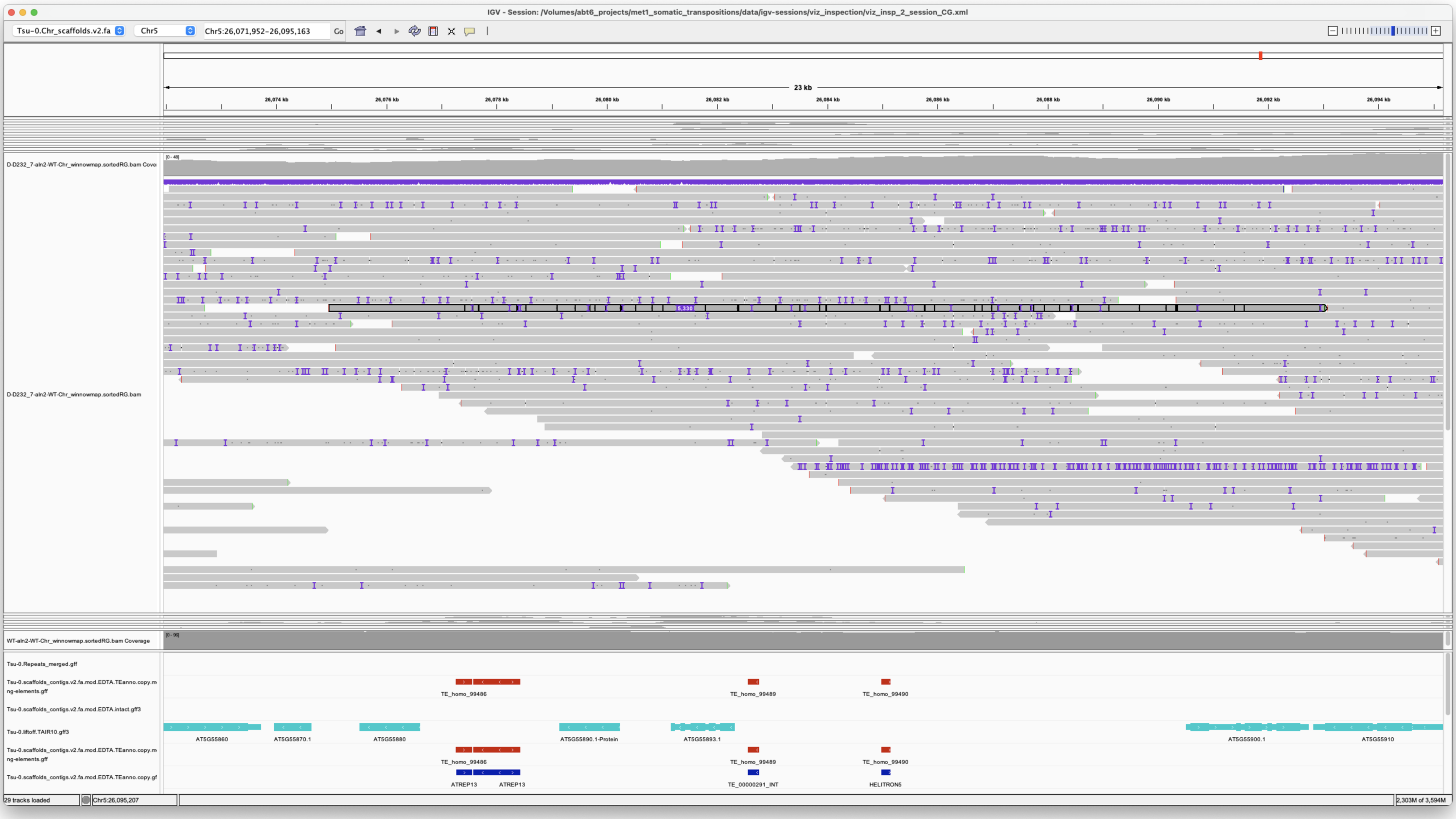


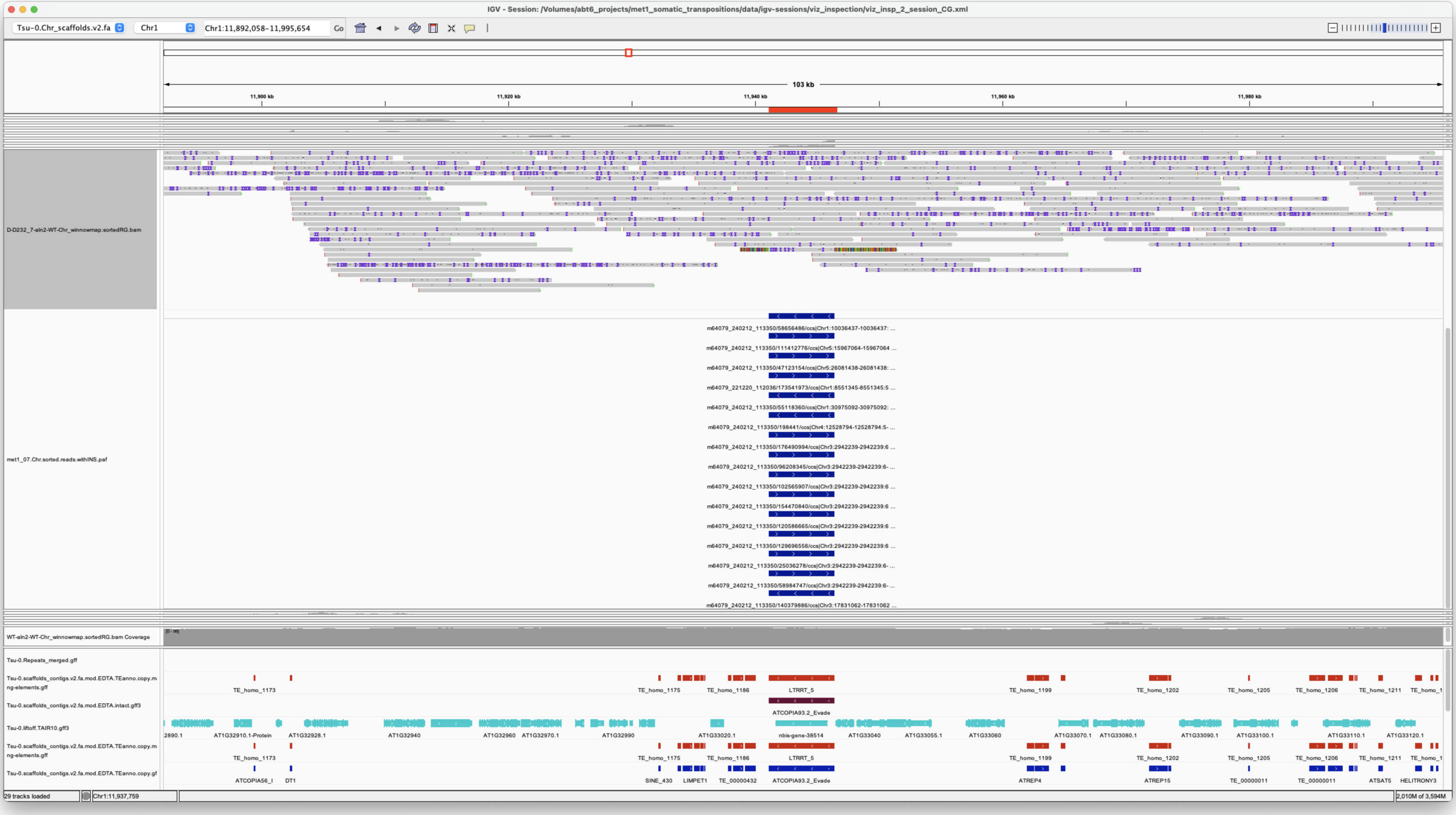


*Confirmed*

> Chr5 26888903 26888905 m64079_221220_112036/28248030/ccs**m64079_221220_112036/30476782/ccs Chr5 875413 876435 Chr5|875414|876433|ID=TE_MANUAL_02;Name=PutativePackTypeCACTAMuDR;classification=DNA/DTC;sequence_ontology=MANUAL;identity=MANUAL;method=MANUAL;ID=TE_MANUAL_02;sequence_ontology=MANUAL**Chr5|875414|876433|ID=TE_MANUAL_02;Name=PutativePackTypeCACTAMuDR;classification=DNA/DTC;sequence_ontology=MANUAL;identity=MANUAL;method=MANUAL;ID=TE_MANUAL_02;sequence_ontology=MANUAL met1_07

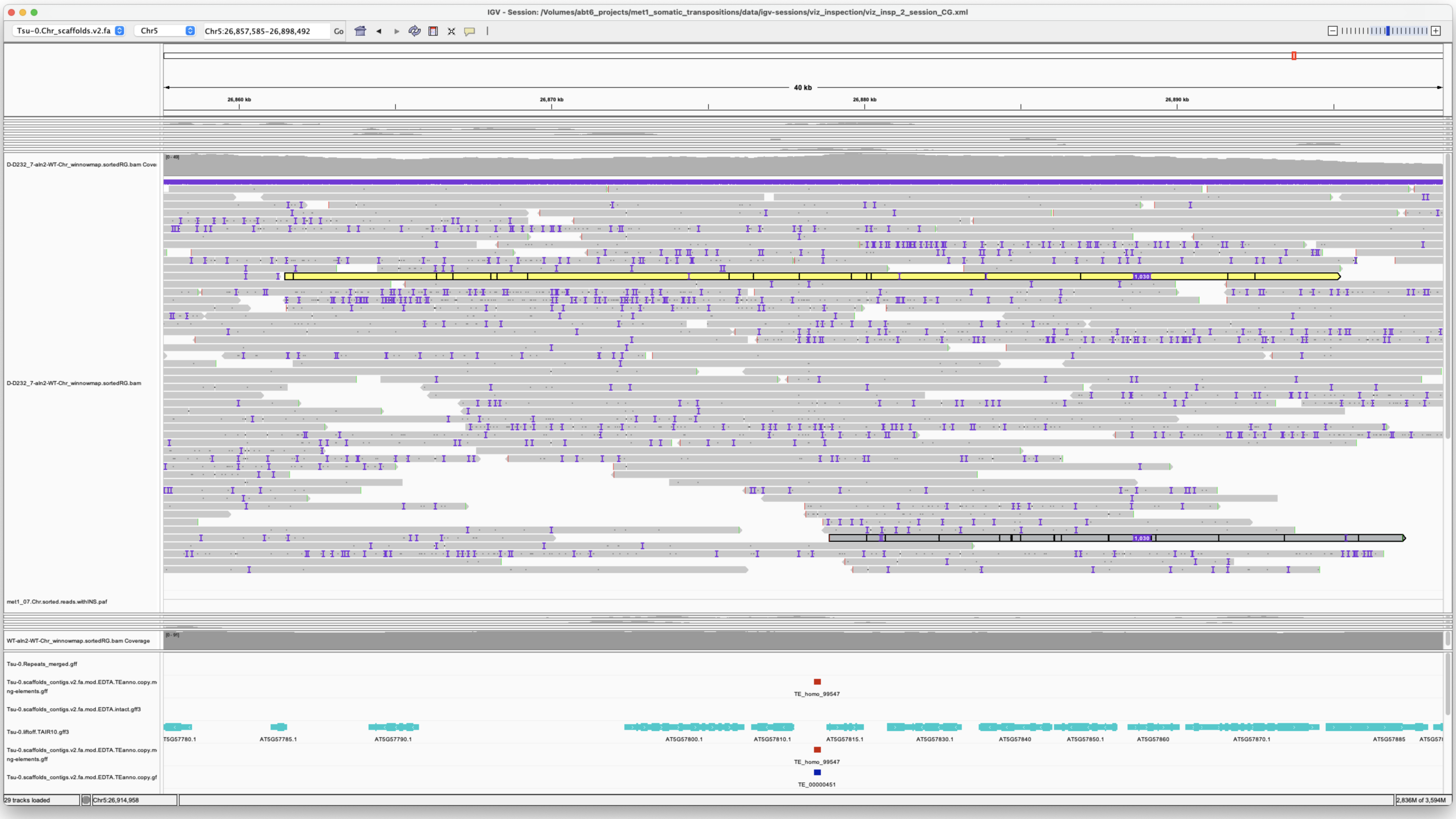


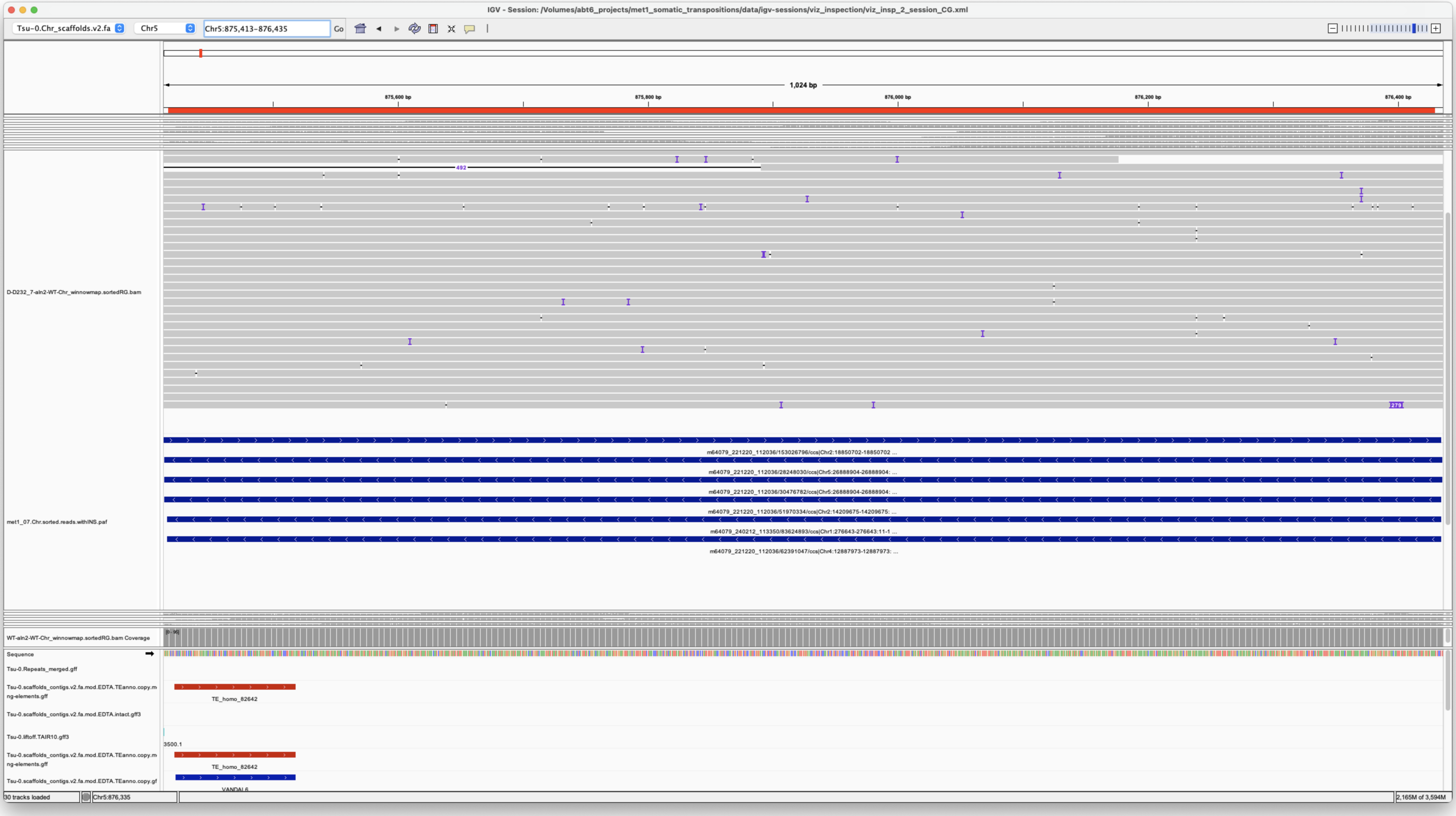


Beautiful

*Confirmed*

---

---

---

#### met1_08

---

> Chr2 14533452 14533452 m64079_221220_112036/101452109/ccs Chr1 11941106 11946441
> Chr1|11941106|11946435|ID=LTRRT_5;Name=ATCOPIA93.2_Evade;Classification=LTR/Copia;Sequence_ontology=SO:0002264;ltr_identity=1.0000;Method=structural;motif=TACA;tsd=ATATG met1_08

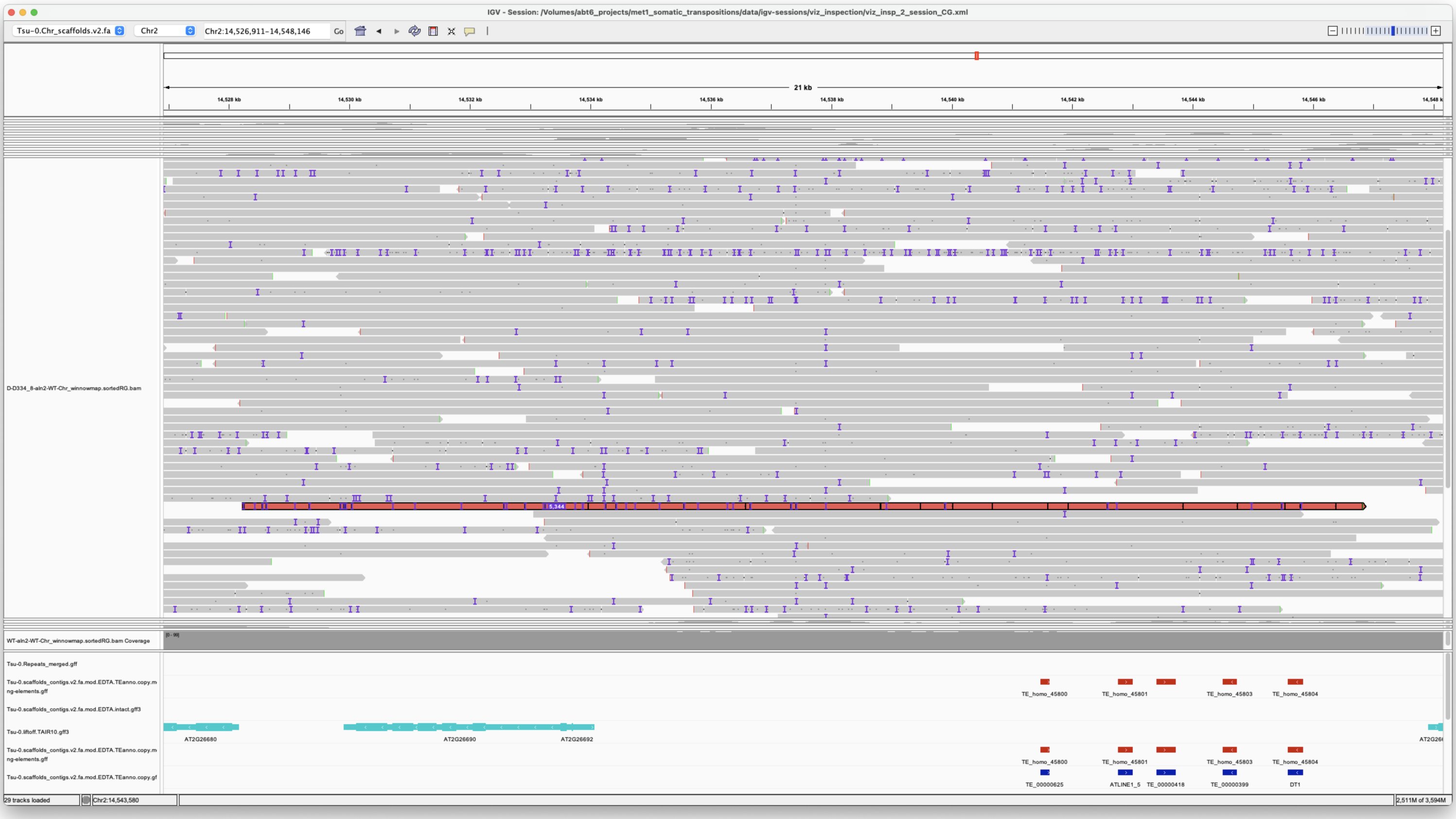


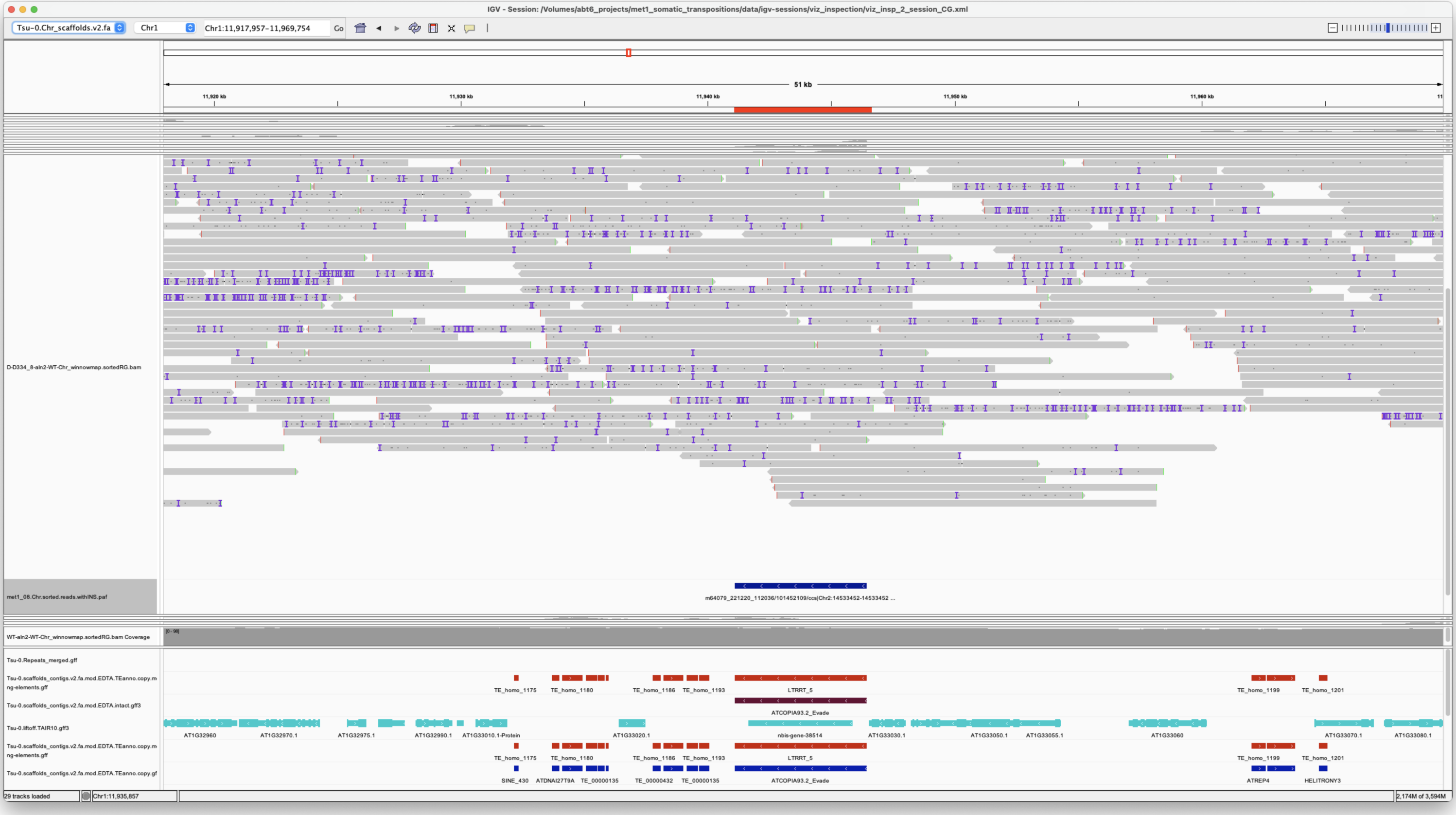


*Confirmed*

> Chr3 6900522 6900522 m64079_221220_112036/152111366/ccs Chr5 875413 876434 Chr5|875414|876433|ID=TE_MANUAL_02;Name=PutativePackTypeCACTAMuDR;classification=DNA/DTC;sequence_ontology=MANUAL;identity=MANUAL;method=MANUAL;ID=TE_MANUAL_02;sequence_ontology=MANUAL met1_08

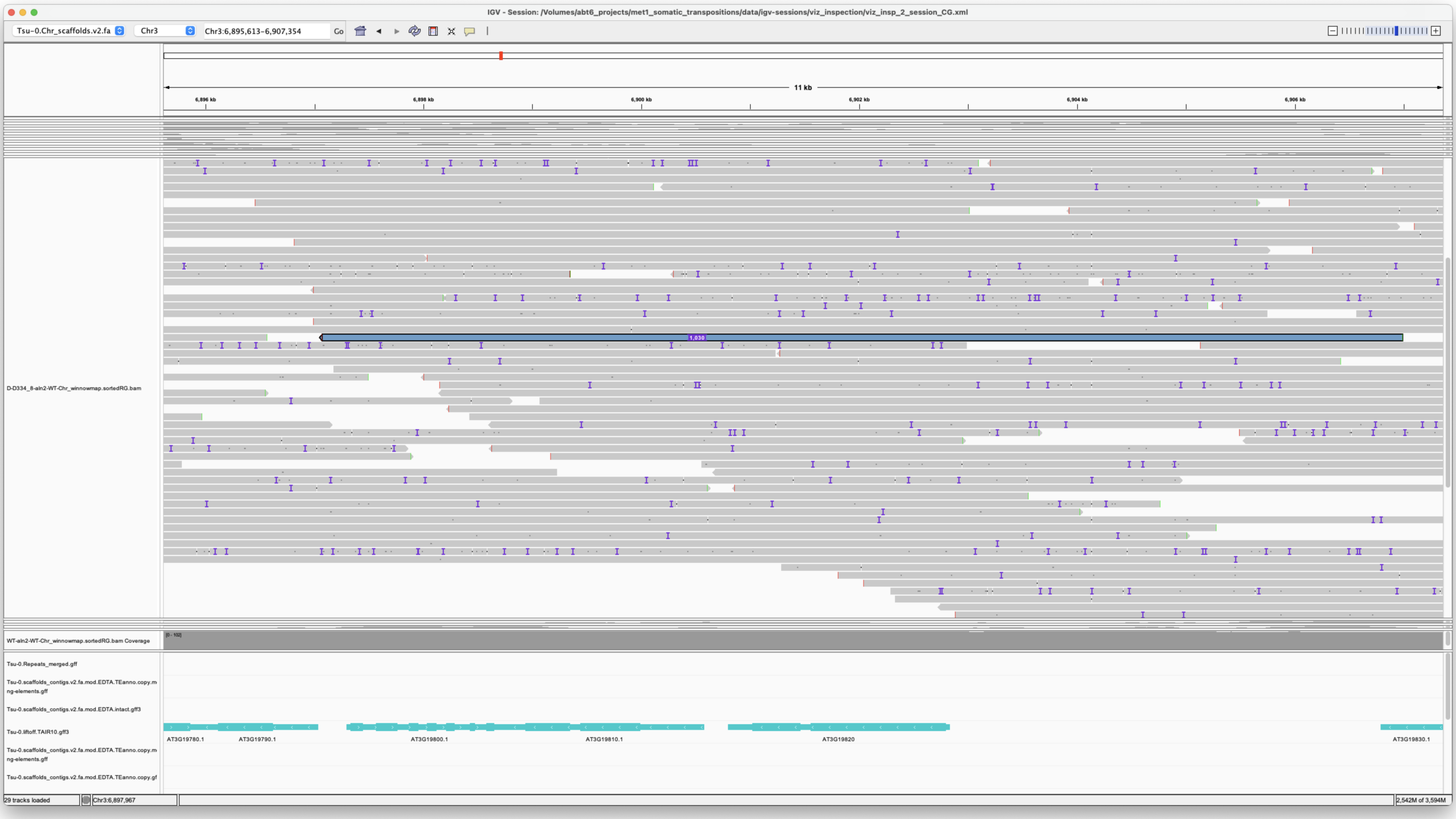


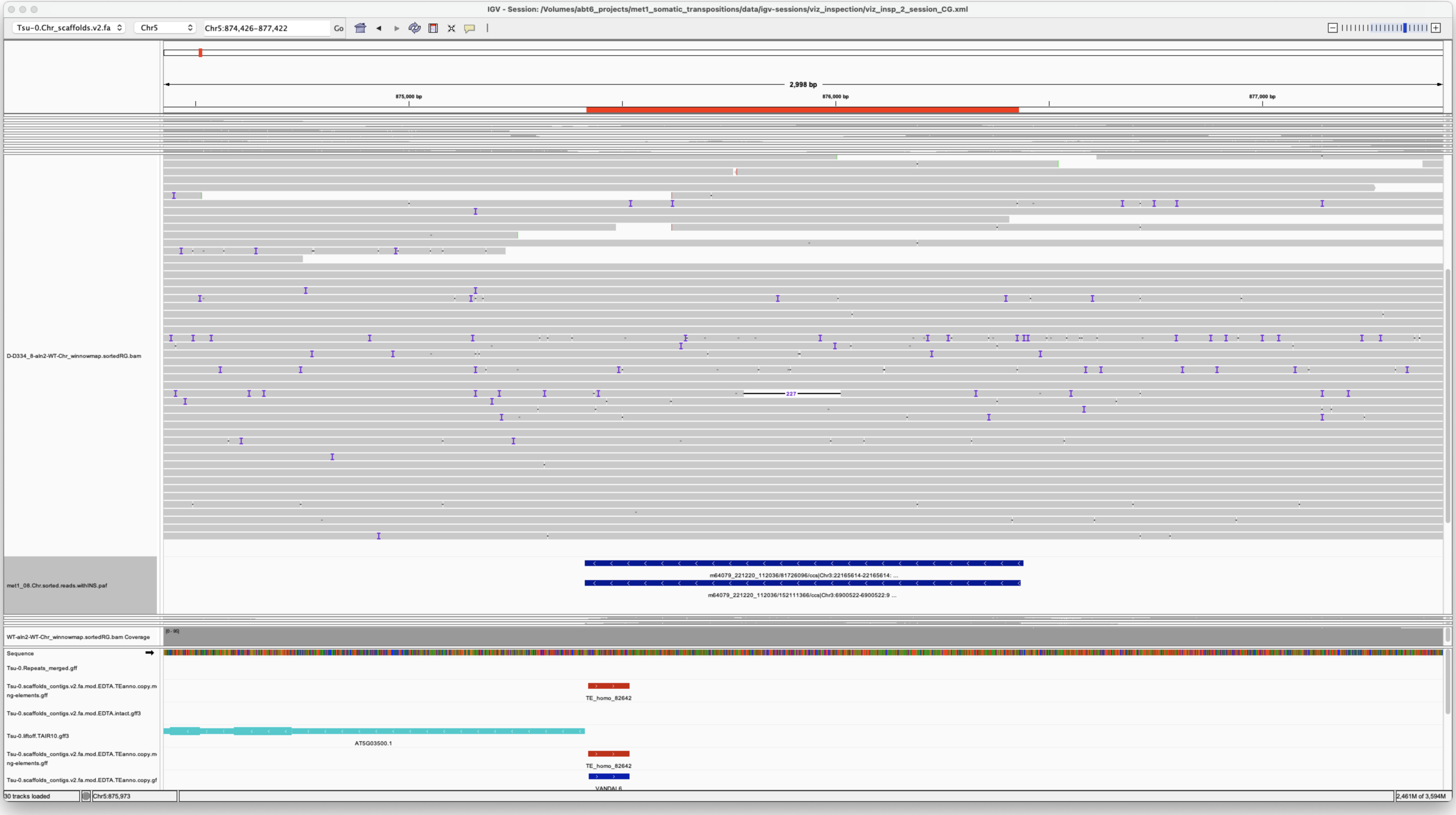


*Confirmed*

> Chr3 22165614 22165614 m64079_221220_112036/81726096/ccs Chr5 875413 876441 Chr5|875414|876433|ID=TE_MANUAL_02;Name=PutativePackTypeCACTAMuDR;classification=DNA/DTC;sequence_ontology=MANUAL;identity=MANUAL;method=MANUAL;ID=TE_MANUAL_02;sequence_ontology=MANUAL met1_08

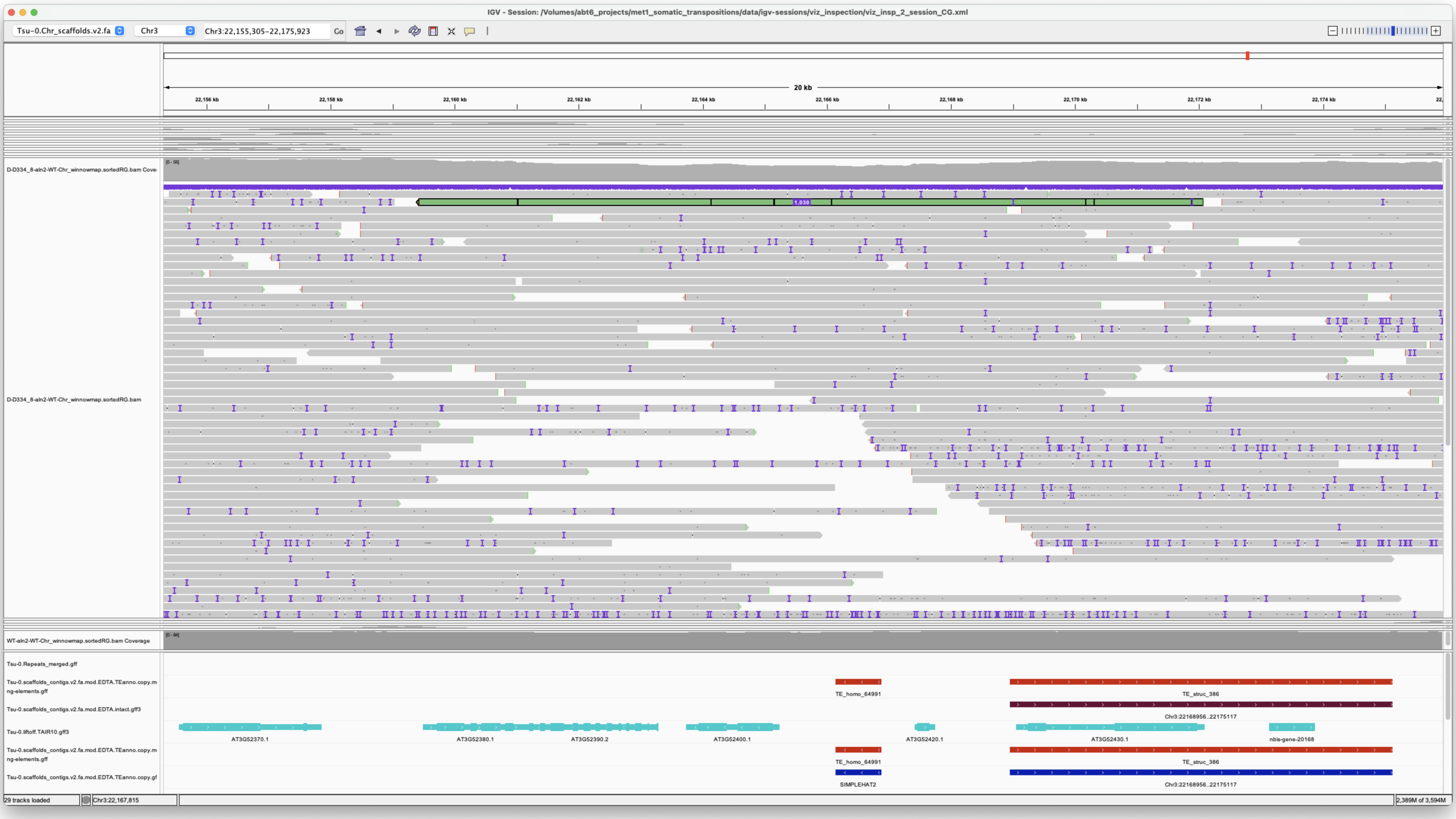


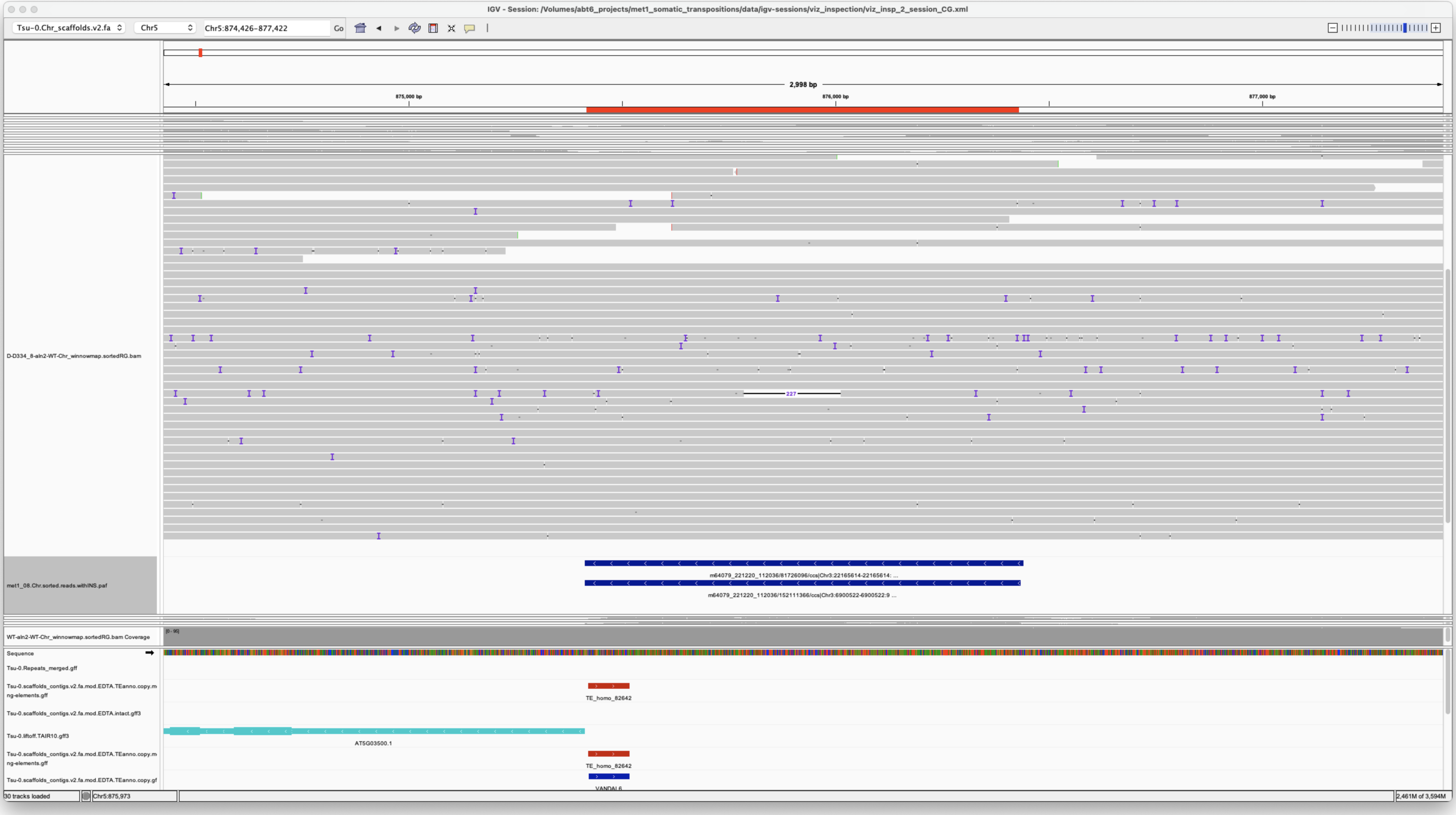


*Confirmed*

Chr5 18982271 18982271 m64079_221220_112036/93391083/ccs Chr5 19152825 19160826 Chr5|19152829|19160825|ID=TE_homo_95640;Name=VANDAL21;classification=DNA/Mutator;sequence_ontology=SO:0002280;identity=0.976;method=homology;ID=TE_homo_98501;sequence_ontology=SO:0002280|ID=TE_homo_95641;Name=VANDAL21;classification=DNA/Mutator;sequence_ontology=SO:0002280;identity=0.966;method=homology;ID=TE_homo_98502;sequence_ontology=SO:0002280 met1_08

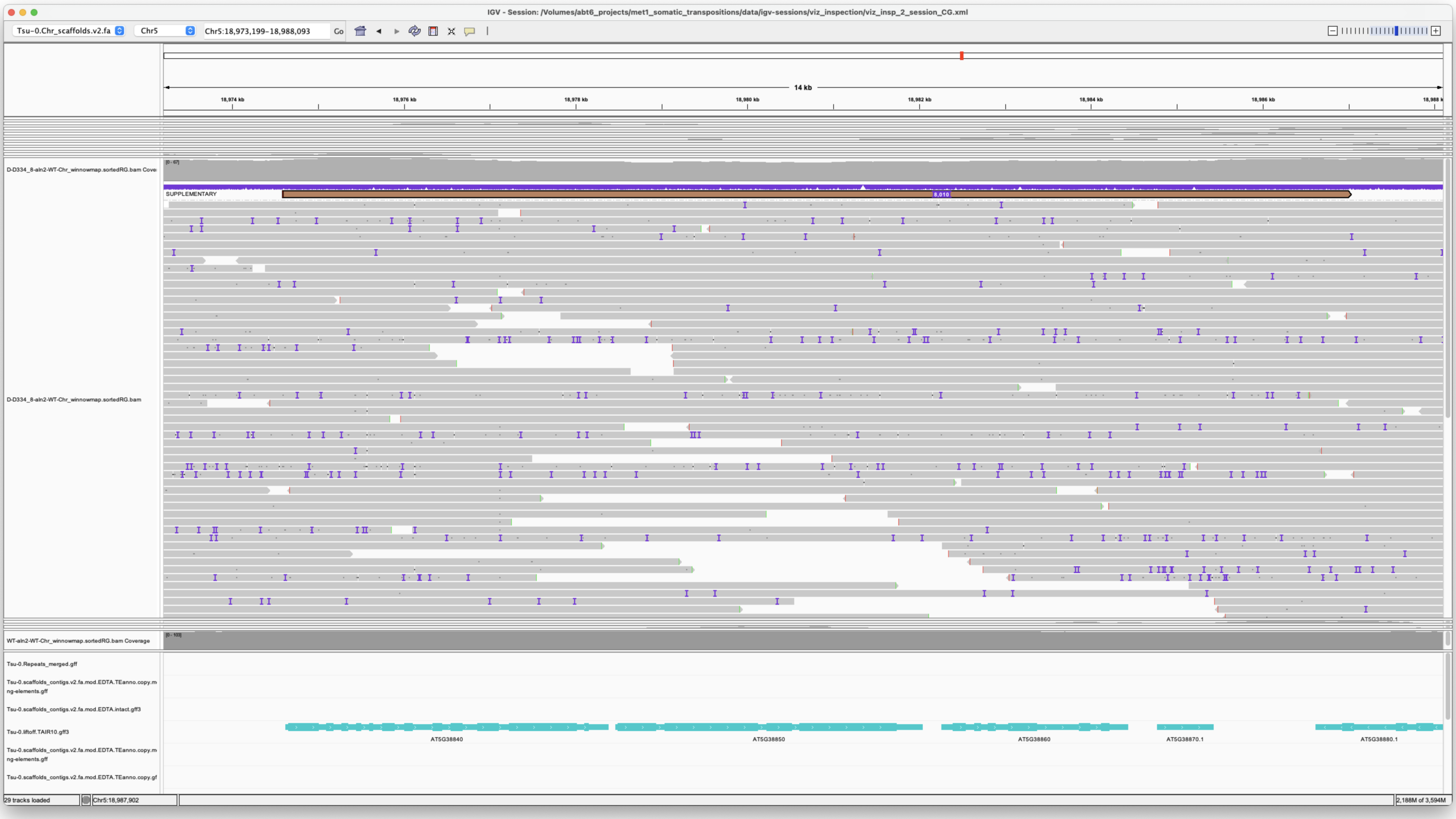


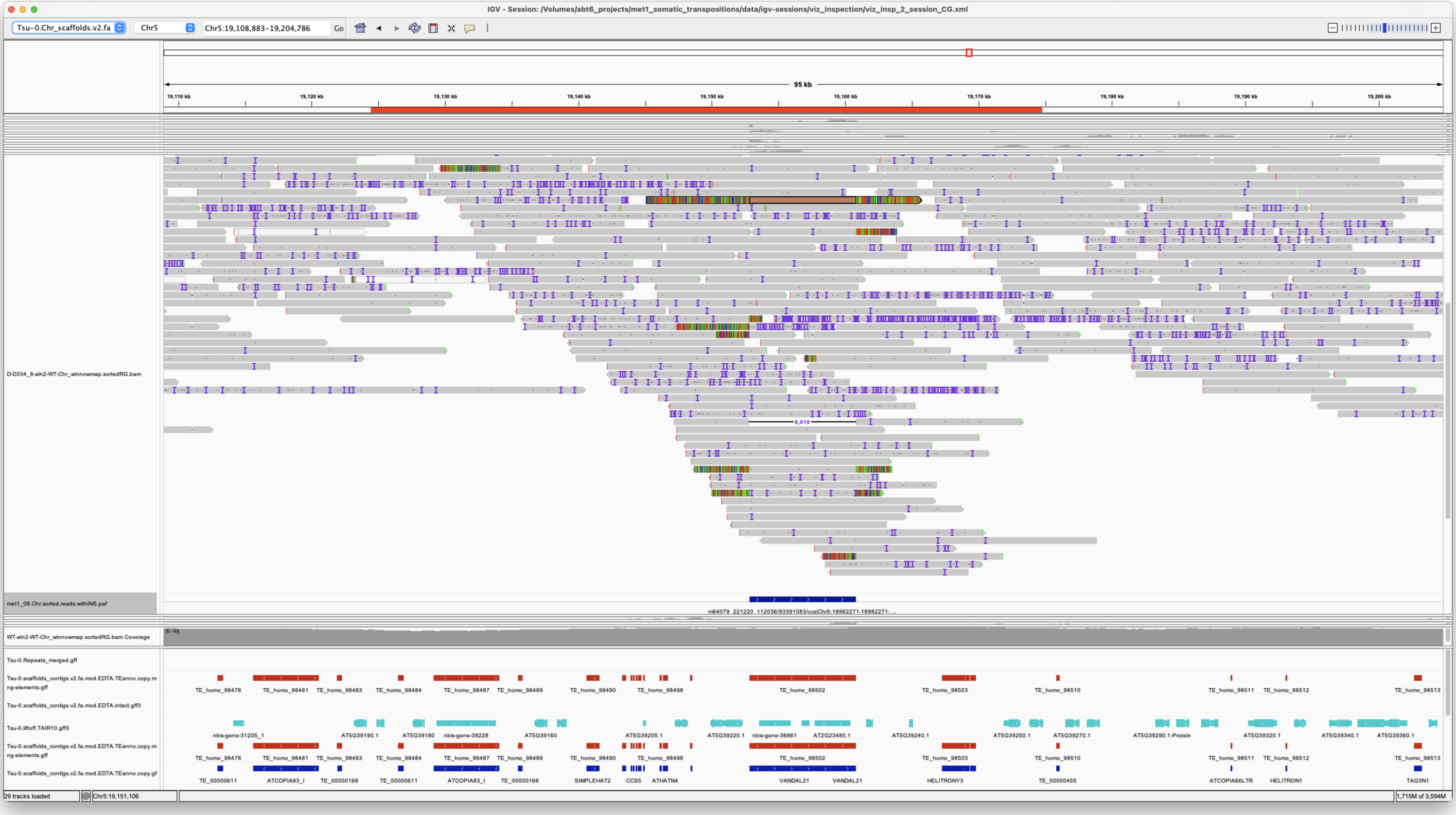


*Confirmed*

#### met1_09

> Chr1 7686553 7686553 m64079_221220_112036/113312127/ccs Chr5 875409 876434 Chr5|875414|876433|ID=TE_MANUAL_02;Name=PutativePackTypeCACTAMuDR;classification=DNA/DTC;sequence_ontology=MANUAL;identity=MANUAL;method=MANUAL;ID=TE_MANUAL_02;sequence_ontology=MANUAL met1_09

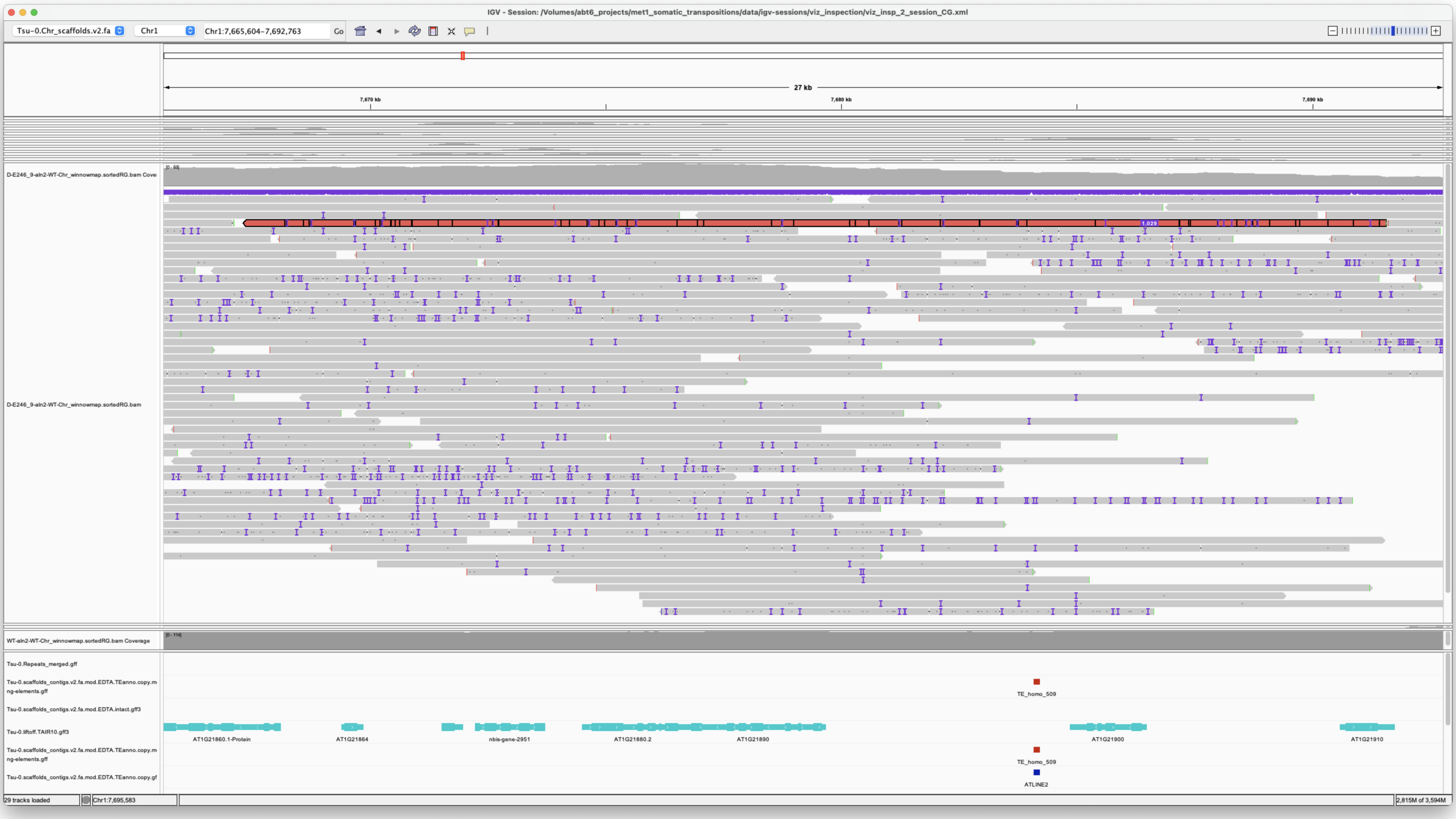


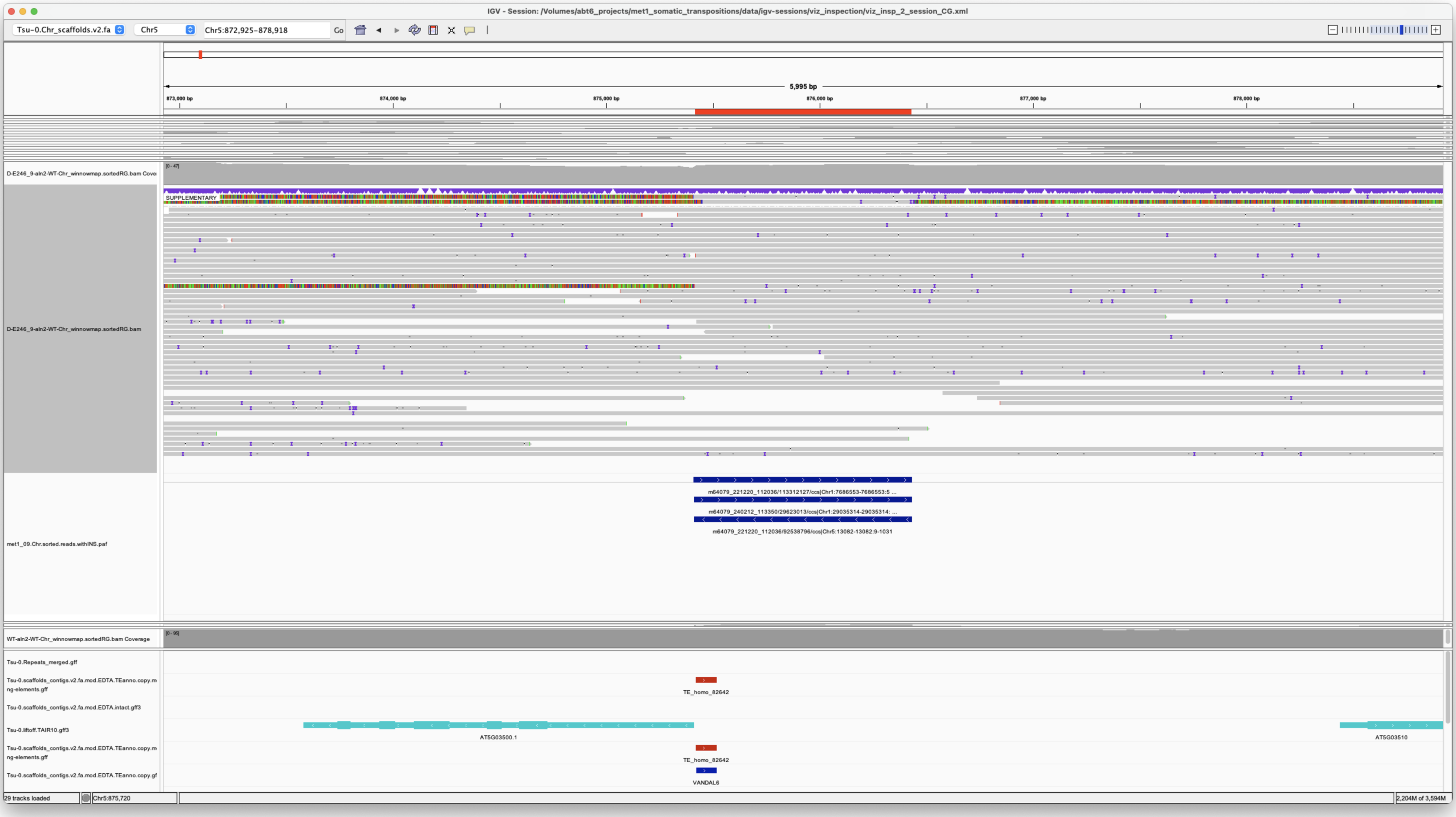


*Confirmed*

Chr1 28846148 28846150 m64079_221220_112036/125503791/ccs Chr4 9036240 9040519 Chr4|9037080|9037164|ID=TE_homo_78576;Name=TE_00000569;classification=Unknown;sequence_ontology=SO:0001050;identity=0.841;method=homology;ID=TE_homo_80707;sequence_ontology=SO:0001050**Chr4|9037221|9037332|ID=TE_homo_78577;Name=TE_00000569;classification=Unknown;sequence_ontology=SO:0001050;identity=0.909;method=homology;ID=TE_homo_80708;sequence_ontology=SO:0001050**Chr4|9037391|9037501|ID=TE_homo_78578;Name=TE_00000569;classification=Unknown;sequence_ontology=SO:0001050;identity=0.927;method=homology;ID=TE_homo_80709;sequence_ontology=SO:0001050**Chr4|9037502|9038015|ID=TE_homo_78579;Name=VANDAL21;classification=DNA/Mutator;sequence_ontology=SO:0002280;identity=0.705;method=homology;ID=TE_homo_80710;sequence_ontology=SO:0002280**Chr4|9038028|9039749|ID=TE_homo_78580;Name=VANDAL21;classification=DNA/MULE-MuDR;sequence_ontology=SO:0002280;identity=0.795;method=homology;ID=TE_homo_80711;sequence_ontology=SO:0002280**Chr4|9039779|9040118|ID=TE_homo_78581;Name=VANDAL21;classification=DNA/MULE-MuDR;sequence_ontology=SO:0002280;identity=0.735;method=homology;ID=TE_homo_80712;sequence_ontology=SO:0002280**Chr4|9040114|9040303|ID=TE_homo_78582;Name=TE_00000579;classification=Unknown;sequence_ontology=SO:0001050;identity=0.937;method=homology;ID=TE_homo_80713;sequence_ontology=SO:0001050 met1_09

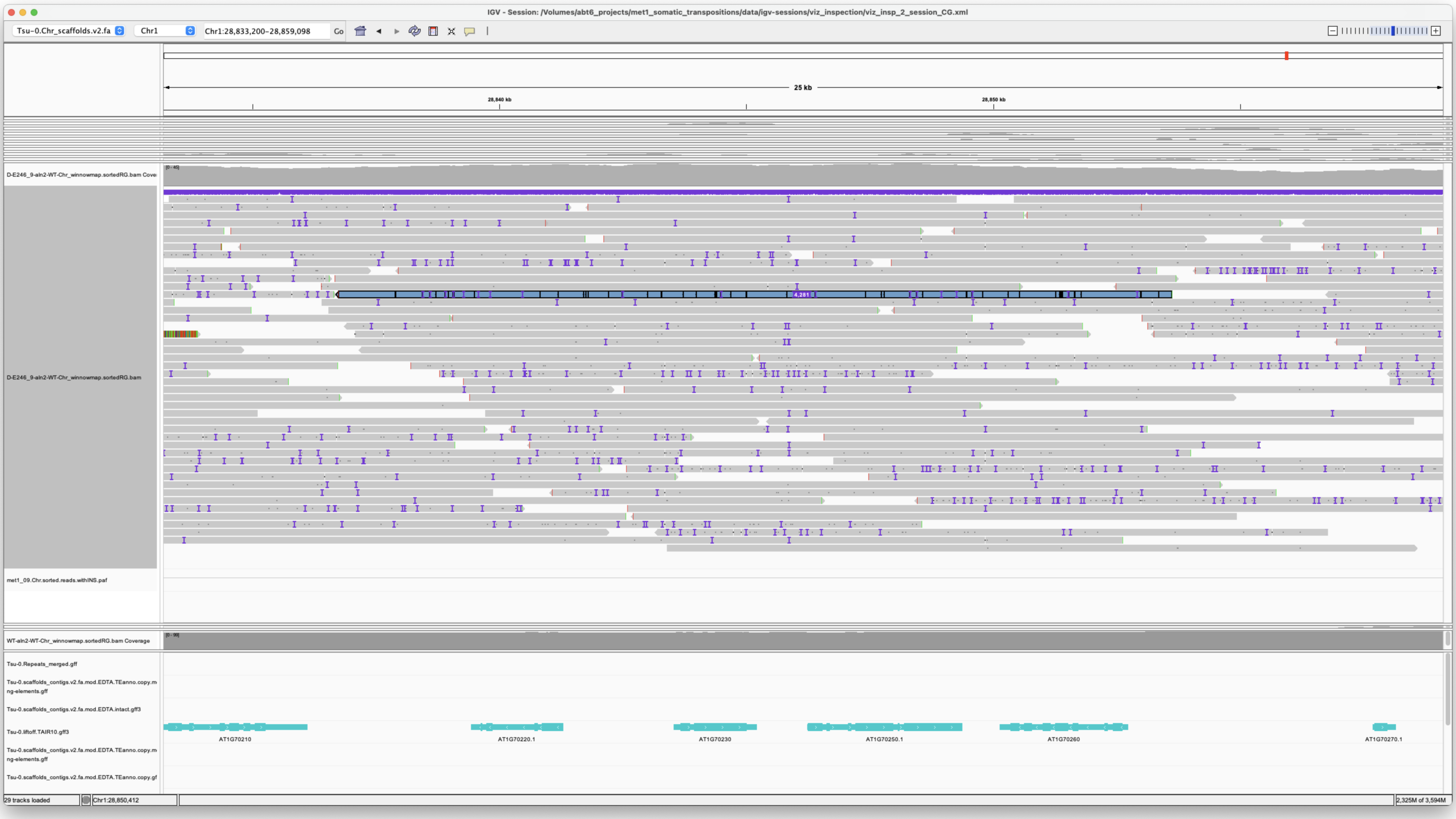


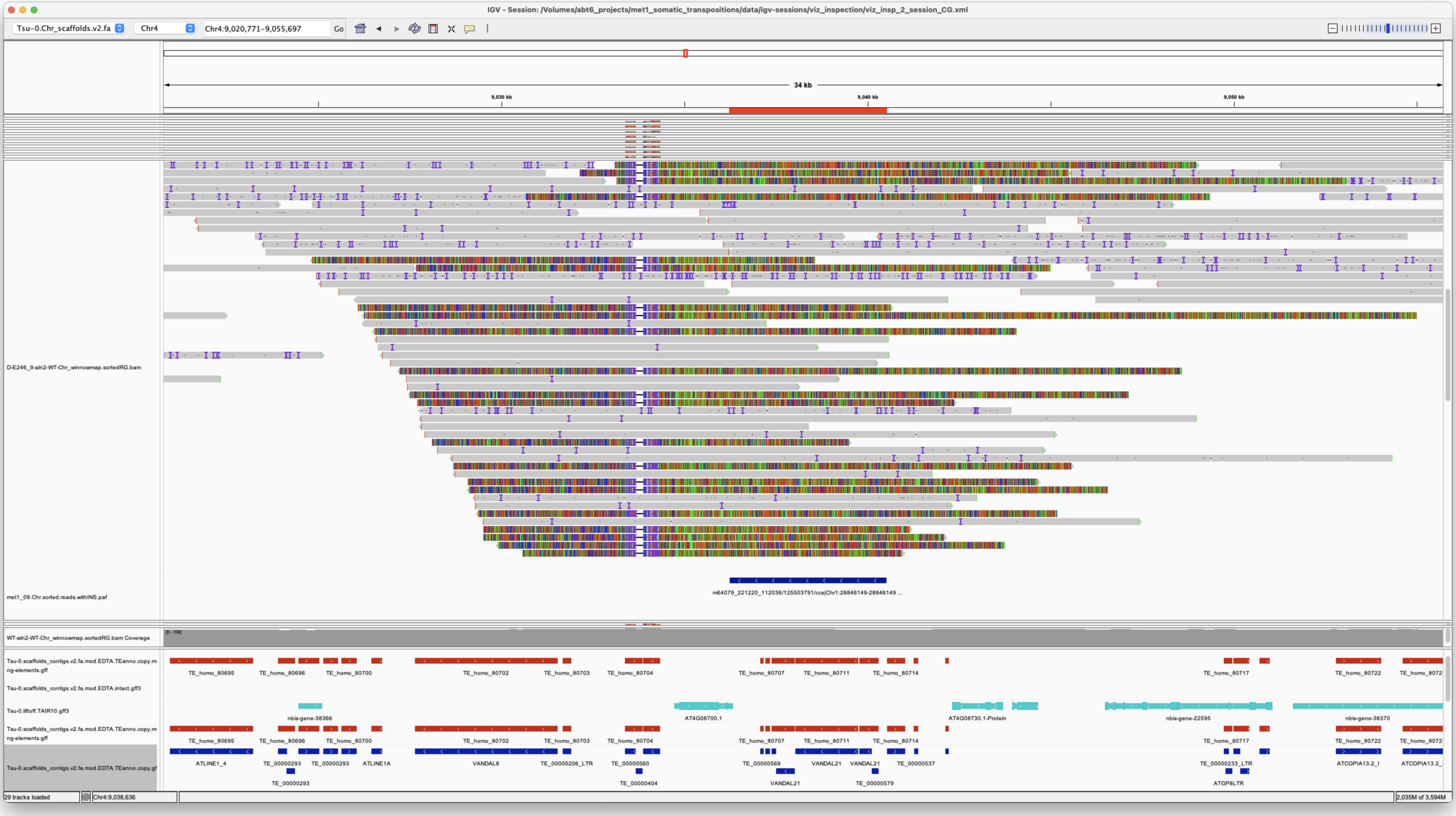


Presence of a difficult region nearby makes difficult to identify transpositions at this locus.
Also this VANDAL21 seems to be incorrectly annotated.

```
!!!To be manually corrected in main annotation.
```

But the presence of an insertion in central configuration is good evidence of mobilization of this element.
Also dotplot shows TSD

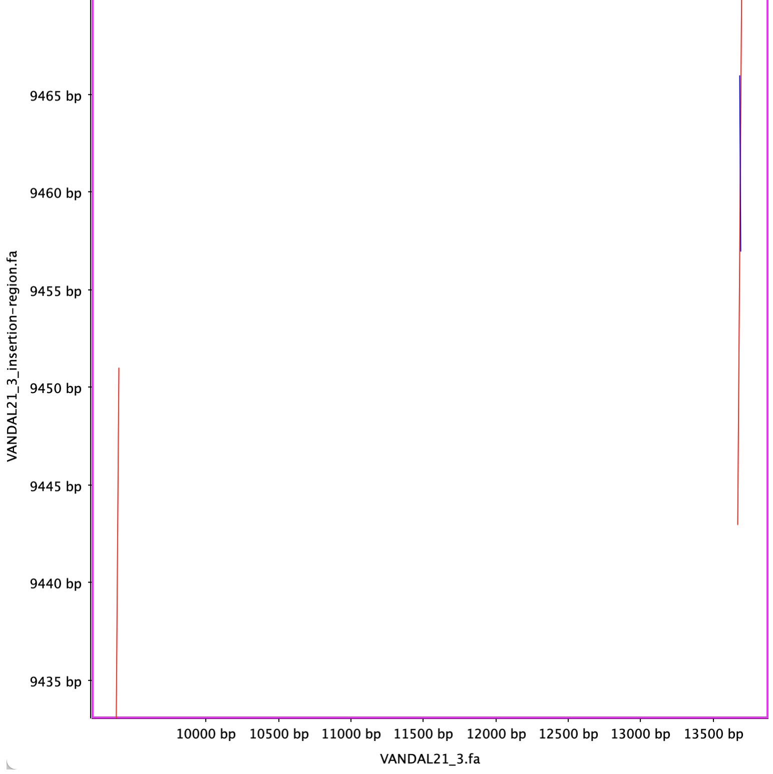


*Confirmed*

Chr1 29035314 29035314 m64079_240212_113350/29623013/ccs Chr5 875413 876434 Chr5|875414|876433|ID=TE_MANUAL_02;Name=PutativePackTypeCACTAMuDR;classification=DNA/DTC;sequence_ontology=MANUAL;identity=MANUAL;method=MANUAL;ID=TE_MANUAL_02;sequence_ontology=MANUAL met1_09

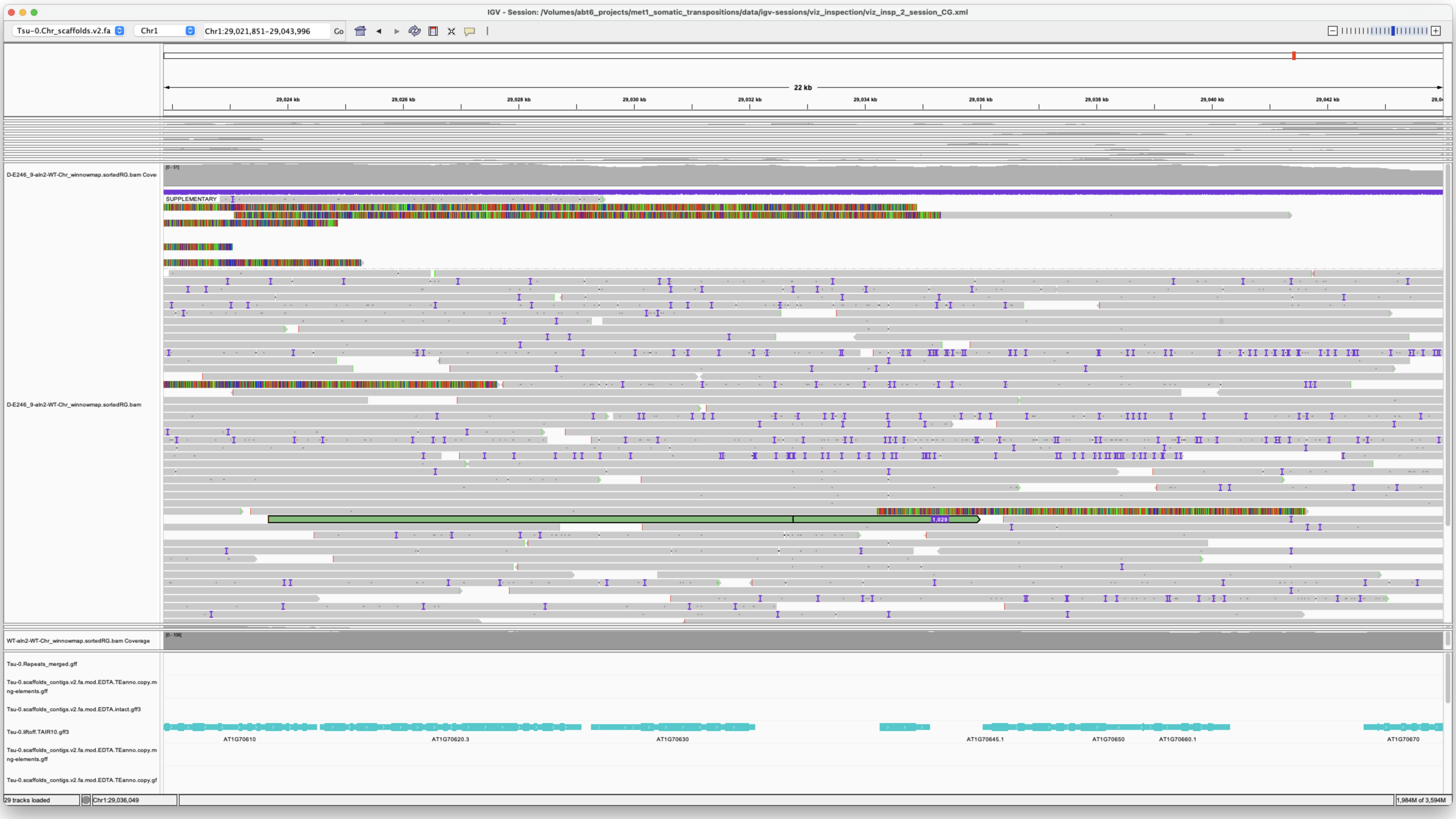

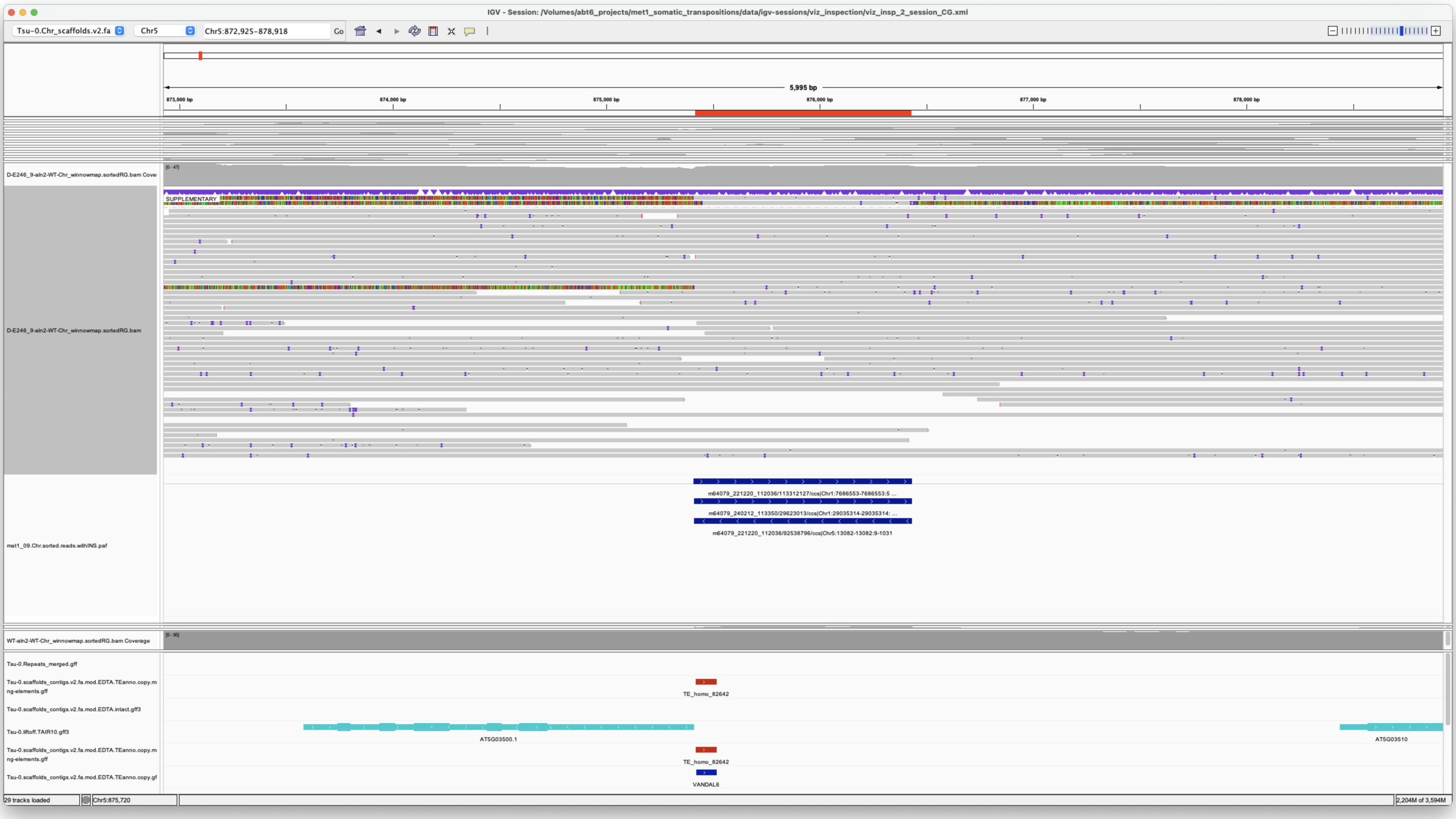

*Confirmed*

Chr5 13082 13082 m64079_221220_112036/92538796/ccs Chr5 875413 876434 Chr5|875414|876433|ID=TE_MANUAL_02;Name=PutativePackTypeCACTAMuDR;classification=DNA/DTC;sequence_ontology=MANUAL;identity=MANUAL;method=MANUAL;ID=TE_MANUAL_02;sequence_ontology=MANUAL met1_09

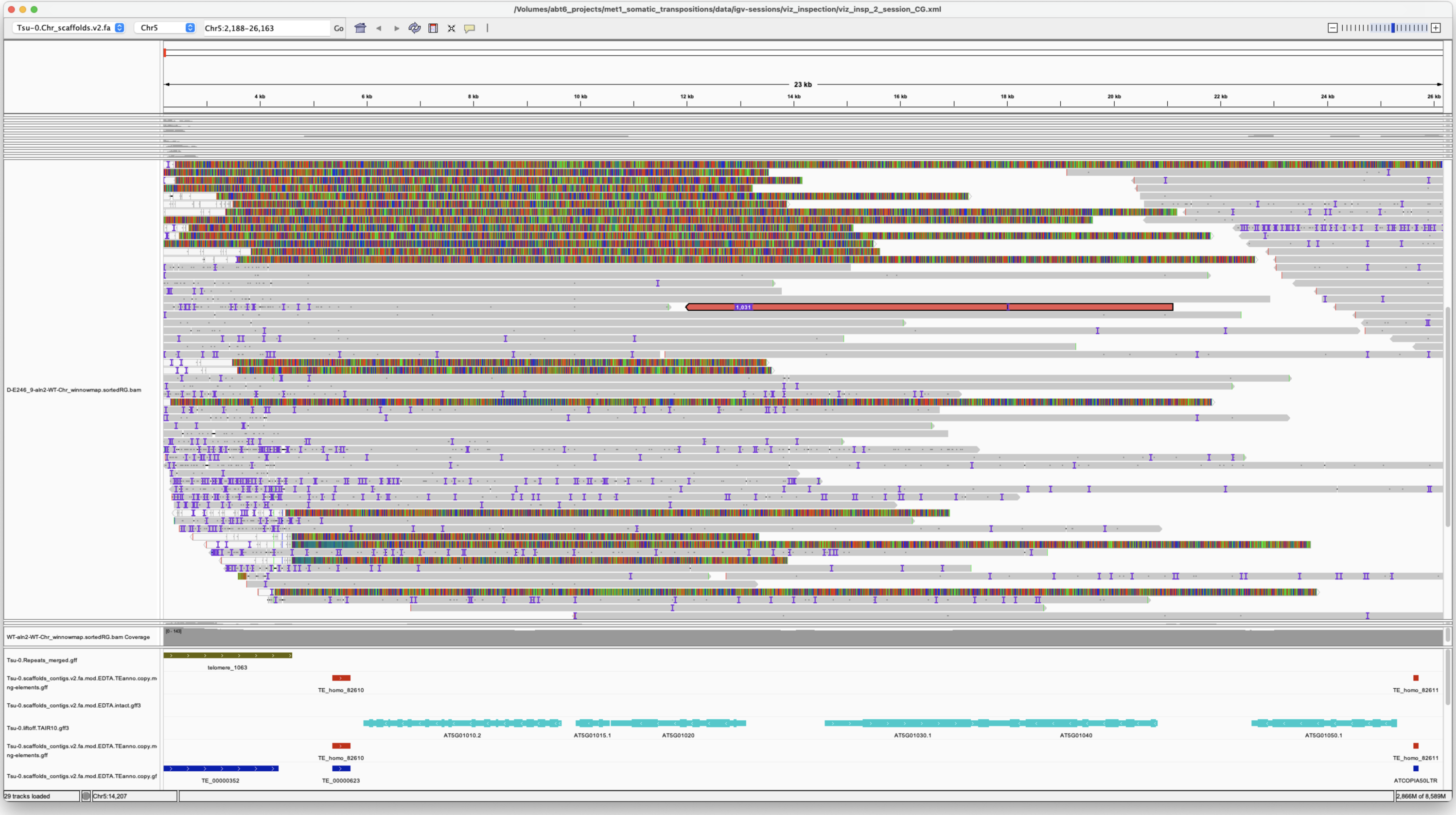

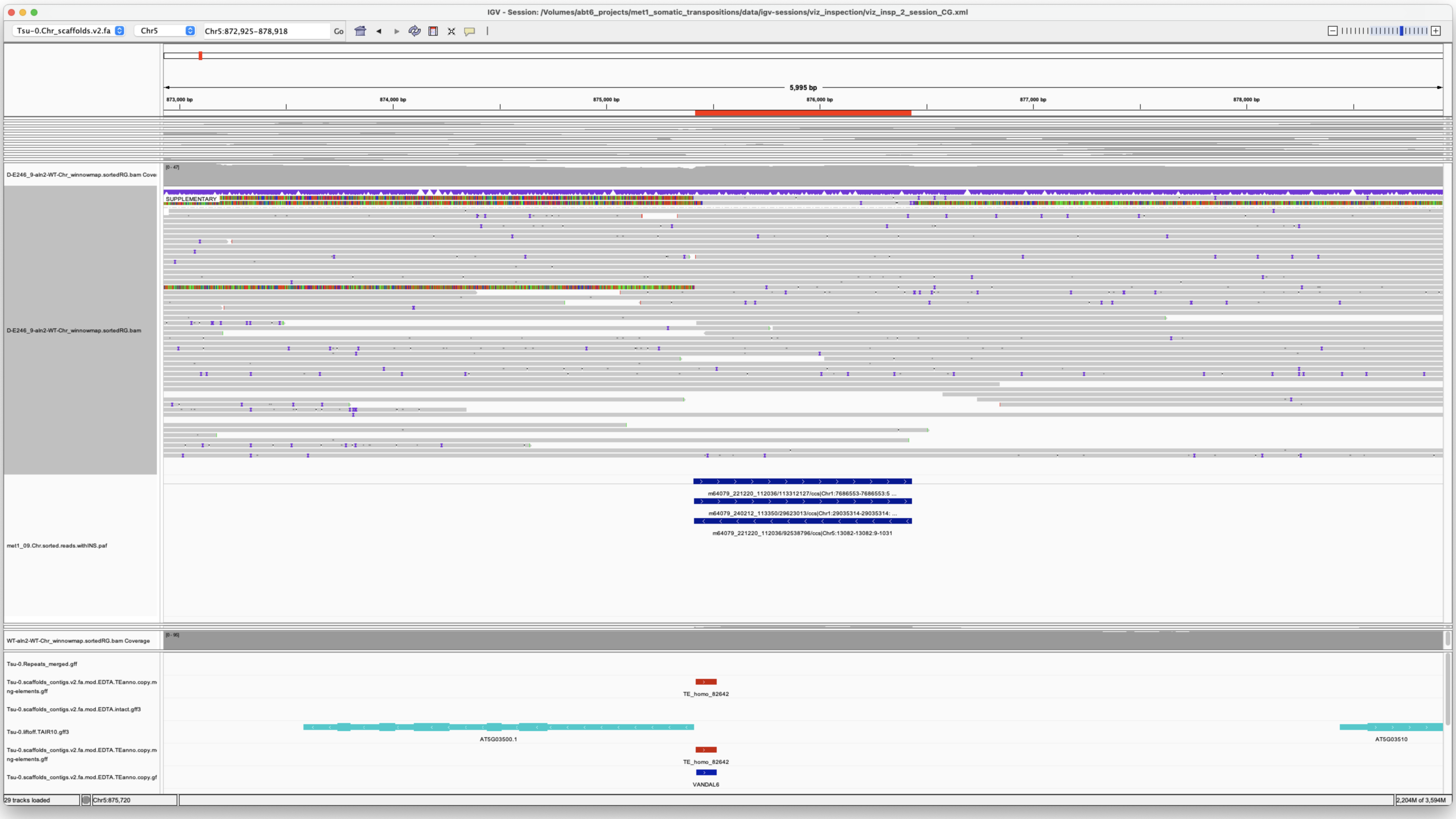


Very telomeric insertion

*Confirmed*

#### met1_10

> Chr1 6485187 6485187 m64079_240212_113350/152699302/ccs Chr5 875413 876434 Chr5|875414|876433|ID=TE_MANUAL_02;Name=PutativePackTypeCACTAMuDR;classification=DNA/DTC;sequence_ontology=MANUAL;identity=MANUAL;method=MANUAL;ID=TE_MANUAL_02;sequence_ontology=MANUAL met1_10

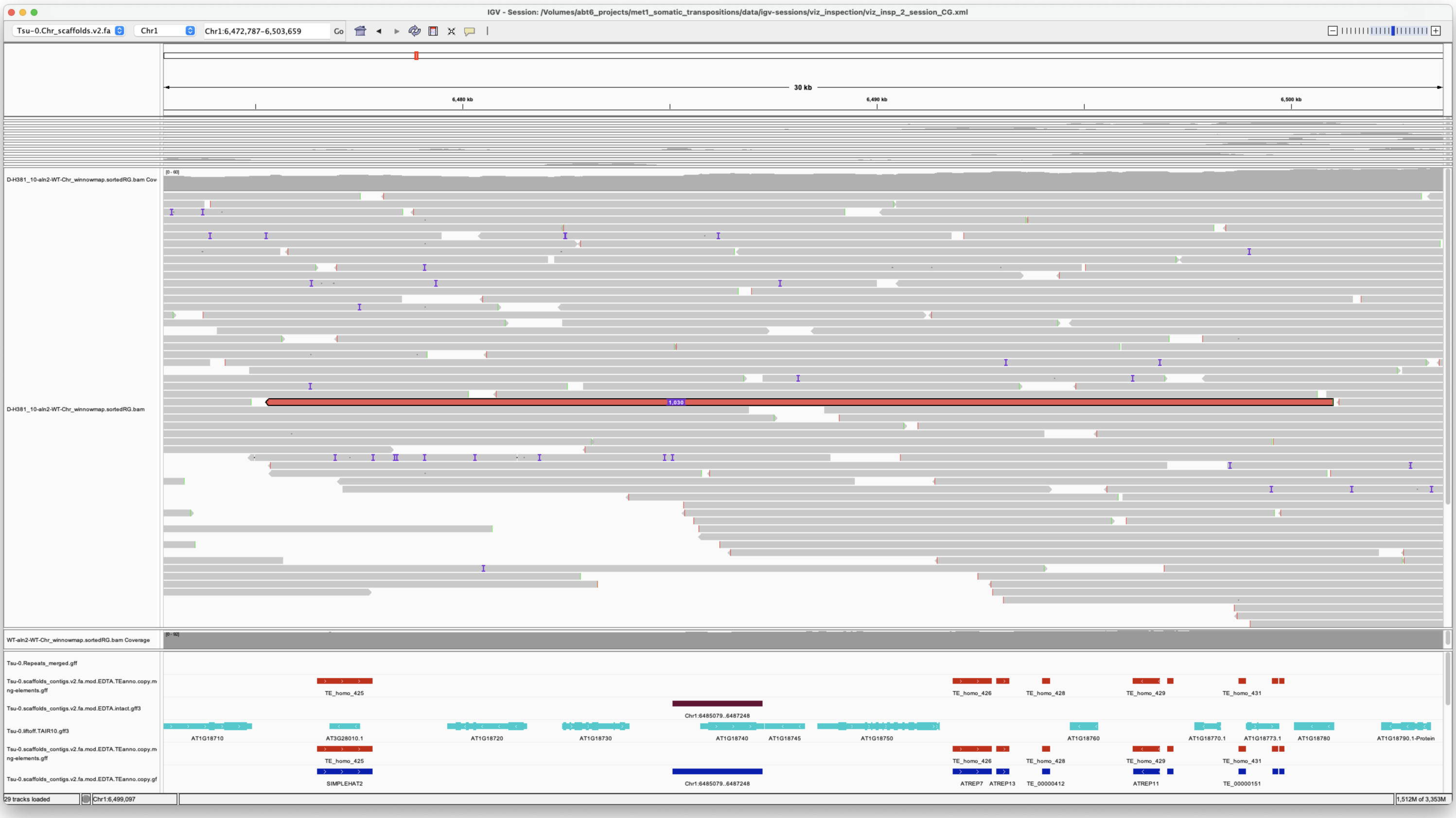

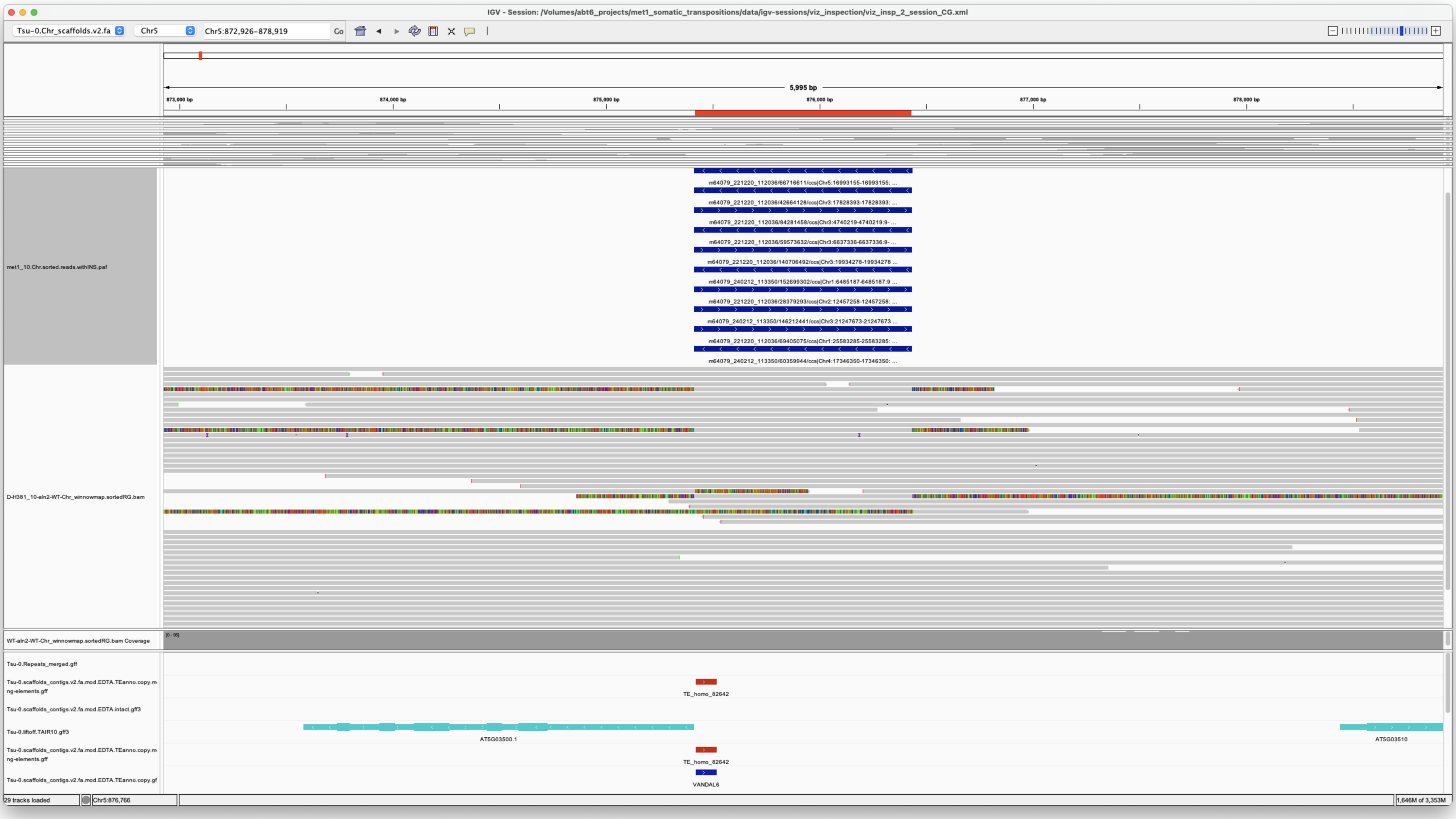


*Confirmed*

> Chr1 9954546 9954546 m64079_221220_112036/69664872/ccs Chr5 21419693 21425021
> Chr5|21419693|21425021|ID=LTRRT_299;Name=ATCOPIA93.2_Evade;Classification=LTR/Copia;Sequence_ontology=SO:0002264;ltr_identity=1.0000;Method=structural;motif=TACA;tsd=GGACA met1_10

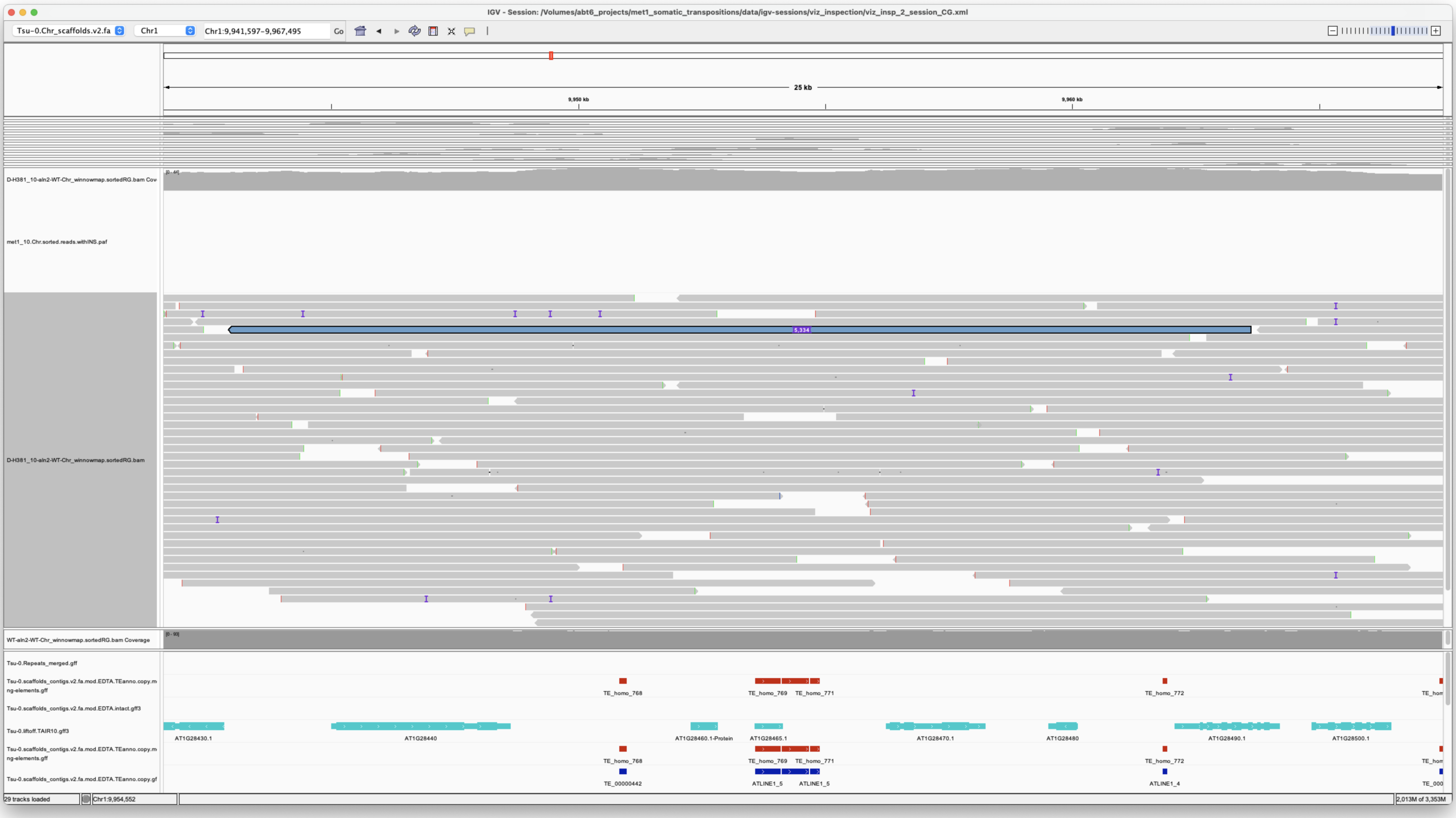

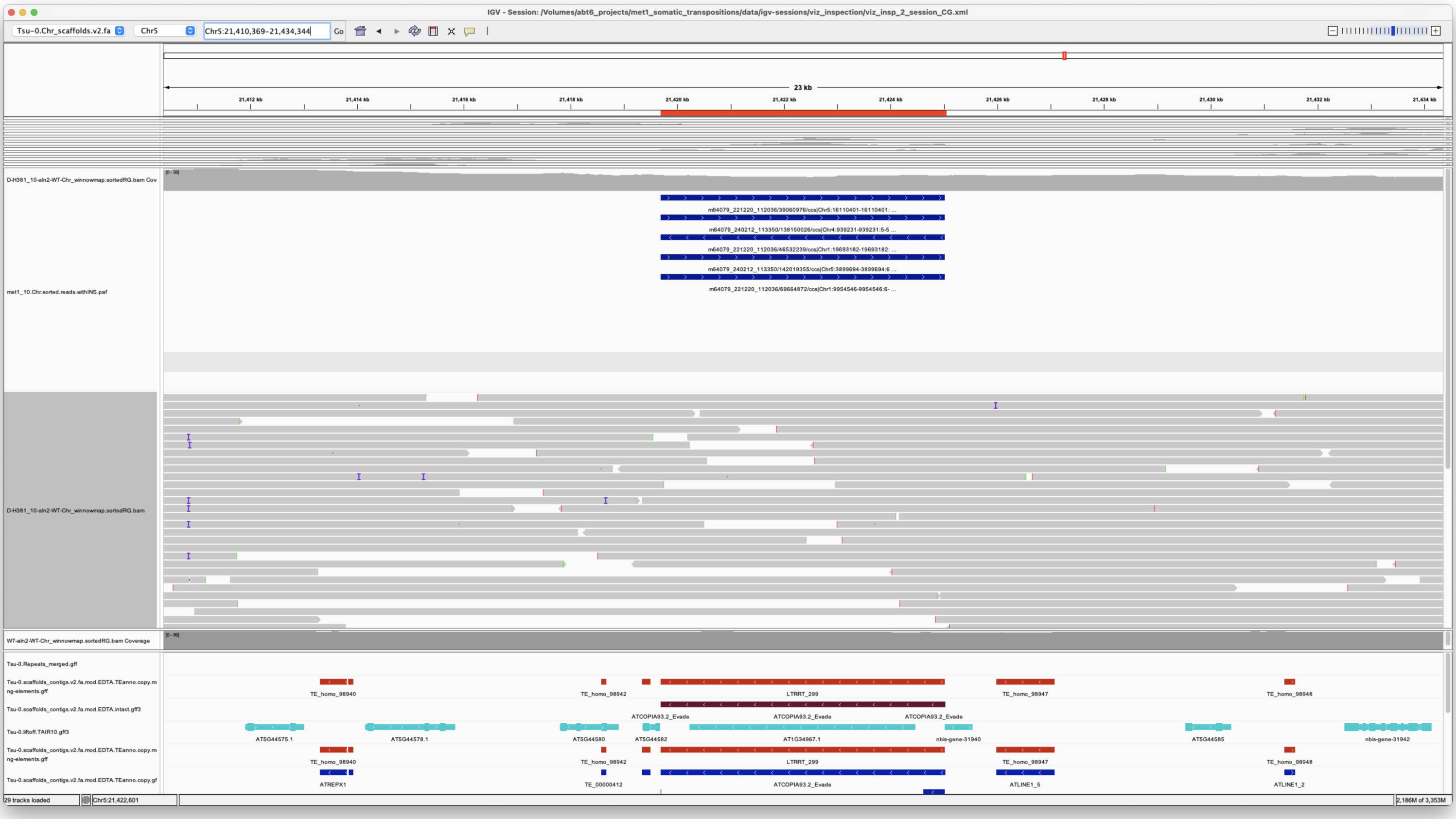


*Confirmed*

---

> Chr1 19693182 19693182 m64079_221220_112036/46532239/ccs Chr5 21419693 21425022
> Chr5|21419693|21425021|ID=LTRRT_299;Name=ATCOPIA93.2_Evade;Classification=LTR/Copia;Sequence_ontology=SO:0002264;ltr_identity=1.0000;Method=structural;motif=TACA;tsd=GGACA met1_10

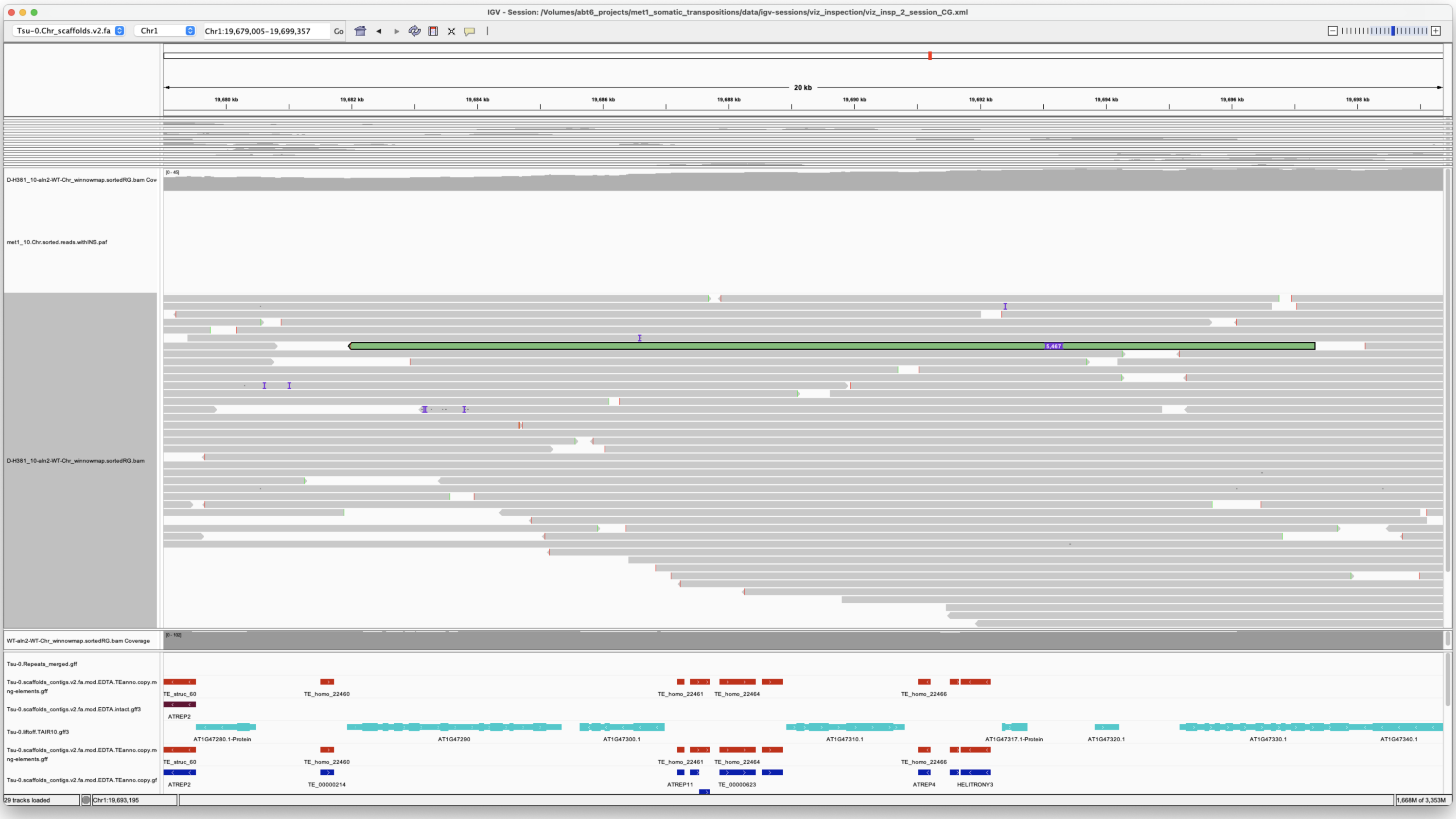


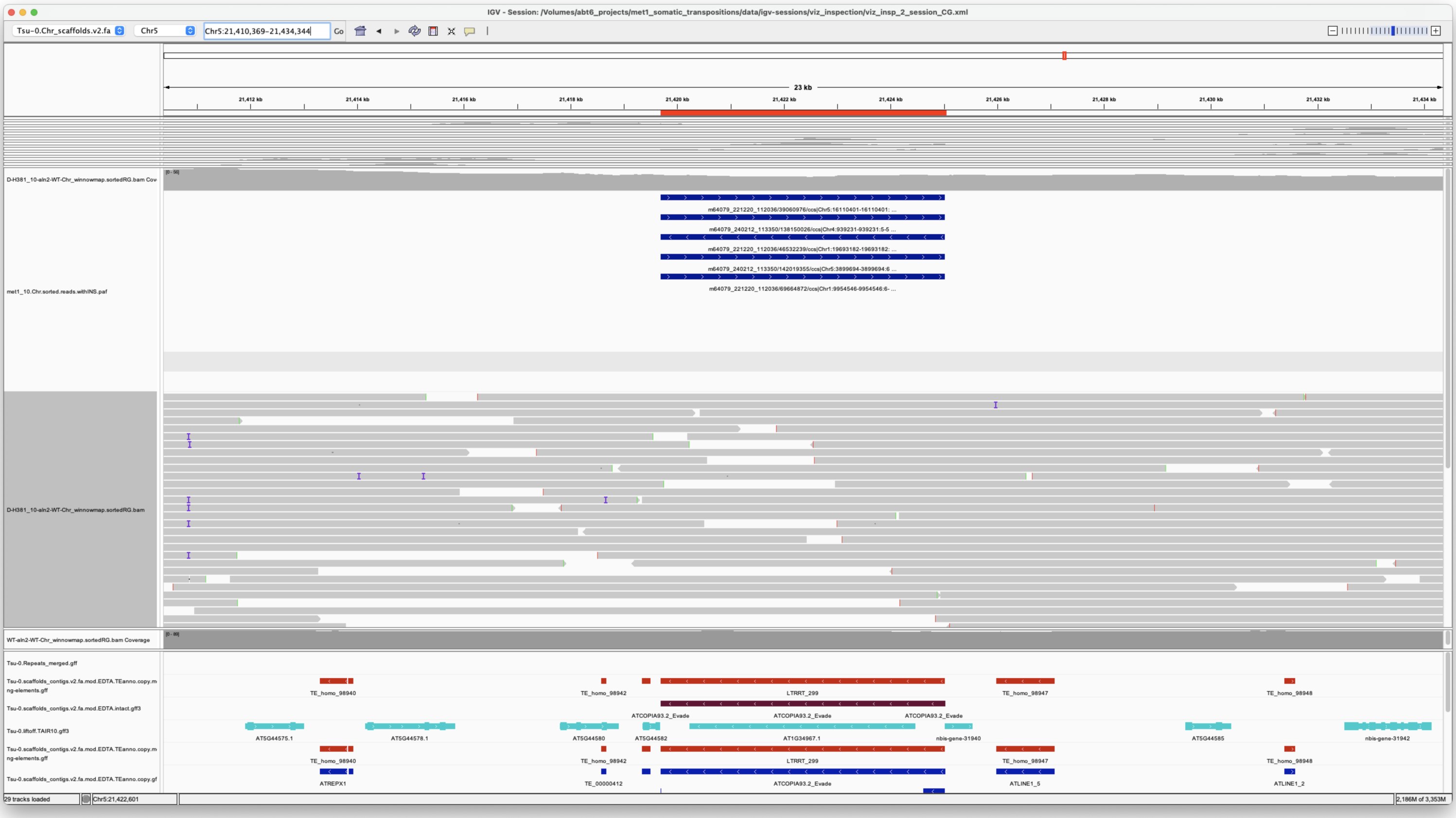


*Confirmed*

---

> Chr1 25583285 25583285 m64079_221220_112036/69405075/ccs Chr5 875413 876433 Chr5|875414|876433|ID=TE_MANUAL_02;Name=PutativePackTypeCACTAMuDR;classification=DNA/DTC;sequence_ontology=MANUAL;identity=MANUAL;method=MANUAL;ID=TE_MANUAL_02;sequence_ontology=MANUAL met1_10

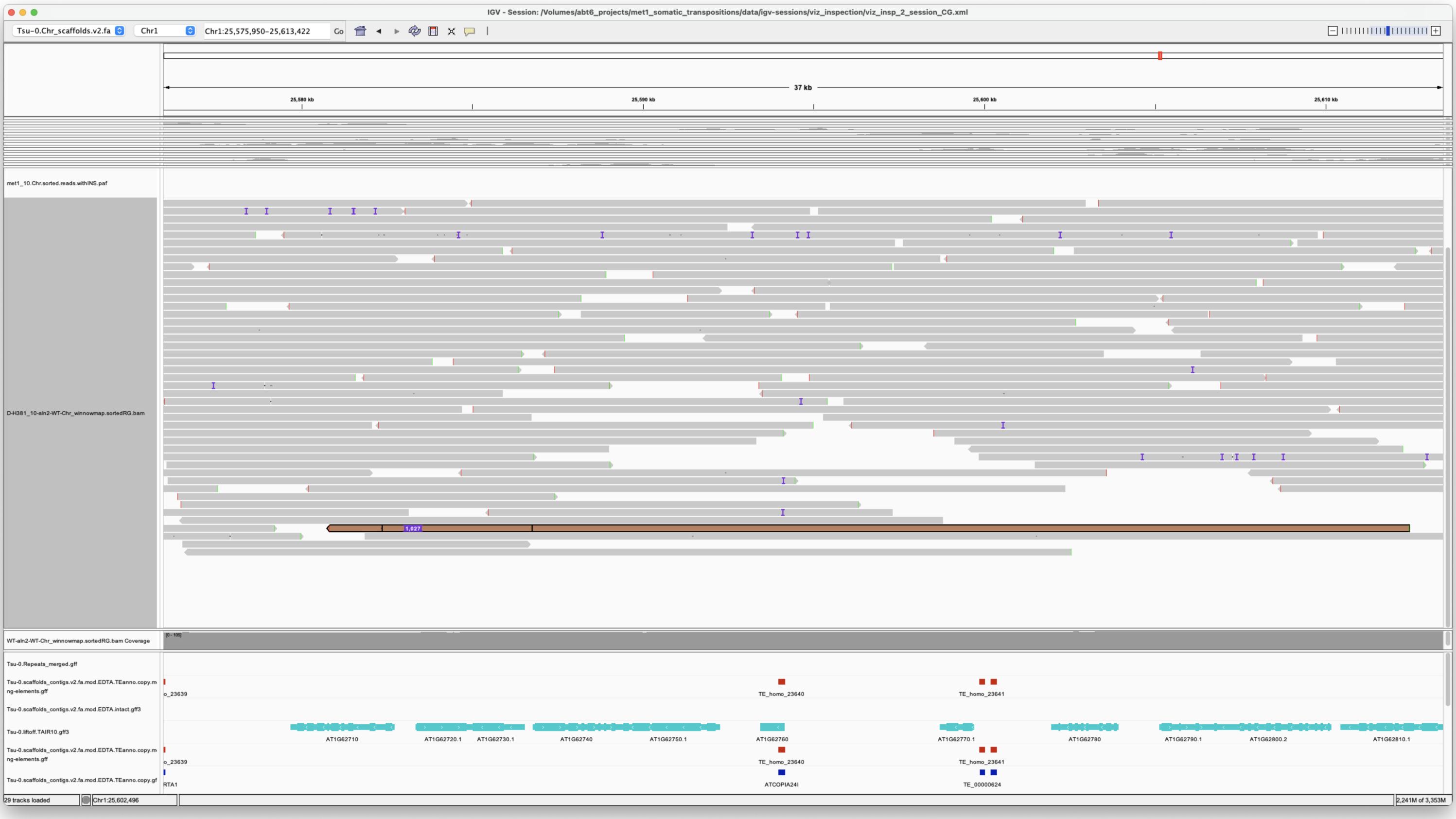

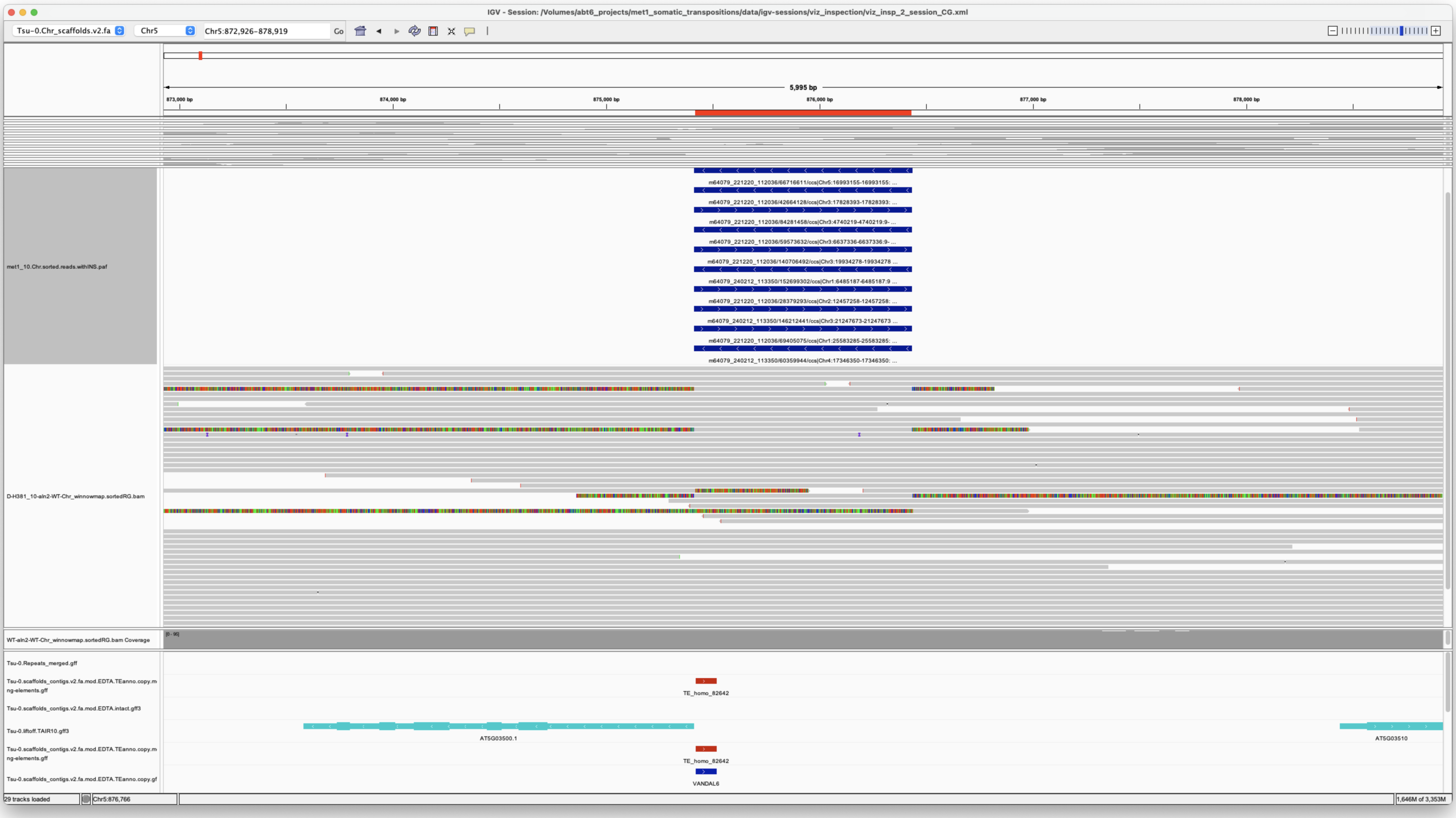

*Confirmed*

Chr2 12457258 12457258 m64079_221220_112036/28379293/ccs Chr5 875413 876433 Chr5|875414|876433|ID=TE_MANUAL_02;Name=PutativePackTypeCACTAMuDR;classification=DNA/DTC;sequence_ontology=MANUAL;identity=MANUAL;method=MANUAL;ID=TE_MANUAL_02;sequence_ontology=MANUAL met1_10

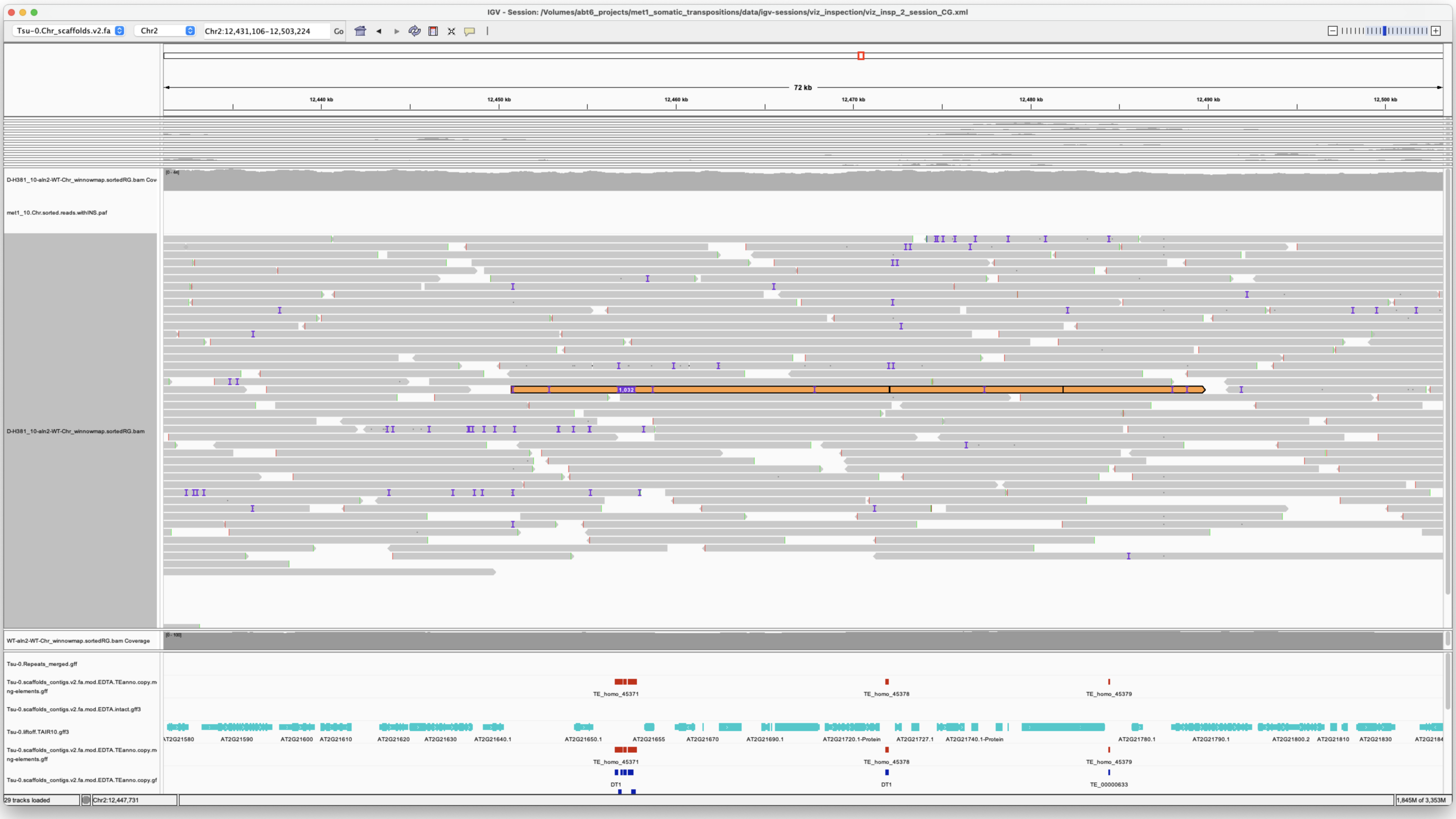

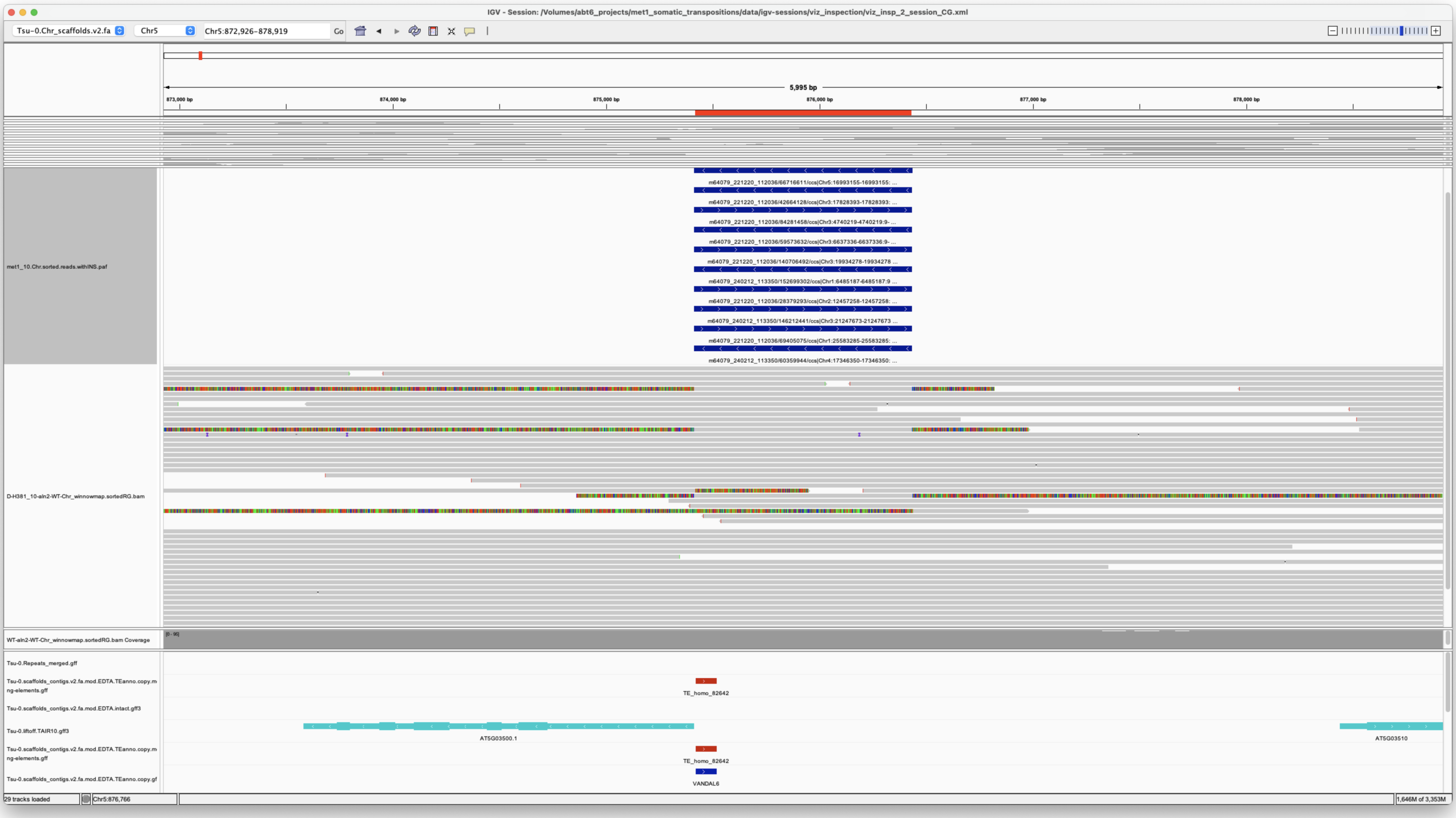

*Confirmed*

---

> Chr2 18394095 18394095 m64079_240212_113350/171707048/ccs Chr3 16344522 16352498 Chr3|16344522|16352496|ID=TE_homo_60420;Name=VANDAL6;classification=DNA/Mutator;sequence_ontology=SO:0002280;identity=0.969;method=homology;ID=TE_homo_62001;sequence_ontology=SO:0002280 met1_10

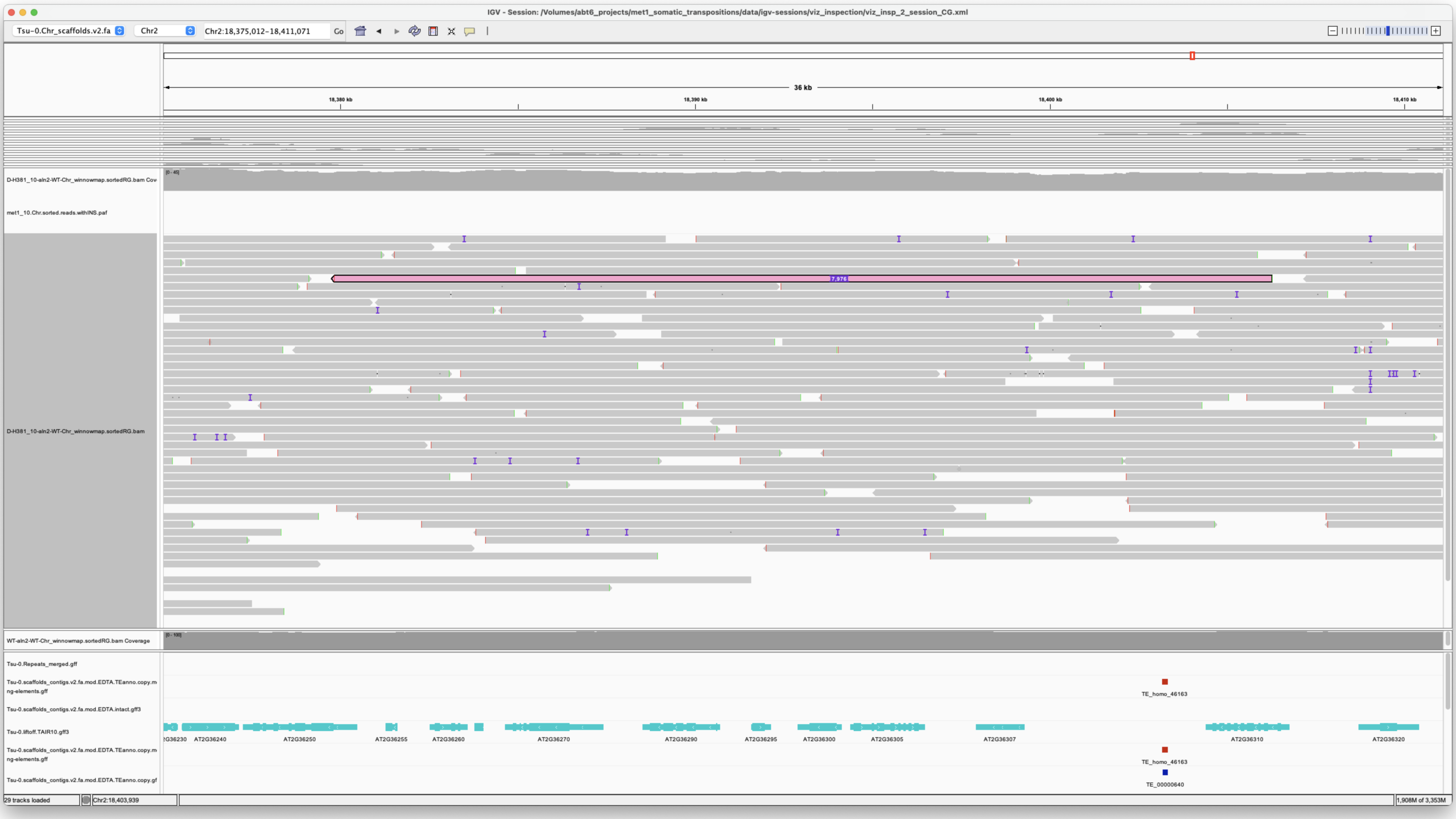


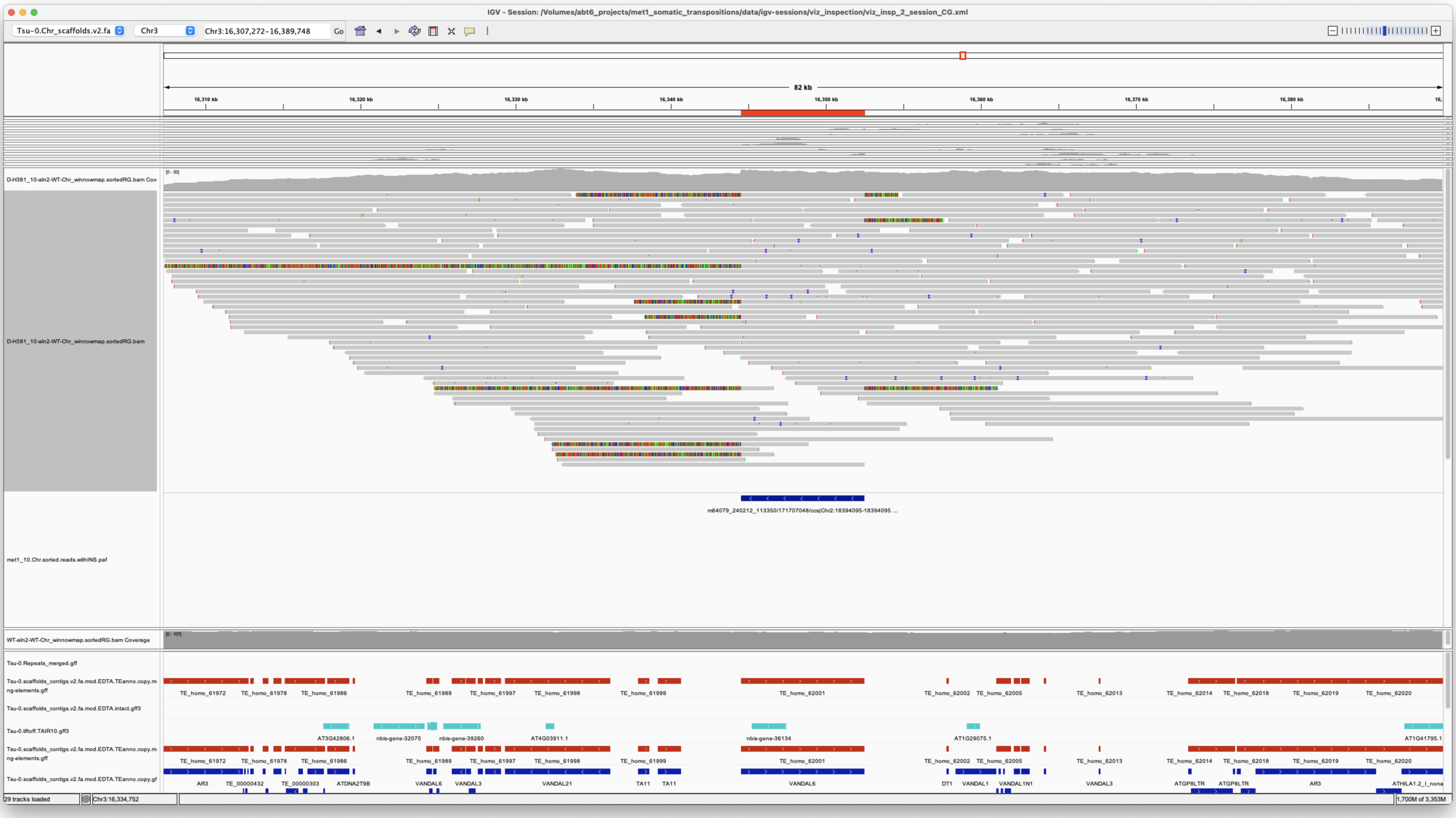


*Confirmed*

---

> Chr3 4740219 4740219 m64079_221220_112036/84281458/ccs Chr5 875413 876434 Chr5|875414|876433|ID=TE_MANUAL_02;Name=PutativePackTypeCACTAMuDR;classification=DNA/DTC;sequence_ontology=MANUAL;identity=MANUAL;method=MANUAL;ID=TE_MANUAL_02;sequence_ontology=MANUAL met1_10

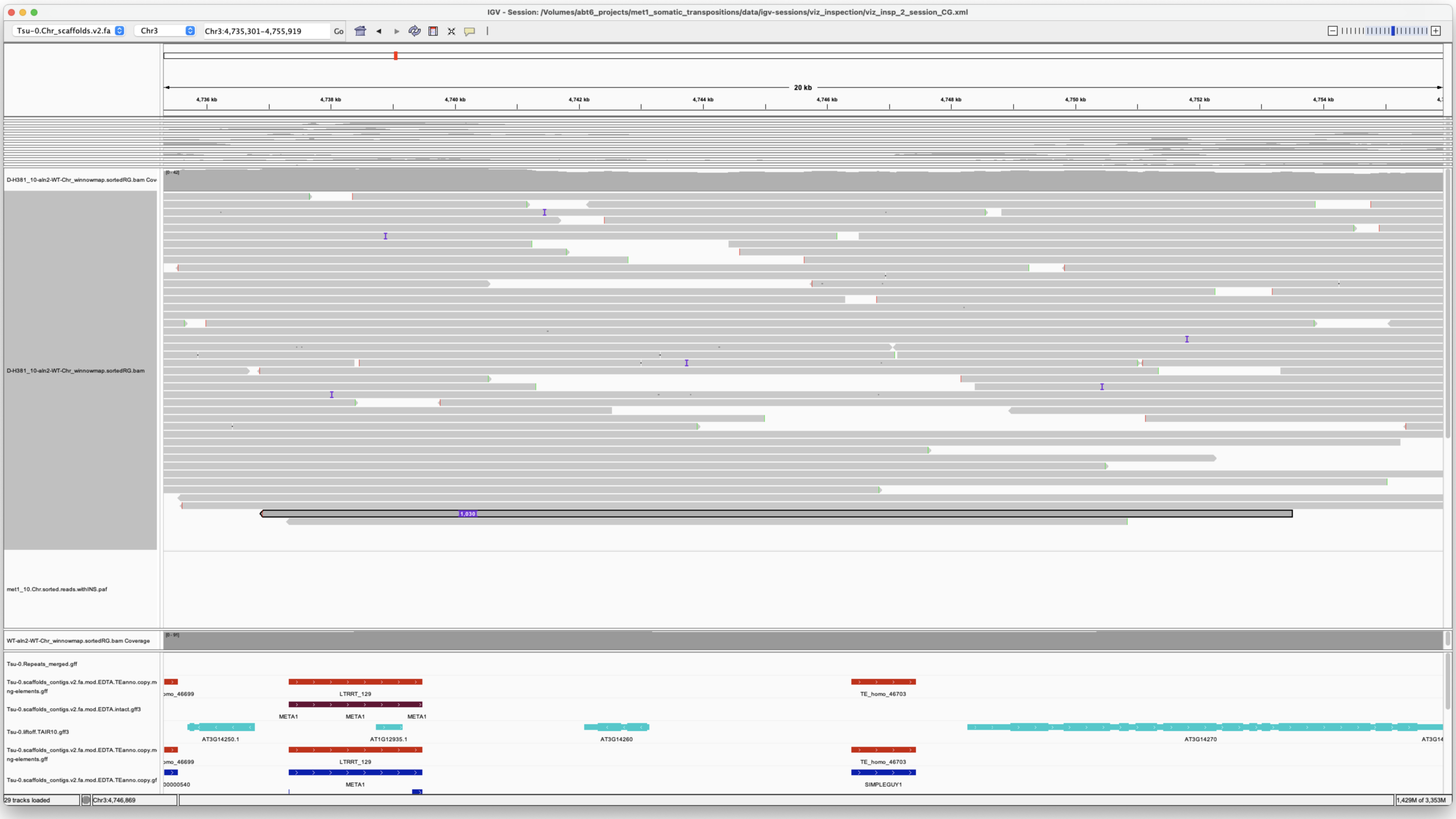


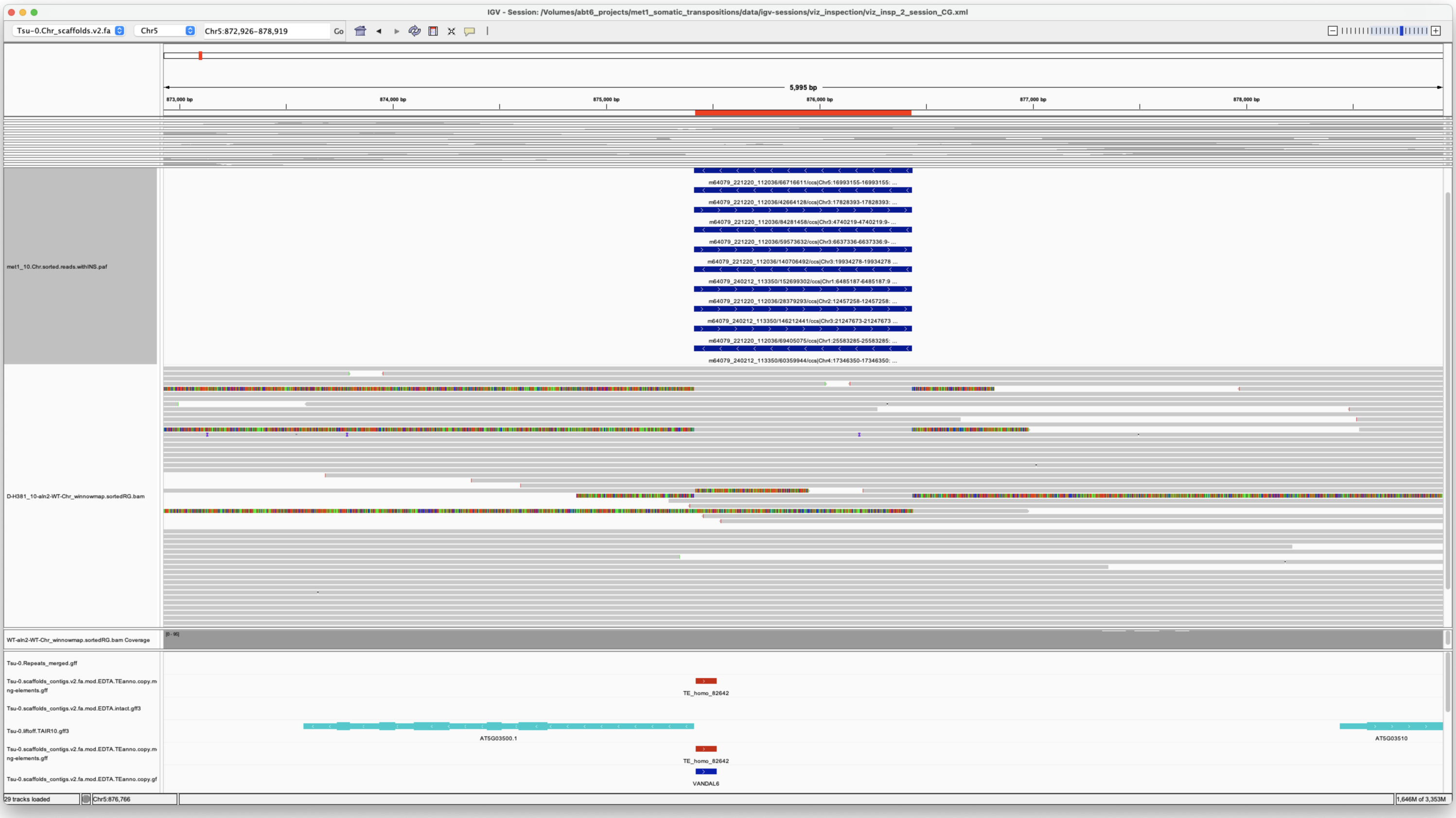


*Confirmed*

---

Chr3 6637336 6637336 m64079_221220_112036/59573632/ccs Chr5 875413 876434 Chr5|875414|876433|ID=TE_MANUAL_02;Name=PutativePackTypeCACTAMuDR;classification=DNA/DTC;sequence_ontology=MANUAL;identity=MANUAL;method=MANUAL;ID=TE_MANUAL_02;sequence_ontology=MANUAL met1_10

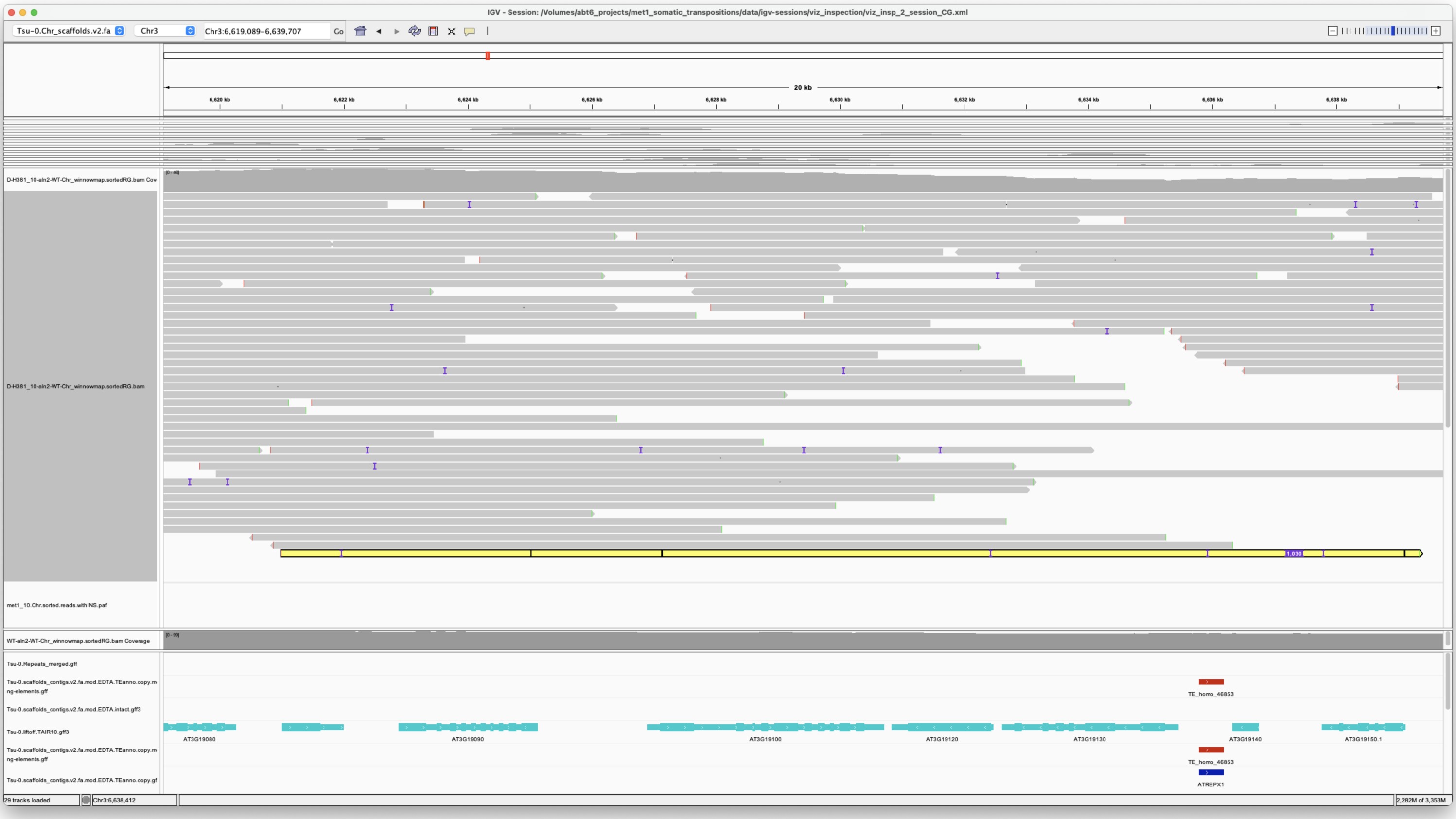

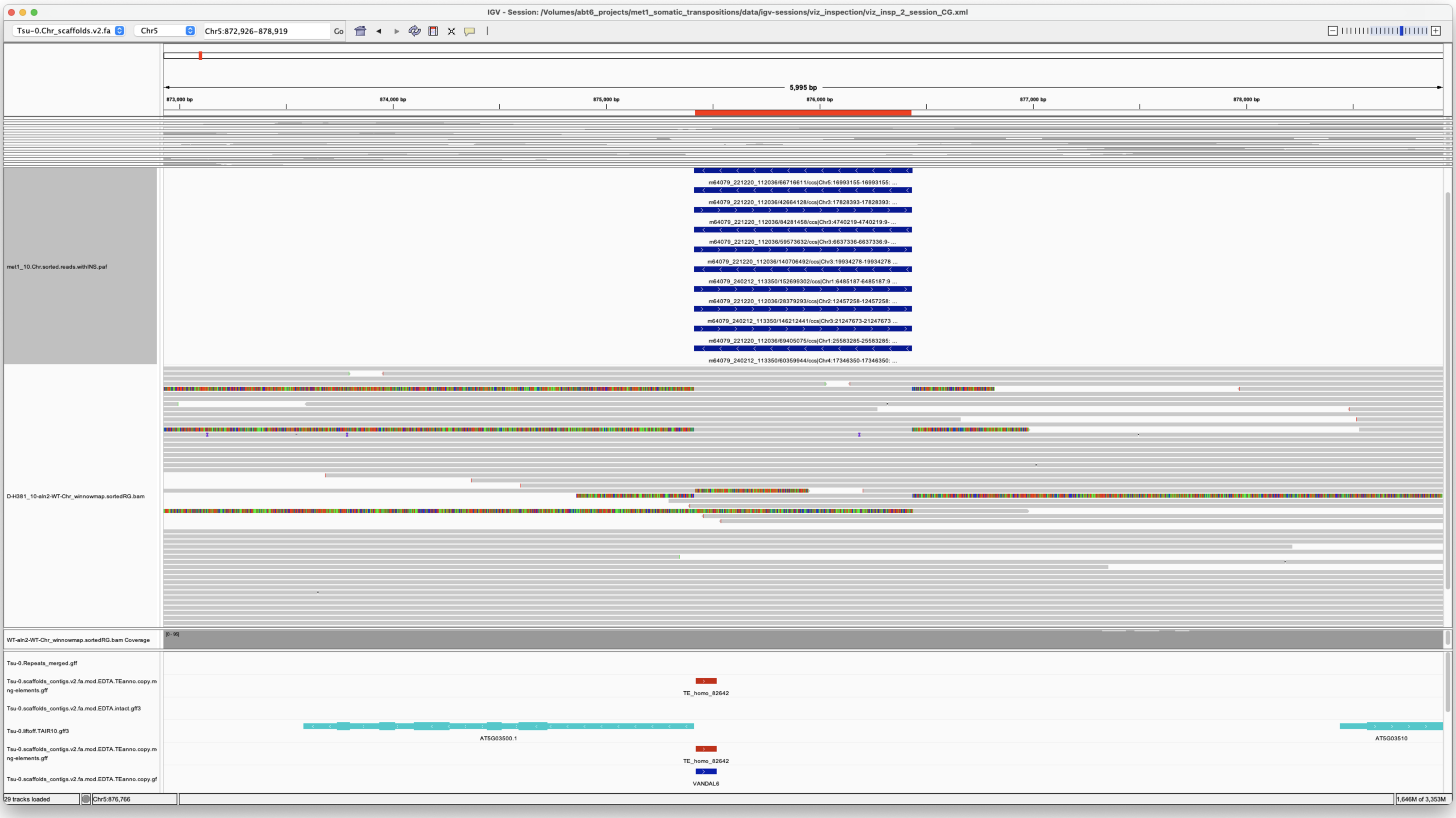

*Confirmed*

> Chr3 17828393 17828393 m64079_221220_112036/42664128/ccs Chr5 875413 876434 Chr5|875414|876433|ID=TE_MANUAL_02;Name=PutativePackTypeCACTAMuDR;classification=DNA/DTC;sequence_ontology=MANUAL;identity=MANUAL;method=MANUAL;ID=TE_MANUAL_02;sequence_ontology=MANUAL met1_10

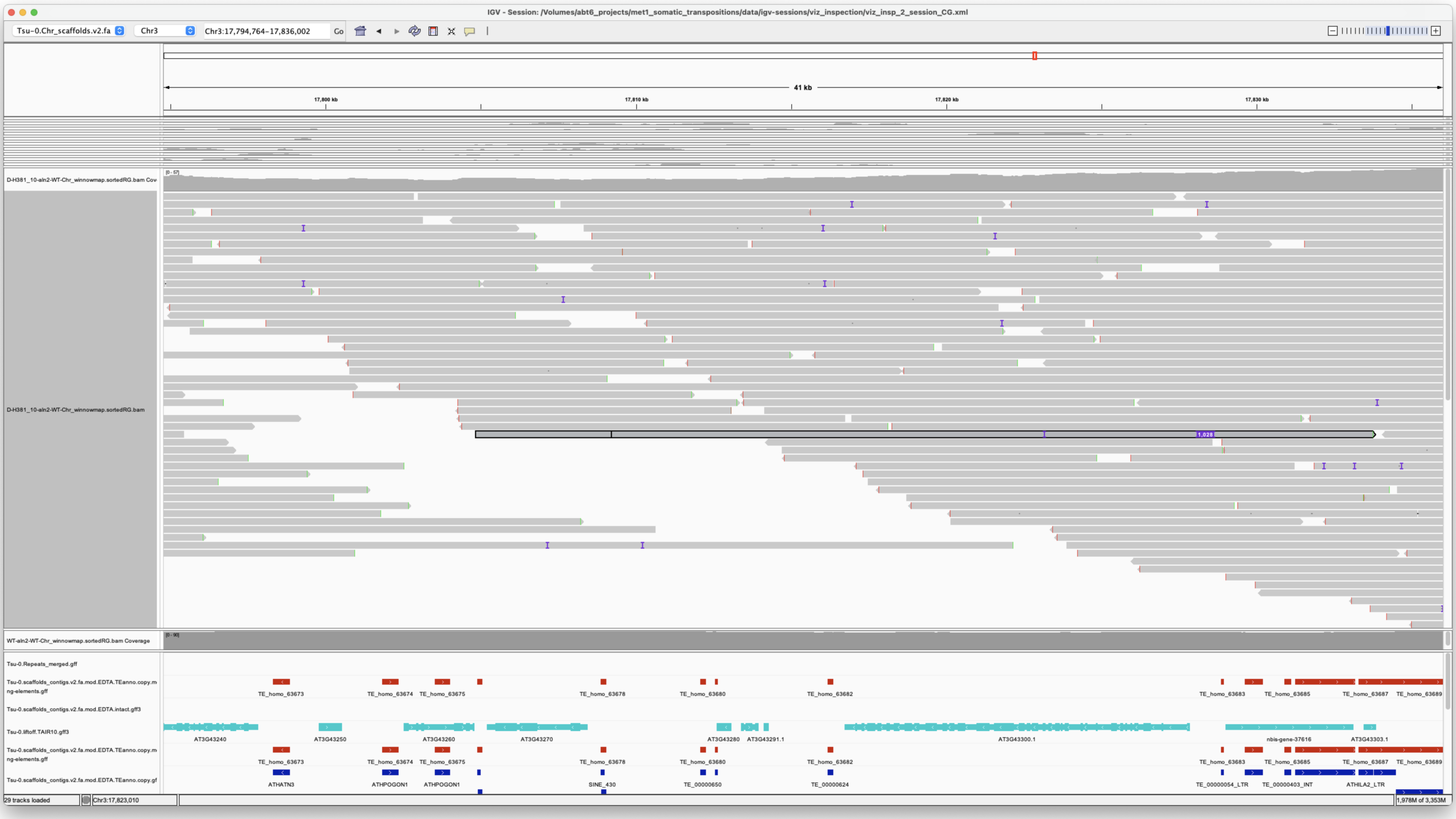


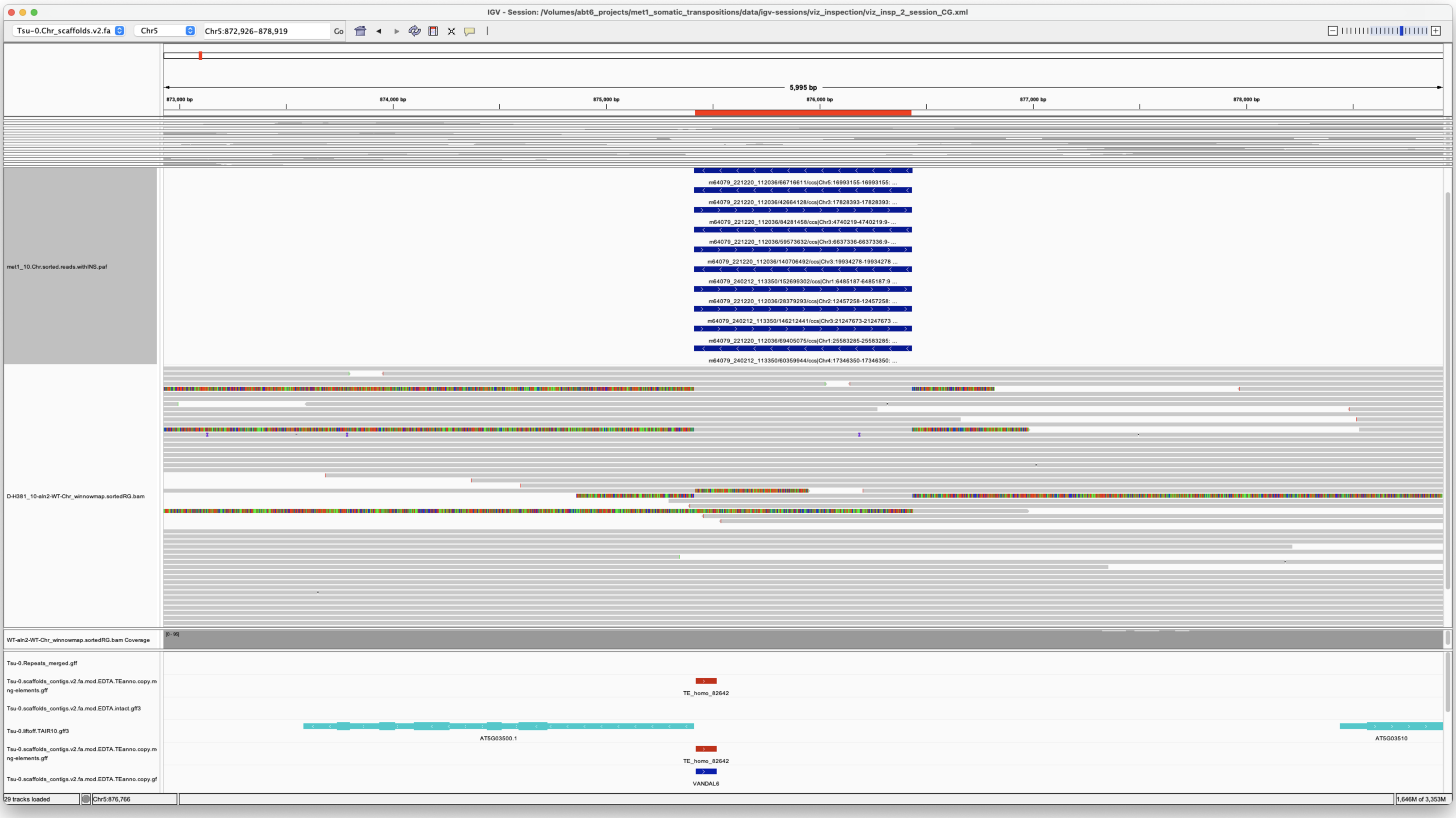


*Confirmed*

---

> Chr3 19934278 19934278 m64079_221220_112036/140706492/ccs Chr5 875413 876434
> Chr5|875414|876433|ID=TE_MANUAL_02;Name=PutativePackTypeCACTAMuDR;classification=DNA/DTC;sequence_ontology=MANUAL;identity=MANUAL;method=MANUAL;ID=TE_MANUAL_02;sequence_ontology=MANUAL met1_10

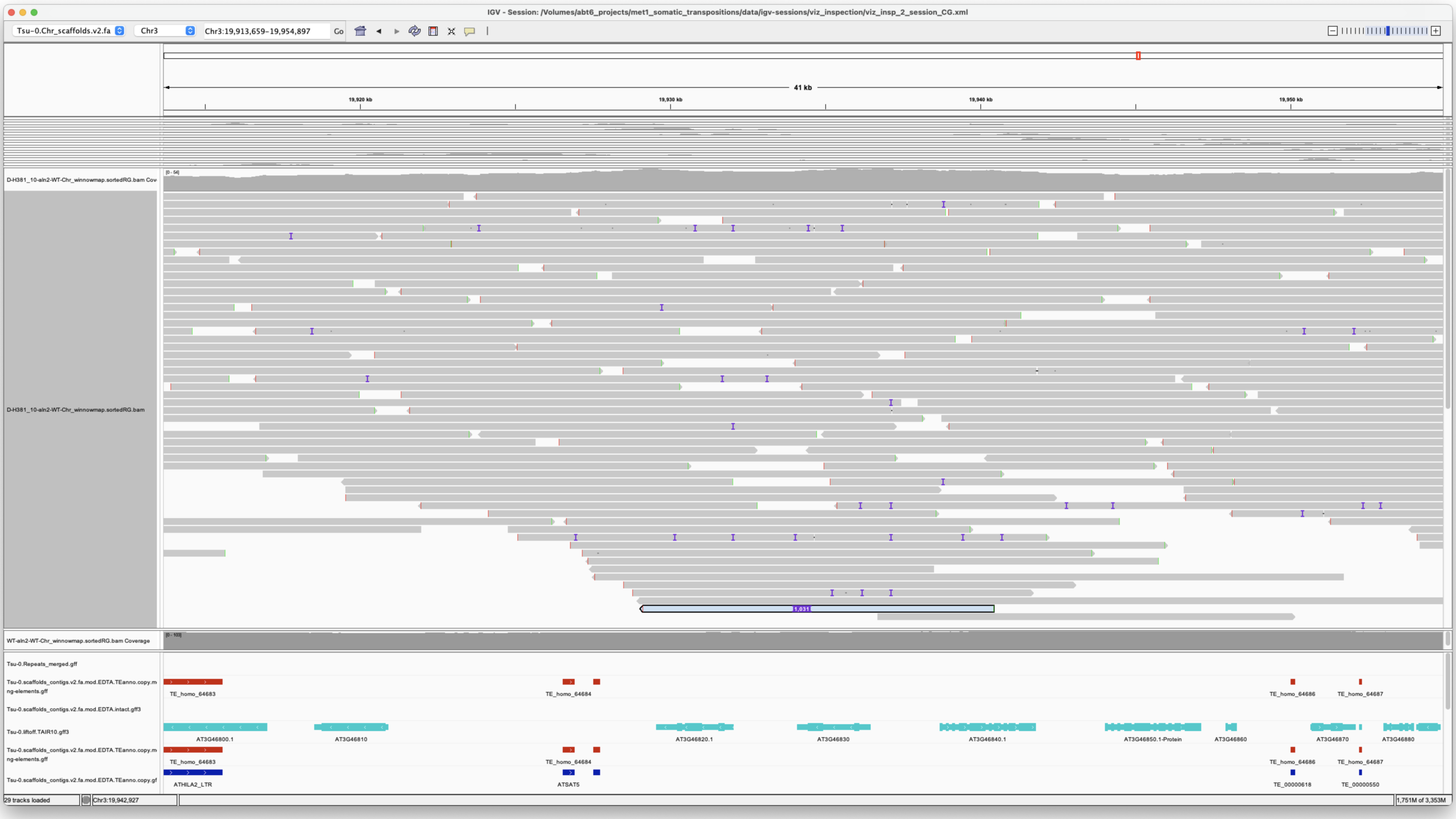

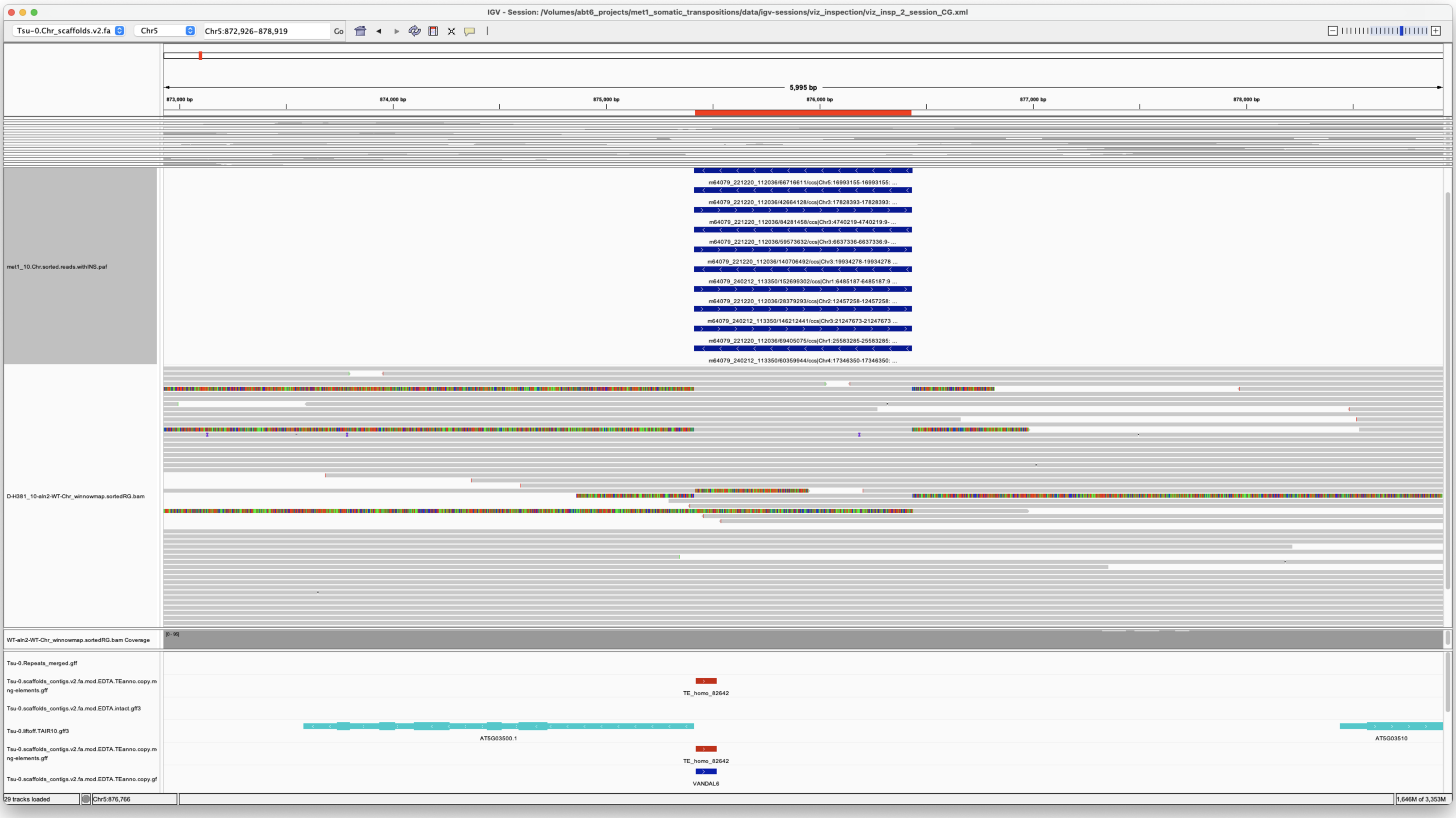

*Confirmed*

---

> Chr3 21247673 21247673 m64079_240212_113350/146212441/ccs Chr5 875413 876433 Chr5|875414|876433|ID=TE_MANUAL_02;Name=PutativePackTypeCACTAMuDR;classification=DNA/DTC;sequence_ontology=MANUAL;identity=MANUAL;method=MANUAL;ID=TE_MANUAL_02;sequence_ontology=MANUAL met1_10

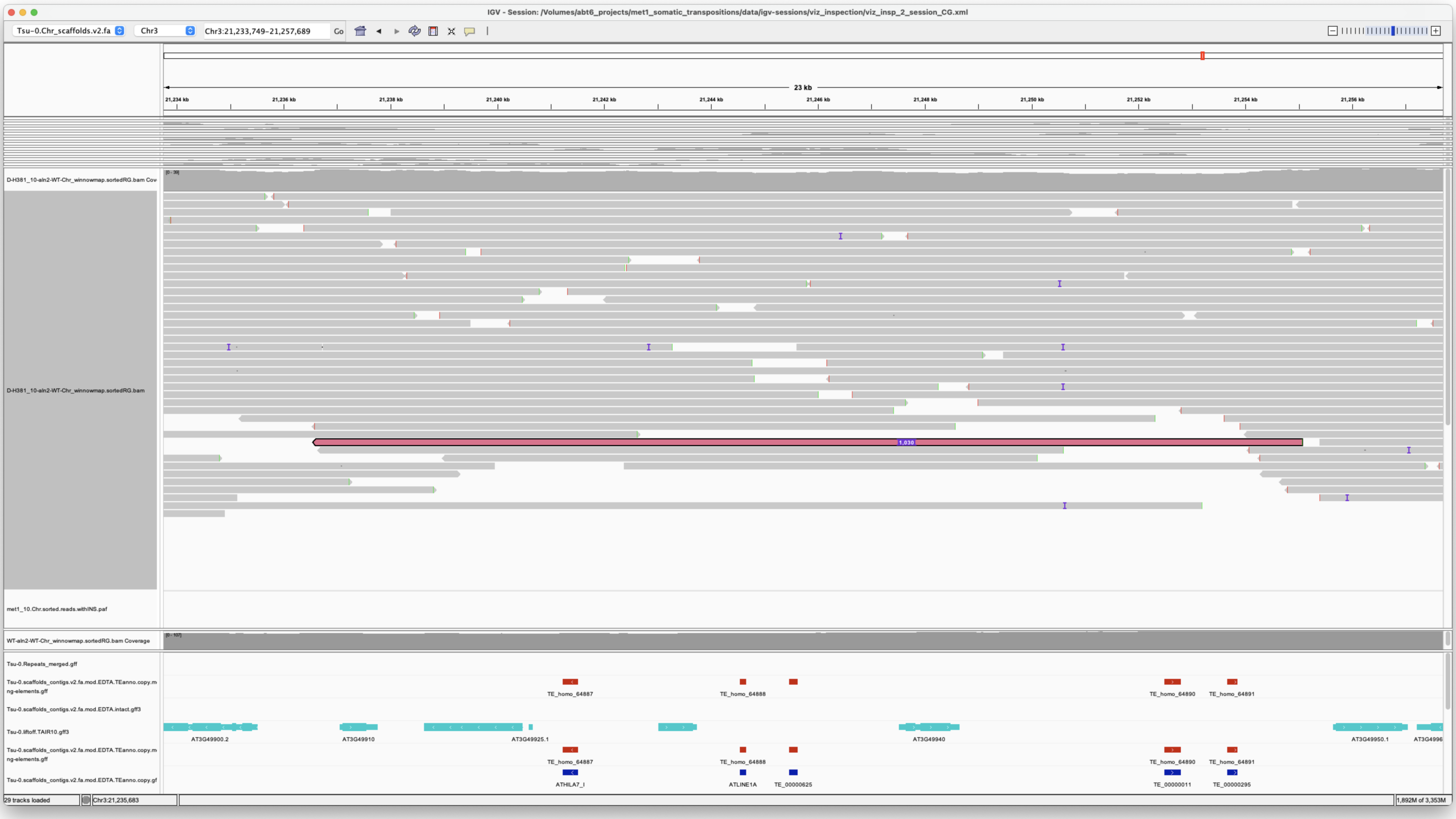

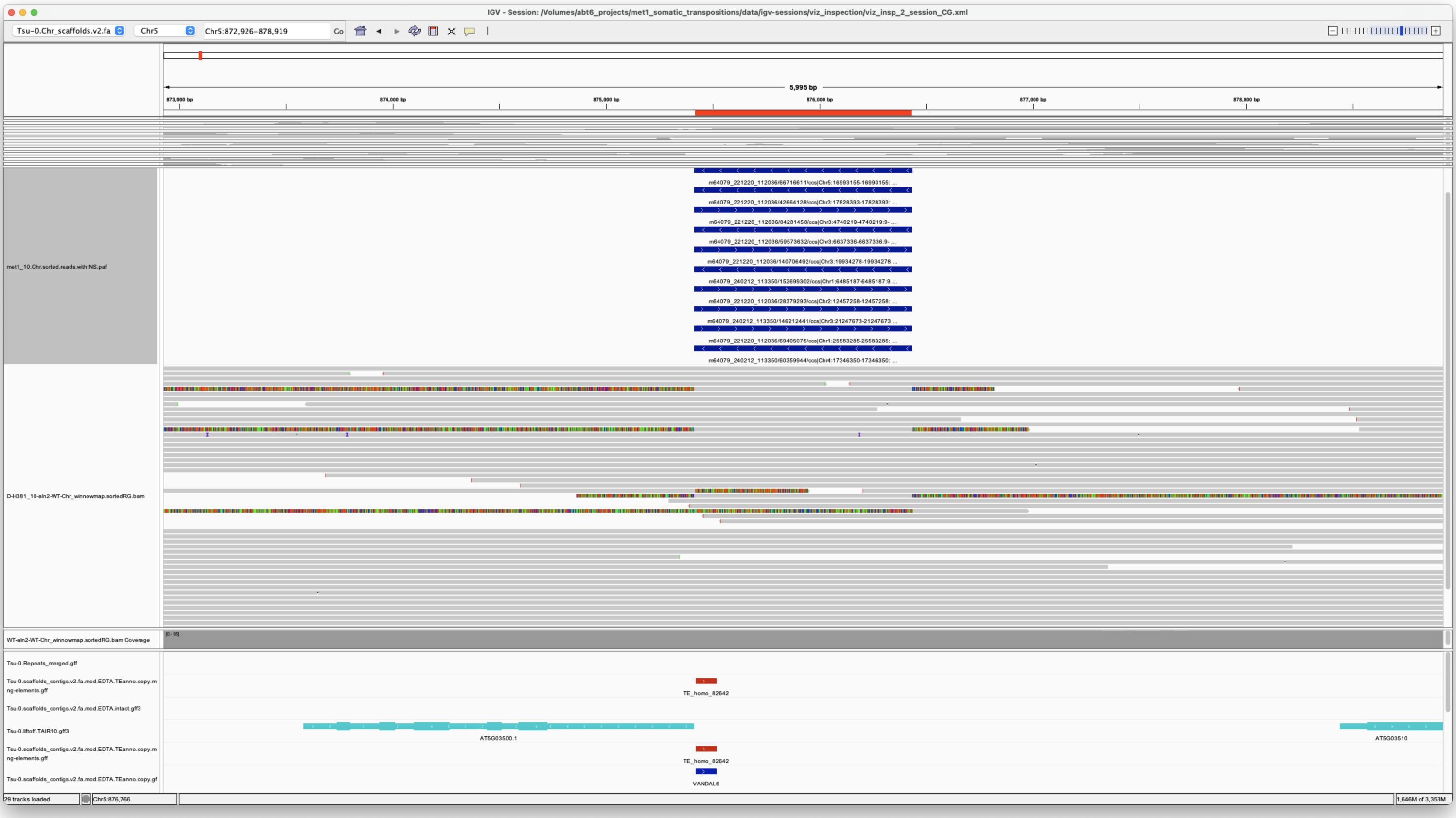

*Confirmed*

Chr4 939231 939231 m64079_240212_113350/138150026/ccs Chr1 11941103 11946436
Chr1|11941106|11946435|ID=LTRRT_5;Name=ATCOPIA93.2_Evade;Classification=LTR/Copia;Sequence_ontology=SO:0002264;ltr_identity=1.0000;Method=structural;motif=TACA;tsd=ATATG met1_10

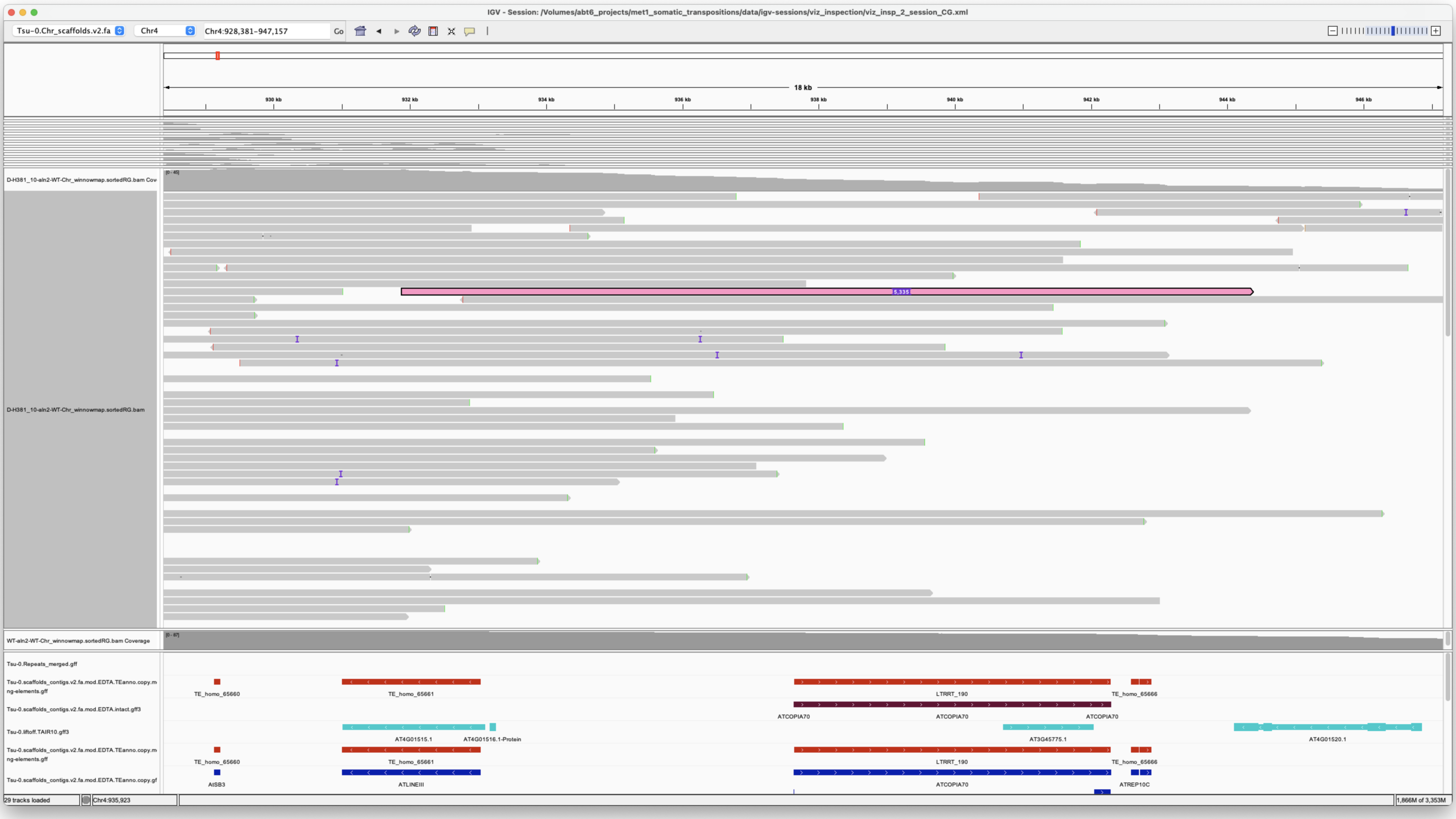


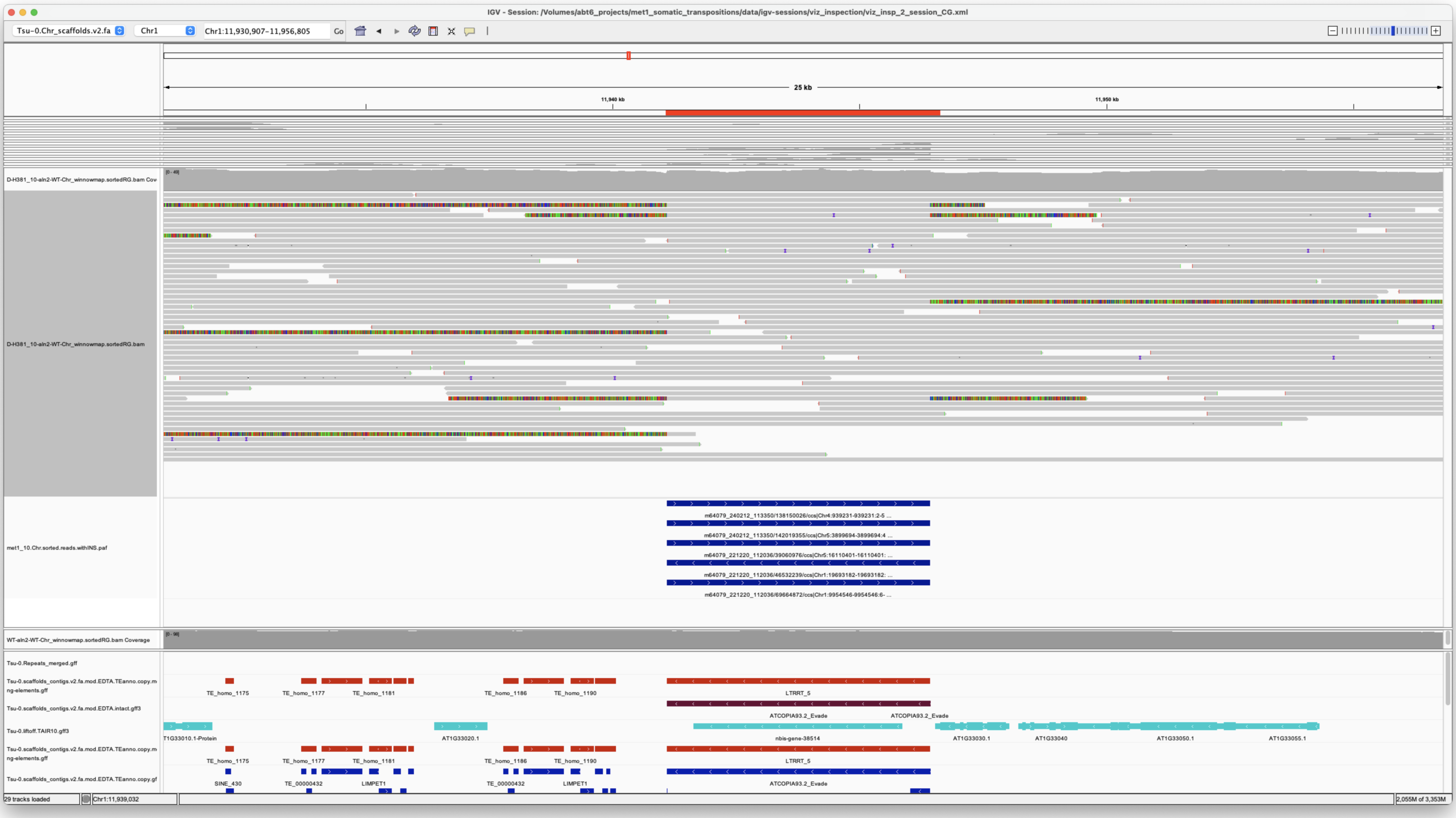


*Confirmed*

> Chr4 17346350 17346350 m64079_240212_113350/60359944/ccs Chr5 875414 876434 Chr5|875414|876433|ID=TE_MANUAL_02;Name=PutativePackTypeCACTAMuDR;classification=DNA/DTC;sequence_ontology=MANUAL;identity=MANUAL;method=MANUAL;ID=TE_MANUAL_02;sequence_ontology=MANUAL met1_10

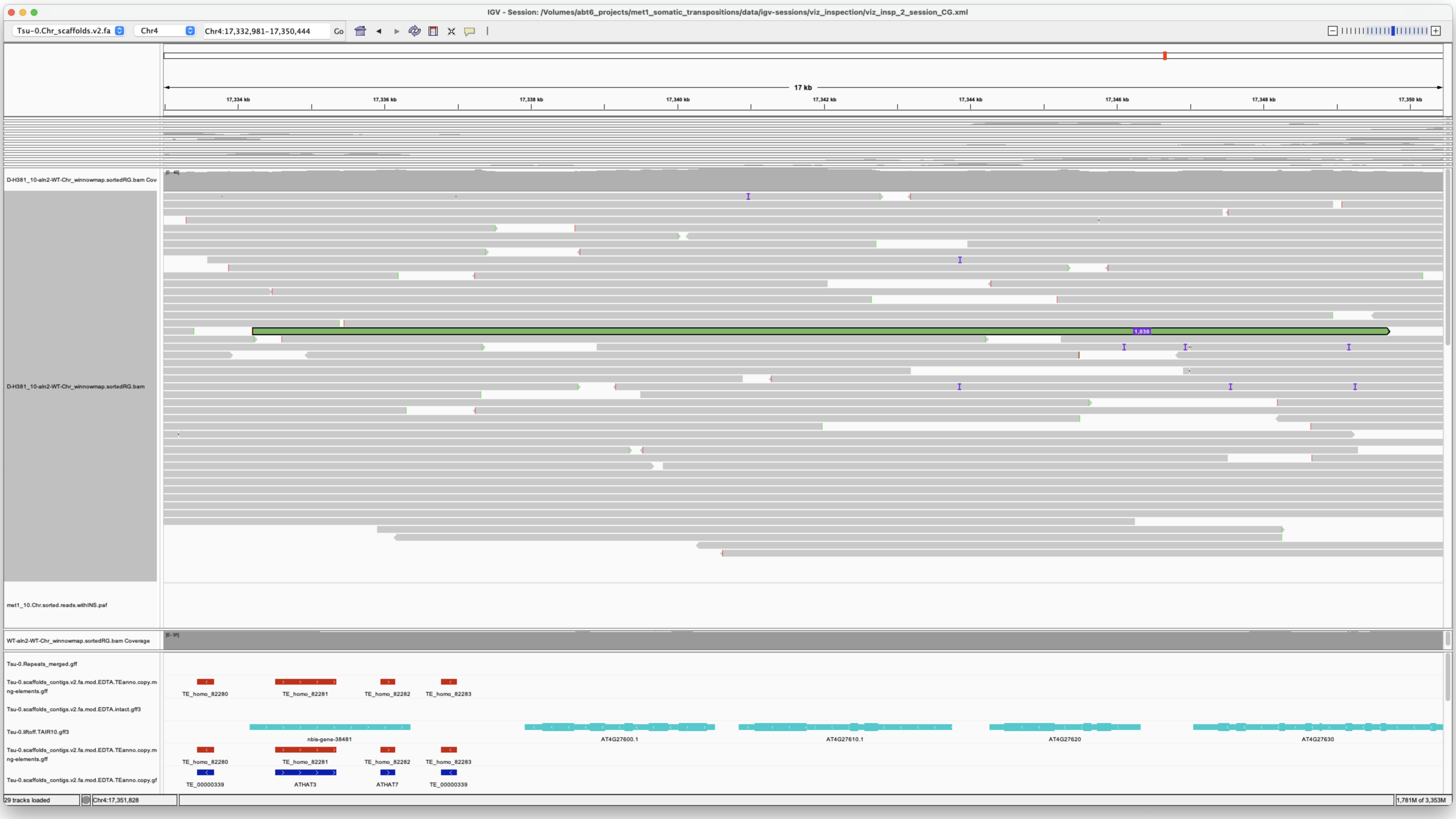


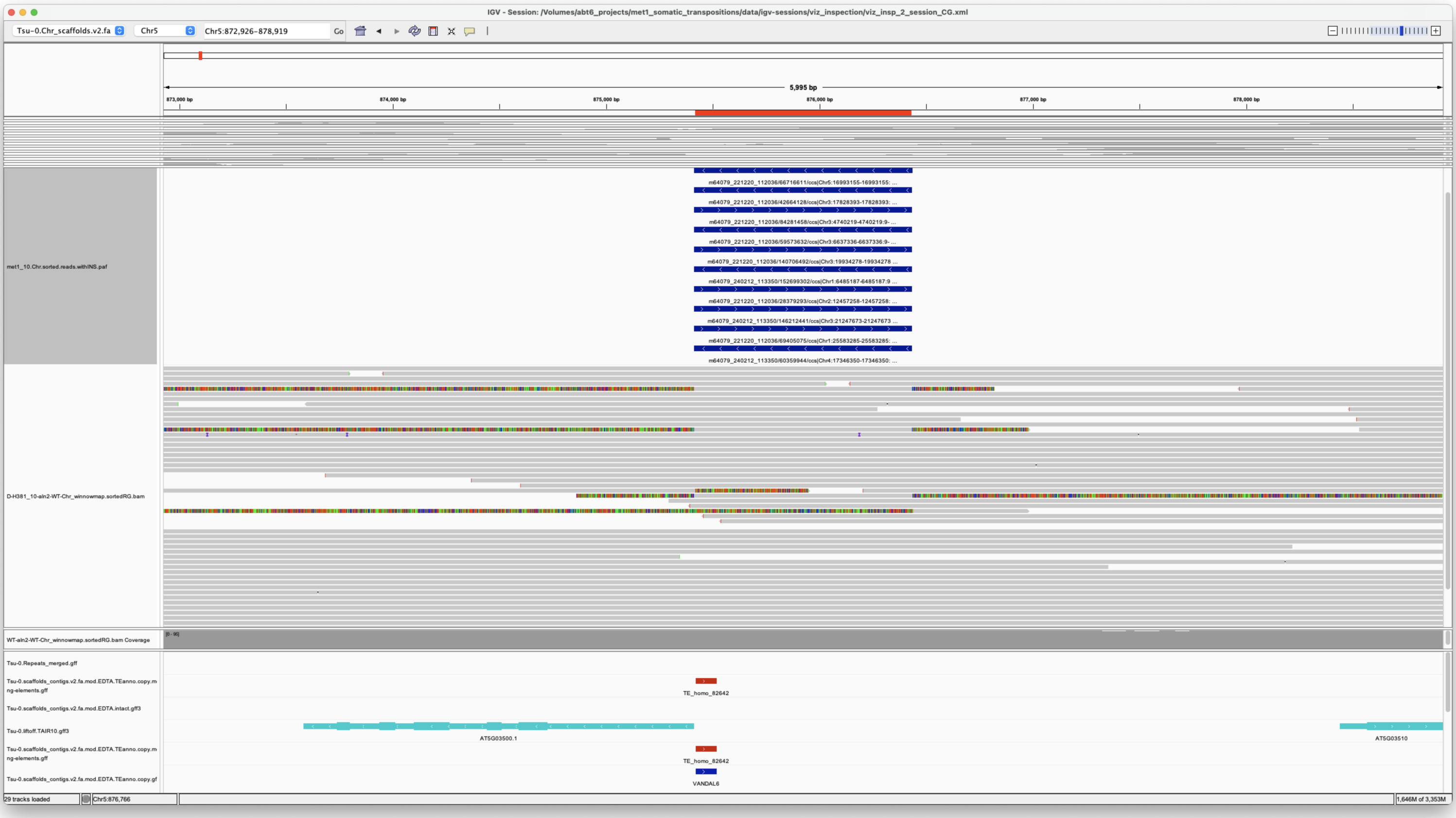


*Confirmed*

---

> Chr5 3899694 3899694 m64079_240212_113350/142019355/ccs Chr1 11941104 11946435 Chr1|11941106|11946435|ID=LTRRT_5;Name=ATCOPIA93.2_Evade;Classification=LTR/Copia;Sequence_ontology=SO:0002264;ltr_identity=1.0000;Method=structural;motif=TACA;tsd=ATATG met1_10

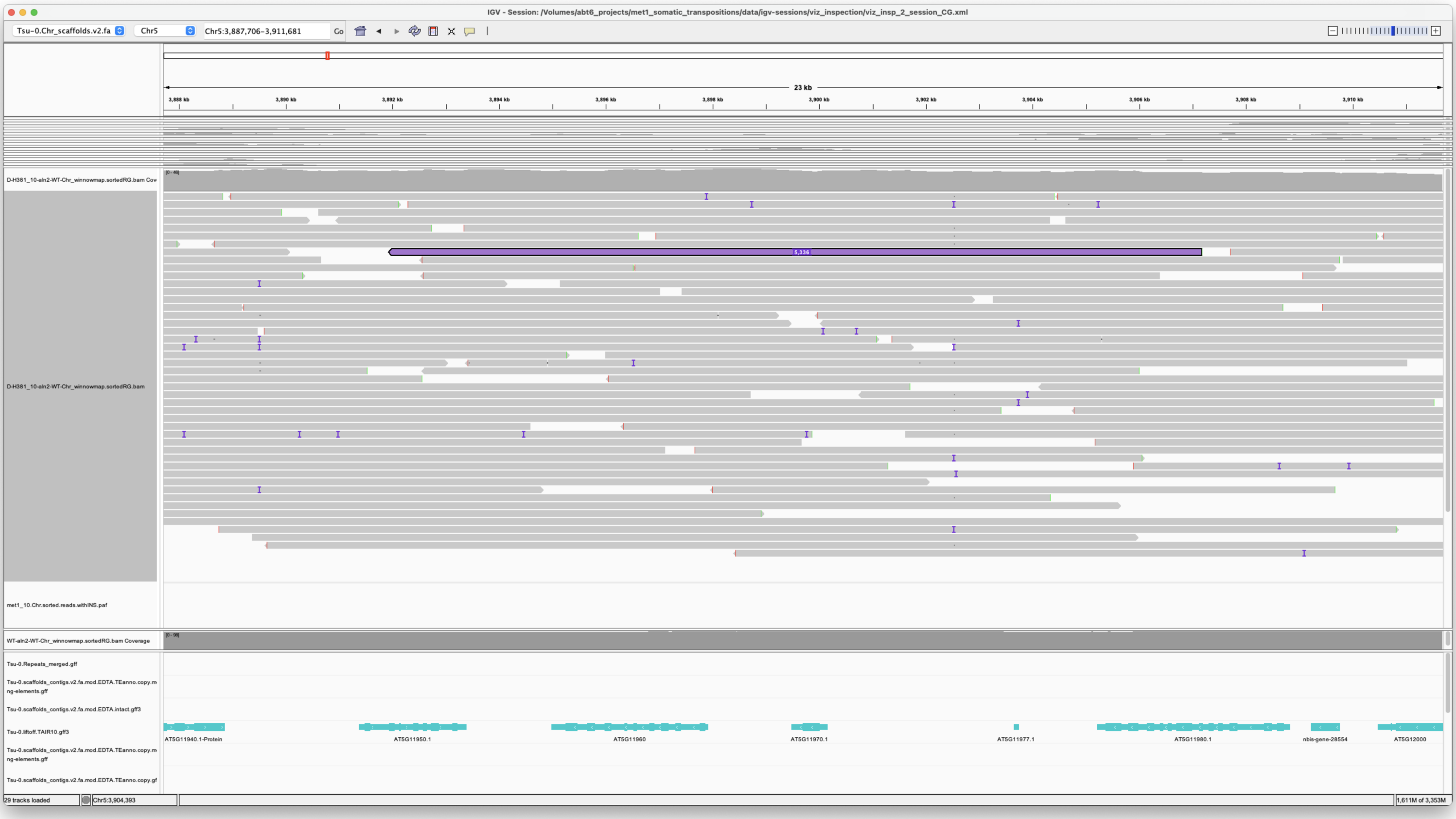


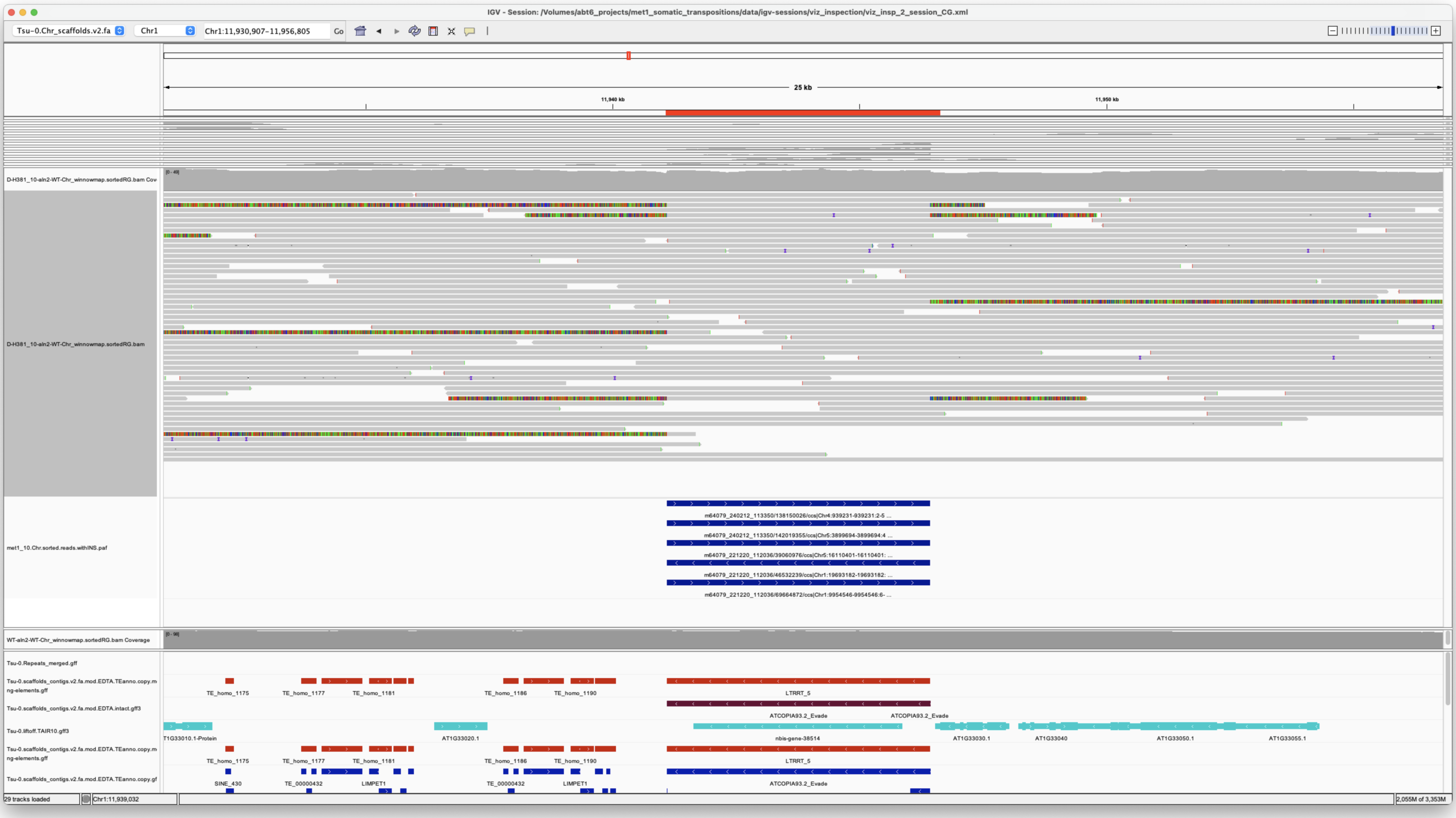


*Confirmed*

---

> Chr5 5803022 5803022 m64079_221220_112036/120455225/ccs Chr5 19152825 19160825 Chr5|19152829|19160825|ID=TE_homo_95640;Name=VANDAL21;classification=DNA/Mutator;sequence_ontology=SO:0002280;identity=0.976;method=homology;ID=TE_homo_98501;sequence_ontology=SO:0002280|ID=TE_homo_95641;Name=VANDAL21;classification=DNA/Mutator;sequence_ontology=SO:0002280;identity=0.966;method=homology;ID=TE_homo_98502;sequence_ontology=SO:0002280 met1_10

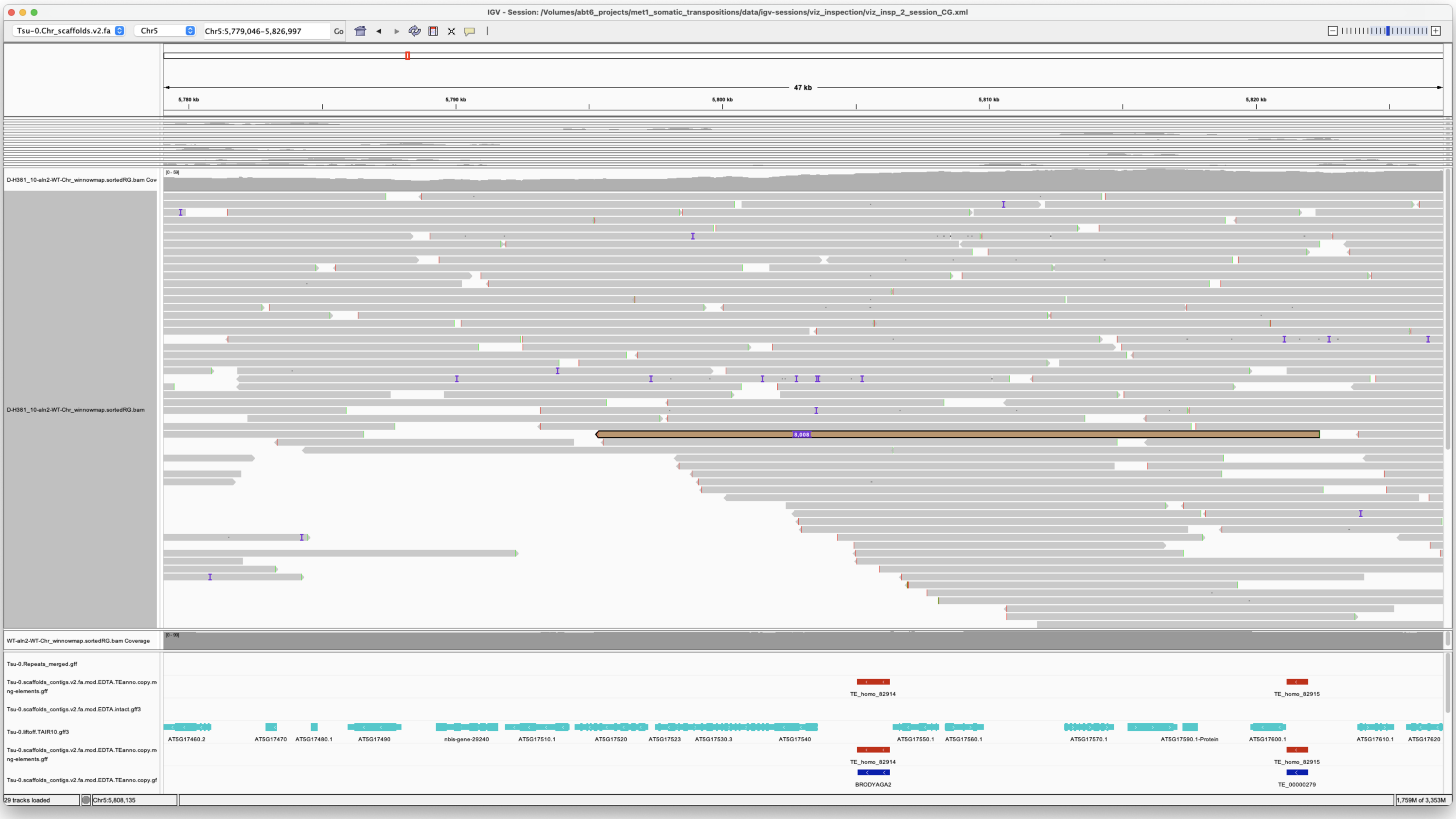


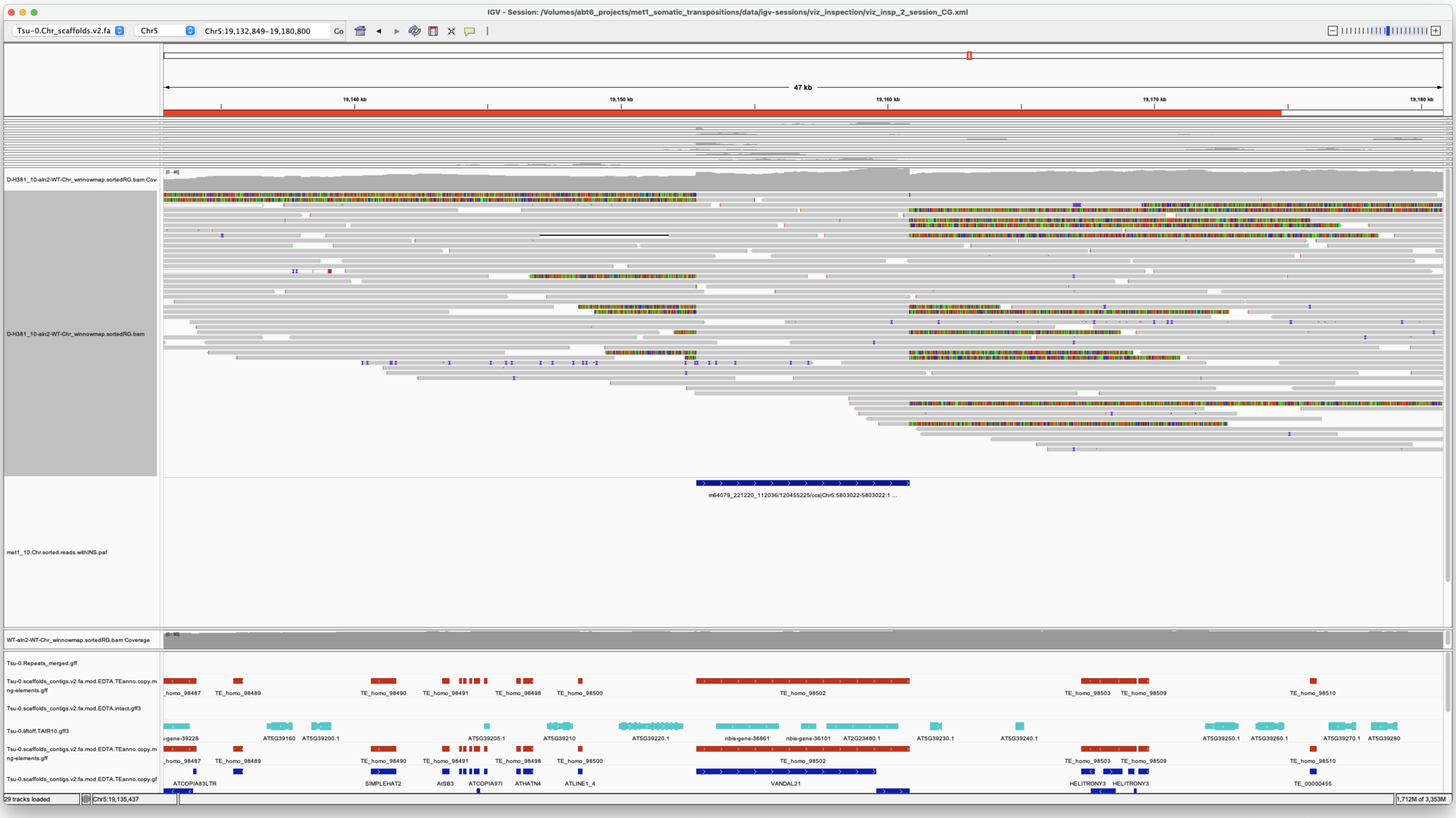


*Confirmed*

---

> Chr5 16110401 16110401 m64079_221220_112036/39060976/ccs Chr1 11941106 11946436
> Chr1|11941106|11946435|ID=LTRRT_5;Name=ATCOPIA93.2_Evade;Classification=LTR/Copia;Sequence_ontology=SO:0002264;ltr_identity=1.0000;Method=structural;motif=TACA;tsd=ATATG met1_10

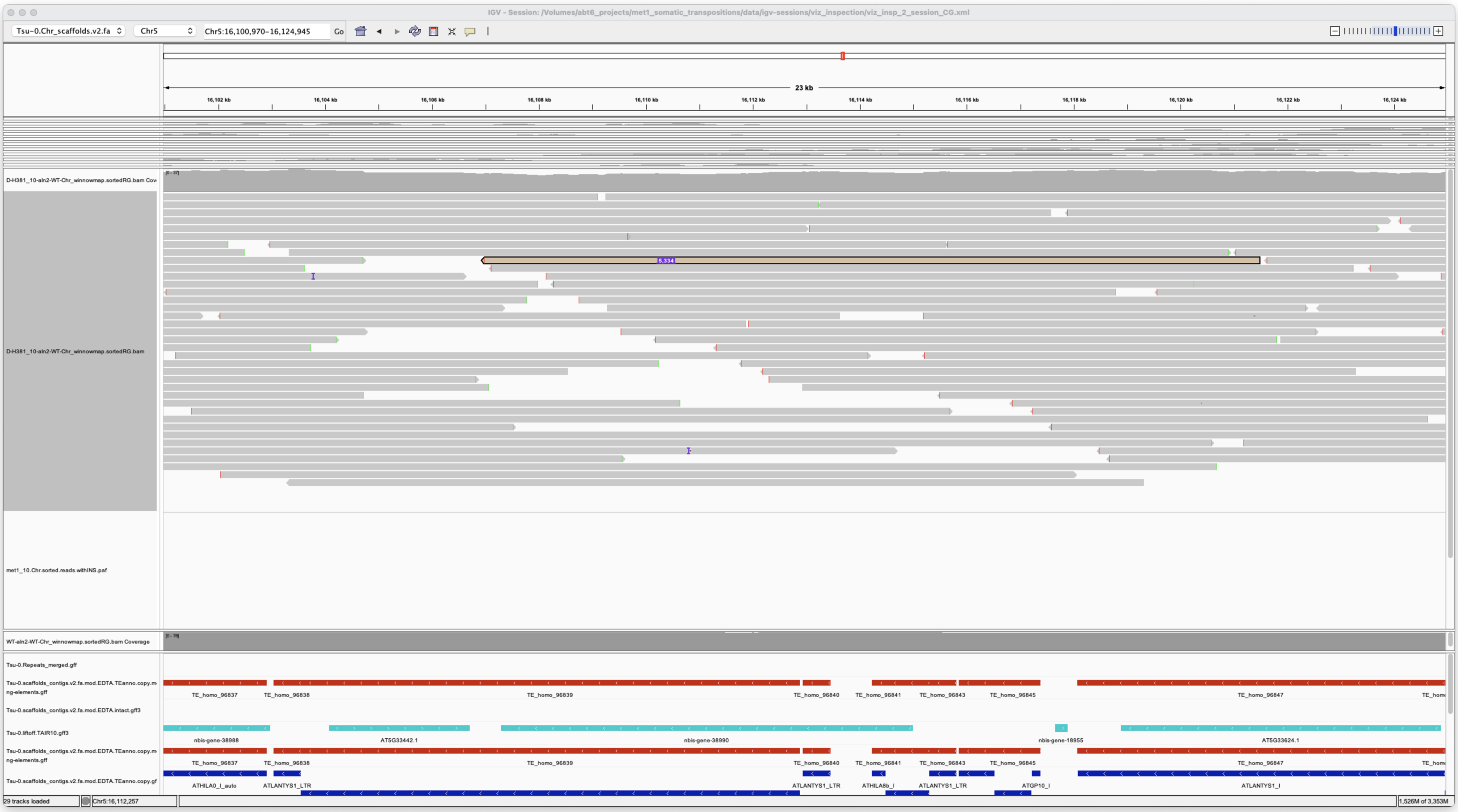


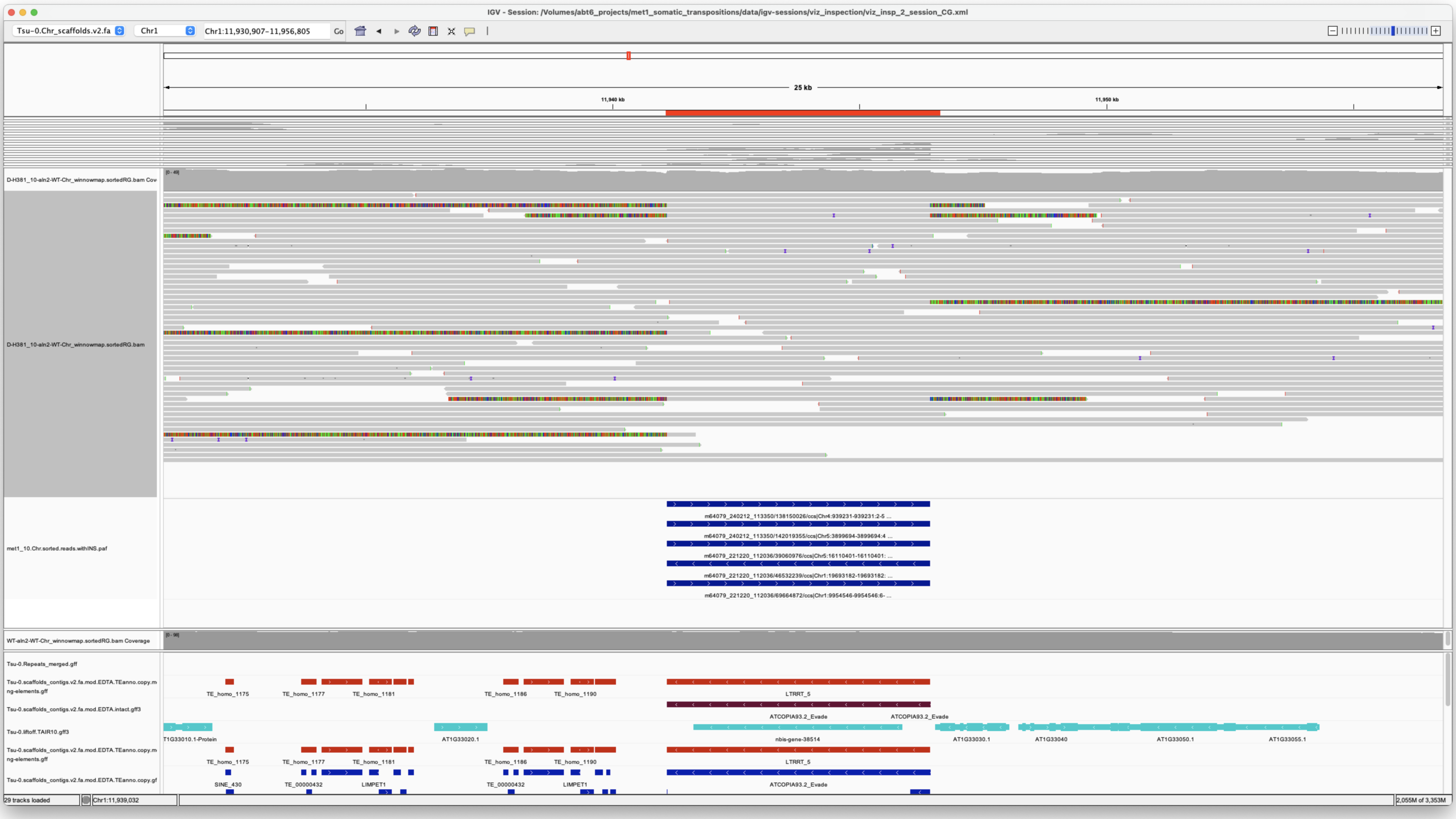


*Confirmed*

> Chr5 16993155 16993155 m64079_221220_112036/66716611/ccs Chr5 875413 876436 Chr5|875414|876433|ID=TE_MANUAL_02;Name=PutativePackTypeCACTAMuDR;classification=DNA/DTC;sequence_ontology=MANUAL;identity=MANUAL;method=MANUAL;ID=TE_MANUAL_02;sequence_ontology=MANUAL met1_10

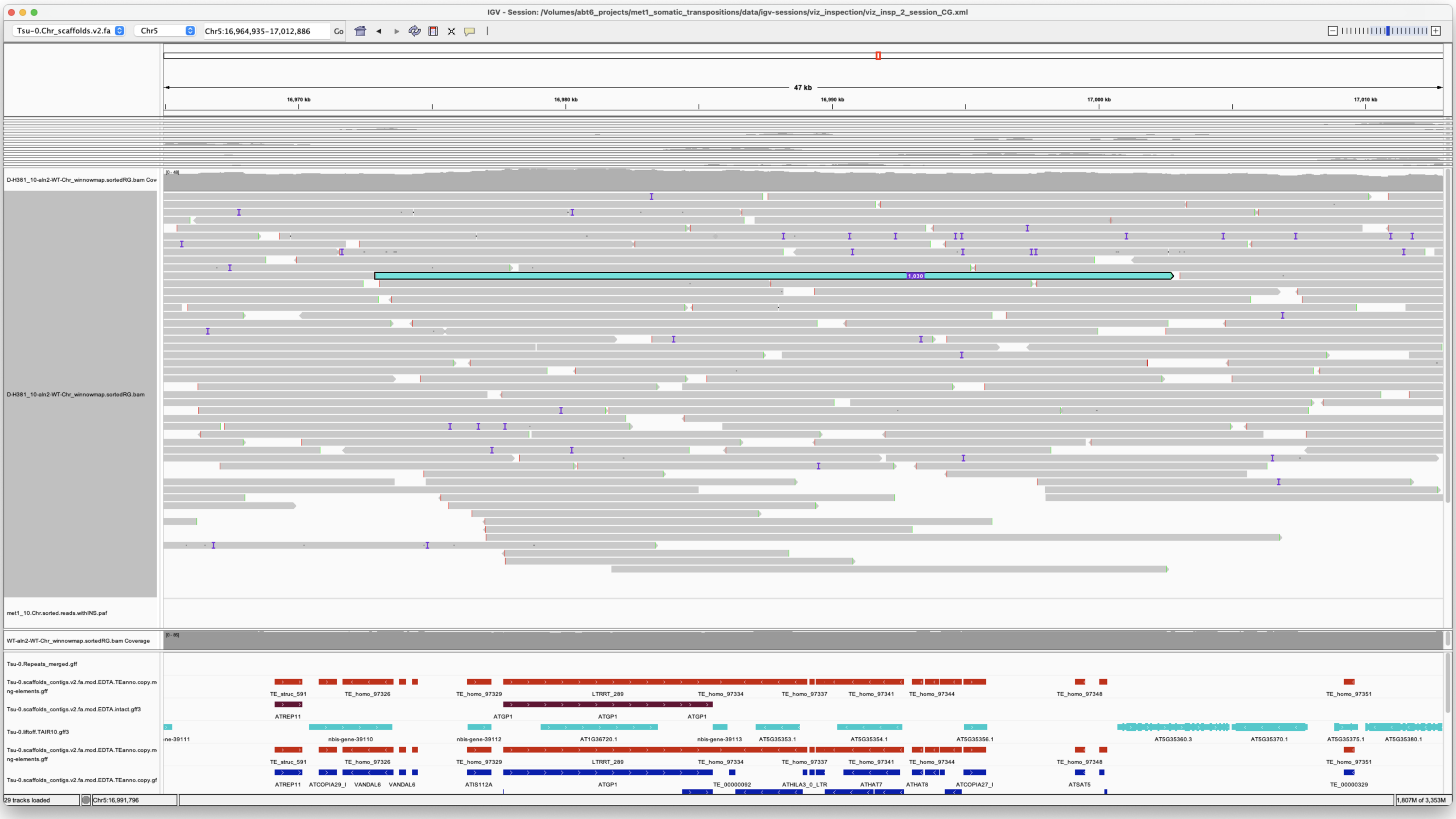
IGV - Session: /Volumes/abt6_projects/met1_somatic_transpositions/data/igv-sessions/viz_inspection/viz_insp_2_session_CG.xml
Tsu-0.Chr_scaffolds.v2.fa
Chr5
Chr5:16,964,935-17,012,886
47 kb
D-H381_10-aln2-WT-Chr_winnowmap.sortedRG.bam
met1_10.Chr.sorted.reads.withINS.paf
WT-aln2-WT-Chr_winnowmap.sortedRG.bam Coverage
Tsu-0.Repeats_merged.gff
Tsu-0.liftoff.TAIR10.gff3
29 tracks loaded

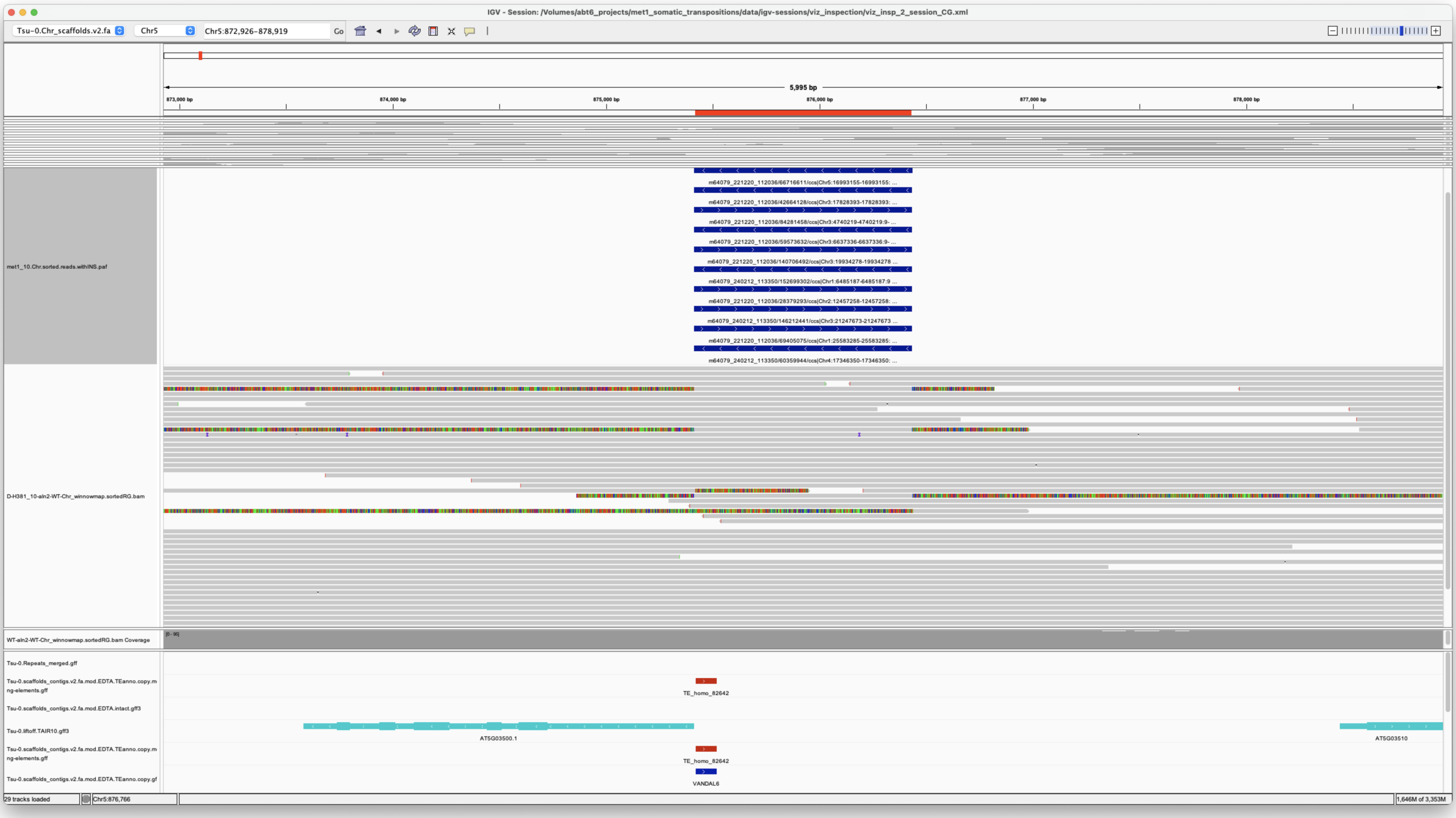
IGV - Session: /Volumes/abt6_projects/met1_somatic_transpositions/data/igv-sessions/viz_inspection/viz_insp_2_session_CG.xml
Tsu-0.Chr_scaffolds.v2.fa
Chr5
Chr5:872,926-878,919
5,995 bp
met1_10.Chr.sorted.reads.withINS.paf
D-H381_10-aln2-WT-Chr_winnowmap.sortedRG.bam
WT-aln2-WT-Chr_winnowmap.sortedRG.bam Coverage
Tsu-0.Repeats_merged.gff
TE_homo_82642
Tsu-0.liftoff.TAIR10.gff3
VANDAL6
29 tracks loaded

*Confirmed*
