## Additional File 4 for "Long-read detection of transposable element mobilization in the soma of hypomethylated *Arabidopsis thaliana* individuals": Synteny-check_Col-0-TEs-in-Tsu-0.pdf

##### Chr1 11148307 11152649 ATCOPIA51

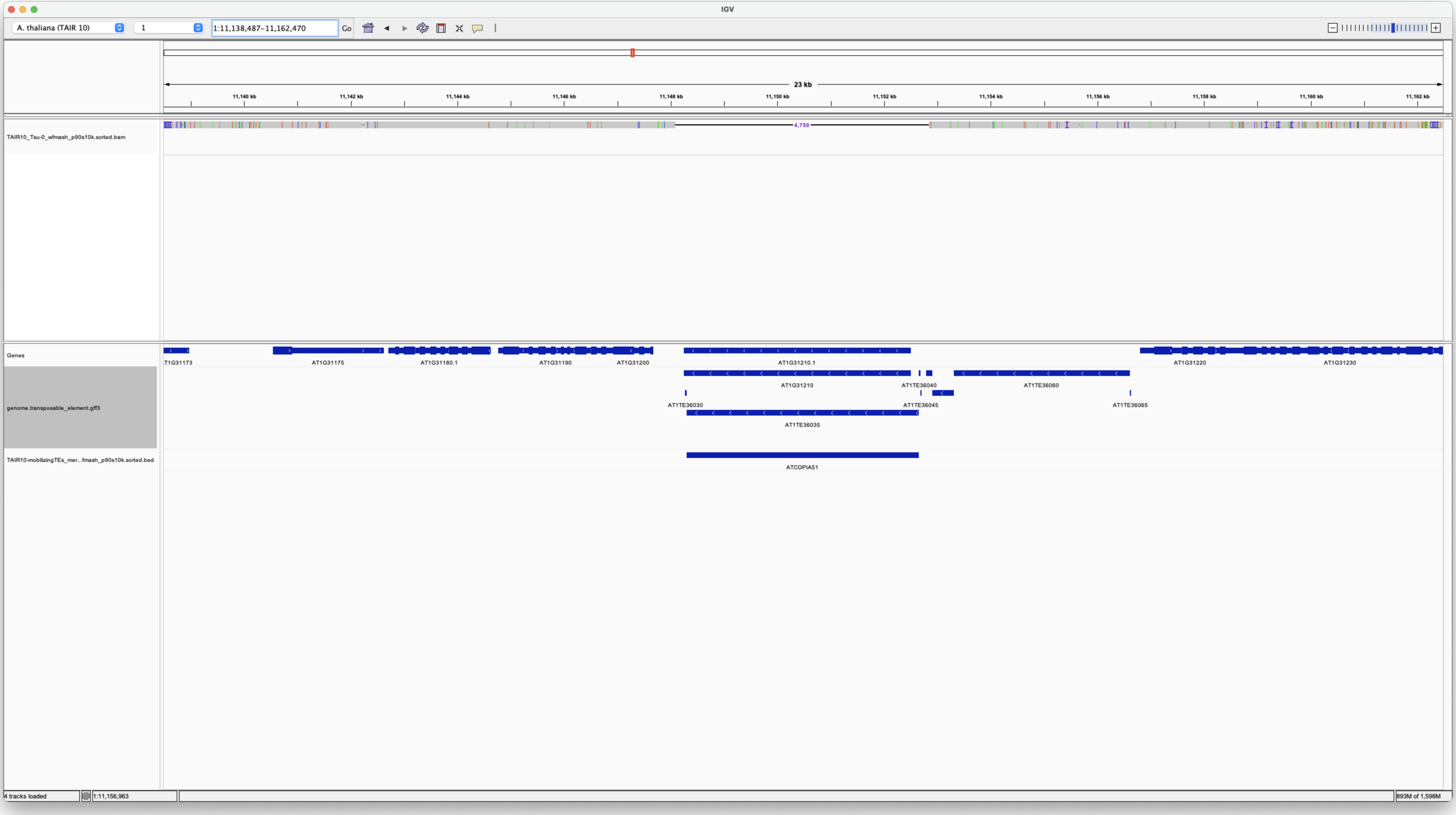


No Tsu-0

##### Chr1 12755059 12760395 ATCOPIA93

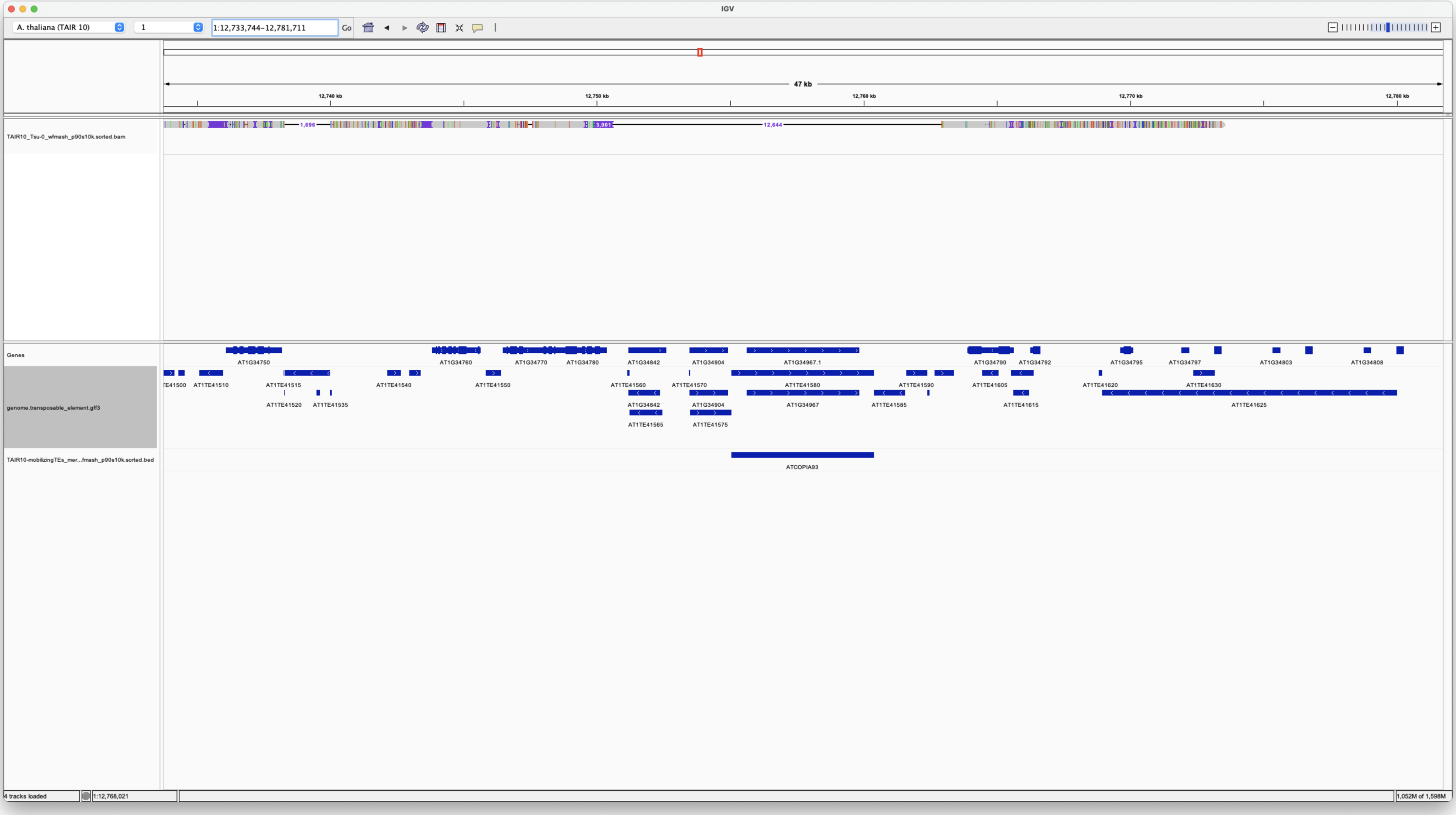


No Tsu-0

##### Chr1 17203925 17206318 ATCOPIA63

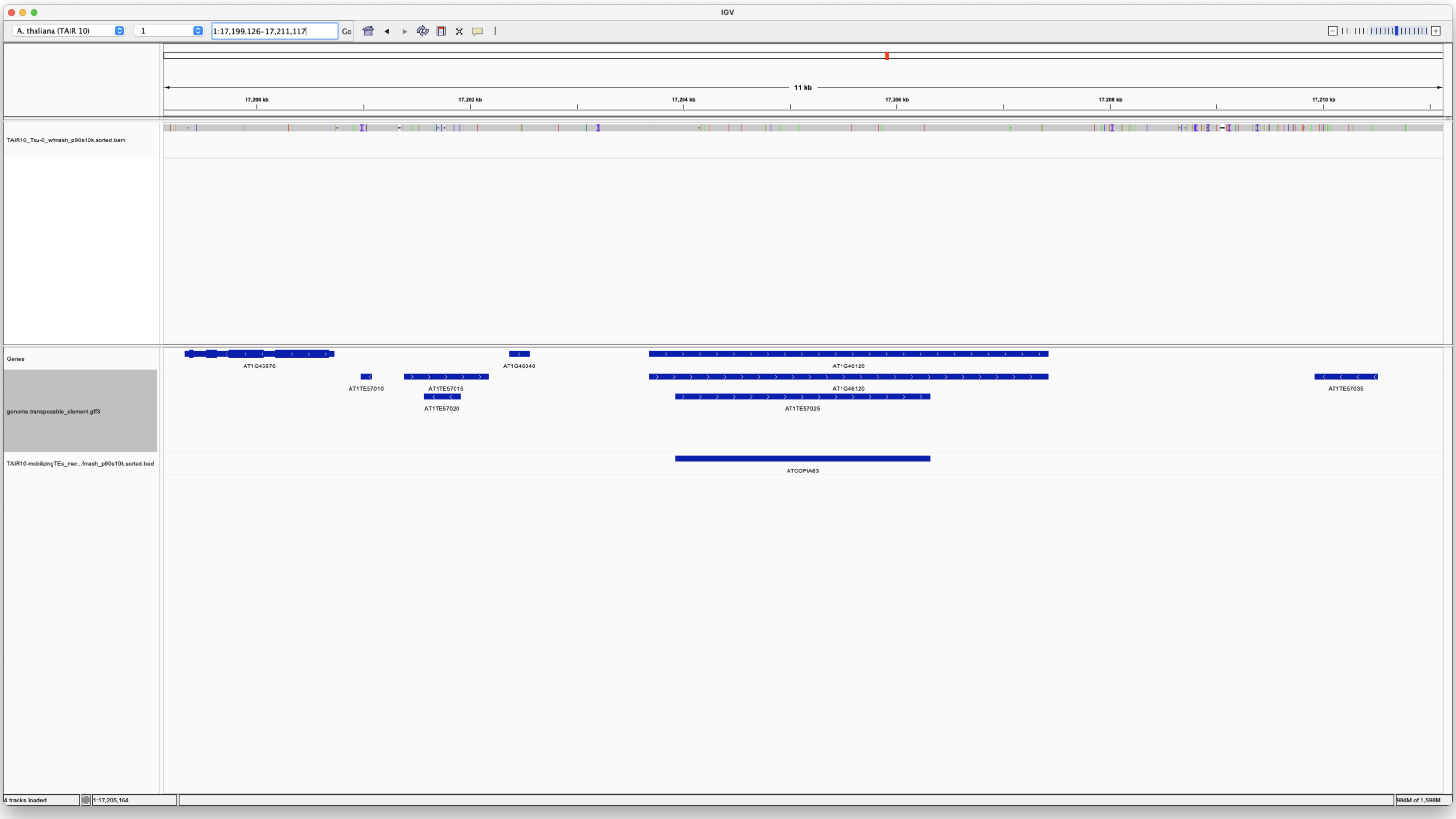


YES Tsu-0

**Chr1 21796937 21801751 ATRE1**

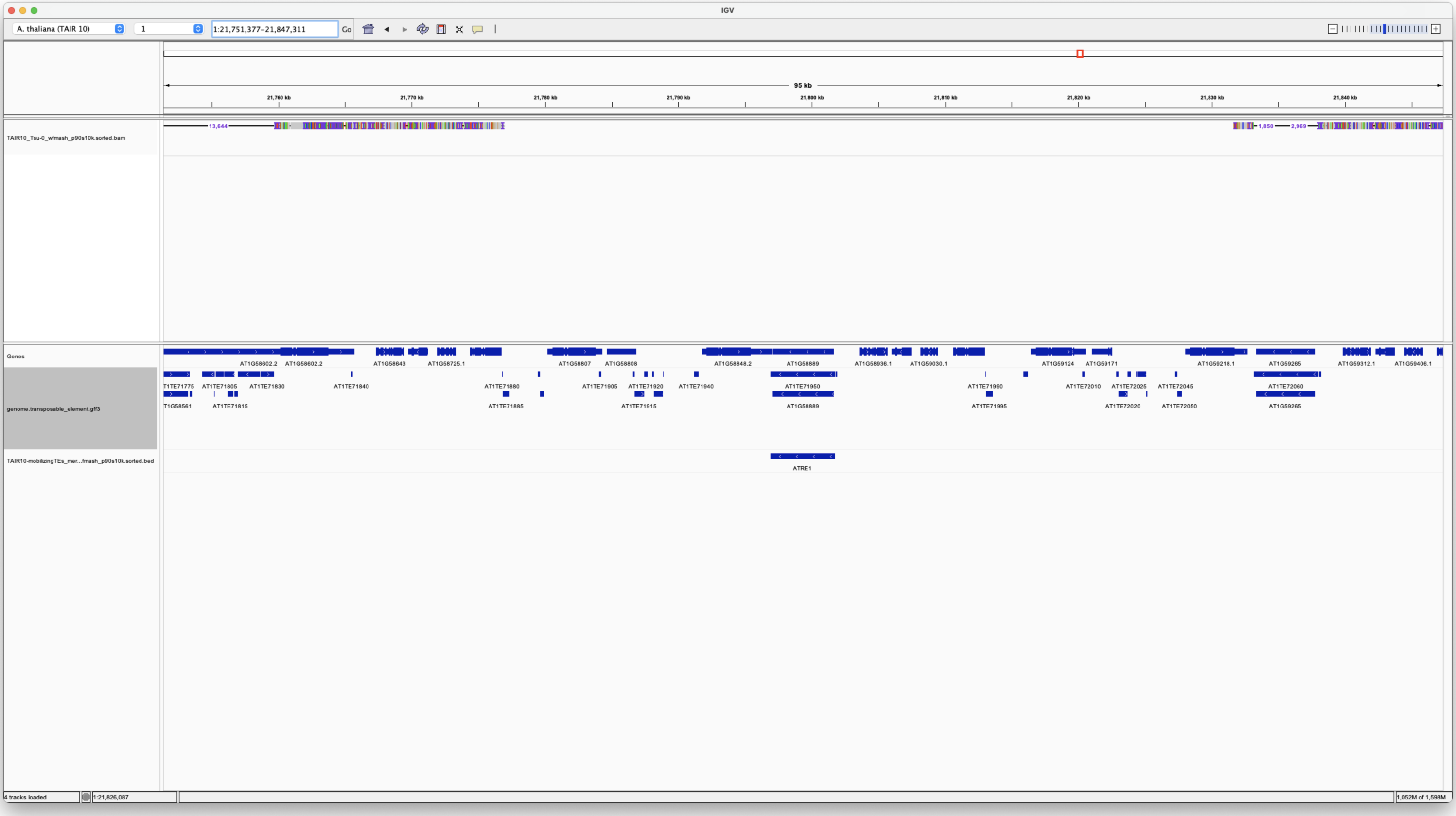


No Tsu-0

**Chr2 10001479 10009658 VANDAL21**

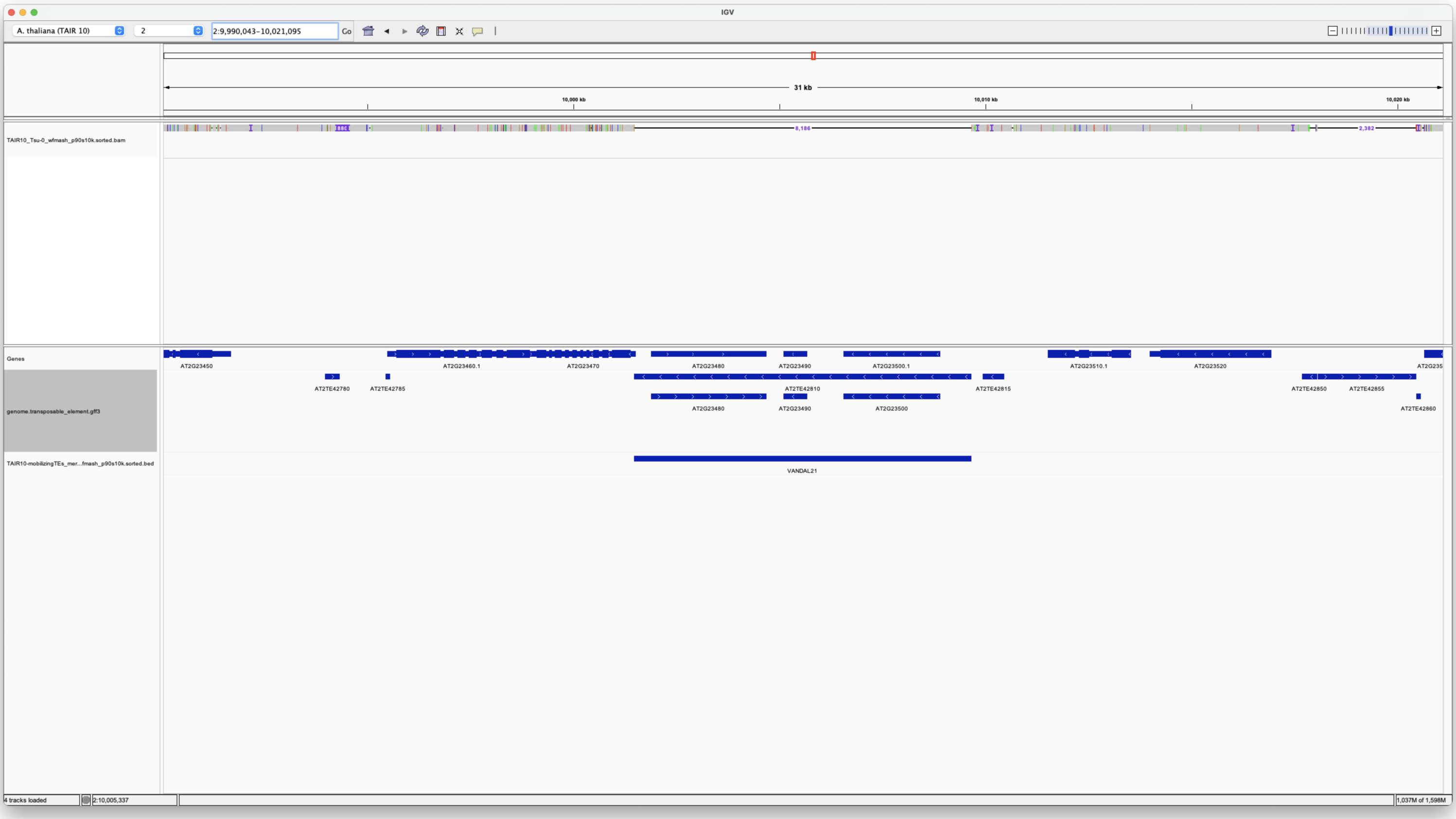


No Tsu-0

##### Chr2 34379 36436 ATGP2N

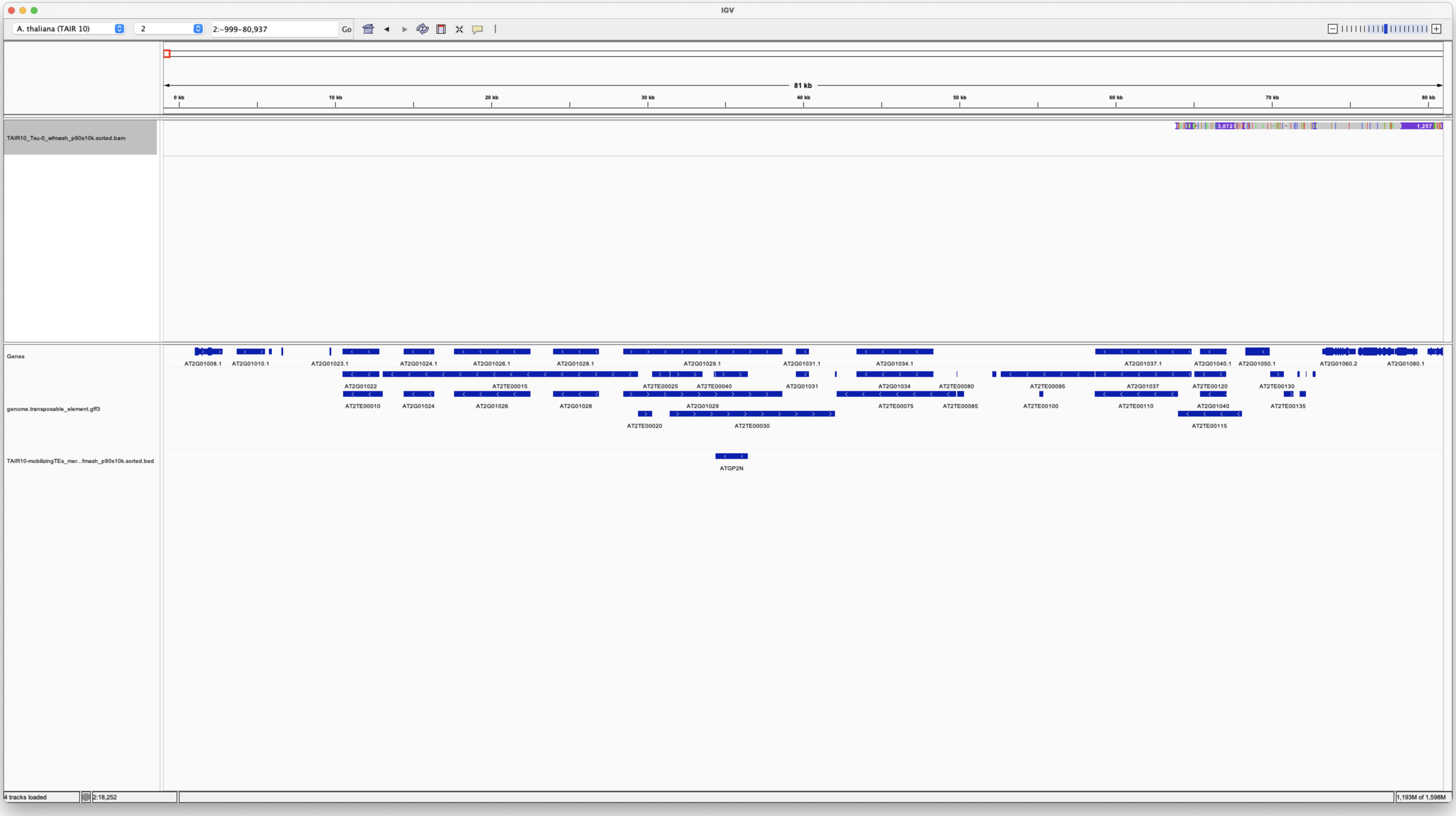


Very telomeric -> dubious

No Tsu-0

##### Chr2 4900802 4909281 ATENSPM3

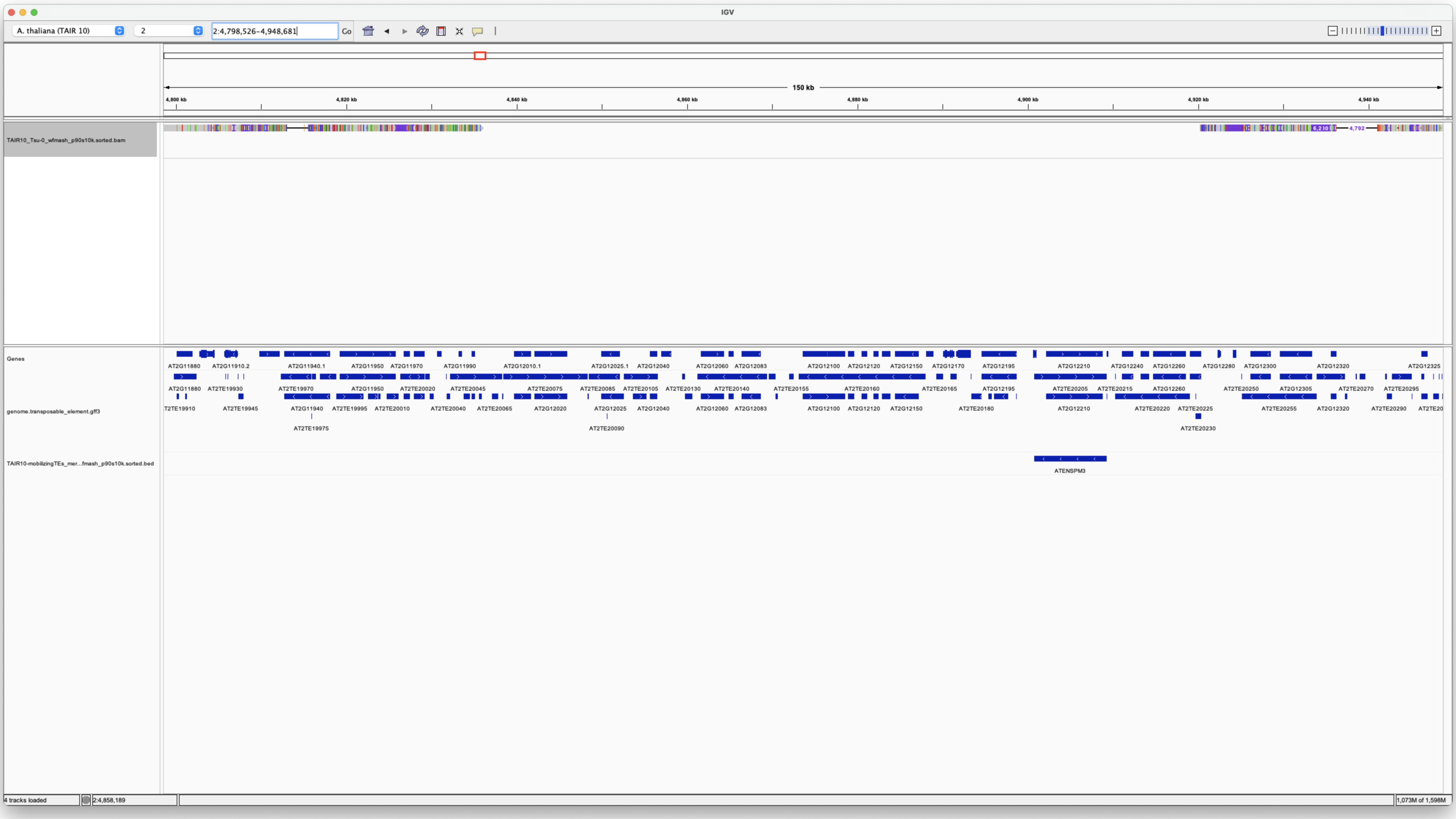


No Tsu-0

**Chr2 5852698 5857428 ATCOPIA13**

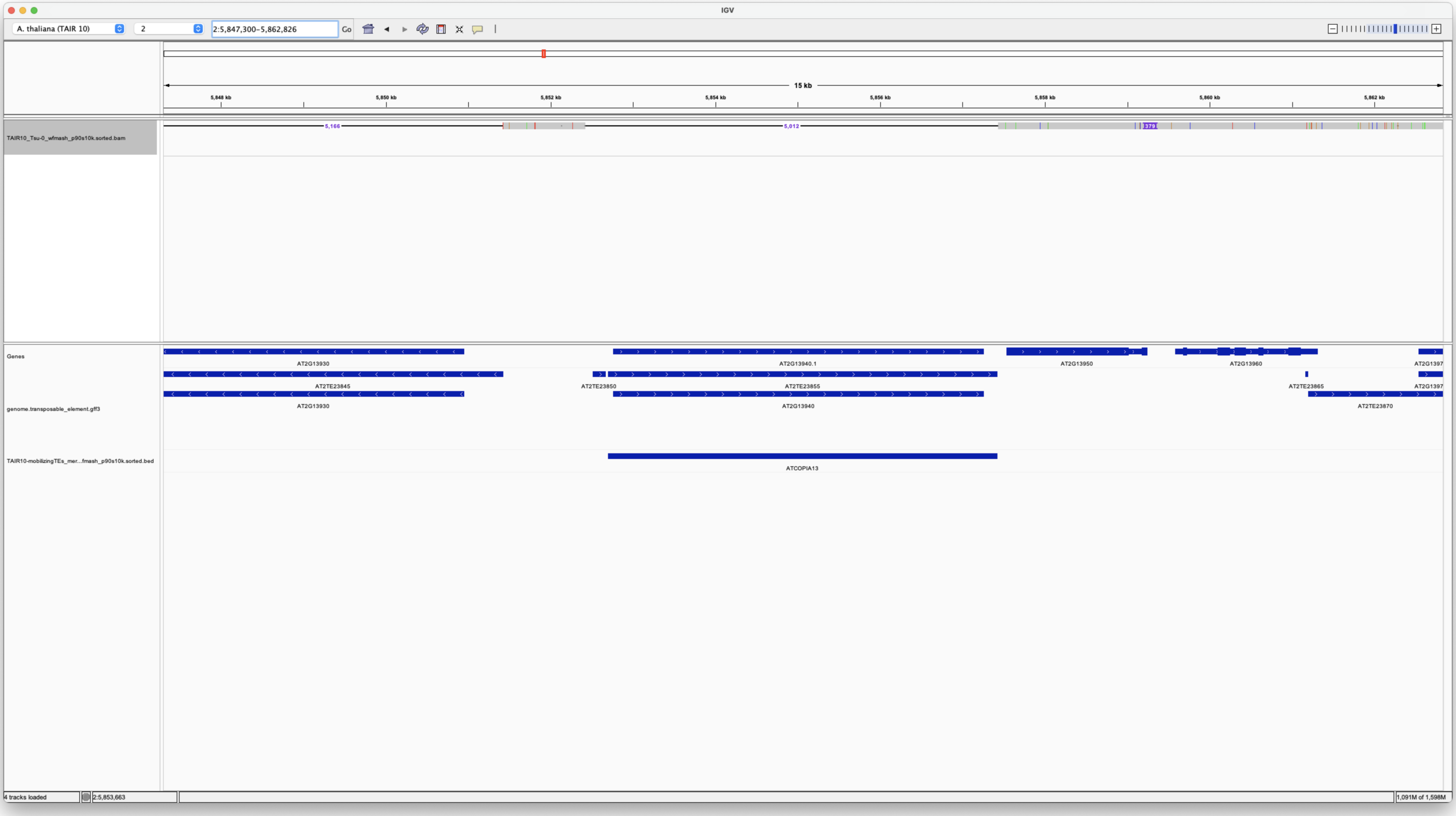


No Tsu-0

**Chr3 15283647 15283911 ATENSPM3**

**Chr3 15283666 15286610 Pack-CACTA2a**

**Chr3 15286419 15286632 ATENSPM3**

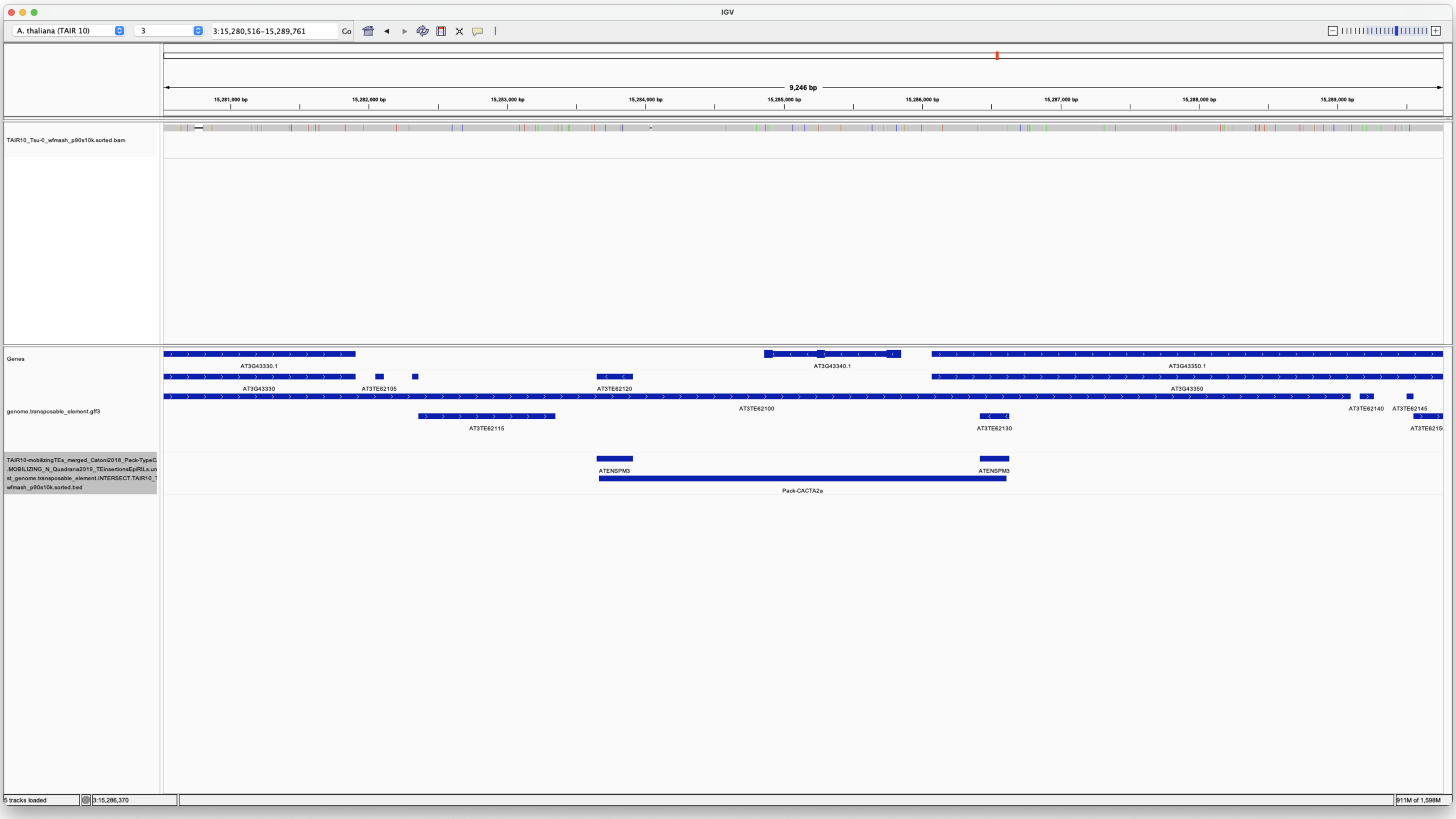

YES Tsu-0

**Chr4 4358128 4358579 ATENSPM3**

**Chr4 4358131 4360293 Pack-CACTA1a**

**Chr4 4359956 4360293 ATENSPM3**

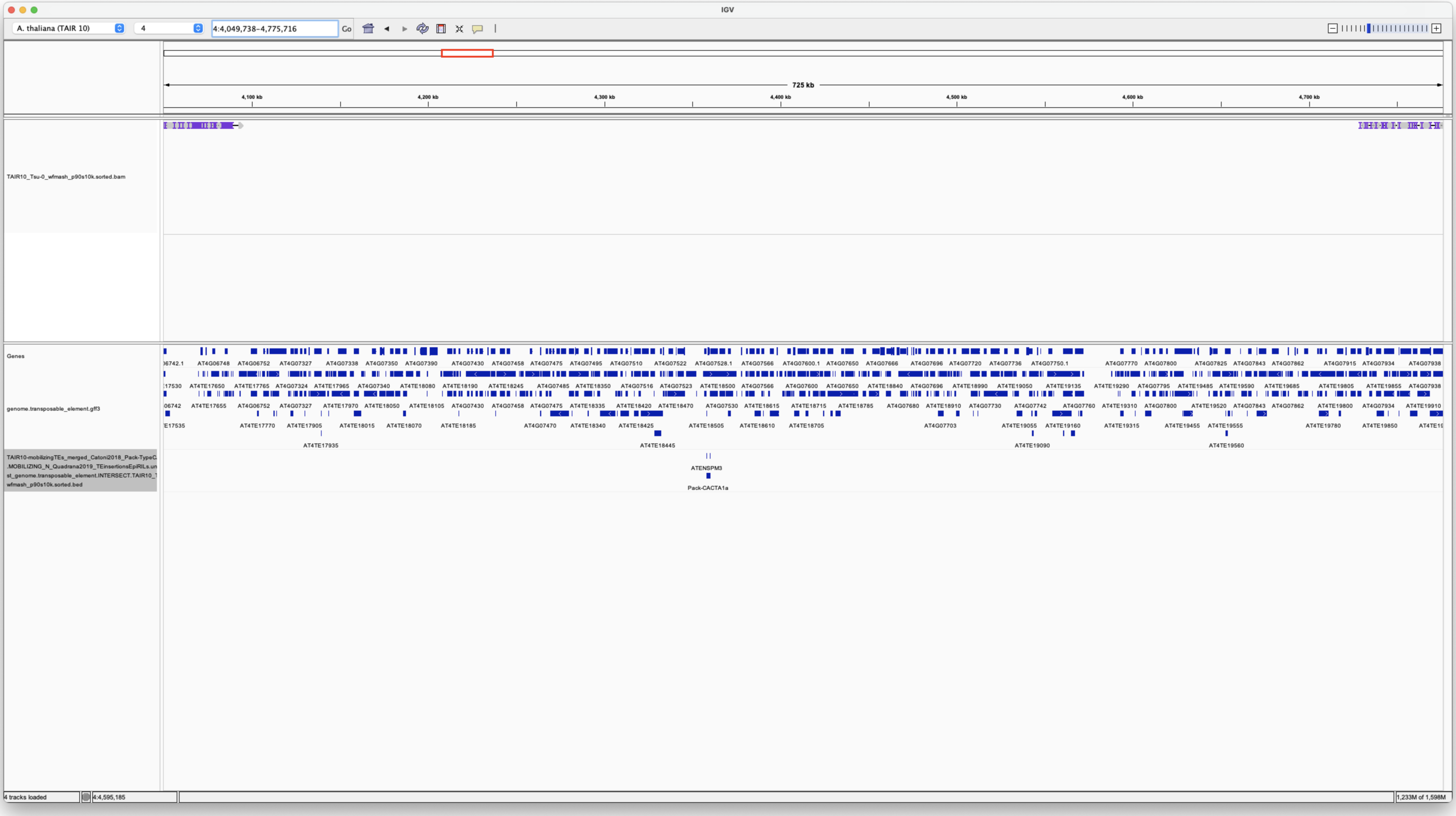

No Tsu-0

**Chr5 13960645 13968612 VANDAL21**

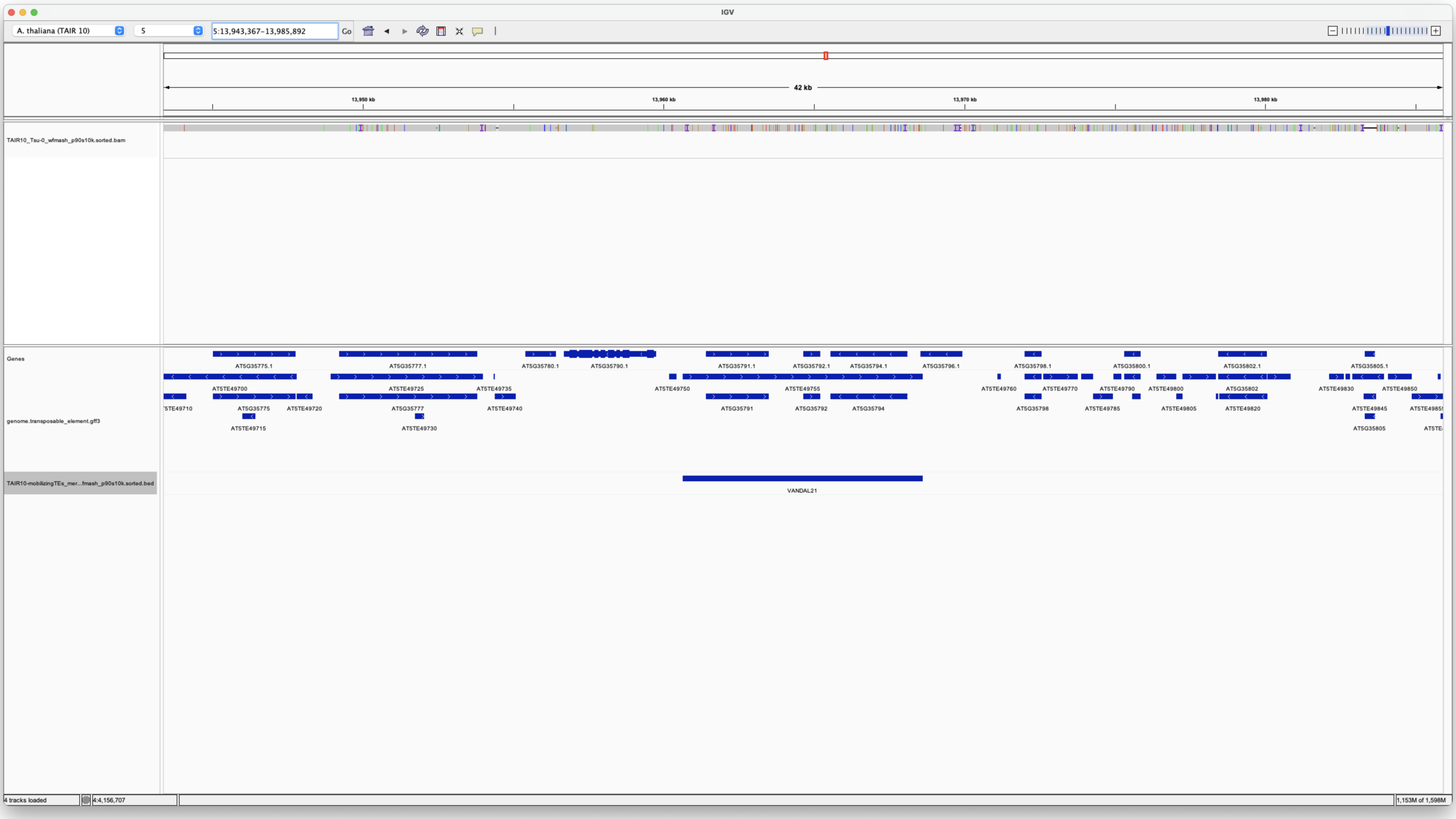

YES Tsu-0

**Chr5 18142155 18146898 ATCOPIA21**

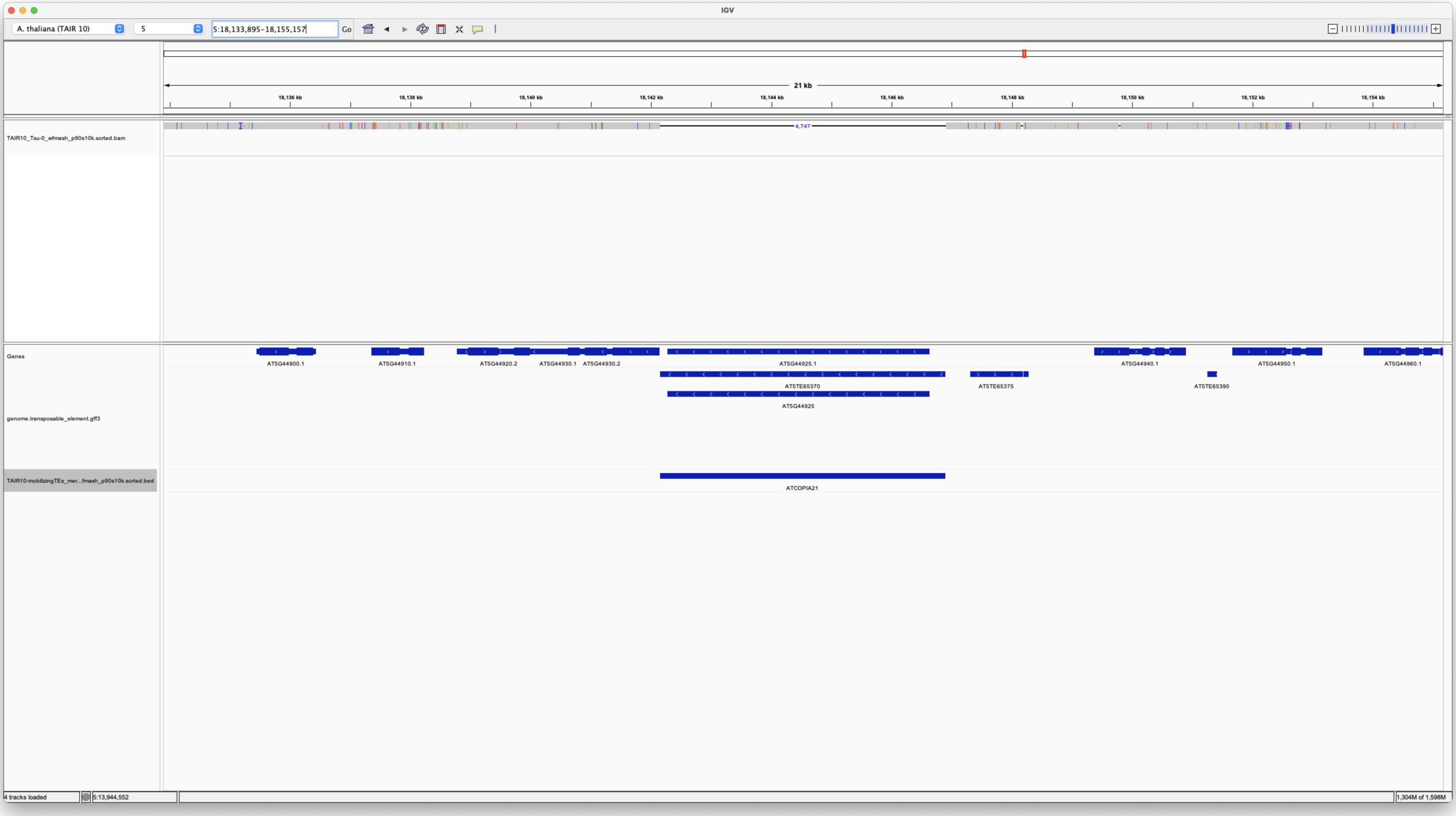

No Tsu-0

**Chr5 18490605 18495234 ATCOPIA31**

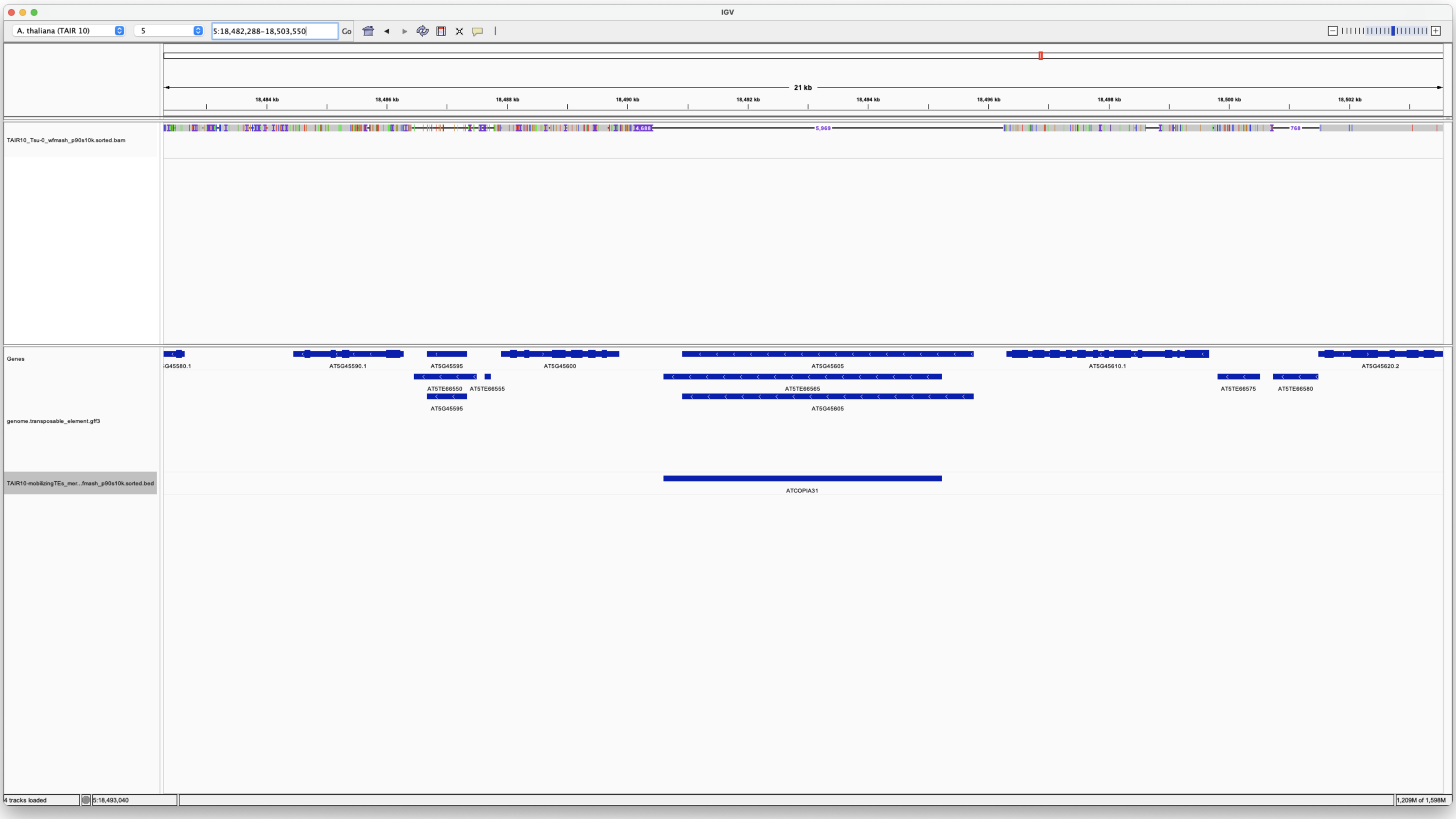


No Tsu-0

**Chr5 5629977 5635310 ATCOPIA93**

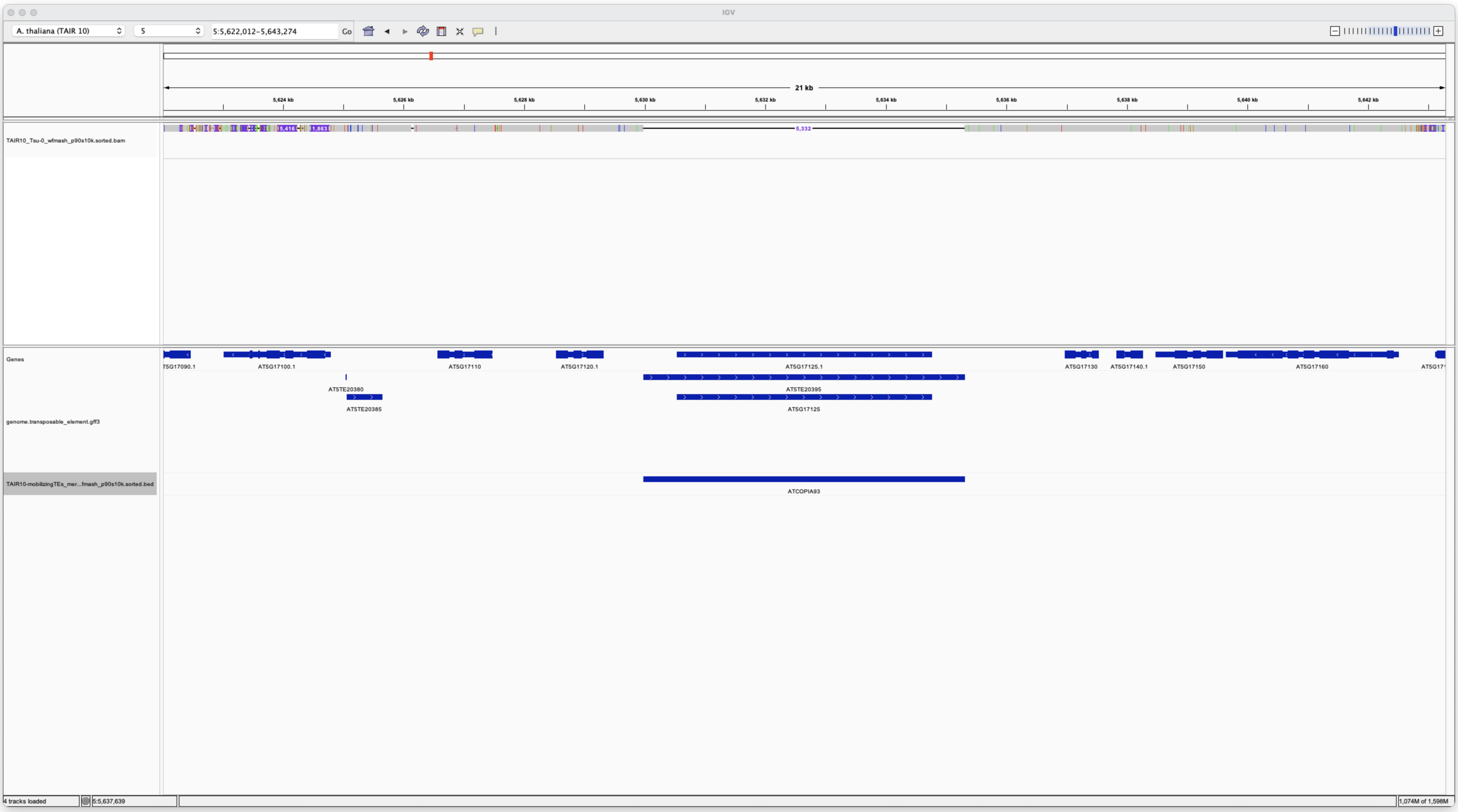


No Tsu-0

**Chr5 9272139 9274532 ATCOPIA63**

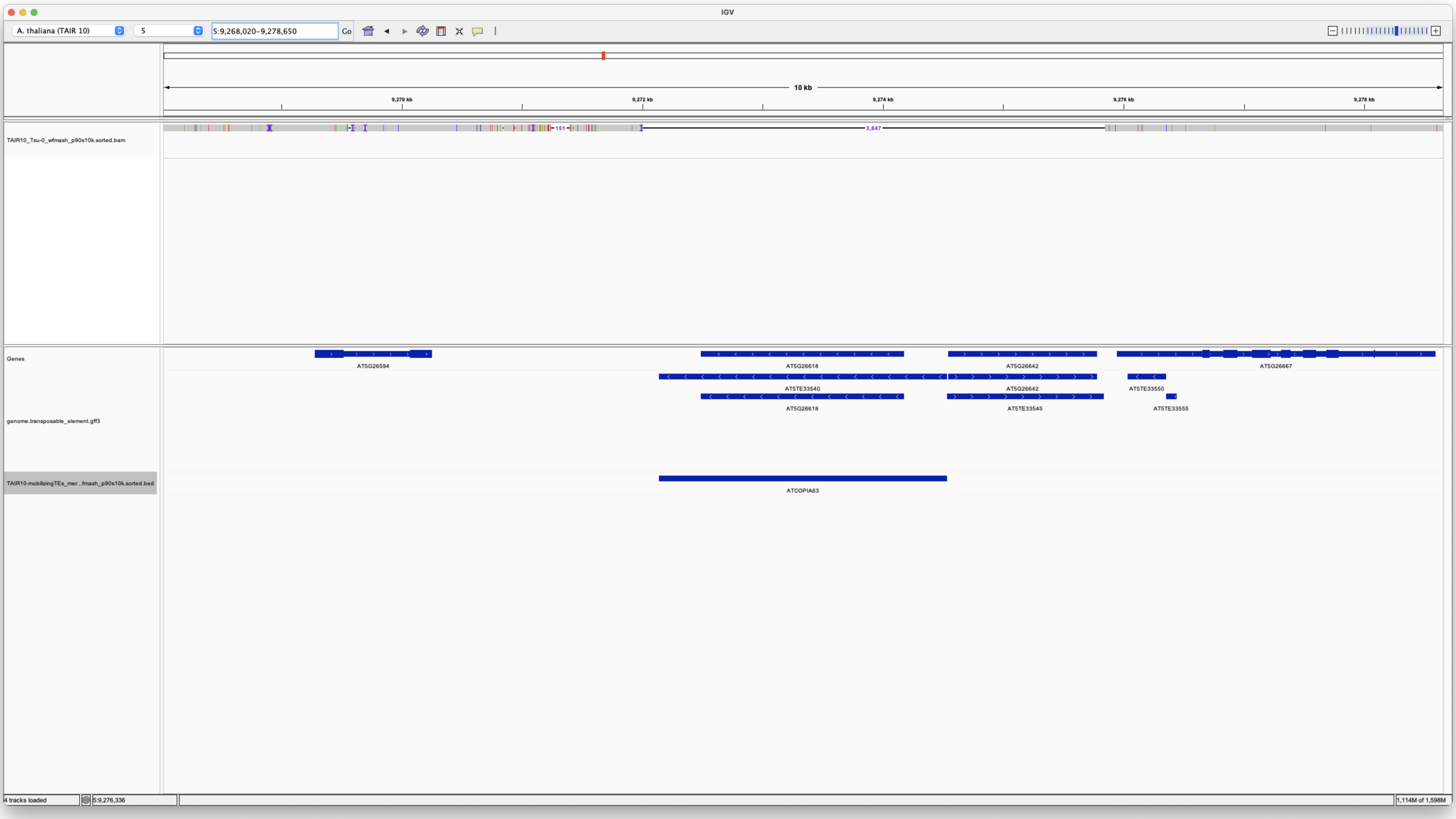


No Tsu-0

---

#### Tsukahara

##### Chr1 13818975 13823475 AT1G36590

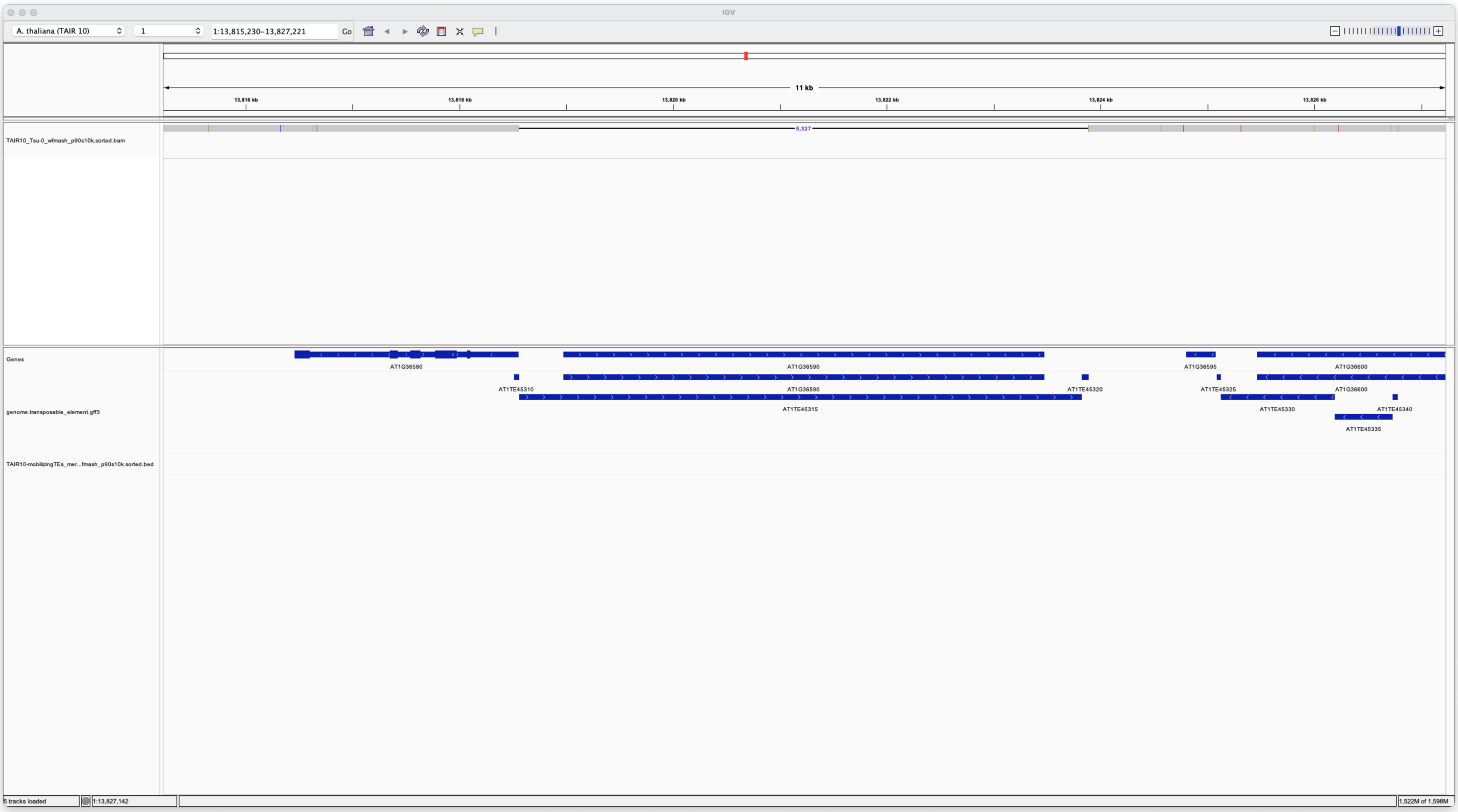


No Tsu-0

##### Chr1 29377751 29380183 AT1G78095

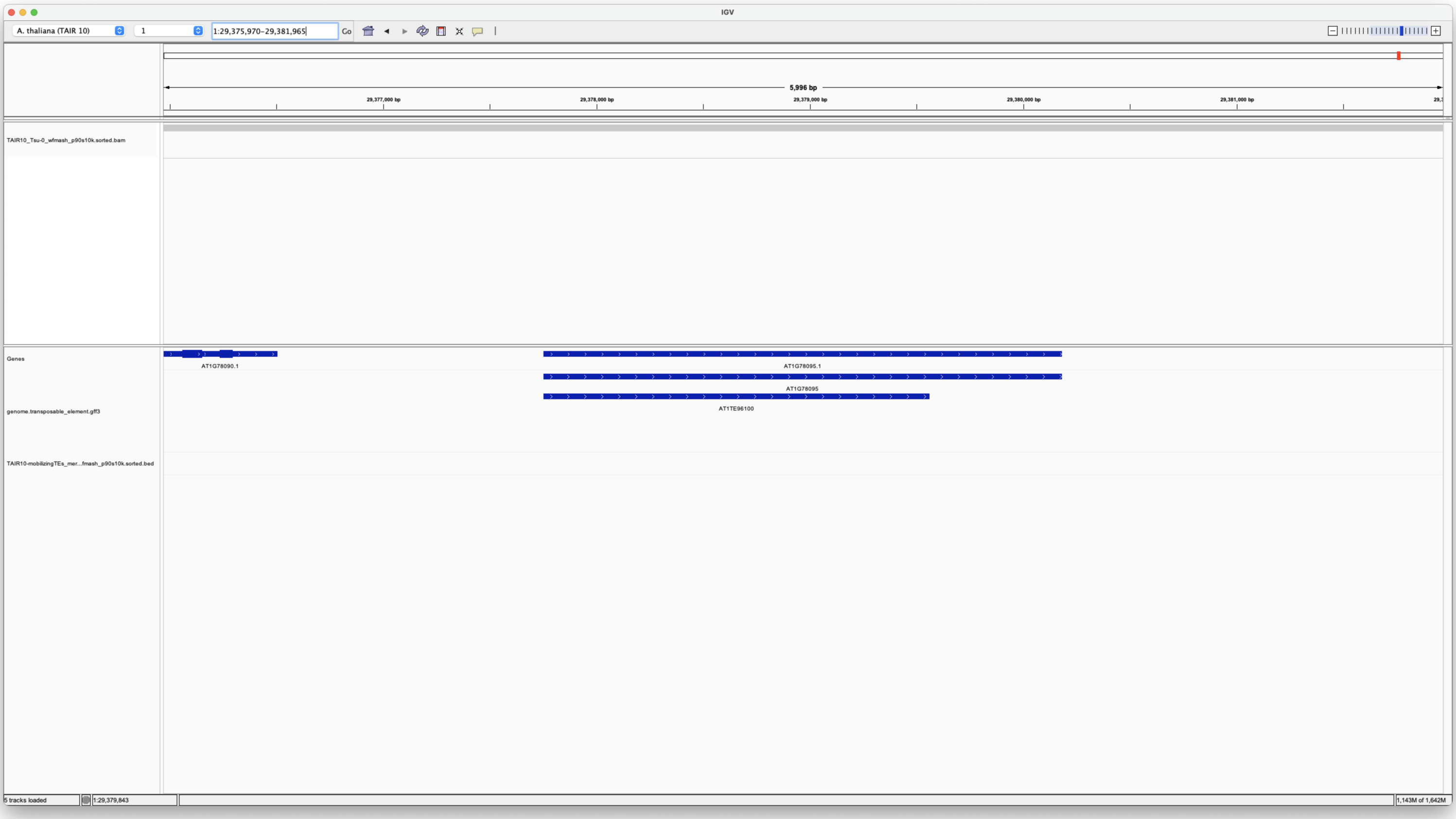


YES Tsu-0

##### Chr3 3807726 3812227 AT3G11970

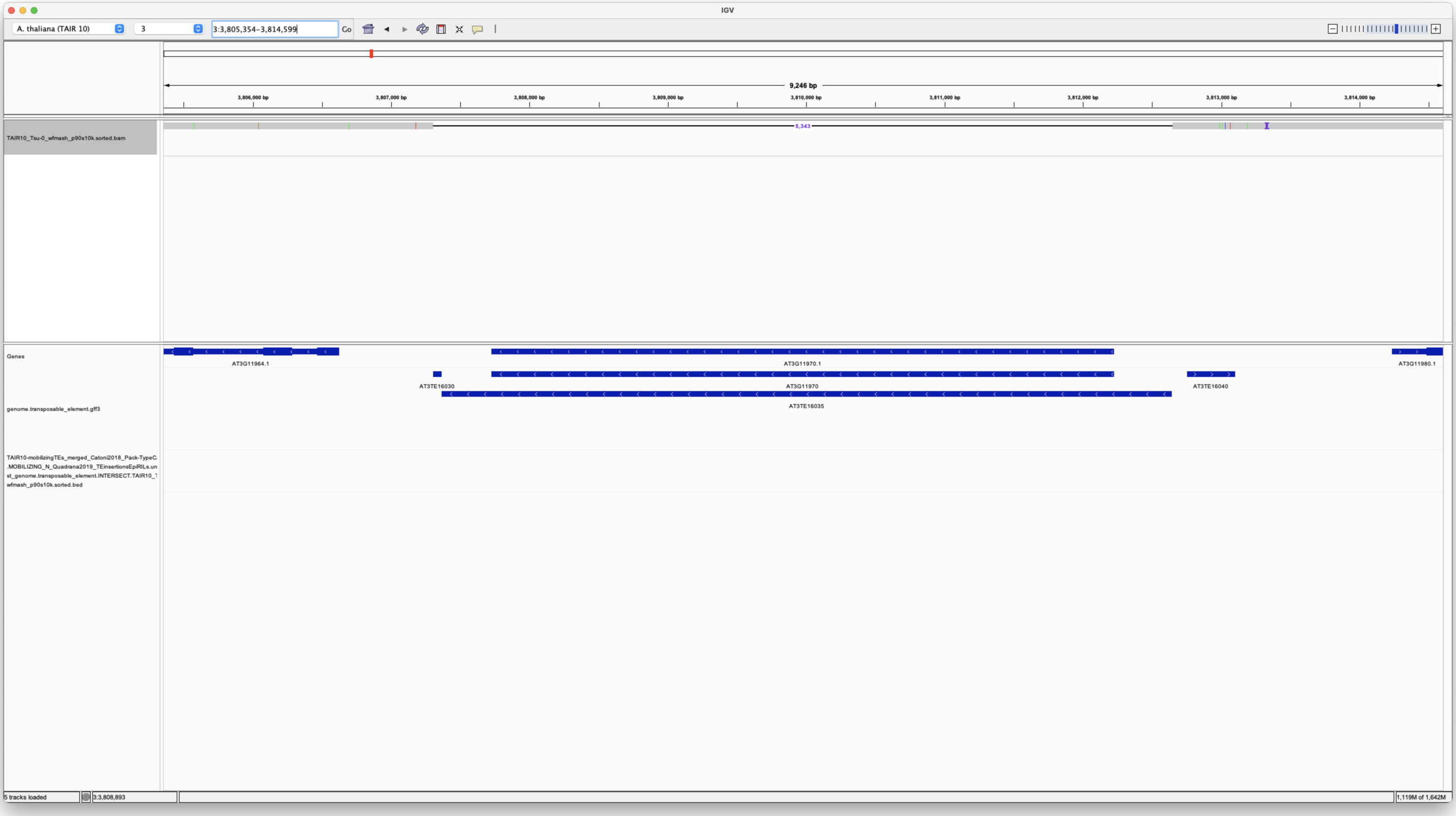


No Tsu-0

##### Chr4 5542430 5544862 AT4G08680

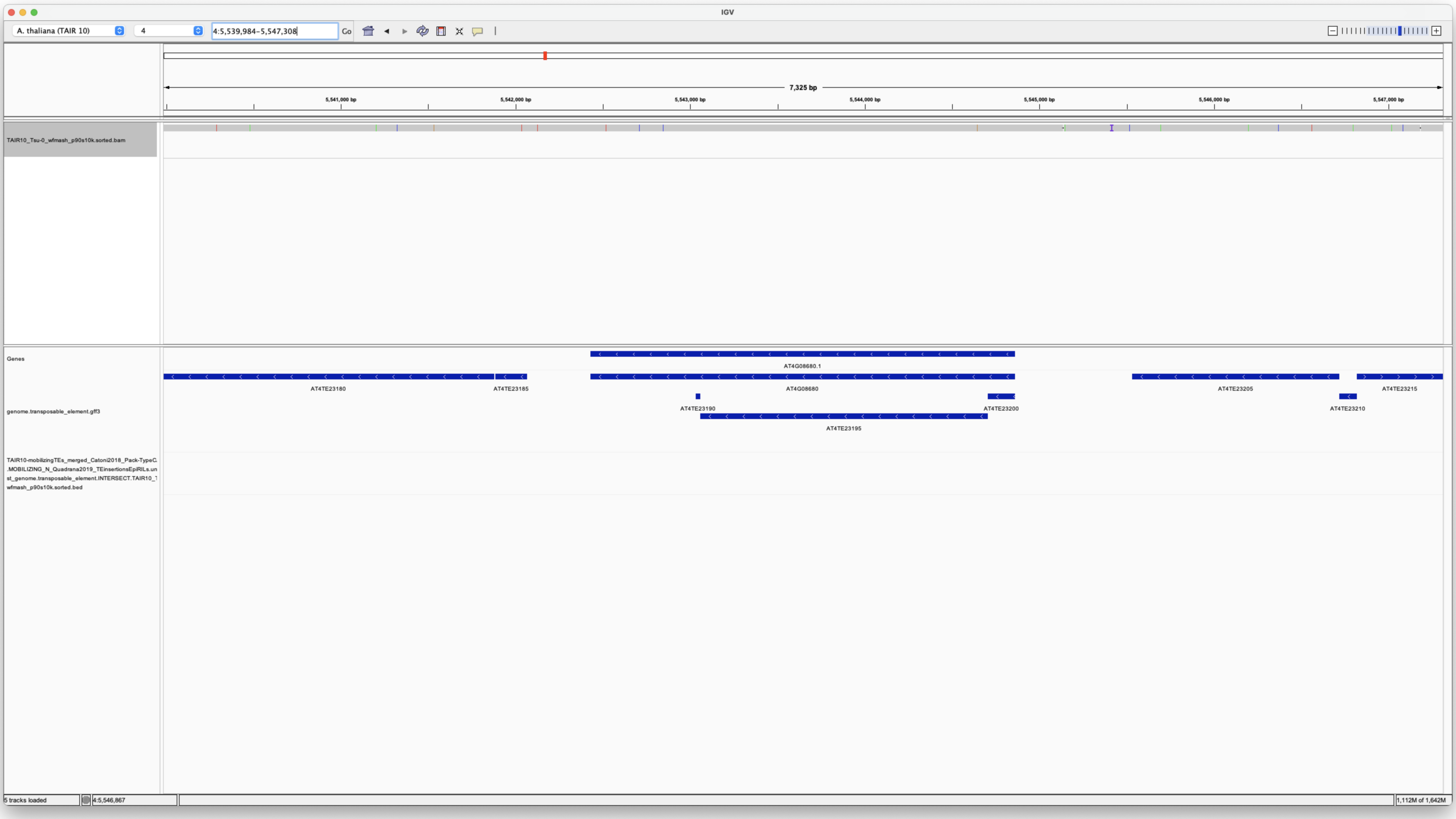


YES Tsu-0

---

#### Miura

##### Chr1 12942824 12947010 AT1TE42210

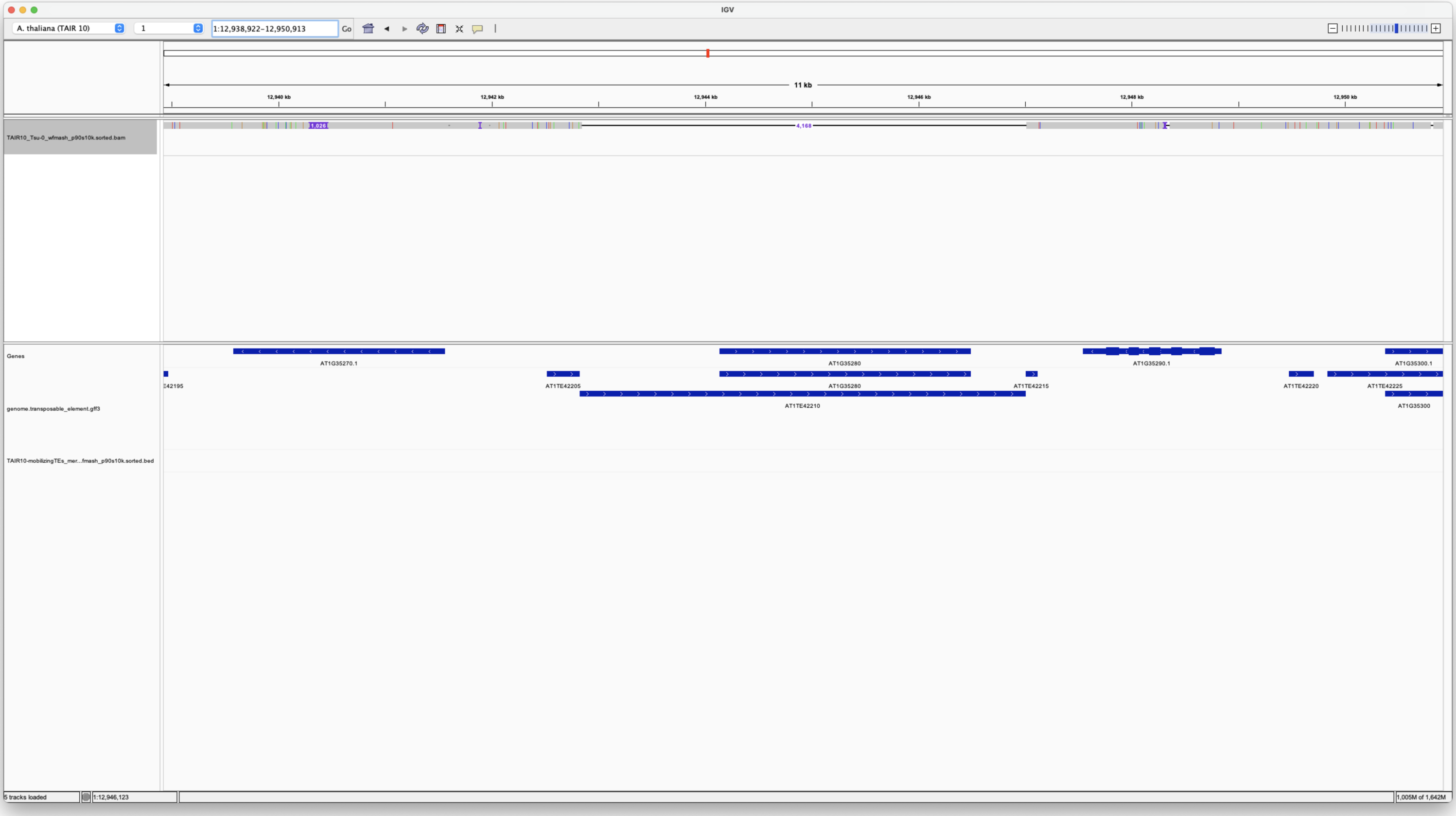


No Tsu-0

---

### ATMU5_Comp

overlaps with ATMU1 (AT4G08680)

##### Quadrana et al., 2018

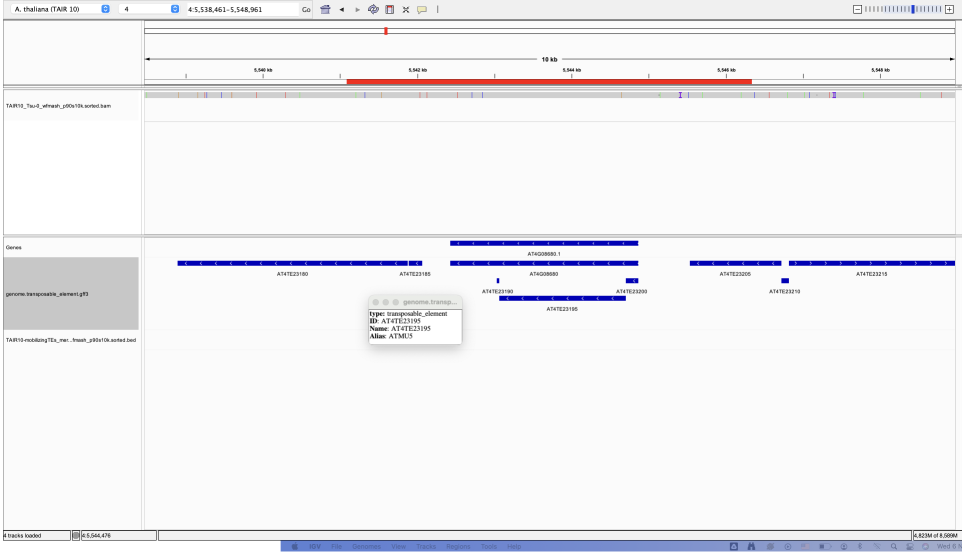

A. thaliana (TAIR 10)
4
4:5,538,461-5,548,961
Go
10 kb
5,540 kb
5,542 kb
5,544 kb
5,546 kb
5,548 kb
TAIR10_Tsu-0_wfmash_p90s10k.sorted.bam
Genes
genome.transposable_element.gff3
TAIR10-mobilizingTEs_mer...fmash_p90s10k.sorted.bed
AT4G08680.1
AT4TE23180
AT4TE23185
AT4G08680
AT4TE23205
AT4TE23215
AT4TE23190
AT4TE23200
AT4TE23210
AT4TE23195
type: transposable_element
ID: AT4TE23195
Name: AT4TE23195
Alias: ATMU5
4 tracks loaded
4:5,544,476
4,823M of 8,589M


Yes Tsu-0
