## Additional File 4 for "Long-read detection of transposable element mobilization in the soma of hypomethylated *Arabidopsis thaliana* individuals": Synteny-check_Tsu-0-TEs-in-Col-0.pdf

#### Chr1 11941106 11946436 ATCOPIA93_Evade

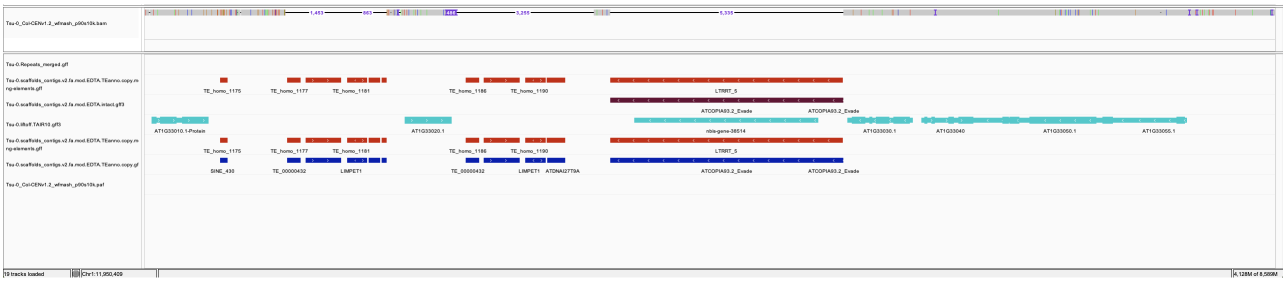

*NON-REF*

### VANDAL21 Chr2:8027483

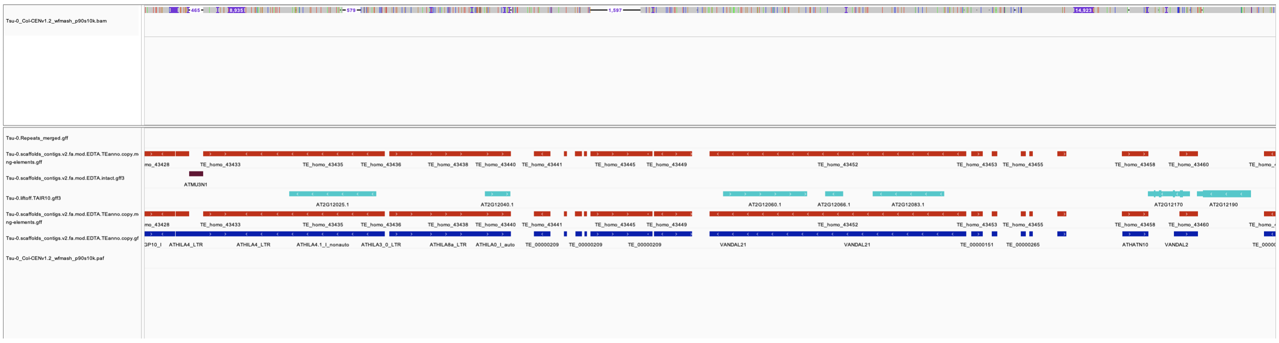

*REF*

### ChimericCACTA Chr2:8256939

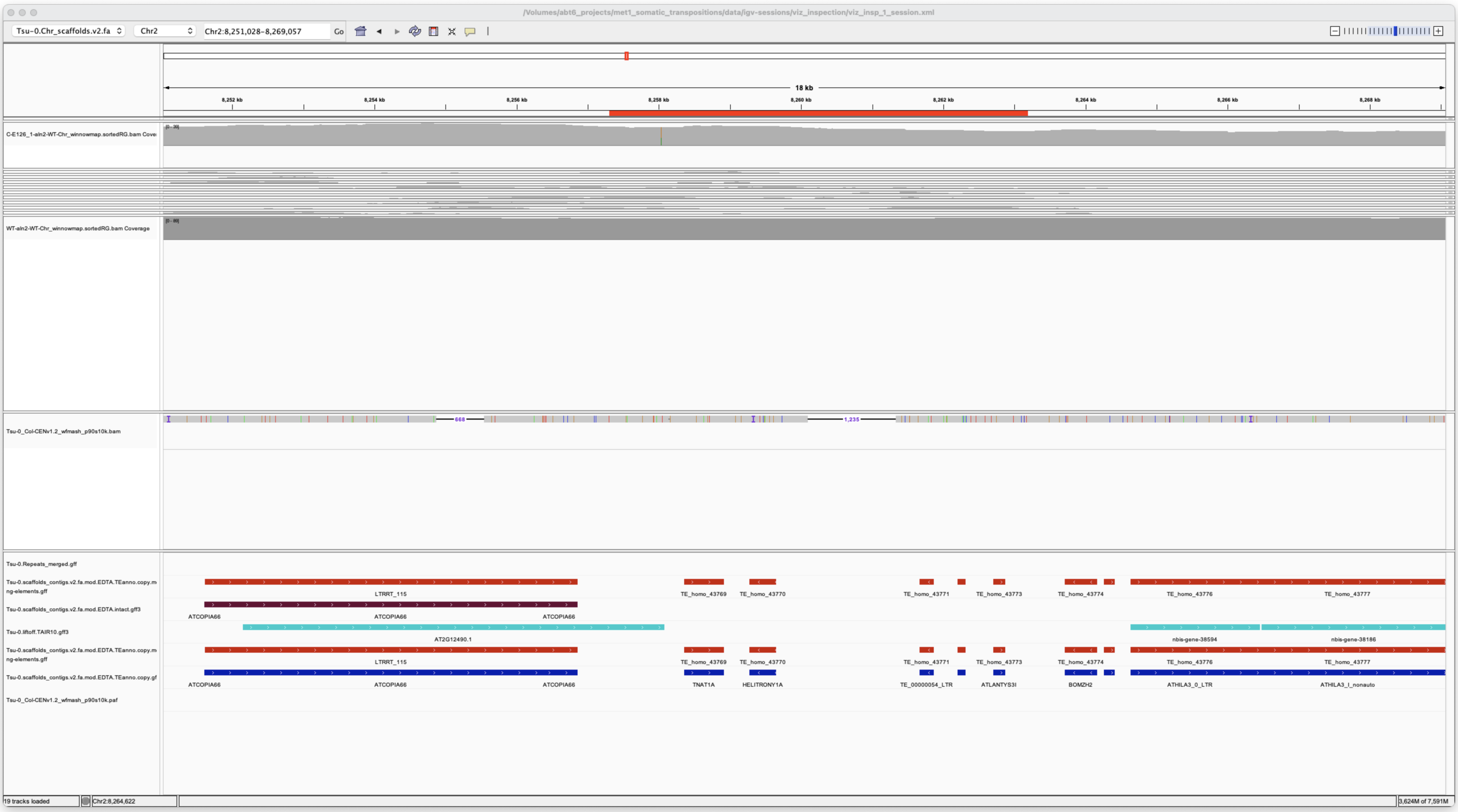

---

*REF*

with deletion in Col-0

### ATENSPM2 Chr2:8337982

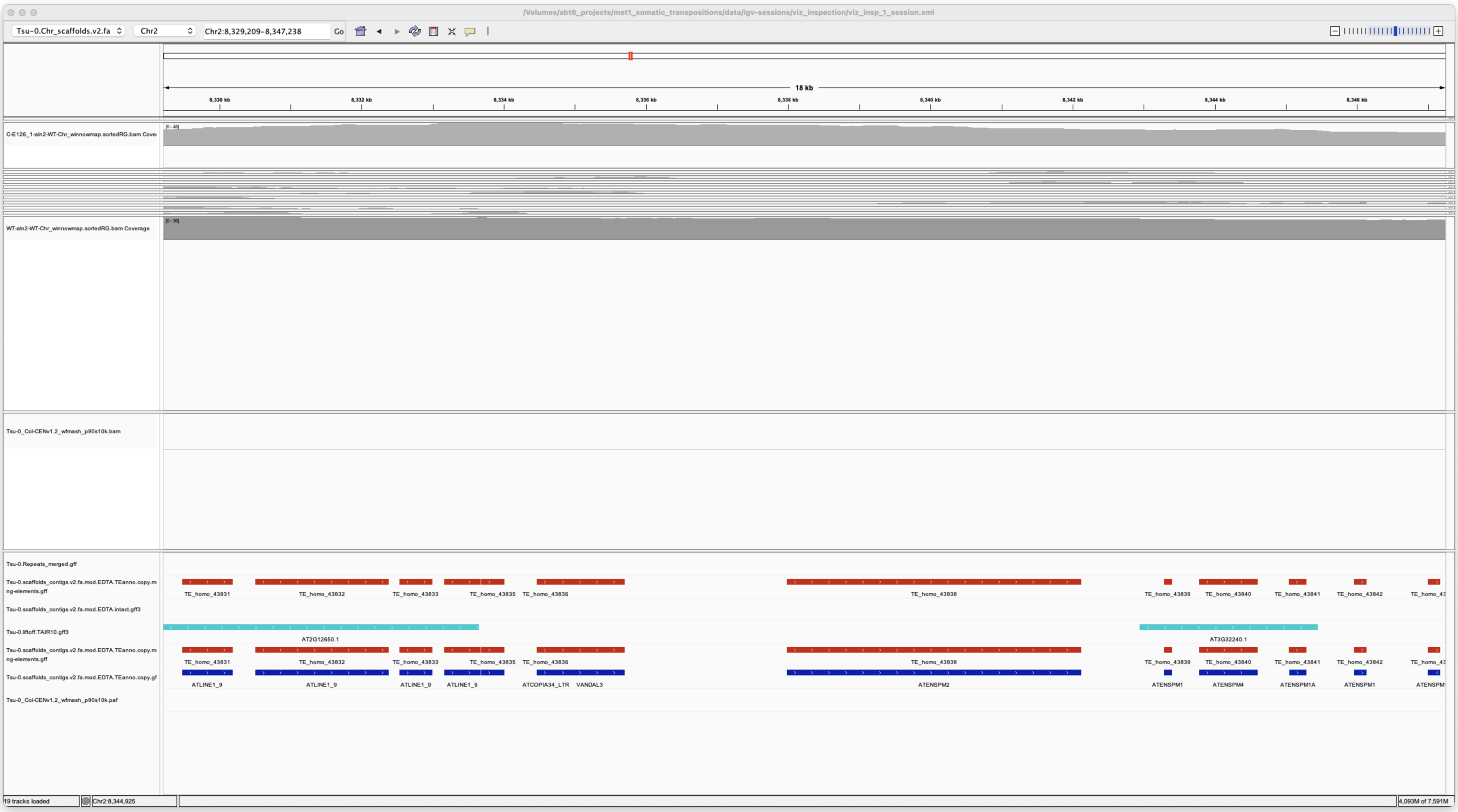


***NON-REF***

**VANDAL6 Chr3:16344522**

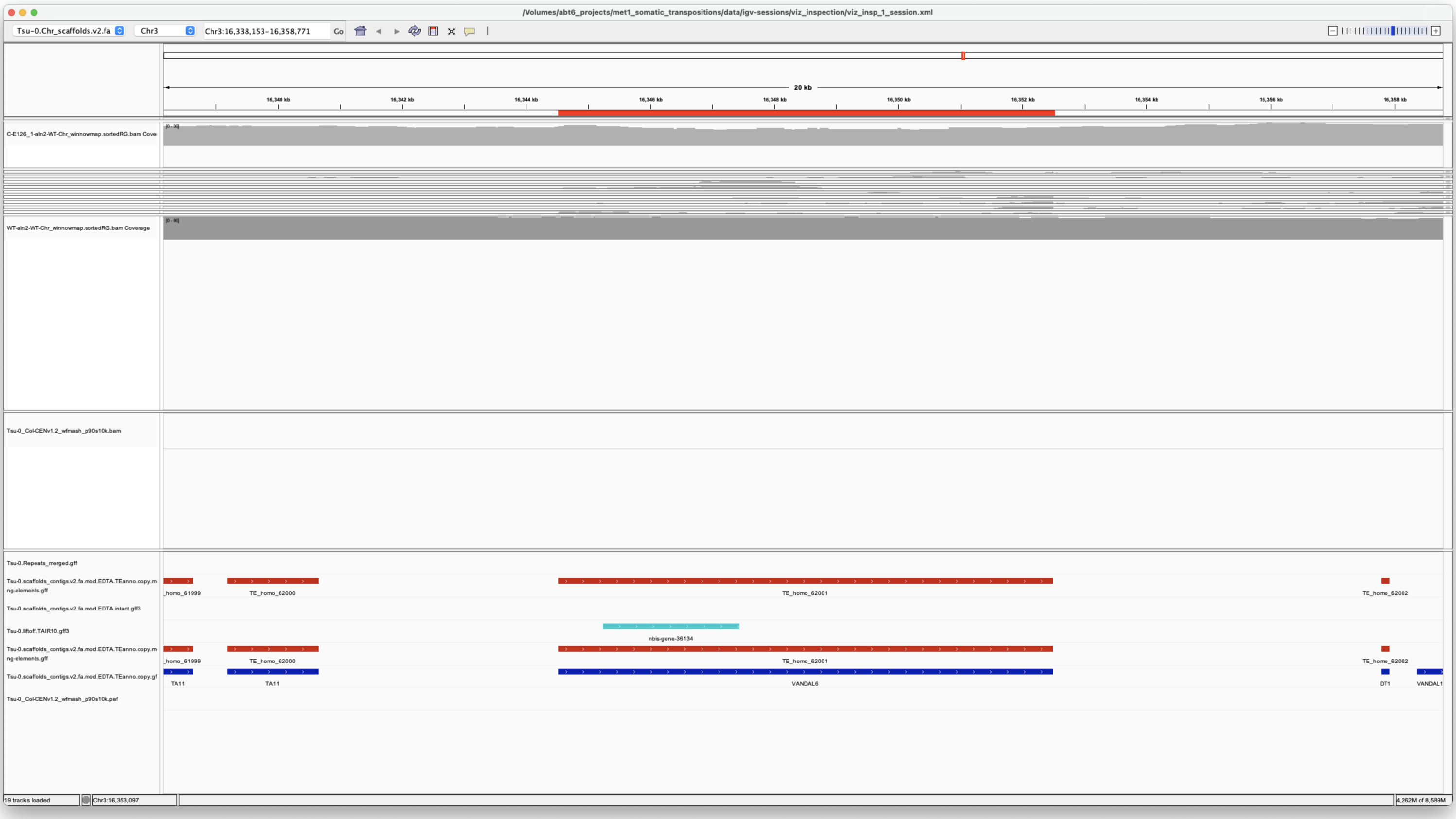

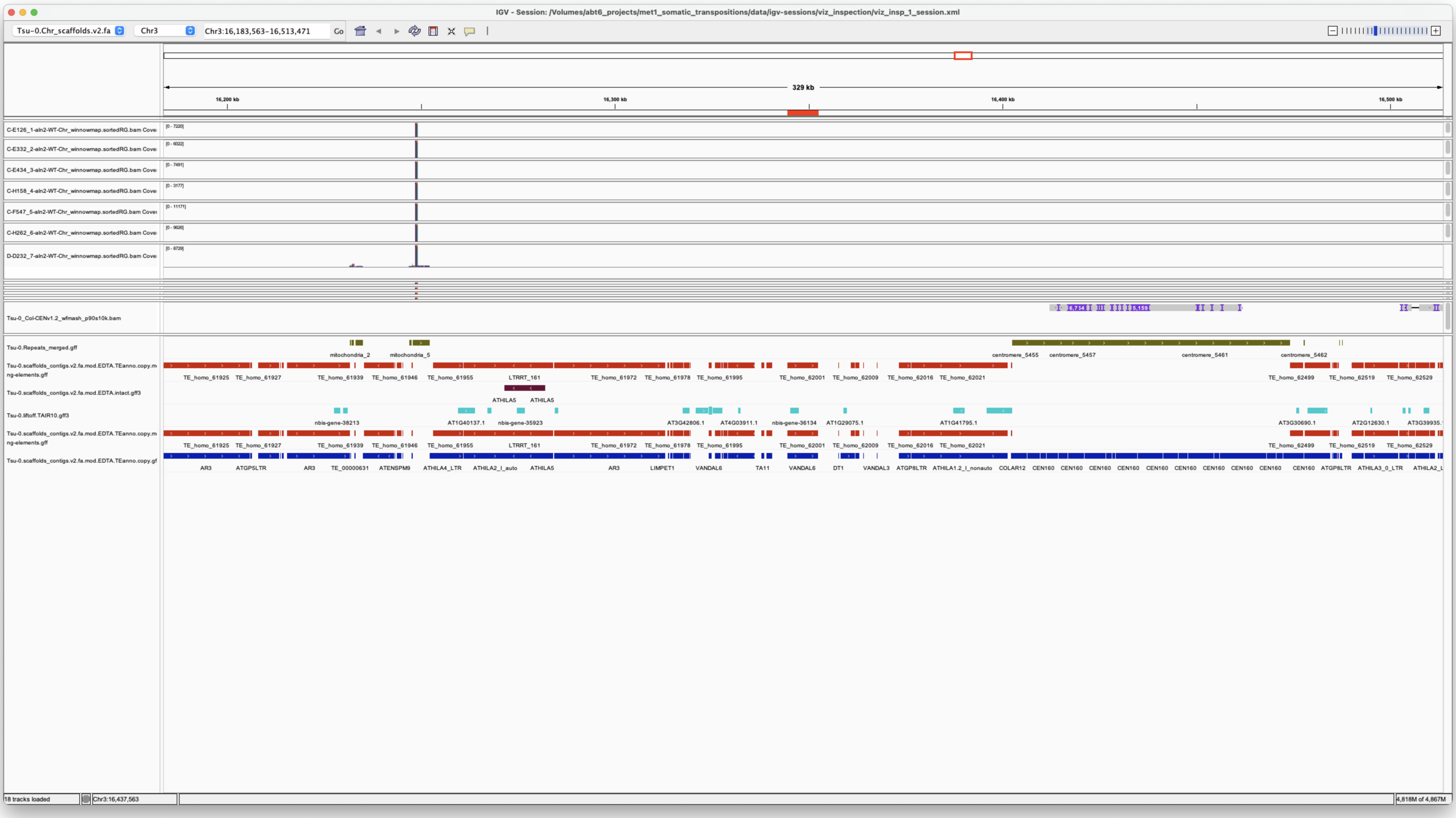

*NON-REF*

**VANDAL6 Chr3:20158137**

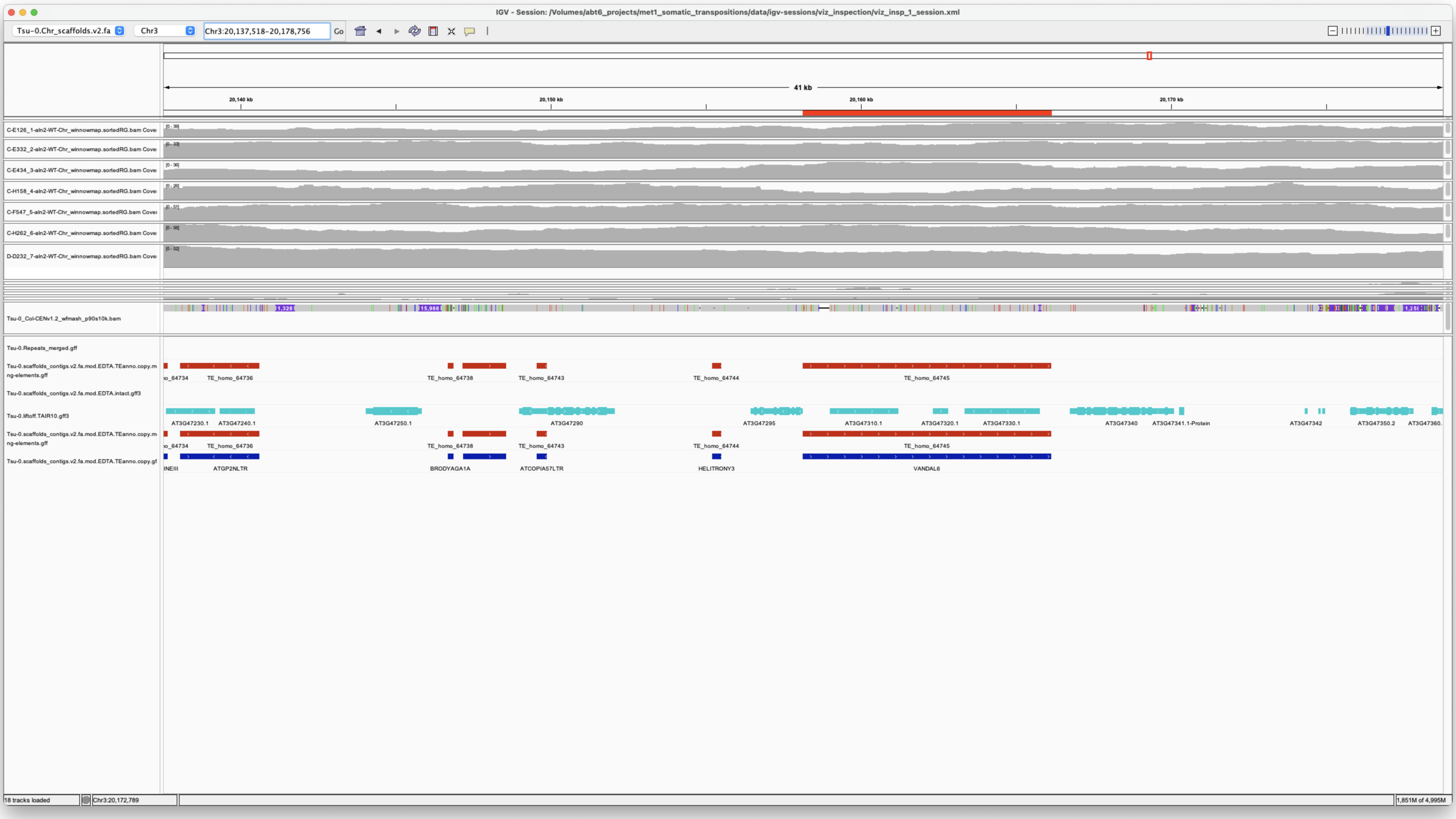


*REF*

**VANDAL21MANUAL Chr4:9036240**

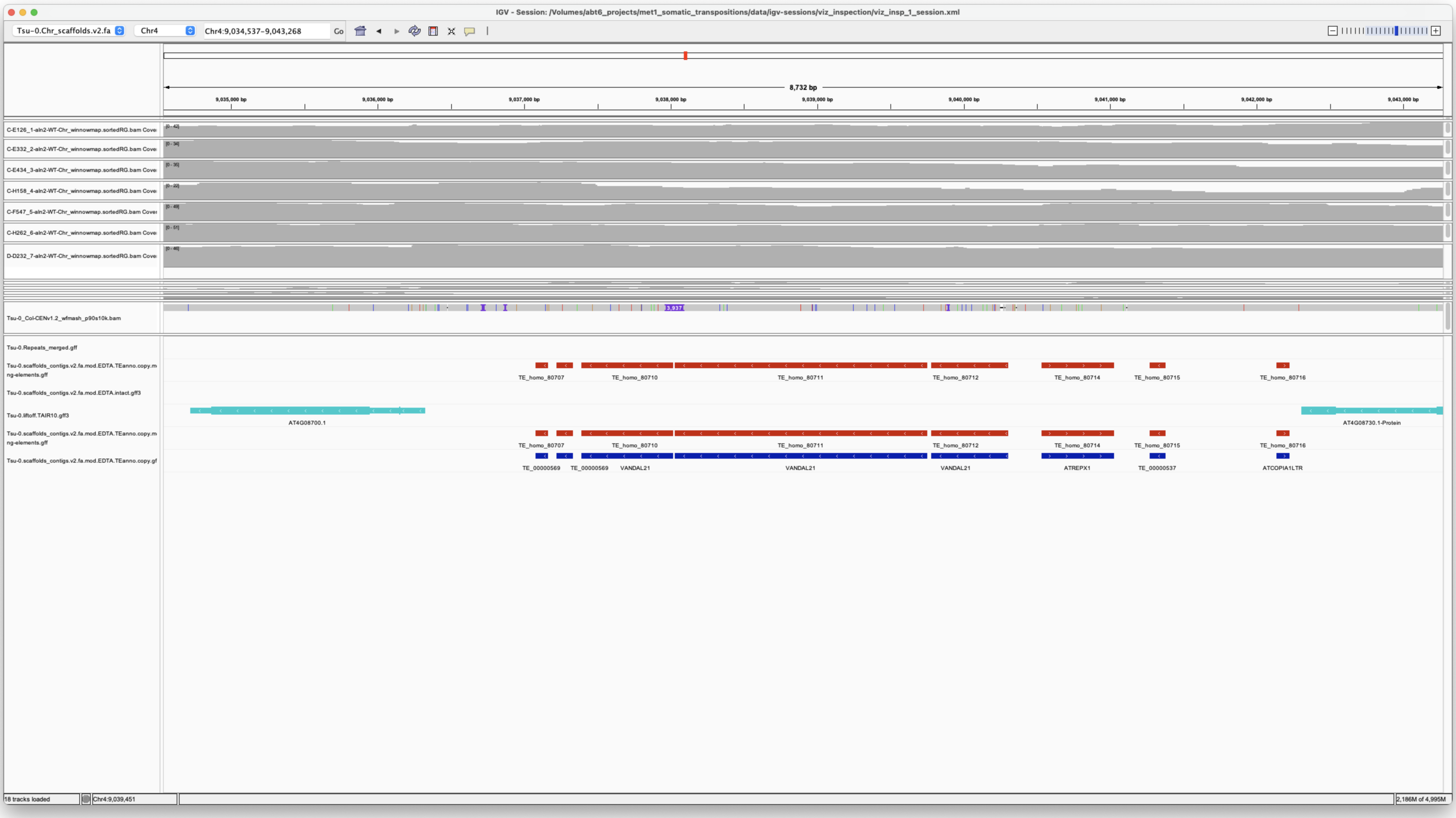


*REF*

Col-0 has an insertion

**VANDAL21 Chr5:19152829**

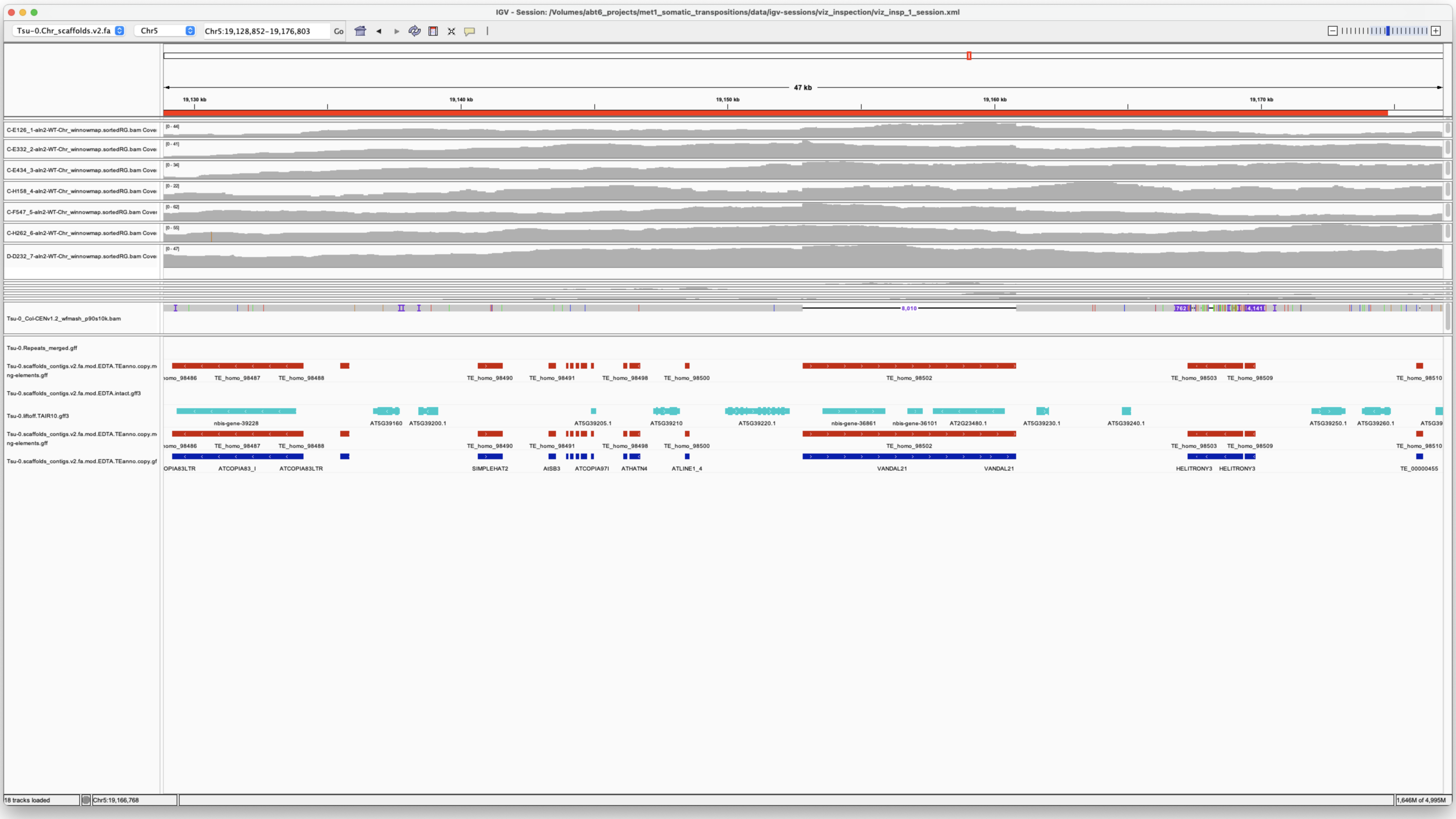

*NON-REF*

**ATCOPIA93_Evade Chr5:21419693**

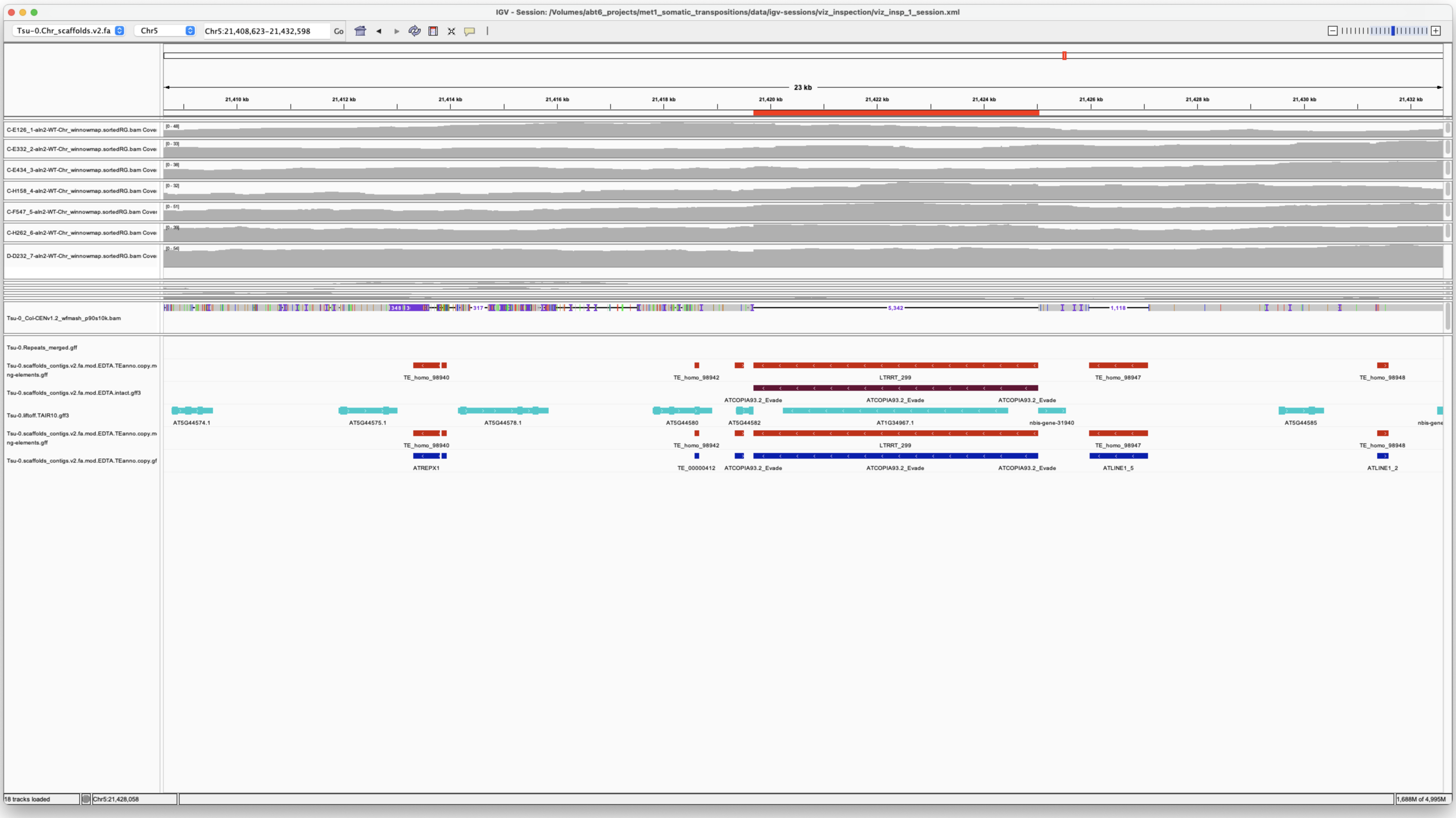

*NON-REF*

**ChimericCACTA Chr5:8256939**

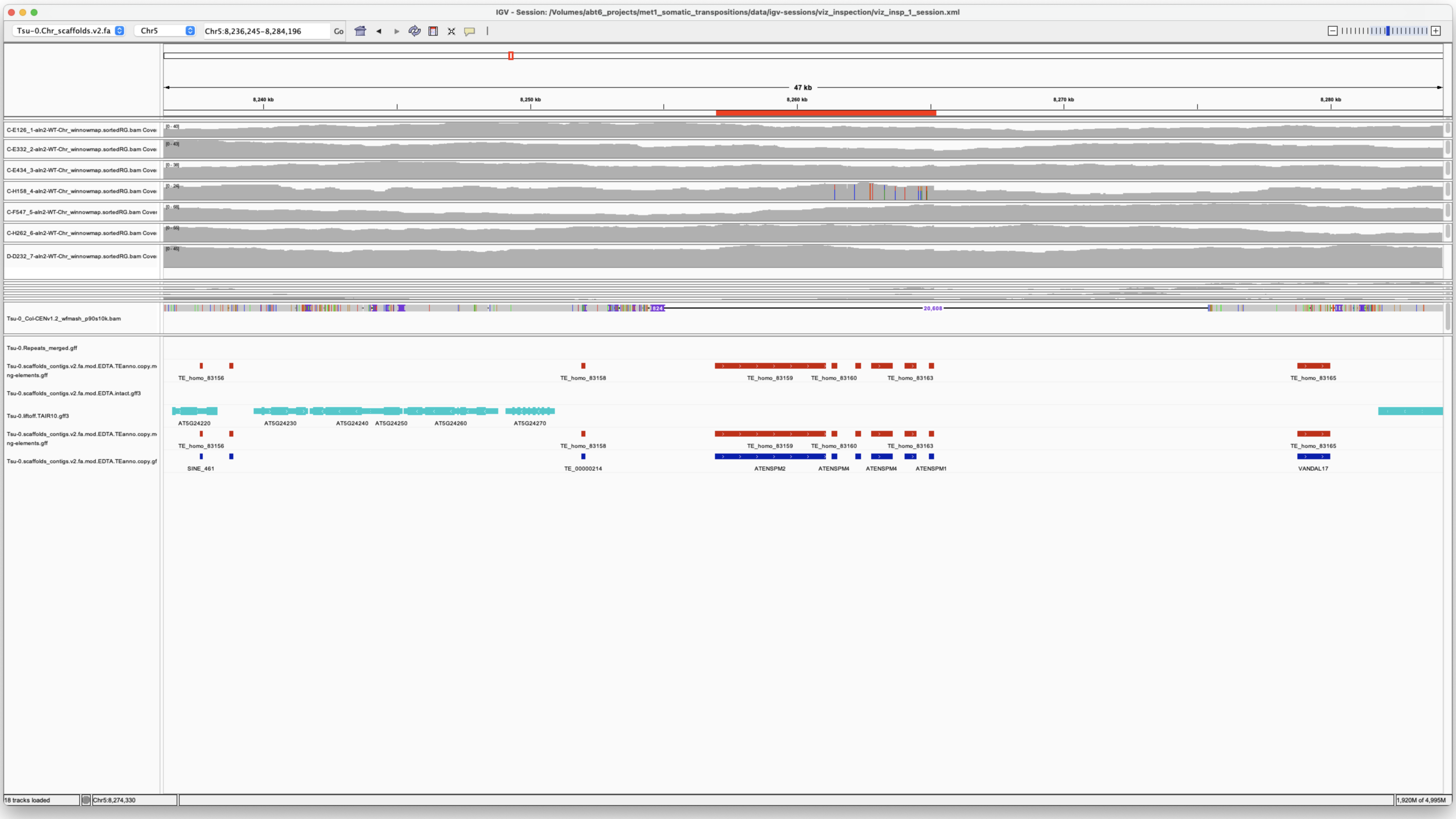


*NON-REF*

**PutativePackTypeCACTAMuDR Chr5:875414**

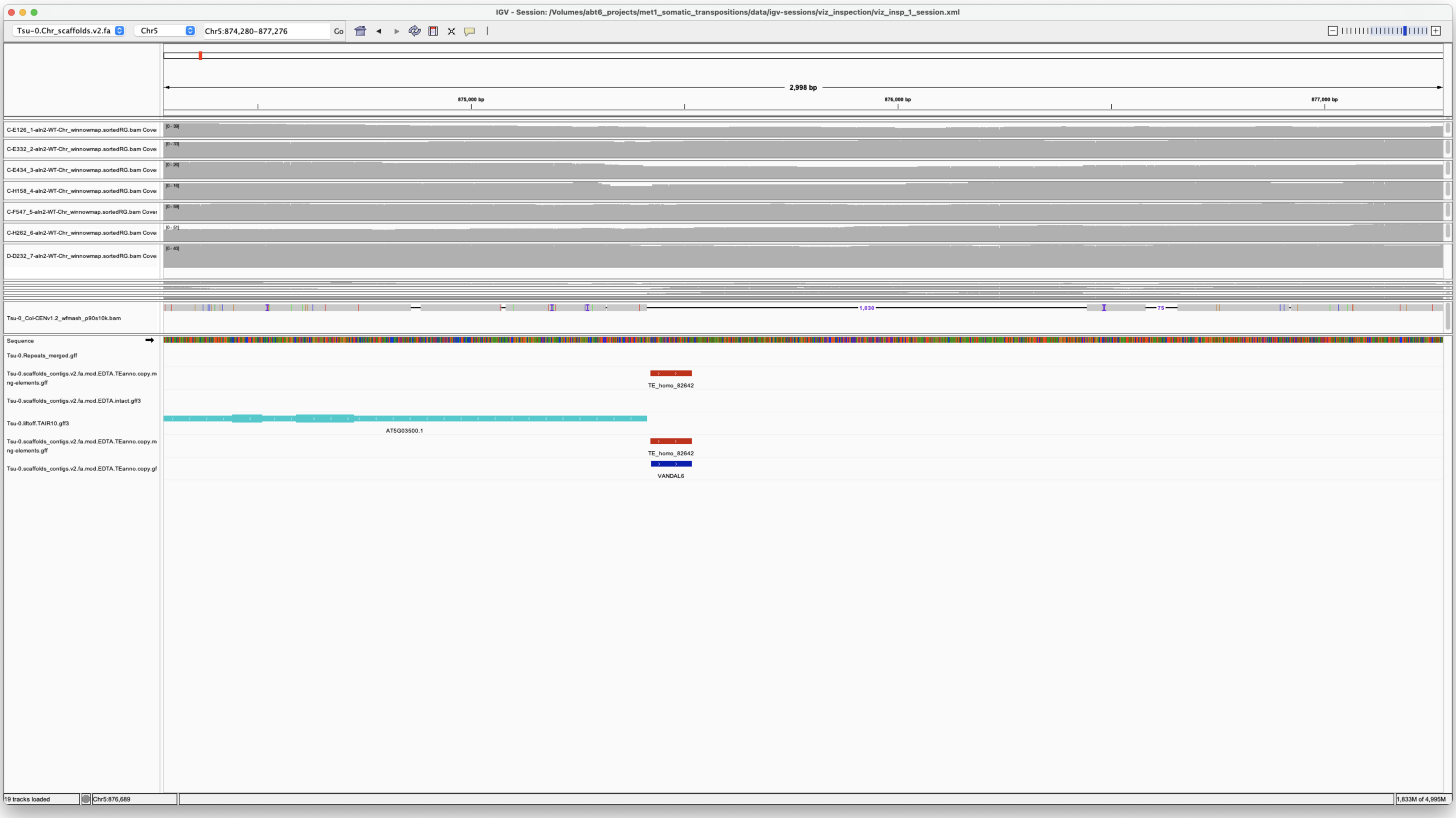


*NON-REF*

**Chr5 24560934 24568727 ChimericCACTA ChimericCACTA-Chr5:24560934**

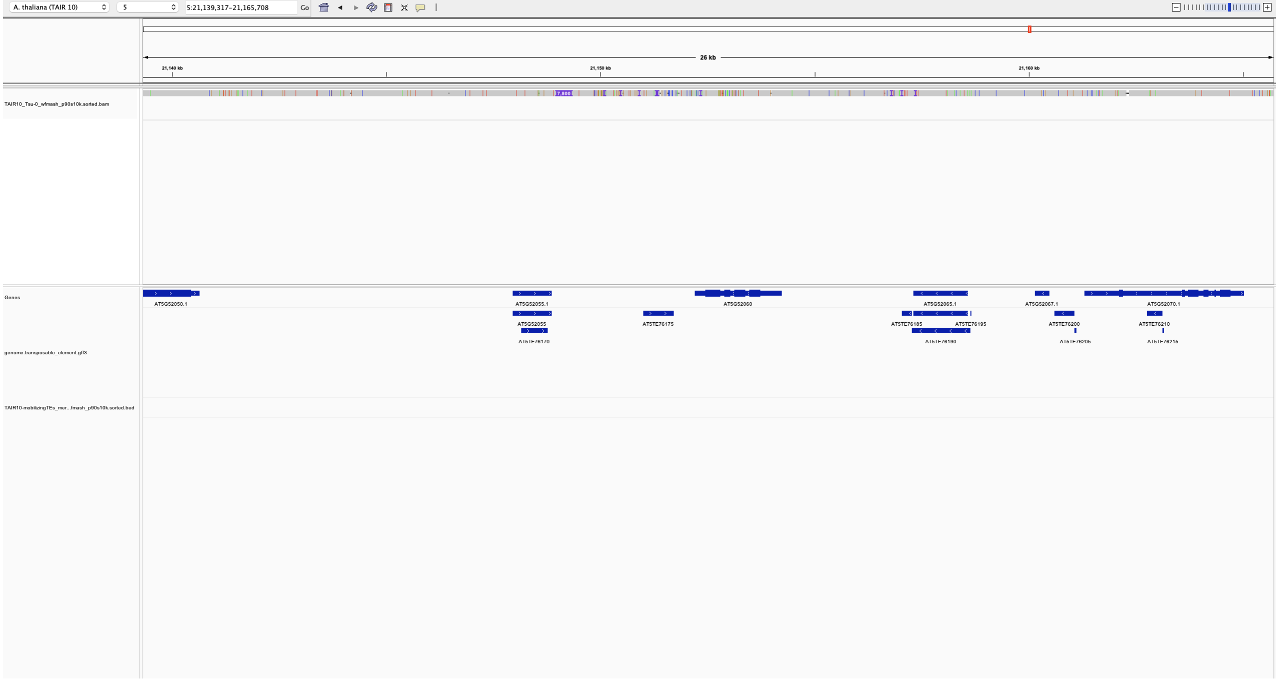
A. thaliana (TAIR 10)
5
5:21,139,317-21,165,708
Go
26 kb
21,140 kb
21,150 kb
21,160 kb
TAIR10_Tsu-0_wfmash_p90s10k.sorted.bam
Genes
genome.transposable_element.gff3
TAIR10-mobilizingTEs_mer...fmash_p90s10k.sorted.bed
AT5G52050.1
AT5G52055.1
AT5G52060
AT5G52065.1
AT5G52067.1
AT5G52070.1
AT5G52055
AT5TE76175
AT5TE76185
AT5TE76195
AT5TE76200
AT5TE76210
AT5TE76170
AT5TE76190
AT5TE76205
AT5TE76215

**
